# dsSurvival: Privacy preserving survival models for federated individual patient meta-analysis in DataSHIELD

**DOI:** 10.1101/2022.01.04.471418

**Authors:** Soumya Banerjee, Ghislain N. Sofack, Thodoris Papakonstantinou, Demetris Avraam, Paul Burton, Daniela Zöller, Tom R.P. Bishop

## Abstract

**Objective:** Achieving sufficient statistical power in a survival analysis usually requires large amounts of data from different sites. Sensitivity of individual-level data, ethical and practical considerations regarding data sharing across institutions could be a potential challenge for achieving this added power. Hence we implemented a federated meta-analysis approach of survival models in DataSHIELD, where only anonymous aggregated data are shared across institutions, while simultaneously allowing for exploratory, interactive modelling.

In this case, meta-analysis techniques to combine analysis results from each site are a solution, but an analytic workflow involving local analysis undertaken at individual studies hinders exploration. Thus, the aim is to provide a framework for performing meta-analysis of Cox regression models across institutions without manual analysis steps for the data providers.

**Results:** We introduce a package (dsSurvival) which allows privacy preserving meta-analysis of survival models, including the calculation of hazard ratios. Our tool can be of great use in biomedical research where there is a need for building survival models and there are privacy concerns about sharing data.

## Introduction

Survival models are widely used in biomedical research for analyzing survival data [1]. These models help researchers compare the effect of exposures on mortality or other outcomes of interest [2]. The Cox proportional hazards model [3] is one of the most popular survival analysis models and was primarily developed to determine the importance of predictors in survival, by using covariate information to make individual predictions [4].

Achieving sufficient power in survival analysis usually requires large amounts of data from several sites or institutions. Multi-site analysis across studies with different population characteristics help us understand how diseases affect different populations and what it is about these populations that cause these differences. However, the number of cases at a single site is often rather small, making statistical analysis challenging. In addition, due to the sensitivity of individual-level biomedical data, ethical and practical considerations related to data transmission, and institutional policies, individual-level data cannot be shared [5].

In consortia, this issue is often addressed by manual analysis in each site, followed by a manual meta-analysis of the analysis results from the individual sites. This process is very time-consuming and error-prone, making exploratory analysis (e.g., for understanding different effect patterns in each site) impractical. As an alternative, the DataSHIELD framework can be used.

DataSHIELD is a framework that enables the remote and privacy preserving analysis of sensitive research data [6]. The framework is based on the programming language R [7] [8]. In each site, specifically requested aggregated anonymous analysis results can be requested, which are then combined in a central analysis server. The requirement is that the analysis be privacy preserving and be conducted across globally distributed cohorts.

We have implemented a meta-analysis approach based on the Cox-model in DataSHIELD using individual patient data that are distributed across several sites, without moving those data to a central site i.e. the individual-level data remain within each site and only non-disclosive aggregated data are shared. Our software package for DataSHIELD allows building of survival models and analyzing results in a federated privacy preserving fashion.

Remote federated meta-analysis allows the analysis to come to the data and enables multiple research groups to collate their data [7] [8]. This is an alternative to literature based meta-analysis since study variables and outcomes can be harmonised [9]. Our package offers considerable advantages over: (1) literature based meta-analysis, which suffers from publication bias as well as restricting the analytic end-points you may wish to use; (2) central pooling of data, which provides important governance challenges and can engender privacy risks; and (3) asking researchers in each location to do local analyses based on a shared analysis plan, which all too often demands numerous emails, with repeated reminders, to disseminate analytic protocols and return results for meta-analysis, which is typically time-consuming and can be error prone. Our tool can be of great use in domains where there is a need for building survival models and there are privacy concerns about sharing data.

## Main text

### Basics of survival analysis

Survival analysis can be used to analyze clinical data if there are records of patient mortality and time to event data. The key quantity is a survival function:

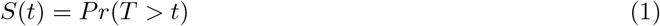

where *S*(*t*) is the survival function, *t* is the current time, and *T* is a random variable denoting time of death. Pr() is the probability that the time of death is greater than time t i.e. the probability of surviving till time *t*.

The instantaneous hazard (*λ*(*t*)) is the probability of death occurring within time period [t, t + *δ* t] given survival till time *t*. This is related to the survival function as follows:

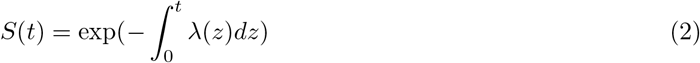

The proportional hazards model assumes that the effect of covariates is proportional to the hazard. This is modelled as follows using the hazard function *λ*(*t*):

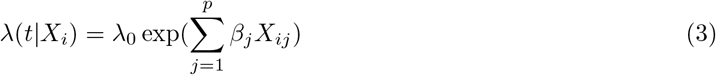

where *λ*_0_ is called the baseline hazard and *λ*(*t*) is the hazard at time *t. β* denotes the parameters and *X_ij_* is the *j*th covariate for the *i* th subject. We aim to meta-analyze these log hazard ratios.

### Implementation

DataSHIELD has a client-server architecture (Fig. 1). There are multiple servers located on separate sites and there is a single analysis client. Assign functions in DataSHIELD perform computation and ultimately create objects that persist on the servers and are not shared with the analysis client. These server-side objects can then be used for subsequent computations. Aggregate functions in DataSHIELD perform computation on each site, check for disclosure risks, and send aggregated results back to a client. The results do not persist on the servers, but can be saved on the client. This is shown in Fig. 1.

**Figure 1.**
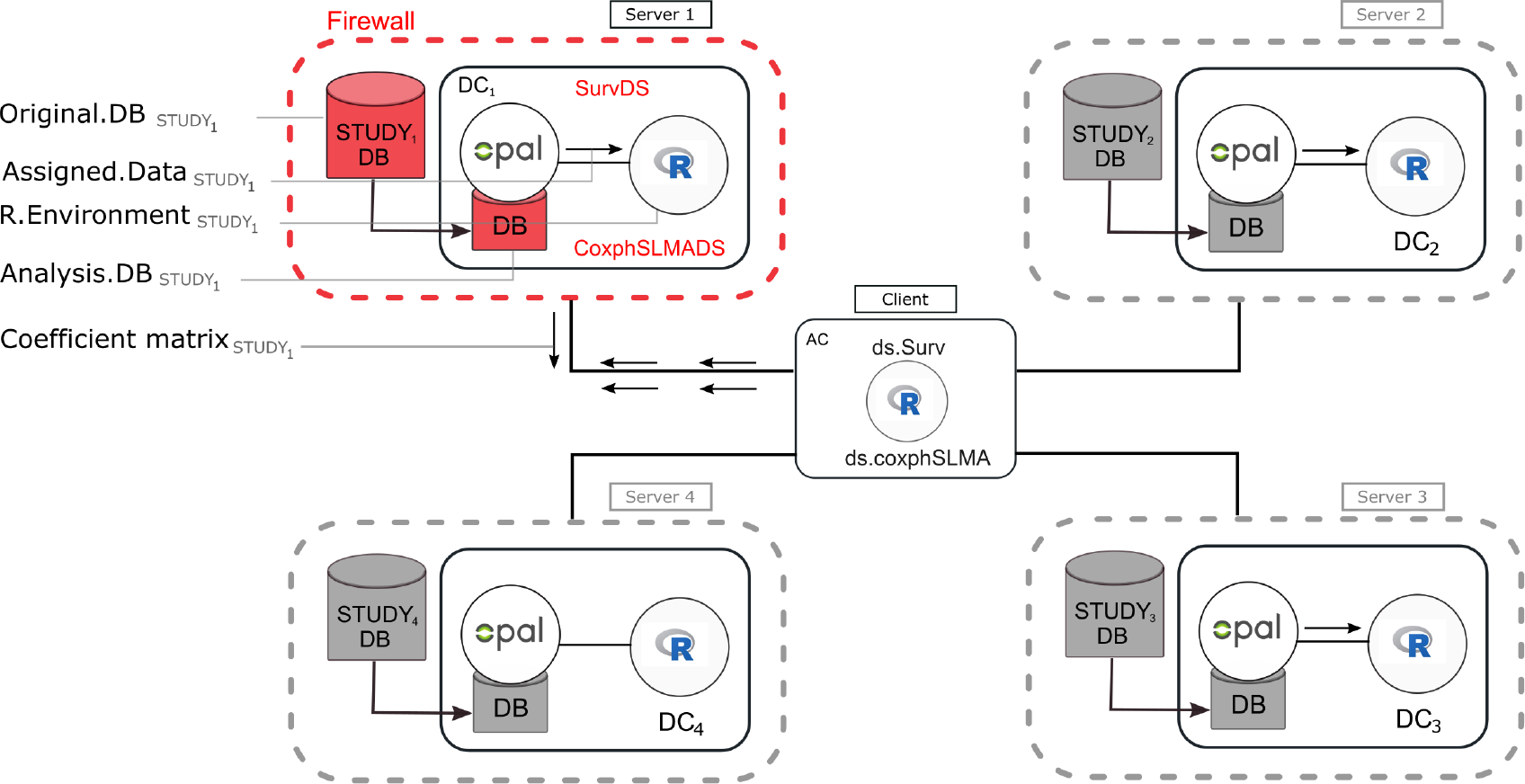
Client-server architecture of DataSHIELD. The diagram shows four study sites/servers (DC) each having data stored in the Original.DB. The analyst (client) sends commands from the analysis computer (AC) to each study site to request the specific data (Assigned.Data) to be analyzed. This could be all the variables or specific variables stored in Analysis.DB. R commands are also sent from the analysis computer to every study telling it to create survival objects and fit the Cox proportional hazard model. Each site responds to instructions sent by creating the survival object and fitting the model. This fitting is carried out in the R environment of each study. The coefficient matrices, standard errors, and odds ratios from each site are then pooled and meta-analyzed using fixed optimization methods, and only non-disclosive statistics are returned to the analyst.

The communication between the client and server for the survival models is shown in Fig. 2 for an assign function (*ds.Surv()* to create survival objects on the servers) and aggregate function (*ds.coxphSLMA()* to perform a meta-analysis of Cox regression models). This shows an asynchronous mode of operation in DataSHIELD where multiple parties (sites) perform secure computation. The client-side package is called dsSurvivalClient and the server-side package is called dsSurvival.

**Figure 2.**
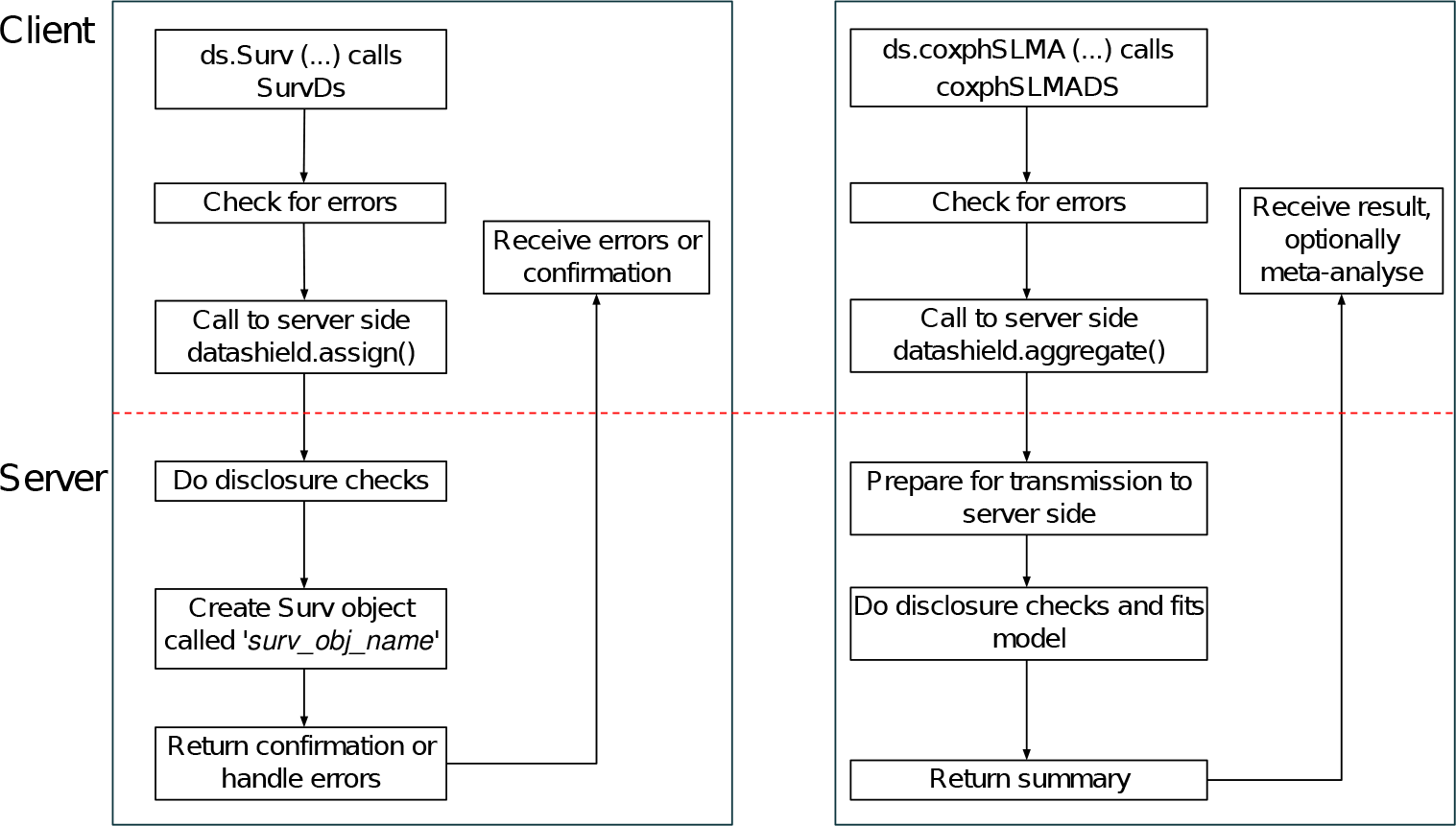
Architecture of client and server side functions for building survival models in dsSurvival. Left panel: An assign function for creating a server-side survival object using *ds.Surv().* Right panel: An aggregate function for a Cox proportional hazards model using *ds.coxphSLMA()*.

The server-side package dsSurvival 1.0.0 contains the functions *SurvDS()* and *coxphSLMADS()*. These functions are configured to reside in modified R environments located behind a firewall at each institution and process the individual-level data at each distinct repository.

The client side package is called dsSurvivalClient and contains the functions *ds.Surv()* and *ds.coxphSLMA()*. These functions reside on the conventional R environment of the analyst. The *ds.Surv()* (assign function) calls the server-side function *SurvDS()* to assign survival objects in each site. This can then be used as the response variable in the *ds.coxphSLMA()* (aggregate) function. The *ds.coxphSLMA()* function calls and controls the corresponding server-side functions *coxphSLMADS()* and performs the regression analysis at different sites. These functions implement study-level meta-analysis (SLMA). The estimates from each site are combined and then pooled using fixed effects or random effects meta-analysis.

### Computational pipeline and use case

We outline the development and code for implementing survival models (Cox regression) and meta-analysis of hazard ratios in our package (dsSurvival).

A tutorial in bookdown format is available here: https://neelsoumya.github.io/dsSurvivalbookdown/

In the following, we demonstrate the computational steps using synthetic data. The first step is using DataSHIELD to connect to the server and loading the survival data. We assume that the reader is familiar with these details. We show the steps using synthetic data. There are 3 data sets that are held on the same server but can be considered to be on separate servers/sites. The variable *EVENT* holds the event information and variables *STARTTIME* and *ENDTIME* hold the time information. There is also age and gender information in variables named *age* and *female*, respectively. We will look at how age and gender affect survival time and then meta-analyze the hazard ratios. For details on how to setup the variables, please see the bookdown above.

#### Algorithm 1: Workflow for performing privacy preserving survival analysis

*%Create survival object*

dsSurvivalClient::ds.Surv(time=‘STARTTIME’, time2=‘ENDTIME’, event = ‘EVENT’, objectname=‘surv_object’, type=‘counting’)

*%use constructed Surv object in ds.coxph.SLMA()*

coxph_model_full = dsSurvivalClient::ds.coxph.SLMA(formula = ‘surv_object ~ D$age+D$female’)

*%Summary of survival objects*

We can also summarize a server-side object of type survival::Surv() to provide a non-disclosive summary of the server-side object.

dsSurvivalClient::ds.coxphSummary(x = ‘coxph_serverside’)

*%Diagnostics for Cox proportional hazards models*

There are functions to test for the assumptions of Cox proportional hazards models.

dsSurvivalClient::ds.coxphSLMAassign(formula = ‘surv_object D$age+D$female’, objectname = ‘coxph_serverside’)

dsSurvivalClient::ds.cox.zphSLMA(fit = ‘coxph_serverside’)

The log-hazard ratios and their standard errors from each study can be found after running *ds.coxphSLMA()*. The hazard ratios can then be meta-analyzed using the *metafor* package [10]. Fig. 3 shows an example forest plot with meta-analysed hazard ratios. The plot shows the log hazard ratios corresponding to age in the survival model.

**Figure 3.**
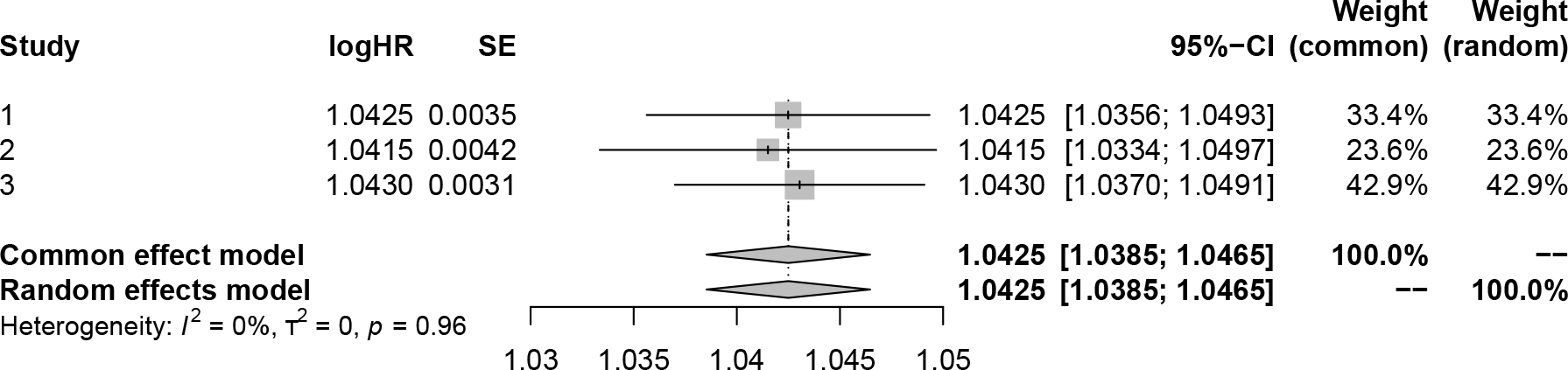
A plot showing the meta-analyzed hazard ratios generated from dsSurvival. A Cox proportional hazards model was fit to synthetic data. The hazard ratios correspond to age in a survival model.

There are two options to generate the survival object. The analyst can generate it separately or inline (for example, by the following command: *dsSurvivalClient::ds.coxph.SLMA(formula = ‘survival::Surgv(time=SURVTIME,event=EVENT) ~ D$age+D$female’)*). If a survival object is generated separately, it is stored on the server and can be used later in an assign function (*ds.coxphSLMAassign()*). This allows the survival model to be stored on the server and can be used later for diagnostics.

### Preserving privacy and disclosure checks

Disclosure checks are an integral part of DataSHIELD and dsSurvival. dsSurvival leverages the DataSHIELD framework to ensure that multiple parties perform secure computation and only the relevant aggregated statistical details are shared. We disallow any Cox models where the number of covariate terms are greater than a fraction (default set to 20%) of the number of data points. The number of data points is the number of entries (for all patients) in the survival data. This fraction can be also be changed by the data custodian or administrator in DataSHIELD. We also deny any access to the baseline hazard function.

### Diagnostics for Cox proportional hazards models

We generate diagnostics for Cox models using the function dsSurvivalClient::ds.cox.zphSLMA(). These diagnostics can allow an analyst to determine if the proportional hazards assumption in Cox proportional hazards models is satisfied. If the p-values returned by dsSurvivalClient::ds.cox.zphSLMA() are greater than 0.05 for a covariate, then the proportional hazards assumption is likely correct for that covariate.

If the proportional hazards assumptions are violated, then the analyst will have to modify the model. Modifications may include introducing strata or using time-dependent covariates.

## Discussion and Conclusion

dsSurvival is a DataSHIELD package for privacy preserving meta-analysis of survival data distributed across different sites. dsSurvival also performs federated calculation of hazard ratios. Its implementation relies exclusively on the distributed algorithm of the DataSHIELD environment. DataSHIELD facilitates important research particularly amongst institutions that are not allowed to transmit patient-level data to an outside server.

Previously building survival models in DataSHIELD involved using approximations like piecewise exponential models. This involves defining time buckets and is an additional burden on the researcher. A lack of familiarity with this approach also makes people less trusting of the results.

Previous work has looked at reducing the dimensions of a survival model and the reduced feature space model is then shared amongst multiple parties [11]. Survival analysis is also possible in DataSHIELD using dsSwissKnife [12]. However, our package offers advantages such as storing the model on the server-side, diagnostics, integration with client-side meta-analysis and future plans to add in more functionality such as survival curves.

We have released an R package for privacy preserving survival analysis in DataSHIELD. Our tool can be of great use in domains where there is a need for building survival models and there are privacy concerns about sharing data. We hope this suite of tools and tutorials will serve as a guideline on how to use survival analysis in a federated environment.

## Supporting information

Client side dsSurvivalClient code

Server side dsSurvival code

Vignette with example code on how to use dsSurvival

Bookdown and documentation for using dsSurvival

Vignette and bookdown with executable code and synthetic data that requires only client side package installation

## Limitations

Our approach implements study-level meta-analysis. This is a computationally faster approach but is also a limitation of our approach. In the future we will implement functionality of iteratively fitting a single model across all studies. We will also develop plotting of privacy preserving survival curves and the ability to have time-dependent covariates in survival models.

## Abbreviations

SLMA: Study Level Meta-Analysis

## Declarations

### Ethics approval and consent to participate

No ethics approval and consent to participate was necessary.

### Consent for publication

Not applicable.

### Availability of data and materials

This study does not generate any data.

A tutorial in bookdown format with code, diagnostics, plots and synthetic data is available here: https://neelsoumya.github.io/dsSurvivalbookdown/

All code is available from the following repositories: https://github.com/neelsoumya/dsSurvivalClient/

https://github.com/neelsoumya/dsSurvival/

### Authors’ contributions

SB and GS carried out the analysis and implementation, participated in the design of the study and drafted the manuscript. TP, DA and PB gave critical comments and edited the manuscript. TB and DZ directed the study. All authors gave final approval for publication.

## Acknowledgements

We acknowledge the help and support of the DataSHIELD technical team. We are especially grateful to Elaine Smith, Stuart Wheater, Patricia Ryser-Welch and Wolfgang Viechtbauer for fruitful discussions and feedback.

