## Supplementary material for "dsSurvival: Privacy preserving survival models for federated individual patient meta-analysis in DataSHIELD": Client side dsSurvivalClient code: build_man_test.R

```
###########################################
# Script to build manuals and test code
#
# Usage:
#   R --no-save < build_man_test.R
#
###########################################

###################
# load libraries
###################
library(devtools)
library(testthat)
library(dsBase)
library(dsBaseClient)
require('DSI')
require('DSOpal')

##################
# build manuals
##################
devtools::build_manual()

########################
# update documentation
########################
devtools::check_man()

##################
# build vignettes
##################
devtools::build_vignettes()

##################
# Testing
##################
devtools::test()

##################
# clean up
##################
gc()
```
