## Supplementary material for "dsSurvival: Privacy preserving survival models for federated individual patient meta-analysis in DataSHIELD": Client side dsSurvivalClient code: CONTRIBUTING.md

# Contributing

When contributing to this repository, please first discuss the change you wish to make via issue,
email, or any other method with the owners of this repository before making a change. 

Please note we have a code of conduct, please follow it in all your interactions with the project.

## Pull Request Process

1. Ensure any install or build dependencies are removed before the end of the layer when doing a 
   build.
2. Update the README.md with details of changes to the interface, this includes new environment 
   variables, exposed ports, useful file locations and container parameters.
3. Increase the version numbers in any examples files and the README.md to the new version that this
   Pull Request would represent. The versioning scheme we use is [SemVer](http://semver.org/).
4. You may merge the Pull Request in once you have the sign-off of two other developers, or if you 
   do not have permission to do that, you may request the second reviewer to merge it for you.

## Code of Conduct

### Our Pledge

In the interest of fostering an open and welcoming environment, we as
contributors and maintainers pledge to making participation in our project and
our community a harassment-free experience for everyone, regardless of age, body
size, disability, ethnicity, gender identity and expression, level of experience,
nationality, personal appearance, race, religion, or sexual identity and
orientation.

### Our Standards

Examples of behavior that contributes to creating a positive environment
include:

* Using welcoming and inclusive language
* Being respectful of differing viewpoints and experiences
* Gracefully accepting constructive criticism
* Focusing on what is best for the community
* Showing empathy towards other community members

Examples of unacceptable behavior by participants include:

* The use of sexualized language or imagery and unwelcome sexual attention or
advances
* Trolling, insulting/derogatory comments, and personal or political attacks
* Public or private harassment
* Publishing others' private information, such as a physical or electronic
  address, without explicit permission
* Other conduct which could reasonably be considered inappropriate in a
  professional setting

### Our Responsibilities

Project maintainers are responsible for clarifying the standards of acceptable
behavior and are expected to take appropriate and fair corrective action in
response to any instances of unacceptable behavior.

Project maintainers have the right and responsibility to remove, edit, or
reject comments, commits, code, wiki edits, issues, and other contributions
that are not aligned to this Code of Conduct, or to ban temporarily or
permanently any contributor for other behaviors that they deem inappropriate,
threatening, offensive, or harmful.

### Scope

This Code of Conduct applies both within project spaces and in public spaces
when an individual is representing the project or its community. Examples of
representing a project or community include using an official project e-mail
address, posting via an official social media account, or acting as an appointed
representative at an online or offline event. Representation of a project may be
further defined and clarified by project maintainers.

### Enforcement

Instances of abusive, harassing, or otherwise unacceptable behavior may be
reported by contacting the project team at []. All
complaints will be reviewed and investigated and will result in a response that
is deemed necessary and appropriate to the circumstances. The project team is
obligated to maintain confidentiality with regard to the reporter of an incident.
Further details of specific enforcement policies may be posted separately.

Project maintainers who do not follow or enforce the Code of Conduct in good
faith may face temporary or permanent repercussions as determined by other
members of the project's leadership.

### Attribution

This Code of Conduct is adapted from the [Contributor Covenant][homepage], version 1.4,
available at [http://contributor-covenant.org/version/1/4][version]

[homepage]: http://contributor-covenant.org
[version]: http://contributor-covenant.org/version/1/4/
