## Supplementary material for "dsSurvival: Privacy preserving survival models for federated individual patient meta-analysis in DataSHIELD": Client side dsSurvivalClient code: dsSurvivalClient_1.0.0.pdf

### Package ‘dsSurvivalClient’

July 5, 2021

**Title** DataSHIELD client functions for building survival models

**Version** 1.0.0

**Author** Soumya Banerjee, Demetris Avraam, Xavier Escriba Montagut, Juan Gonzalez, Paul Burton and Tom R P Bishop <>

**Maintainer** Soumya Banerjee, Demetris Avraam, Xavier Escriba Montagut, Juan Gonzalez, Paul Burton and Tom R P Bishop <>

**Description** DataSHIELD client functions for building survival models.

**License** GPL-3

**Depends** R (¿= 3.5.0),
DSI (¿= 1.1.0)

**Imports** fields,
metafor,
ggplot2,
gridExtra,
data.table,
dsBaseClient

**RoxygenNote** 7.1.1

**Encoding** UTF-8

#### R topics documented:

---

`ds.cox.zphSLMA` *Tests the proportional hazards assumption for a Cox proportional hazards model*

---

##### Description

This function tests the proportional hazards assumption for a Cox proportional hazards model.

##### Usage

```
ds.cox.zphSLMA(
  fit = NULL,
  transform = "km",
  terms = TRUE,
  singledf = FALSE,
  global = TRUE,
  datasources = NULL
)
```

##### Arguments

| | |
|---|---|
| `fit` | character string (potentially including `*` symbol without spaces) specifying the name of the fitted server-side Cox proportioanl hazards model that has been created using ds.coxphSLMAassign() |
| `transform` | character string specifying how the survival times should be transformed before the test is performed. Possible values are "km", "rank", "identity" or a function of one argument. |
| `terms` | logical if TRUE, do a test for each term in the model rather than for each separate covariate. For a factor variable with k levels, for instance, this would lead to a k-1 degree of freedom test. The plot for such variables will be a single curve evaluating the linear predictor over time. |
| `singledf` | logical use a single degree of freedom test for terms that have multiple coefficients, i.e., the test that corresponds most closely to the plot. If terms=FALSE this argument has no effect. |
| `global` | logical should a global chi-square test be done, in addition to the per-variable or per-term tests tests. |
| `datasources` | a list of `DSConnection-class` objects obtained after login. If the `datasources` argument is not specified the default set of connections will be used: see `datashield.connections_default`. For more information see **Details**. |

##### Details

This is a function that performs diagnostics on a fitted Cox proportional hazards model.

Server function called: `cox.zphSLMADS`.

#### Value

`cox.zphSLMADS` returns to the client-side the diagnostics of the Cox proportional hazards model

#### Author(s)

Soumya Banerjee and Tom Bishop, 2021

#### Examples

```
## Not run:

  ## Version 1.0.0

  # connecting to the Opal servers

  require('DSI')
  require('DSOpal')
  require('dsBaseClient')

  builder <- DSI::newDSLoginBuilder()
  builder$append(server = "study1",
                 url = "http://192.168.56.100:8080/",
                 user = "administrator", password = "datashield_test&",
                 table = "SURVIVAL.EXPAND_NO_MISSING1", driver = "OpalDriver")
  builder$append(server = "study2",
                 url = "http://192.168.56.100:8080/",
                 user = "administrator", password = "datashield_test&",
                 table = "SURVIVAL.EXPAND_NO_MISSING2", driver = "OpalDriver")
  builder$append(server = "study3",
                 url = "http://192.168.56.100:8080/",
                 user = "administrator", password = "datashield_test&",
                 table = "SURVIVAL.EXPAND_NO_MISSING3", driver = "OpalDriver")
  logindata <- builder$build()

  connections <- DSI::datashield.login(logins = logindata, assign = TRUE, symbol = "D")

  # make sure that the outcome is numeric
  ds.asNumeric(x.name = "D$cens",
            newobj = "EVENT",
            datasources = connections)

  ds.asNumeric(x.name = "D$survtime",
            newobj = "SURVTIME",
            datasources = connections)

  dsSurvivalClient::ds.Surv(time='SURVTIME', event='EVENT', objectname='surv_object')

  dsSurvivalClient::ds.coxph.SLMA(formula = 'surv_object ~  D$female',
            dataName = 'D', datasources = connections)

  # clear the Datashield R sessions and logout
```

```
  datashield.logout(connections)

## End(Not run)
```

---

ds.coxph.SLMA *Performs survival analysis using Cox proportional hazards model*

---

#### Description

Passes a formula to a server side environment and returns the summary of Cox proportional hazards model from the server.

#### Usage

```
ds.coxph.SLMA(
  formula = NULL,
  dataName = NULL,
  weights = NULL,
  init = NULL,
  ties = "efron",
  singular.ok = TRUE,
  model = FALSE,
  x = FALSE,
  y = TRUE,
  control = NULL,
  combine_with_metafor = FALSE,
  datasources = NULL
)
```

#### Arguments

`formula` character string (potentially including `*` symbol without spaces) specifying the formula that you want to pass to the server-side. For more information see **Details**.

`dataName` character string of name of data frame

`weights` vector of case weights

`init` vector of initial values of the iteration.

`ties` character string specifying the method for tie handling. The Efron approximation is used as the default. Other options are 'breslow' and 'exact'.

`singular.ok` logical value indicating how to handle collinearity in the model matrix. Default is TRUE. If TRUE, the program will automatically skip over columns of the X matrix that are linear combinations of earlier columns. In this case the coefficients of such columns will be NA and the variance matrix will contain zeros.

`model` logical value. If TRUE, the model frame is returned in component model.

x
: logical value. If TRUE, the x matrix is returned in component x.

y
: logical value. If TRUE, the response vector is returned in component y.

control
: object of class survival::coxph.control() specifying iteration limit and other control options. Default is survival::coxph.control()

combine_with_metafor
: logical If TRUE the estimates and standard errors for each regression coefficient are pooled across studies using random-effects meta-analysis under maximum likelihood (ML), restricted maximum likelihood (REML) or fixed-effects meta-analysis (FE). Default is FALSE.

datasources
: a list of DSConnection-class objects obtained after login. If the datasources argument is not specified the default set of connections will be used: see datashield.connections_default.

#### Details

This is a function that performs survival analysis using the Cox proportional hazards model.

Server function called: `coxphSLMADS`.

#### Value

`coxphSLMADS` returns to the client-side a summary of the Cox proportional hazards model

#### Author(s)

Soumya Banerjee and Tom Bishop, 2021

#### Examples

```
## Not run:

  ## Version 1.0.0

  # connecting to the Opal servers

  require('DSI')
  require('DSOpal')
  require('dsBaseClient')

  builder <- DSI::newDSLoginBuilder()
  builder$append(server = "study1",
                 url = "http://192.168.56.100:8080/",
                 user = "administrator", password = "datashield_test&",
                 table = "SURVIVAL.EXPAND_NO_MISSING1", driver = "OpalDriver")
  builder$append(server = "study2",
                 url = "http://192.168.56.100:8080/",
                 user = "administrator", password = "datashield_test&",
                 table = "SURVIVAL.EXPAND_NO_MISSING2", driver = "OpalDriver")
  builder$append(server = "study3",
                 url = "http://192.168.56.100:8080/",
                 user = "administrator", password = "datashield_test&",
```

```
                 table = "SURVIVAL.EXPAND_NO_MISSING3", driver = "OpalDriver")
  logindata <- builder$build()

  connections <- DSI::datashield.login(logins = logindata, assign = TRUE, symbol = "D")

  # make sure that the outcome is numeric
  ds.asNumeric(x.name = "D$cens",
            newobj = "EVENT",
            datasources = connections)

  ds.asNumeric(x.name = "D$survtime",
            newobj = "SURVTIME",
            datasources = connections)

  dsSurvivalClient::ds.Surv(time='SURVTIME', event='EVENT', objectname='surv_object')

  dsSurvivalClient::ds.coxph.SLMA(formula = 'surv_object ~  D$female',
            dataName = 'D', datasources = connections)

  # clear the Datashield R sessions and logout
  datashield.logout(connections)

## End(Not run)
```

---

ds.coxphSLMAassign *Performs survival analysis using Cox proportional hazards model*

---

#### Description

Passes a formula to a server side environment and stores the Cox proportional hazards model from the server.

#### Usage

```
ds.coxphSLMAassign(
  formula = NULL,
  dataName = NULL,
  weights = NULL,
  init = NULL,
  ties = "efron",
  singular.ok = TRUE,
  model = FALSE,
  x = FALSE,
  y = TRUE,
  control = NULL,
  datasources = NULL,
  objectname = NULL
)
```

#### Arguments

- `formula` — character string (potentially including * symbol without spaces) specifying the formula that you want to pass to the server-side. For more information see **Details**.
- `dataName` — character string of name of data frame
- `weights` — vector of case weights
- `init` — vector of initial values of the iteration.
- `ties` — character string specifying the method for tie handling. The Efron approximation is used as the default. Other options are 'breslow' and 'exact'.
- `singular.ok` — logical value indicating how to handle collinearity in the model matrix. Default is TRUE. If TRUE, the program will automatically skip over columns of the X matrix that are linear combinations of earlier columns. In this case the coefficients of such columns will be NA and the variance matrix will contain zeros.
- `model` — logical value. If TRUE, the model frame is returned in component model.
- `x` — logical value. If TRUE, the x matrix is returned in component x.
- `y` — logical value. If TRUE, the response vector is returned in component y.
- `control` — object of class survival::coxph.control() specifying iteration limit and other control options. Default is survival::coxph.control()
- `datasources` — a list of `DSConnection-class` objects obtained after login. If the `datasources` argument is not specified the default set of connections will be used: see `datashield.connections_default`.
- `objectname` — character name of server-side variable to store the Cox model

#### Details

This is a function that performs survival analysis using the Cox proportional hazards model.

Server function called: `coxphSLMAassignDS`.

#### Author(s)

Soumya Banerjee and Tom Bishop, 2021

#### Examples

```
## Not run:

  ## Version 1.0.0

  # connecting to the Opal servers

  require('DSI')
  require('DSOpal')
  require('dsBaseClient')

  builder <- DSI::newDSLoginBuilder()
```

```
  builder$append(server = "study1",
                 url = "http://192.168.56.100:8080/",
                 user = "administrator", password = "datashield_test&",
                 table = "SURVIVAL.EXPAND_NO_MISSING1", driver = "OpalDriver")
  builder$append(server = "study2",
                 url = "http://192.168.56.100:8080/",
                 user = "administrator", password = "datashield_test&",
                 table = "SURVIVAL.EXPAND_NO_MISSING2", driver = "OpalDriver")
  builder$append(server = "study3",
                 url = "http://192.168.56.100:8080/",
                 user = "administrator", password = "datashield_test&",
                 table = "SURVIVAL.EXPAND_NO_MISSING3", driver = "OpalDriver")
  logindata <- builder$build()

  connections <- DSI::datashield.login(logins = logindata, assign = TRUE, symbol = "D")

  # make sure that the outcome is numeric
  ds.asNumeric(x.name = "D$cens",
            newobj = "EVENT",
            datasources = connections)

  ds.asNumeric(x.name = "D$survtime",
            newobj = "SURVTIME",
            datasources = connections)

  dsSurvivalClient::ds.Surv(time='SURVTIME', event='EVENT', objectname='surv_object')

  dsSurvivalClient::ds.coxph.SLMA(formula = 'surv_object ~  D$female',
            dataName = 'D', datasources = connections)

  dsSurvivalClient::ds.coxphSLMAassign(formula = 'surv_object ~  D$female',
             dataName = 'D', datasources = connections,
             objectname = 'coxph_serverside')

  # clear the Datashield R sessions and logout
  datashield.logout(connections)

## End(Not run)
```

---

ds.coxphSummary — *Returns a summary of a server-side Cox proportional hazards model*

---

#### Description

This function returns a summary of server-side for a Cox proportional hazards model.

#### Usage

```
ds.coxphSummary(x = NULL, datasources = NULL)
```

#### Arguments

x
: character string (potentially including * symbol without spaces) specifying the name of the fitted server-side Cox proportioanl hazards model that has been created using ds.coxphSLMAassign()

datasources
: a list of `DSConnection-class` objects obtained after login. If the `datasources` argument is not specified the default set of connections will be used: see `datashield.connections_default`. For more information see **Details**.

#### Details

This is a function that returns a summary of a fitted Cox proportional hazards model.

Server function called: `coxphSummaryDS`.

#### Value

`coxphSummaryDS` returns to the client-side the summary of the Cox proportional hazards model

#### Author(s)

Soumya Banerjee and Tom Bishop, 2021

#### Examples

```
## Not run:

  ## Version 1.0.0

  # connecting to the Opal servers

  require('DSI')
  require('DSOpal')
  require('dsBaseClient')

  builder <- DSI::newDSLoginBuilder()
  builder$append(server = "study1",
                 url = "http://192.168.56.100:8080/",
                 user = "administrator", password = "datashield_test&",
                 table = "SURVIVAL.EXPAND_NO_MISSING1", driver = "OpalDriver")
  builder$append(server = "study2",
                 url = "http://192.168.56.100:8080/",
                 user = "administrator", password = "datashield_test&",
                 table = "SURVIVAL.EXPAND_NO_MISSING2", driver = "OpalDriver")
  builder$append(server = "study3",
                 url = "http://192.168.56.100:8080/",
                 user = "administrator", password = "datashield_test&",
                 table = "SURVIVAL.EXPAND_NO_MISSING3", driver = "OpalDriver")
  logindata <- builder$build()

  connections <- DSI::datashield.login(logins = logindata, assign = TRUE, symbol = "D")
```

```
  # make sure that the outcome is numeric
  ds.asNumeric(x.name = "D$cens",
            newobj = "EVENT",
            datasources = connections)

  ds.asNumeric(x.name = "D$survtime",
            newobj = "SURVTIME",
            datasources = connections)

  dsSurvivalClient::ds.Surv(time='SURVTIME', event='EVENT', objectname='surv_object')

  dsSurvivalClient::ds.coxphSLMAassign(formula = 'surv_object ~  D$female',
             dataName = 'D', datasources = connections,
objectname = 'coxph_serverside')

  dsSurvivalClient::ds.coxphSummary(x = 'coxph_serverside')

  # clear the Datashield R sessions and logout
  datashield.logout(connections)

## End(Not run)
```

---

ds.Surv — *Creates a server-side Survival object. This is used as a response variable in in survival models and Cox proportional hazards models.*

---

#### Description

Creates a server side Survival object of type survival::Surv()

#### Usage

```
ds.Surv(
  time = NULL,
  event = NULL,
  time2 = NULL,
  type = NULL,
  origin = 0,
  objectname = NULL,
  datasources = NULL
)
```

#### Arguments

| | |
|---|---|
| time | character string specifying the server-side start time or follow up timeparameter that has the start time element or follow-up time for survival analysis. |

| | |
|---|---|
| `event` | character string of name of server side event parameter for use in survival analysis |
| `time2` | character string specifying the server-side stop time parameter that has the stop time element for survival analysis. For more information see **Details**. |
| `type` | character string specifying the type of censoring. Possible values are "right", "left", "counting", "interval", "interval2", or "mstate" |
| `origin` | numeric, used for counting process data and is the hazard function origin. The origin parameter is used with time-dependent strata in order to align the subjects properly when they cross over from one strata to another. This parameter has rarely proven useful. |
| `objectname` | character string of name of new server-side object which will store object of class survival::Surv() |
| `datasources` | a list of `DSConnection-class` objects obtained after login. If the `datasources` argument is not specified the default set of connections will be used: see `datashield.connections_default`. |

#### Details

This is a function that Creates a server side Survival object of type survival::Surv(). This can be used to perform survival analysis using the Cox proportional hazards model.

Server function called: `SurvDS`.

#### Value

`SurvDS` returns to the client-side a Surv() obejct for use in the Cox proportional hazards model

#### Author(s)

Soumya Banerjee and Tom Bishop, 2021

#### Examples

```
## Not run:

  ## Version 1.0.0

  # connecting to the Opal servers

  require('DSI')
  require('DSOpal')
  require('dsBaseClient')

  builder <- DSI::newDSLoginBuilder()
  builder$append(server = "study1",
                 url = "http://192.168.56.100:8080/",
                 user = "administrator", password = "datashield_test&",
                 table = "SURVIVAL.EXPAND_NO_MISSING1", driver = "OpalDriver")
```

```
builder$append(server = "study2",
               url = "http://192.168.56.100:8080/",
               user = "administrator", password = "datashield_test&",
               table = "SURVIVAL.EXPAND_NO_MISSING2", driver = "OpalDriver")
builder$append(server = "study3",
               url = "http://192.168.56.100:8080/",
               user = "administrator", password = "datashield_test&",
               table = "SURVIVAL.EXPAND_NO_MISSING3", driver = "OpalDriver")
logindata <- builder$build()

connections <- DSI::datashield.login(logins = logindata, assign = TRUE, symbol = "D")

# make sure that the outcome is numeric
ds.asNumeric(x.name = "D$cens",
          newobj = "EVENT",
          datasources = connections)

ds.asNumeric(x.name = "D$survtime",
          newobj = "SURVTIME",
          datasources = connections)

# create start time variable
ds.asNumeric(x.name = "D$starttime",
          newobj = "STARTTIME",
          datasources = connections)

# create end time variable
ds.asNumeric(x.name = "D$endtime",
          newobj = "ENDTIME",
          datasources = connections)

# create a server-side survival object
dsSurvivalClient::ds.Surv(time='STARTTIME', time2='ENDTIME',
 event = 'EVENT', objectname='surv_object')

# create a Cox proportional hazards model using the created survival object
dsSurvivalClient::ds.coxph.SLMA(formula = 'surv_object~D$age+D$female')

# clear the Datashield R sessions and logout
datashield.logout(connections)

## End(Not run)
```

---

`ds.survfit` *Creates a server-side Survival fit (survfit) object for use in Cox proportional hazards model.*

---

#### Description

Creates a server side Survival fit (survfit) object,

#### Usage

```
ds.survfit(formula = NULL, objectname = NULL, datasources = NULL)
```

#### Arguments

- `formula` character string specifying the formula to be used in survival::survfit() on the server-side. For more information see **Details**.
- `objectname` character string of name of new server-side object which will store object of class survival::Surv()
- `datasources` a list of `DSConnection-class` objects obtained after login. If the `datasources` argument is not specified the default set of connections will be used: see `datashield.connections_default`.

#### Details

This is a function that creates a server side survfit object. This is to be used in plotting results from survival analysis using the Cox proportional hazards model.

Server function called: `survfitDS`.

#### Value

`SurvDS` returns to the client-side a Surv() obejct for use in the Cox proportional hazards model

#### Author(s)

Soumya Banerjee and Tom Bishop, 2021

#### Examples

```
## Not run:

  ## Version 1.0.0

  # connecting to the Opal servers

  require('DSI')
  require('DSOpal')
  require('dsBaseClient')

  builder <- DSI::newDSLoginBuilder()
  builder$append(server = "study1",
                 url = "http://192.168.56.100:8080/",
                 user = "administrator", password = "datashield_test&",
                 table = "SURVIVAL.EXPAND_NO_MISSING1", driver = "OpalDriver")
  builder$append(server = "study2",
                 url = "http://192.168.56.100:8080/",
                 user = "administrator", password = "datashield_test&",
                 table = "SURVIVAL.EXPAND_NO_MISSING2", driver = "OpalDriver")
  builder$append(server = "study3",
```

```
                 url = "http://192.168.56.100:8080/",
                 user = "administrator", password = "datashield_test&",
                 table = "SURVIVAL.EXPAND_NO_MISSING3", driver = "OpalDriver")
  logindata <- builder$build()

  connections <- DSI::datashield.login(logins = logindata, assign = TRUE, symbol = "D")

  # make sure that the outcome is numeric
  ds.asNumeric(x.name = "D$cens",
            newobj = "EVENT",
            datasources = connections)

  ds.asNumeric(x.name = "D$survtime",
            newobj = "SURVTIME",
            datasources = connections)

  dsSurvivalClient::ds.Surv('SURVTIME', 'EVENT', 'surv_object')
  dsSurvivalClient::ds.coxph.SLMA(formula = 'surv_object~D$age+D$female')
  dsSurvivalClient::ds.survfit(formula='surv_object',object='survfit_object')

  # clear the Datashield R sessions and logout
  datashield.logout(connections)

## End(Not run)
```
