## Supplementary material for "dsSurvival: Privacy preserving survival models for federated individual patient meta-analysis in DataSHIELD": Client side dsSurvivalClient code: NEWS.md

v1.0.0: A basic first release of survival models in DataSHIELD. This release has Cox proportional hazards models, summaries of models, diagnostics and the ability to meta-analyze hazard ratios. There is also capability to generate forest plots of meta-analyzed hazard ratios. This release supports study-level meta-analysis (SLMA).


A shiny graphical user interface for building survival models in DataSHIELD has also been created by Xavier Escriba Montagut and Juan Gonzalez. It uses dsSurvival and dsSurvivalClient.


* https://github.com/isglobal-brge/ShinyDataSHIELD

* https://isglobal-brge.github.io/ShinyDataSHIELD_bookdown/

* https://isglobal-brge.github.io/ShinyDataSHIELD_bookdown/functionalities.html#survival-analysis

* https://datashield-demo.obiba.org/


v2.0.0: Forthcoming. This will have privacy preserving survival curves.
