## Supplementary material for "dsSurvival: Privacy preserving survival models for federated individual patient meta-analysis in DataSHIELD": Client side dsSurvivalClient code: paper.bib

@article{Viechtbauer2010,
abstract = {Epigenetic biomarkers of aging (the “epigenetic clock”) have the potential to address puzzling findings surrounding mortality rates and incidence of cardio-metabolic disease such as: (1) women consistently exhibiting lower mortality than men despite having higher levels of morbidity; (2) racial/ethnic groups having different mortality rates even after adjusting for socioeconomic differences; (3) the black/white mortality cross-over effect in late adulthood; and (4) Hispanics in the United States having a longer life expectancy than Caucasians despite having a higher burden of traditional cardio-metabolic risk factors. We analyzed blood, saliva, and brain samples from seven different racial/ethnic groups. We assessed the intrinsic epigenetic age acceleration of blood (independent of blood cell counts) and the extrinsic epigenetic aging rates of blood (dependent on blood cell counts and tracks the age of the immune system). In blood, Hispanics and Tsimane Amerindians have lower intrinsic but higher extrinsic epigenetic aging rates than Caucasians. African-Americans have lower extrinsic epigenetic aging rates than Caucasians and Hispanics but no differences were found for the intrinsic measure. Men have higher epigenetic aging rates than women in blood, saliva, and brain tissue. Epigenetic aging rates are significantly associated with sex, race/ethnicity, and to a lesser extent with CHD risk factors, but not with incident CHD outcomes. These results may help elucidate lower than expected mortality rates observed in Hispanics, older African-Americans, and women.},
archivePrefix = {arXiv},
arxivId = {arXiv:1501.0228},
author = {Viechtbauer, Wolfgang},
doi = {10.18637/jss.v036.i03},
eprint = {arXiv:1501.0228},
isbn = {1548-7660},
issn = {1548-7660},
journal = {J. Stat. Softw.},
mendeley-groups = {Periphery{\_}project,cam{\_}project},
month = {aug},
number = {3},
pages = {1--48},
pmid = {18291371},
title = {{Conducting Meta-Analyses in R with the metafor Package}},
url = {http://www.jstatsoft.org/v36/i03/},
volume = {36},
year = {2010}
}
