## Supplementary material for "dsSurvival: Privacy preserving survival models for federated individual patient meta-analysis in DataSHIELD": Client side dsSurvivalClient code: paper.md

A sample submission file from:

# https://joss.readthedocs.io/en/latest/submitting.html

---
title: 'Gala: A Python package for galactic dynamics'
tags:
  - Python
  - astronomy
  - dynamics
  - galactic dynamics
  - milky way
authors:
  - name: Adrian M. Price-Whelan^[co-first author] # note this makes a footnote saying 'co-first author'
    orcid: 0000-0003-0872-7098
    affiliation: "1, 2" # (Multiple affiliations must be quoted)
  - name: Author Without ORCID^[co-first author] # note this makes a footnote saying 'co-first author'
    affiliation: 2
  - name: Author with no affiliation^[corresponding author]
    affiliation: 3
affiliations:
 - name: Lyman Spitzer, Jr. Fellow, Princeton University
   index: 1
 - name: Institution Name
   index: 2
 - name: Independent Researcher
   index: 3
date: 13 August 2017
bibliography: paper.bib

# Optional fields if submitting to a AAS journal too, see this blog post:
# https://blog.joss.theoj.org/2018/12/a-new-collaboration-with-aas-publishing
aas-doi: 10.3847/xxxxx <- update this with the DOI from AAS once you know it.
aas-journal: Astrophysical Journal <- The name of the AAS journal.
---

# Summary

The forces on stars, galaxies, and dark matter under external gravitational
fields lead to the dynamical evolution of structures in the universe. The orbits
of these bodies are therefore key to understanding the formation, history, and
future state of galaxies. The field of "galactic dynamics," which aims to model
the gravitating components of galaxies to study their structure and evolution,
is now well-established, commonly taught, and frequently used in astronomy.
Aside from toy problems and demonstrations, the majority of problems require
efficient numerical tools, many of which require the same base code (e.g., for
performing numerical orbit integration).

# Statement of need

`Gala` is an Astropy-affiliated Python package for galactic dynamics. Python
enables wrapping low-level languages (e.g., C) for speed without losing
flexibility or ease-of-use in the user-interface. The API for `Gala` was
designed to provide a class-based and user-friendly interface to fast (C or
Cython-optimized) implementations of common operations such as gravitational
potential and force evaluation, orbit integration, dynamical transformations,
and chaos indicators for nonlinear dynamics. `Gala` also relies heavily on and
interfaces well with the implementations of physical units and astronomical
coordinate systems in the `Astropy` package [@astropy] (`astropy.units` and
`astropy.coordinates`).

`Gala` was designed to be used by both astronomical researchers and by
students in courses on gravitational dynamics or astronomy. It has already been
used in a number of scientific publications [@Pearson:2017] and has also been
used in graduate courses on Galactic dynamics to, e.g., provide interactive
visualizations of textbook material [@Binney:2008]. The combination of speed,
design, and support for Astropy functionality in `Gala` will enable exciting
scientific explorations of forthcoming data releases from the *Gaia* mission
[@gaia] by students and experts alike.

# Mathematics

Single dollars ($) are required for inline mathematics e.g. $f(x) = e^{\pi/x}$

Double dollars make self-standing equations:

$$\Theta(x) = \left\{\begin{array}{l}
0\textrm{ if } x < 0\cr
1\textrm{ else}
\end{array}\right.$$

You can also use plain \LaTeX for equations
\begin{equation}\label{eq:fourier}
\hat f(\omega) = \int_{-\infty}^{\infty} f(x) e^{i\omega x} dx
\end{equation}
and refer to \autoref{eq:fourier} from text.

# Citations

Citations to entries in paper.bib should be in
[rMarkdown](http://rmarkdown.rstudio.com/authoring_bibliographies_and_citations.html)
format.

If you want to cite a software repository URL (e.g. something on GitHub without a preferred
citation) then you can do it with the example BibTeX entry below for @fidgit.

For a quick reference, the following citation commands can be used:
- `@author:2001`  ->  "Author et al. (2001)"
- `[@author:2001]` -> "(Author et al., 2001)"
- `[@author1:2001; @author2:2001]` -> "(Author1 et al., 2001; Author2 et al., 2002)"

# Figures

Figures can be included like this:
![Caption for example figure.\label{fig:example}](figure.png)
and referenced from text using \autoref{fig:example}.

Figure sizes can be customized by adding an optional second parameter:
![Caption for example figure.](figure.png){ width=20% }

# Acknowledgements

We acknowledge contributions from Brigitta Sipocz, Syrtis Major, and Semyeong
Oh, and support from Kathryn Johnston during the genesis of this project.

# References
