## Supplementary material for "dsSurvival: Privacy preserving survival models for federated individual patient meta-analysis in DataSHIELD": Client side dsSurvivalClient code: ds.cox.zphSLMA.R

```
#' @title Tests the proportional hazards assumption for a Cox proportional hazards model
#' @description This function tests the proportional hazards assumption 
#'	for a Cox proportional hazards model.
#' @details This is a function that performs diagnostics on a fitted Cox 
#' 	proportional hazards model. 
#' 
#' Server function called: \code{cox.zphSLMADS}. 
#' 
#' @param fit character string (potentially including \code{*} symbol without spaces) 
#' specifying the name of the fitted server-side Cox proportioanl hazards model
#'	 that has been created using ds.coxphSLMAassign()
#' @param transform character string specifying how the survival times should be transformed
#' 	before the test is performed. Possible values are "km", "rank", "identity" or a
#' 	function of one argument.
#' @param terms logical if TRUE, do a test for each term in the model rather than for each
#'	separate covariate. For a factor variable with k levels, for instance, this would lead
#'	to a k-1 degree of freedom test. The plot for such variables will be a single curve
#'	evaluating the linear predictor over time.
#' @param singledf logical use a single degree of freedom test for terms that have multiple 
#'	coefficients, i.e., the test that corresponds most closely to the plot. If terms=FALSE
#'	this argument has no effect.
#' @param global logical should a global chi-square test be done, in addition to the per-variable
#'	or per-term tests tests. 
#' @param datasources a list of \code{\link{DSConnection-class}} objects obtained after login. 
#' If the \code{datasources} argument is not specified
#' the default set of connections will be used: see \code{\link{datashield.connections_default}}.
#' For more information see \strong{Details}. 
#' @return \code{cox.zphSLMADS} returns to the client-side the diagnostics of
#' 	the Cox proportional hazards model
#' @author Soumya Banerjee and Tom Bishop, 2021
#' @examples
#' \dontrun{
#'
#'   ## Version 1.0.0
#'   
#'   # connecting to the Opal servers
#' 
#'   require('DSI')
#'   require('DSOpal')
#'   require('dsBaseClient')
#'
#'   builder <- DSI::newDSLoginBuilder()
#'   builder$append(server = "study1", 
#'                  url = "http://192.168.56.100:8080/", 
#'                  user = "administrator", password = "datashield_test&", 
#'                  table = "SURVIVAL.EXPAND_NO_MISSING1", driver = "OpalDriver")
#'   builder$append(server = "study2", 
#'                  url = "http://192.168.56.100:8080/", 
#'                  user = "administrator", password = "datashield_test&", 
#'                  table = "SURVIVAL.EXPAND_NO_MISSING2", driver = "OpalDriver")
#'   builder$append(server = "study3",
#'                  url = "http://192.168.56.100:8080/", 
#'                  user = "administrator", password = "datashield_test&", 
#'                  table = "SURVIVAL.EXPAND_NO_MISSING3", driver = "OpalDriver")
#'   logindata <- builder$build()
#'   
#'   connections <- DSI::datashield.login(logins = logindata, assign = TRUE, symbol = "D") 
#'   
#'   # make sure that the outcome is numeric 
#'   ds.asNumeric(x.name = "D$cens",
#'             newobj = "EVENT",
#'             datasources = connections)
#'
#'   ds.asNumeric(x.name = "D$survtime",
#'             newobj = "SURVTIME",
#'             datasources = connections)
#'
#'   dsSurvivalClient::ds.Surv(time='SURVTIME', event='EVENT', objectname='surv_object')
#'
#'   dsSurvivalClient::ds.coxph.SLMA(formula = 'surv_object ~  D$female', 
#'             dataName = 'D', datasources = connections)
#'   
#'   # clear the Datashield R sessions and logout
#'   datashield.logout(connections)
#' }
#'
#' @export
ds.cox.zphSLMA <- function(fit = NULL,
			   transform = 'km',
			   terms = TRUE,
			   singledf = FALSE,
			   global = TRUE,
			   datasources = NULL
			  )
{
   
   # look for DS connections
   # if one not provided then get current
   if(is.null(datasources))
   {
      datasources <- datashield.connections_find()
   }
      
   # verify that 'fit' was set
   if(is.null(fit))
   {
      stop(" Please provide a valid name for a server-side Cox proportional hazards model that has been fit to data !", call.=FALSE)
   }
   
   calltext <- call("cox.zphSLMADS", fit, transform, terms, singledf, global)

   # call aggregate function
   output <- datashield.aggregate(datasources, calltext)
  
   # return summary of coxph model
   return(output)
	
}
#ds.cox.zphSLMA
```
