## Supplementary material for "dsSurvival: Privacy preserving survival models for federated individual patient meta-analysis in DataSHIELD": Client side dsSurvivalClient code: ds.coxph.SLMA.R

```
#' @title Performs survival analysis using Cox proportional hazards model
#' @description Passes a formula to a server side environment and returns the summary of 
#' Cox proportional hazards model from the server. 
#' @details This is a function that performs survival analysis using the Cox 
#' proportional hazards model. 
#' 
#' Server function called: \code{coxphSLMADS}. 
#' 
#' @param formula character string (potentially including \code{*} symbol without spaces) 
#' specifying the formula that you want to pass to the server-side.
#' For more information see \strong{Details}. 
#' @param dataName character string of name of data frame
#' @param weights vector of case weights
#' @param init vector of initial values of the iteration.
#' @param ties character string specifying the method for tie handling. The Efron approximation is
#'	used as the default. Other options are 'breslow' and 'exact'.
#' @param singular.ok logical value indicating how to handle collinearity in the model matrix.
#'	Default is TRUE. If TRUE, the program will automatically skip over columns of the X matrix
#'	that are linear combinations of earlier columns. In this case the coefficients of such
#'	columns will be NA and the variance matrix will contain zeros. 
#' @param model logical value. If TRUE, the model frame is returned in component model.
#' @param x logical value. If TRUE, the x matrix is returned in component x.
#' @param y logical value. If TRUE, the response vector is returned in component y.
#' @param control object of class survival::coxph.control() specifying iteration limit and 
#'		other control options. Default is survival::coxph.control()
#' @param combine_with_metafor logical If TRUE the
#' 	estimates and standard errors for each regression coefficient are pooled across
#' 	studies using random-effects meta-analysis under maximum likelihood (ML),
#' 	restricted maximum likelihood (REML) or fixed-effects meta-analysis (FE). Default is FALSE. 
#' @param datasources a list of \code{\link{DSConnection-class}} objects obtained after login. 
#' If the \code{datasources} argument is not specified
#' the default set of connections will be used: see \code{\link{datashield.connections_default}}.
#' @return \code{coxphSLMADS} returns to the client-side a summary of 
#' the Cox proportional hazards model
#' @author Soumya Banerjee and Tom Bishop, 2021
#' @examples
#' \dontrun{
#'
#'   ## Version 1.0.0
#'   
#'   # connecting to the Opal servers
#' 
#'   require('DSI')
#'   require('DSOpal')
#'   require('dsBaseClient')
#'
#'   builder <- DSI::newDSLoginBuilder()
#'   builder$append(server = "study1", 
#'                  url = "http://192.168.56.100:8080/", 
#'                  user = "administrator", password = "datashield_test&", 
#'                  table = "SURVIVAL.EXPAND_NO_MISSING1", driver = "OpalDriver")
#'   builder$append(server = "study2", 
#'                  url = "http://192.168.56.100:8080/", 
#'                  user = "administrator", password = "datashield_test&", 
#'                  table = "SURVIVAL.EXPAND_NO_MISSING2", driver = "OpalDriver")
#'   builder$append(server = "study3",
#'                  url = "http://192.168.56.100:8080/", 
#'                  user = "administrator", password = "datashield_test&", 
#'                  table = "SURVIVAL.EXPAND_NO_MISSING3", driver = "OpalDriver")
#'   logindata <- builder$build()
#'   
#'   connections <- DSI::datashield.login(logins = logindata, assign = TRUE, symbol = "D") 
#'   
#'   # make sure that the outcome is numeric 
#'   ds.asNumeric(x.name = "D$cens",
#'             newobj = "EVENT",
#'             datasources = connections)
#'
#'   ds.asNumeric(x.name = "D$survtime",
#'             newobj = "SURVTIME",
#'             datasources = connections)
#'
#'   dsSurvivalClient::ds.Surv(time='SURVTIME', event='EVENT', objectname='surv_object')
#'
#'   dsSurvivalClient::ds.coxph.SLMA(formula = 'surv_object ~  D$female', 
#'             dataName = 'D', datasources = connections)
#'   
#'   # clear the Datashield R sessions and logout
#'   datashield.logout(connections)
#' }
#'
#' @export
ds.coxph.SLMA <- function(formula = NULL,
			  dataName = NULL,
			  weights = NULL,
			  init = NULL,
			  ties = 'efron',
			  singular.ok = TRUE,
			  model = FALSE,
			  x = FALSE,
			  y = TRUE,
			  control = NULL,
			  combine_with_metafor = FALSE,
			  datasources = NULL)
{
   
   # look for DS connections
   # if one not provided then get current
   if(is.null(datasources))
   {
      datasources <- datashield.connections_find()
   }
   
   # if the argument 'dataName' is set, check that the data frame is defined (i.e. exists) on the server site
   if(!(is.null(dataName)))
   {
      # TODO: cannot find function isDefined but is is inds.glmerSLMA
      # defined <- isDefined(datasources, dataName)
   }
   
   # verify that 'formula' was set
   if(is.null(formula))
   {
      stop(" Please provide a valid survival formula!", call.=FALSE)
   }
   
   
   formula = stats::as.formula(formula)
   
   ####################################################################	
   # Logic for parsing formula: since this need to be passed
   #     to parser, we need to remove special symbols
   #     On the server-side function (coxphSLMADS) this needs
   #     to be reconstructed
   #     formula as text, then split at pipes to avoid triggering parser
   ####################################################################
   formula <- Reduce(paste, deparse(formula))
   formula <- gsub("survival::Surv(", "sssss", formula, fixed = TRUE)
   formula <- gsub("|", "xxx", formula, fixed = TRUE)
   formula <- gsub("(", "yyy", formula, fixed = TRUE)
   formula <- gsub(")", "zzz", formula, fixed = TRUE)
   formula <- gsub("/", "ppp", formula, fixed = TRUE)
   formula <- gsub(":", "qqq", formula, fixed = TRUE)
   formula <- gsub(",", "rrr", formula, fixed = TRUE)
   formula <- gsub(" ", "",    formula, fixed = TRUE)
   formula <- gsub("=", "lll", formula, fixed = TRUE)
   # "survival::Surv(time=SURVTIME,event=EVENT)~D$female"
   # gets converted to EVENTzzz ~ D$female
   cat(formula)
	
   # convert to formula otherwise we get parser error
   formula <- stats::as.formula(formula)
   #formula <- strsplit(x = formurand()la, split="|", fixed=TRUE)[[1]]
   
   
   ####################################################################
   # Logic for parsing control argument
   ####################################################################
   if (!is.null(control))
   {	
        # everything needs to be passed as formula to server
	#	otherwise will not go through parser
	#	and a formula needs a ~ something
	#	so introduce dummy ~ something and remove
	#	it on server side   
	control <- paste0(control, "~bbbb")  
	   
        control <- Reduce(paste, deparse(control))
        control <- gsub("survival::coxph.control(", "aaaaa", control, fixed =  TRUE)
        control <- gsub("|", "xxx", control, fixed = TRUE)
        control <- gsub("(", "yyy", control, fixed = TRUE)
        control <- gsub(")", "zzz", control, fixed = TRUE)
        control <- gsub("/", "ppp", control, fixed = TRUE)
        control <- gsub(":", "qqq", control, fixed = TRUE)
	control <- gsub(",", "rrr", control, fixed = TRUE)
        control <- gsub(" ", "",    control, fixed = TRUE)
        control <- gsub("=", "lll", control, fixed = TRUE)
	
	control <- stats::as.formula(control)   
   }	   
	
	
   calltext <- call("coxphSLMADS", formula=formula, dataName, weights, init, ties, singular.ok, model, x, y, control)
   
   # call aggregate function
   output <- datashield.aggregate(datasources, calltext)
  
   # return summary of coxph model
   if (combine_with_metafor == FALSE)
   {	   
       # do not combine with metafor return summary of Cox model	   
       return(output)
   }
   else
   {
       ###############################	   
       # combine with metafor
       ###############################

       # get number of studies
       numstudies <- length(datasources)

       # get the max number of coefficients in model
       # numcoefficients <- length( output[[1]]$coefficients[,1] )
       # create a variable to store max number of coefficients	   
       numcoefficients_max <- 0
  
       # for each study find out the number of coefficients and then get max	   
       for (g in 1:numstudies)
       {
	   # if the number of coefficients in the g th study is greater than max,
	   #     then make it the new max    
           if (length(output[[g]]$coefficients[,1]) > numcoefficients_max)
	   {
               numcoefficients_max <- length(output[[g]]$coefficients[,1])
           }
       }	   

       # assign this max number of coefficients to the variable numcoefficients
       numcoefficients <- numcoefficients_max	   
	   
       # initialize matrices to store coefficient values and standard errors
       betamatrix <- matrix(NA, nrow = numcoefficients, ncol = numstudies)
       sematrix   <- matrix(NA, nrow = numcoefficients, ncol = numstudies)	   
	   
       # for each study store these values
       for (k in 1:numstudies)
       {
           betamatrix[,k] <- output[[k]]$coefficients[,1]
           sematrix[,k]   <- output[[k]]$coefficients[,2]
       }
	   
       # create a list to store all RMA metafor values
       SLMA.pooled.ests.matrix <- matrix(NA, nrow = numcoefficients, ncol = 6)
       # create meaningful column names	   
       dimnames(SLMA.pooled.ests.matrix) <- list(dimnames(betamatrix)[[1]],
                                                 c("pooled.ML","se.ML","pooled.REML","se.REML","pooled.FE","se.FE")
					         )
       
       # call metafor::rma() for each study and call with ML, REML and FE options
       for(p in 1:numcoefficients)
       {
           rma.ML  <- metafor::rma(yi = as.matrix(betamatrix)[p,], sei = as.matrix(sematrix)[p,], method = "ML")
           rma.REML<- metafor::rma(yi = as.matrix(betamatrix)[p,], sei = as.matrix(sematrix)[p,], method = "REML")
           rma.FE  <- metafor::rma(yi = as.matrix(betamatrix)[p,], sei = as.matrix(sematrix)[p,], method = "FE")
    
           SLMA.pooled.ests.matrix[p,1] <- rma.ML$beta
           SLMA.pooled.ests.matrix[p,2] <- rma.ML$se
    
           SLMA.pooled.ests.matrix[p,3] <- rma.REML$beta
           SLMA.pooled.ests.matrix[p,4] <- rma.REML$se
    
           SLMA.pooled.ests.matrix[p,5] <- rma.FE$beta
           SLMA.pooled.ests.matrix[p,6] <- rma.FE$se
       }	   
	
       # return this SLMA pooled metafor::rma() list
       return (list(output = output,
		    betamatrix = betamatrix,
		    sematrix = sematrix,
		    SLMA.pooled.ests.matrix = SLMA.pooled.ests.matrix
		   )
	      )
	   
   }	   

	
}
#ds.coxph.SLMA
```
