## Supplementary material for "dsSurvival: Privacy preserving survival models for federated individual patient meta-analysis in DataSHIELD": Client side dsSurvivalClient code: ds.Surv.R

```
#' @title Creates a server-side Survival object. This is used as a response variable in 
#'	in survival models and Cox proportional hazards models.
#' @description Creates a server side Survival object of type survival::Surv()
#' @details This is a function that Creates a server side Survival object of type 
#' 	survival::Surv(). This can be used to perform survival analysis using the Cox 
#' 	proportional hazards model. 
#' 
#' Server function called: \code{SurvDS}. 
#' 
#' @param time character string  
#' specifying the server-side start time or follow up timeparameter that has the 
#' 	start time element or follow-up time for survival analysis.
#' @param event character string of name of server side event parameter for
#'    use in survival analysis
#' @param time2 character string  
#' specifying the server-side stop time parameter that has the stop time element for survival analysis.
#' For more information see \strong{Details}. 
#' @param type character string specifying the type of censoring. Possible values are "right", "left",
#'	"counting", "interval", "interval2", or "mstate"
#' @param origin numeric, used for counting process data and is the hazard function origin.
#'	The origin parameter is used with time-dependent strata in order to align the subjects
#'	properly when they cross over from one strata to another. This parameter has rarely
#'	proven useful.
#' @param objectname character string of name of new server-side object which will
#'  	store object of class survival::Surv()
#' @param datasources a list of \code{\link{DSConnection-class}} objects obtained after login. 
#' If the \code{datasources} argument is not specified
#' the default set of connections will be used: see \code{\link{datashield.connections_default}}.
#' @return \code{SurvDS} returns to the client-side a Surv() obejct for use in
#' the Cox proportional hazards model
#' @author Soumya Banerjee and Tom Bishop, 2021
#' @examples
#' \dontrun{
#'
#'   ## Version 1.0.0
#'   
#'   # connecting to the Opal servers
#' 
#'   require('DSI')
#'   require('DSOpal')
#'   require('dsBaseClient')
#'
#'   builder <- DSI::newDSLoginBuilder()
#'   builder$append(server = "study1", 
#'                  url = "http://192.168.56.100:8080/", 
#'                  user = "administrator", password = "datashield_test&", 
#'                  table = "SURVIVAL.EXPAND_NO_MISSING1", driver = "OpalDriver")
#'   builder$append(server = "study2", 
#'                  url = "http://192.168.56.100:8080/", 
#'                  user = "administrator", password = "datashield_test&", 
#'                  table = "SURVIVAL.EXPAND_NO_MISSING2", driver = "OpalDriver")
#'   builder$append(server = "study3",
#'                  url = "http://192.168.56.100:8080/", 
#'                  user = "administrator", password = "datashield_test&", 
#'                  table = "SURVIVAL.EXPAND_NO_MISSING3", driver = "OpalDriver")
#'   logindata <- builder$build()
#'   
#'   connections <- DSI::datashield.login(logins = logindata, assign = TRUE, symbol = "D") 
#'   
#'   # make sure that the outcome is numeric 
#'   ds.asNumeric(x.name = "D$cens",
#'             newobj = "EVENT",
#'             datasources = connections)
#'
#'   ds.asNumeric(x.name = "D$survtime",
#'             newobj = "SURVTIME",
#'             datasources = connections)
#'
#'   # create start time variable
#'   ds.asNumeric(x.name = "D$starttime",
#'             newobj = "STARTTIME",
#'             datasources = connections)
#'
#'   # create end time variable
#'   ds.asNumeric(x.name = "D$endtime",
#'             newobj = "ENDTIME",
#'             datasources = connections)
#'
#'   # create a server-side survival object
#'   dsSurvivalClient::ds.Surv(time='STARTTIME', time2='ENDTIME', 
#'			   event = 'EVENT', objectname='surv_object')
#'
#'   # create a Cox proportional hazards model using the created survival object	
#'   dsSurvivalClient::ds.coxph.SLMA(formula = 'surv_object~D$age+D$female')
#'   
#'   # clear the Datashield R sessions and logout
#'   datashield.logout(connections)
#' }
#'
#' @export
ds.Surv <- function(time = NULL, 
		    event = NULL,
		    time2 = NULL,  
		    type = NULL, # c('right', 'left', 'interval', 'counting', 'interval2', 'mstate'),
		    origin = 0, # NULL, 
		    objectname = NULL, 
		    datasources = NULL)
{
   
   # look for DS connections
   # if one not provided then get current
   if(is.null(datasources))
   {
      datasources <- datashield.connections_find()
   }
   
   # if the argument 'event' is set, check that the data frame is defined (i.e. exists) on the server site
   if(!(is.null(event)))
   {
      # TODO: cannot find function isDefined but is is inds.glmerSLMA
      # defined <- isDefined(datasources, event)
   }
      
   # verify that 'time' was set
   if(is.null(time))
   {
      stop(" Please provide a valid survival start time or follow-up time parameter", call.=FALSE)
   }

   # verify that 'stop' was set	
   #if(is.null(stop))
   #{
   #   stop(" Please provide a valid survival stop time parameter", call.=FALSE)
   #}

   # verify that 'event' was set
   if(is.null(event))
   {
      stop(" Please provide a valid survival event parameter", call.=FALSE)
   }

   # verify that 'objectname' was set
   if(is.null(objectname))
   {
      stop(" Please provide a valid objectname to store the server-side survival::Surv() object", call.=FALSE)
   }

   
   # call the server side function
   calltext <- call("SurvDS", time, time2, event, type, origin) # SurvDS
   
   # call aggregate function
   output <- DSI::datashield.assign(conns = datasources, value = calltext, symbol = objectname) # 'surv_object') 
 
   # return summary of coxph model
   # output <- NULL
   return(output)
	
}
#ds.Surv
```
