## Supplementary material for "dsSurvival: Privacy preserving survival models for federated individual patient meta-analysis in DataSHIELD": Client side dsSurvivalClient code: README.md

# dsSurvivalClient

[![License](https://img.shields.io/badge/license-GPLv3-blue.svg)](https://www.gnu.org/licenses/gpl-3.0.html)

## Introduction

This is a standalone package for building survival functions (client side) for DataSHIELD (a platform for federated analysis of private data). These are client side functions for building survival models, Cox proportional hazards models and Cox regression models.

A tutorial in bookdown format with executable code is available here:

https://neelsoumya.github.io/dsSurvival_bookdown/


DataSHIELD is a platform for federated analysis of private data. DataSHIELD has a client-server architecture and this package has a client side and server side component.

* The server side package is called dsSurvival:

    * https://github.com/neelsoumya/dsSurvival

* The client side package is called dsSurvivalClient:

    * https://github.com/neelsoumya/dsSurvivalClient


## Quick start

Install R 

   https://www.r-project.org/

and R Studio 

   https://www.rstudio.com/products/rstudio/download/preview/


Install the following packages:


```r 
install.packages('devtools')
library(devtools)
devtools::install_github('neelsoumya/dsSurvivalClient')
devtools::install_github('datashield/dsBaseClient@6.1.1')
install.packages('rmarkdown')
install.packages('knitr')
install.packages('tinytex')
install.packages('metafor')
install.packages('DSOpal')
install.packages('DSI')
install.packages('opalr')
```


Follow the tutorial in bookdown format with executable code:

https://neelsoumya.github.io/dsSurvival_bookdown/


## Installation

* Install R Studio and the development environment as described below:

    * https://data2knowledge.atlassian.net/wiki/spaces/DSDEV/pages/12943461/Getting+started


* Install the virtual machines as described below:

    * https://data2knowledge.atlassian.net/wiki/spaces/DSDEV/pages/931069953/Installation+Training+Hub-+DataSHIELD+v6

    * https://data2knowledge.atlassian.net/wiki/spaces/DSDEV/pages/1657634881/Testing+100+VM

    * https://data2knowledge.atlassian.net/wiki/spaces/DSDEV/pages/1657634898/Tutorial+6.1.0+100+VM

* Install dsBase and dsSurvival on Opal server in the Virtual Machine (type neelsoumya/dsSurvival and main in the textboxes) as shown in the screenshot below

![Screenshot of installation of package in VM](Capture_VM_install_screenshot.PNG)


```r

install.packages('devtools')

library(devtools)

devtools::install_github('neelsoumya/dsBaseClient')

devtools::install_github('neelsoumya/dsSurvivalClient')

```

If you want to use a certain release then you can do the following

```r

library(devtools)

devtools::install_github('neelsoumya/dsSurvivalClient@v1.0.0')

```

If you want to try privacy preserving survival curves (work in progress and to be available in v2.0), then you can do the following

```r

library(devtools)

devtools::install_github('neelsoumya/dsSurvivalClient', ref = 'privacy_survival_curves')

```


## Usage

A tutorial in bookdown format is available here: 

https://neelsoumya.github.io/dsSurvival_bookdown/


A screenshot of meta-analyzed hazard ratios from a survival model is shown below.

![Meta-analyzed hazard ratios from survival models](screenshot_survival_models.png)


## Release notes

v1.0.0: A basic release of survival models in DataSHIELD. This release has Cox proportional hazards models, summaries of models, diagnostics and the ability to meta-analyze hazard ratios. There is also capability to generate forest plots of meta-analyzed hazard ratios. This release supports study-level meta-analysis.

A shiny graphical user interface for building survival models in DataSHIELD has also been created by Xavier Escriba Montagut and Juan Gonzalez. It calls dsSurvival and dsSurvivalClient.


* https://github.com/isglobal-brge/ShinyDataSHIELD

* https://isglobal-brge.github.io/ShinyDataSHIELD_bookdown/

* https://isglobal-brge.github.io/ShinyDataSHIELD_bookdown/functionalities.html#survival-analysis

* https://datashield-demo.obiba.org/


v2.0.0: Forthcoming. This release will have privacy preserving survival curves.


## Acknowledgements

We acknowledge the help and support of the DataSHIELD technical team.
We are especially grateful to Elaine Smith, Stuart Wheater, Yannick Marcon, Paul Burton, Demetris Avraam, Patricia Ryser-Welch and Wolfgang Viechtbauer for fruitful discussions and feedback.


## Contact

* Soumya Banerjee, Demetris Avraam, Paul Burton, Xavier Escriba Montagut, Juan Gonzalez, Tom R. P. Bishop and DataSHIELD technical team

*

* DataSHIELD 

    * DataSHIELD is a platform that enables the non-disclosive analysis of distributed sensitive data 

    * https://www.datashield.ac.uk
    
    
## Citation

If you use this code, please cite the following DOI

[![DOI](https://zenodo.org/badge/362161720.svg)](https://zenodo.org/badge/latestdoi/362161720)

or see the CITATION.cff file edited using

https://citation-file-format.github.io/cff-initializer-javascript/
