## Supplementary material for "dsSurvival: Privacy preserving survival models for federated individual patient meta-analysis in DataSHIELD": Client side dsSurvivalClient code: init_discordant_datasets.R

```
init.discordant.dataset.simple <- function(variables)
{
    if (ds.test_env$secure_login_details)
    {
      if (ds.test_env$driver == "OpalDriver")
      {
        builder <- DSI::newDSLoginBuilder(.silent = TRUE)
        builder$append(server = "discordant1", url = ds.test_env$ip_address_1, user = ds.test_env$user_1, password = ds.test_env$password_1, table = "DISCORDANT.DISCORDANT_STUDY1")
        builder$append(server = "discordant2", url = ds.test_env$ip_address_2, user = ds.test_env$user_2, password = ds.test_env$password_2, table = "DISCORDANT.DISCORDANT_STUDY2")
        builder$append(server = "discordant3", url = ds.test_env$ip_address_3, user = ds.test_env$user_3, password = ds.test_env$password_3, table = "DISCORDANT.DISCORDANT_STUDY3")
        ds.test_env$login.data <- builder$build()
      }
      else 
      {
          ds.test_env$login.data <- DSLite::setupDISCORDANTTest("dsBase", env = ds.test_env)
      }
      ds.test_env$stats.var <- variables
    }
}

connect.discordant.dataset.simple <- function(variables)
{
    log.out.data.server()
    source("connection_to_datasets/login_details.R")
    init.discordant.dataset.simple(variables)
    log.in.data.server()
}

disconnect.discordant.dataset.simple <- function()
{
    log.out.data.server()
}
```
