## Supplementary material for "dsSurvival: Privacy preserving survival models for federated individual patient meta-analysis in DataSHIELD": Client side dsSurvivalClient code: init_local_settings.R

```
#this file stores some settings for the continuous integration and local testing.

init.ip.address <- function()
{
   file.name <- init.local.settings()
   if (file.exists(file.name))
   {
      content <- read.csv(file.name, header = FALSE)
      ip.address <- as.character(content[[1]][1])
   }
   else
   {
      ip.address <- "127.0.0.1"
   }
   return (ip.address)
}


init.local.settings <- function()
{
  path <- getwd()
  sub.folder.name <- "/connection_to_datasets/"
  file.name <- "local_settings.csv"
  return(paste(path, sub.folder.name,file.name, sep=""))
  
}
```
