## Supplementary material for "dsSurvival: Privacy preserving survival models for federated individual patient meta-analysis in DataSHIELD": Client side dsSurvivalClient code: init_studies_datasets.R

```
init.studies.dataset.cnsim <- function(variables)
{
    if (ds.test_env$secure_login_details)
    {
      if (ds.test_env$driver == "OpalDriver")
      {
        builder <- DSI::newDSLoginBuilder(.silent = TRUE)
        builder$append(server = "sim1", url = ds.test_env$ip_address_1, user = ds.test_env$user_1, password = ds.test_env$password_1, table = "CNSIM.CNSIM1")
        builder$append(server = "sim2", url = ds.test_env$ip_address_2, user = ds.test_env$user_2, password = ds.test_env$password_2, table = "CNSIM.CNSIM2")
        builder$append(server = "sim3", url = ds.test_env$ip_address_3, user = ds.test_env$user_3, password = ds.test_env$password_3, table = "CNSIM.CNSIM3")
        ds.test_env$login.data <- builder$build()
      }
      else 
      {
         ds.test_env$login.data <- DSLite::setupCNSIMTest("dsBase", env = ds.test_env)
      }
      ds.test_env$stats.var <- variables
    }
}

init.studies.dataset.dasim <- function(variables)
{
    if (ds.test_env$secure_login_details)
    {
      if (ds.test_env$driver == "OpalDriver")
      {
        builder <- DSI::newDSLoginBuilder(.silent = TRUE)
        builder$append(server = "sim1", url = ds.test_env$ip_address_1, user = ds.test_env$user_1, password = ds.test_env$password_1, table = "DASIM.DASIM1")
        builder$append(server = "sim2", url = ds.test_env$ip_address_2, user = ds.test_env$user_2, password = ds.test_env$password_2, table = "DASIM.DASIM2")
        builder$append(server = "sim3", url = ds.test_env$ip_address_3, user = ds.test_env$user_3, password = ds.test_env$password_3, table = "DASIM.DASIM3")
        ds.test_env$login.data <- builder$build()
      }
      else 
      {
        ds.test_env$login.data <- DSLite::setupDASIMTest("dsBase", env = ds.test_env)
      }
      ds.test_env$stats.var <- variables
    }
}

init.studies.dataset.survival <- function(variables)
{
    if (ds.test_env$secure_login_details)
    {
      if (ds.test_env$driver == "OpalDriver")
      {
        builder <- DSI::newDSLoginBuilder(.silent = TRUE)
        builder$append(server = "survival1", url = ds.test_env$ip_address_1, user = ds.test_env$user_1, password = ds.test_env$password_1, table = "SURVIVAL.EXPAND_WITH_MISSING1")
        builder$append(server = "survival2", url = ds.test_env$ip_address_2, user = ds.test_env$user_2, password = ds.test_env$password_2, table = "SURVIVAL.EXPAND_WITH_MISSING2")
        builder$append(server = "survival3", url = ds.test_env$ip_address_3, user = ds.test_env$user_3, password = ds.test_env$password_3, table = "SURVIVAL.EXPAND_WITH_MISSING3")
        ds.test_env$login.data <- builder$build()
      }
      else 
      {
        ds.test_env$login.data <- DSLite::setupSURVIVALTest("dsBase", env = ds.test_env)
      }
      ds.test_env$stats.var <- variables
    }
}


init.studies.dataset.survival_nomissing <- function(variables)
{
  # this function loads the required data and variables
  # variables has list of variables to be loaded
  # function to connect to survival no missing dataset SURVIVAL.EXPAND_NO_MISSING1
  if (ds.test_env$secure_login_details)
  {
    if (ds.test_env$driver == "OpalDriver")
    {
      builder <- DSI::newDSLoginBuilder(.silent = TRUE)
      builder$append(server = "survival1", url = ds.test_env$ip_address_1, user = ds.test_env$user_1, password = ds.test_env$password_1, table = "SURVIVAL.EXPAND_NO_MISSING1")
      builder$append(server = "survival2", url = ds.test_env$ip_address_2, user = ds.test_env$user_2, password = ds.test_env$password_2, table = "SURVIVAL.EXPAND_NO_MISSING2")
      builder$append(server = "survival3", url = ds.test_env$ip_address_3, user = ds.test_env$user_3, password = ds.test_env$password_3, table = "SURVIVAL.EXPAND_NO_MISSING3")
      ds.test_env$login.data <- builder$build()
    }
    else 
    {
      ds.test_env$login.data <- DSLite::setupSURVIVALTest("dsBase", env = ds.test_env)
    }
    ds.test_env$stats.var <- variables
  }
}


add_server_side_var_survival <- function()
{
      # add server-side variables for survival analysis
      # assumes that init.studies.dataset.survival_nomissing() already called
      # make sure that the outcome is numeric 
      ds.asNumeric(x.name = "D$cens", # NOTE: not cens
                   newobj = "EVENT", datasources = ds.test_env$connections )

      ds.asNumeric(x.name = "D$survtime", # NOTE: not survtime
                  newobj = "SURVTIME", datasources = ds.test_env$connections )


      ds.asNumeric(x.name = "D$age.60", # NOTE: not survtime
                   newobj = "AGE", datasources = ds.test_env$connections )
    
      # create start time variable
      ds.asNumeric(x.name = "D$starttime",
                     newobj = "STARTTIME",
                     datasources = ds.test_env$connections)

      # create end time variable
      ds.asNumeric(x.name = "D$endtime",
                     newobj = "ENDTIME",
                     datasources = ds.test_env$connections)

      return(ds.ls(datasources = ds.test_env$connections))
    
}


init.studies.dataset.cluster.int <- function(variables)
{
  if (ds.test_env$secure_login_details)
  {
    #reading data from local files
    ds.test_env$local.values.1 <- read.csv("data_files/CLUSTER/CLUSTER_INT1.csv", header = TRUE)
    ds.test_env$local.values.2 <- read.csv("data_files/CLUSTER/CLUSTER_INT2.csv", header = TRUE)
    ds.test_env$local.values.3 <- read.csv("data_files/CLUSTER/CLUSTER_INT3.csv", header = TRUE)
    ds.test_env$local.values   <- rbind(ds.test_env$local.values.1,ds.test_env$local.values.2,ds.test_env$local.values.3)
    if (ds.test_env$driver == "OpalDriver")
    {
      builder <- DSI::newDSLoginBuilder(.silent = TRUE)
      builder$append(server = "cluster.int1", url = ds.test_env$ip_address_1, user = ds.test_env$user_1, password = ds.test_env$password_1, table = "CLUSTER.CLUSTER_INT1")
      builder$append(server = "cluster.int2", url = ds.test_env$ip_address_2, user = ds.test_env$user_2, password = ds.test_env$password_2, table = "CLUSTER.CLUSTER_INT2")
      builder$append(server = "cluster.int3", url = ds.test_env$ip_address_3, user = ds.test_env$user_3, password = ds.test_env$password_3, table = "CLUSTER.CLUSTER_INT3")
      ds.test_env$login.data <- builder$build()
    }
    else 
    {
      #to do
      #ds.test_env$login.data <- DSLite::setupCLUSTERTest("dsBase", env = ds.test_env)
    }
    ds.test_env$stats.var <- variables
  }
}

init.studies.dataset.cluster.slo <- function(variables)
{
  if (ds.test_env$secure_login_details)
  {
    #reading data from local files
    ds.test_env$local.values.1 <- read.csv("data_files/CLUSTER/CLUSTER_SLO1.csv", header = TRUE)
    ds.test_env$local.values.2 <- read.csv("data_files/CLUSTER/CLUSTER_SLO2.csv", header = TRUE)
    ds.test_env$local.values.3 <- read.csv("data_files/CLUSTER/CLUSTER_SLO3.csv", header = TRUE)
    ds.test_env$local.values   <- rbind(ds.test_env$local.values.1,ds.test_env$local.values.2,ds.test_env$local.values.3)
    if (ds.test_env$driver == "OpalDriver")
    {
      builder <- DSI::newDSLoginBuilder(.silent = TRUE)
      builder$append(server = "cluster.slo1", url = ds.test_env$ip_address_1, user = ds.test_env$user_1, password = ds.test_env$password_1, table = "CLUSTER.CLUSTER_SLO1")
      builder$append(server = "cluster.slo2", url = ds.test_env$ip_address_2, user = ds.test_env$user_2, password = ds.test_env$password_2, table = "CLUSTER.CLUSTER_SLO2")
      builder$append(server = "cluster.slo3", url = ds.test_env$ip_address_3, user = ds.test_env$user_3, password = ds.test_env$password_3, table = "CLUSTER.CLUSTER_SLO3")
      ds.test_env$login.data <- builder$build()
    }
    else 
    {
      #to do
      #ds.test_env$login.data <- DSLite::setupCLUSTERTest("dsBase", env = ds.test_env)
    }
    ds.test_env$stats.var <- variables

  }
}

init.studies.dataset.anthro <- function(variables)
{
  if (ds.test_env$secure_login_details)
  {
    #reading data from local files
    ds.test_env$local.values.1 <- read.csv("data_files/ANTHRO/anthro1.csv", header = TRUE)
    ds.test_env$local.values.2 <- read.csv("data_files/ANTHRO/anthro2.csv", header = TRUE)
    ds.test_env$local.values.3 <- read.csv("data_files/ANTHRO/anthro3.csv", header = TRUE)
    ds.test_env$local.values   <- rbind(ds.test_env$local.values.1,ds.test_env$local.values.2,ds.test_env$local.values.3)
    if (ds.test_env$driver == "OpalDriver")
    {
      builder <- DSI::newDSLoginBuilder(.silent = TRUE)
      builder$append(server = "study1", url = ds.test_env$ip_address_1, user = ds.test_env$user_1, password = ds.test_env$password_1, table = "ANTHRO.anthro1")
      builder$append(server = "study2", url = ds.test_env$ip_address_2, user = ds.test_env$user_2, password = ds.test_env$password_2, table = "ANTHRO.anthro2")
      builder$append(server = "study3", url = ds.test_env$ip_address_3, user = ds.test_env$user_3, password = ds.test_env$password_3, table = "ANTHRO.anthro3")
      ds.test_env$login.data <- builder$build()
    }
    else 
    {
      #to do
      #ds.test_env$login.data <- DSLite::setupCLUSTERTest("dsBase", env = ds.test_env)
    }
    ds.test_env$stats.var <- variables

  }
}

connect.studies.dataset.cnsim <- function(variables)
{
    log.out.data.server()
    source("connection_to_datasets/login_details.R")
    init.studies.dataset.cnsim(variables)
    log.in.data.server()
}

connect.studies.dataset.dasim <- function(variables)
{
    log.out.data.server()
    source("connection_to_datasets/login_details.R")
    init.studies.dataset.dasim(variables)
    log.in.data.server()
}

connect.studies.dataset.survival <- function(variables)
{
    log.out.data.server()
    source("connection_to_datasets/login_details.R")
    init.studies.dataset.survival(variables)
    log.in.data.server()
}

connect.studies.dataset.survival_nomissing <- function(variables)
{
  # variables is list of columns to be added 
  log.out.data.server()
  source("connection_to_datasets/login_details.R")
  init.studies.dataset.survival_nomissing(variables)
  log.in.data.server()
}


connect.studies.dataset.cluster.int <- function(variables)
{
  log.out.data.server()
  source("connection_to_datasets/login_details.R")
  init.studies.dataset.cluster.int(variables)
  log.in.data.server()
}

connect.studies.dataset.cluster.slo <- function(variables)
{
  log.out.data.server()
  source("connection_to_datasets/login_details.R")
  init.studies.dataset.cluster.slo(variables)
  log.in.data.server()
}

connect.studies.dataset.anthro <- function(variables)
{
  log.out.data.server()
  source("connection_to_datasets/login_details.R")
  init.studies.dataset.anthro(variables)
  log.in.data.server()
}

disconnect.studies.dataset.cnsim <- function()
{
    log.out.data.server()
}

disconnect.studies.dataset.dasim <- function()
{
    log.out.data.server()
}

disconnect.studies.dataset.survival <- function()
{
    log.out.data.server()
}

disconnect.studies.dataset.cluster.int <- function()
{
  log.out.data.server()
}

disconnect.studies.dataset.cluster.slo <- function()
{
  log.out.data.server()
}

disconnect.studies.dataset.anthro <- function()
{
  log.out.data.server()
}
```
