## Supplementary material for "dsSurvival: Privacy preserving survival models for federated individual patient meta-analysis in DataSHIELD": Client side dsSurvivalClient code: init_testing_datasets.R

```
# Purpose: This script provides all the functions to connect to testing data. The latter are stored 
# in data_files folder of test_that and on the virtual machine as TESTING.DATASET1, TESTING.DATASET2, 
# TESTING.DATASET3.
# Author: Patricia Ryser-Welch, DataSHIELD team

# Connect to three servers and the three datasets. Four local variables named ds.test_env$local.values.1,
# ds.test_env$local.values.2, ds.test_env$local.values.3 and ds.test_env$local.values are created.
init.testing.datasets <- function()
{
  log.out.data.server()
  if (ds.test_env$secure_login_details)
  {
    #reading data from local files 
    ds.test_env$local.values.1 <- read.csv("data_files/TESTING/DATASET1.csv", header = TRUE)
    ds.test_env$local.values.2 <- read.csv("data_files/TESTING/DATASET2.csv", header = TRUE)
    ds.test_env$local.values.3 <- read.csv("data_files/TESTING/DATASET3.csv", header = TRUE)
    ds.test_env$local.values   <- rbind(ds.test_env$local.values.1,ds.test_env$local.values.2,ds.test_env$local.values.3)
    if (ds.test_env$driver == "OpalDriver") 
    {
      builder <- DSI::newDSLoginBuilder(.silent = TRUE)
      builder$append(server = "study1", url = ds.test_env$ip_address_1, user = ds.test_env$user_1, password = ds.test_env$password_1, table = "TESTING.DATASET1")
      builder$append(server = "study2", url = ds.test_env$ip_address_2, user = ds.test_env$user_2, password = ds.test_env$password_2, table = "TESTING.DATASET2")
      builder$append(server = "study3", url = ds.test_env$ip_address_3, user = ds.test_env$user_3, password = ds.test_env$password_3, table = "TESTING.DATASET3")
      ds.test_env$login.data <- builder$build()
    }
    else 
    {
      ds.test_env$login.data <- DSLite::setupDATASETTest("dsBase", env = ds.test_env)
    }
    ds.test_env$stats.var <- list('ID','CHARACTER', 'LOGICAL','NA_VALUES','INTEGER','NULL_VALUES',
                                  'NON_NEGATIVE_INTEGER','POSITIVE_INTEGER','NEGATIVE_INTEGER',
                                  'NUMERIC', 'NON_NEGATIVE_NUMERIC','POSITIVE_NUMERIC','NEGATIVE_NUMERIC','FACTOR_CHARACTER', 
                                  'FACTOR_INTEGER','IDENTIFIER','CATEGORY','IDENTIFIER','CATEGORY', 'NUMERIC_ONE_CHANGE', 'INTEGER_ONE_CHANGE')
  }
}

# Connect to one server and the three datasets. One local variables named ds.test_env$local.values.3 is created.
init.dataset.3 <- function()
{
  log.out.data.server()
  if (ds.test_env$secure_login_details)
  {
    ds.test_env$local.values.3 <- read.csv("data_files/TESTING/DATASET3.csv", header = TRUE)
    if (ds.test_env$driver == "OpalDriver")
    {
      builder <- DSI::newDSLoginBuilder(.silent = TRUE)
      builder$append(server = "study3", url = ds.test_env$ip_address_3, user = ds.test_env$user_3, password = ds.test_env$password_3, table = "TESTING.DATASET3")
      ds.test_env$login.data <- builder$build()
    }
    else
    {
      login.data <- DSLite::setupDATASETTest("dsBase", env = ds.test_env)
      ds.test_env$login.data <- subset(login.data, server=="study3")
    }
    ds.test_env$stats.var <- list('ID','CHARACTER', 'LOGICAL','NA_VALUES','INTEGER','NULL_VALUES',
                                  'NON_NEGATIVE_INTEGER','POSITIVE_INTEGER','NEGATIVE_INTEGER',
                                  'NUMERIC', 'NON_NEGATIVE_NUMERIC','POSITIVE_NUMERIC','NEGATIVE_NUMERIC','FACTOR_CHARACTER', 
                                  'FACTOR_INTEGER','IDENTIFIER','CATEGORY','IDENTIFIER','CATEGORY', 'NUMERIC_ONE_CHANGE', 'INTEGER_ONE_CHANGE')
  }
}

# Connect to one server and the three datasets. One local variables named ds.test_env$local.values.2 is created.
init.dataset.2 <- function()
{
  log.out.data.server()
  if (ds.test_env$secure_login_details)
  {
    ds.test_env$local.values.2 <- read.csv("data_files/TESTING/DATASET2.csv", header = TRUE)
    if (ds.test_env$driver == "OpalDriver")
    {
      builder <- DSI::newDSLoginBuilder(.silent = TRUE)
      builder$append(server = "study2", url = ds.test_env$ip_address_2, user = ds.test_env$user_2, password = ds.test_env$password_2, table = "TESTING.DATASET2")
      ds.test_env$login.data <- builder$build()
    }
    else
    {
      login.data <- DSLite::setupDATASETTest("dsBase", env = ds.test_env)
      ds.test_env$login.data <- subset(login.data, server=="study2")
    } 
    ds.test_env$stats.var <- list('ID','CHARACTER', 'LOGICAL','NA_VALUES','INTEGER','NULL_VALUES',
                                  'NON_NEGATIVE_INTEGER','POSITIVE_INTEGER','NEGATIVE_INTEGER',
                                  'NUMERIC', 'NON_NEGATIVE_NUMERIC','POSITIVE_NUMERIC','NEGATIVE_NUMERIC','FACTOR_CHARACTER', 
                                  'FACTOR_INTEGER','IDENTIFIER','CATEGORY','IDENTIFIER','CATEGORY', 'NUMERIC_ONE_CHANGE', 'INTEGER_ONE_CHANGE')
  }
}

# Connect to one server and the three datasets. One local variables named ds.test_env$local.values.2is created.
init.dataset.1 <- function()
{
  log.out.data.server()
  if (ds.test_env$secure_login_details)
  {
    ds.test_env$local.values.1 <- read.csv("data_files/TESTING/DATASET1.csv", header = TRUE)
    if (ds.test_env$driver == "OpalDriver")
    {
      builder <- DSI::newDSLoginBuilder(.silent = TRUE)
      builder$append(server = "study1", url = ds.test_env$ip_address_1, user = ds.test_env$user_1, password = ds.test_env$password_1, table = "TESTING.DATASET1")
      ds.test_env$login.data <- builder$build()
    }
    else
    {
      login.data <- DSLite::setupDATASETTest("dsBase", env = ds.test_env)
      ds.test_env$login.data <- subset(login.data, server=="study1")
    }  
    ds.test_env$stats.var <- list('ID','CHARACTER', 'LOGICAL','NA_VALUES','INTEGER','NULL_VALUES',
                                  'NON_NEGATIVE_INTEGER','POSITIVE_INTEGER','NEGATIVE_INTEGER',
                                  'NUMERIC', 'NON_NEGATIVE_NUMERIC','POSITIVE_NUMERIC','NEGATIVE_NUMERIC','FACTOR_CHARACTER', 
                                  'FACTOR_INTEGER','IDENTIFIER','CATEGORY','IDENTIFIER','CATEGORY', 'NUMERIC_ONE_CHANGE', 'INTEGER_ONE_CHANGE')
  }
}

#####FACTOR_LEVELS

# Connect to one server and the two studies. One local variables named ds.test_env$local.values.2is created.
init.testing.dataset.factor_levels <- function()
{
  log.out.data.server()
  if (ds.test_env$secure_login_details)
  {
    ds.test_env$local.values.1 <- read.csv("data_files/FACTOR_LEVELS/FACTOR_LEVELS_DATASET1.csv", header = TRUE)
    ds.test_env$local.values.2 <- read.csv("data_files/FACTOR_LEVELS/FACTOR_LEVELS_DATASET2.csv", header = TRUE)
    ds.test_env$local.values.3 <- read.csv("data_files/FACTOR_LEVELS/FACTOR_LEVELS_DATASET3.csv", header = TRUE)
    ds.test_env$local.values   <- rbind(ds.test_env$local.values.1,ds.test_env$local.values.2,ds.test_env$local.values.3)
    if (ds.test_env$driver == "OpalDriver")
    {
      builder <- DSI::newDSLoginBuilder(.silent = TRUE)
      builder$append(server = "GROUP1", url = ds.test_env$ip_address_1, user = ds.test_env$user_1, password = ds.test_env$password_1, table = "FACTOR_LEVELS.FACTOR_LEVELS1")
      builder$append(server = "GROUP2", url = ds.test_env$ip_address_1, user = ds.test_env$user_1, password = ds.test_env$password_1, table = "FACTOR_LEVELS.FACTOR_LEVELS2")
      builder$append(server = "GROUP3", url = ds.test_env$ip_address_1, user = ds.test_env$user_1, password = ds.test_env$password_1, table = "FACTOR_LEVELS.FACTOR_LEVELS3")
      ds.test_env$login.data <- builder$build()
    }
    else
    {
      login.data <- DSLite::setupDATASETTest("dsBase", env = ds.test_env)
      ds.test_env$login.data <- subset(login.data, server=="study1")
    }  
    ds.test_env$stats.var <- list('ID', 'COLOURS', 'COLOURS.NUMBERS', 'POSITIVE.NUMBERS', 'NEGATIVE.NUMBERS', 'NUMBERS',
                                  'POSITIVE.DECIMAL', 'NEGATIVE.DECIMAL', 'DECIMAL', 'PLANETS.CHARACTERS')
  }
}

init.testing.dataset.factor_levels.1 <- function()
{
  log.out.data.server()
  if (ds.test_env$secure_login_details)
  {
    ds.test_env$local.values.1 <- read.csv("data_files/FACTOR_LEVELS/FACTOR_LEVELS_DATASET1.csv", header = TRUE)
    ds.test_env$local.values   <- rbind(ds.test_env$local.values.1)
    if (ds.test_env$driver == "OpalDriver")
    {
      builder <- DSI::newDSLoginBuilder(.silent = TRUE)
      builder$append(server = "GROUP1", url = ds.test_env$ip_address_1, user = ds.test_env$user_1, password = ds.test_env$password_1, table = "FACTOR_LEVELS.FACTOR_LEVELS1")
      ds.test_env$login.data <- builder$build()
    }
    else
    {
      login.data <- DSLite::setupDATASETTest("dsBase", env = ds.test_env)
      ds.test_env$login.data <- subset(login.data, server=="study1")
    }  
    ds.test_env$stats.var <- list('ID', 'COLOURS', 'COLOURS.NUMBERS', 'POSITIVE.NUMBERS', 'NEGATIVE.NUMBERS', 'NUMBERS',
                                  'POSITIVE.DECIMAL', 'NEGATIVE.DECIMAL', 'DECIMAL', 'PLANETS.CHARACTERS')
  }
}

log.in.data.server <- function()
{
  #print(ds.test_env$login.data)
  ds.test_env$connections <- datashield.login(logins=ds.test_env$login.data, assign=TRUE,variables=ds.test_env$stats.var)
}


log.out.data.server <- function()
{
  if (!is.null(ds.test_env) && !is.null(ds.test_env$connections))
  {
    datashield.logout(ds.test_env$connections)
  }
  rm(list = ls())
  gc()
}

connect.all.datasets <- function()
{
   log.out.data.server()
   source("connection_to_datasets/login_details.R")
   init.testing.datasets()
   log.in.data.server()
}

connect.dataset.1 <- function()
{
  log.out.data.server()
  source("connection_to_datasets/login_details.R")
  init.dataset.1()
  log.in.data.server()
}

connect.dataset.2 <- function()
{
  log.out.data.server()
  source("connection_to_datasets/login_details.R")
  init.dataset.2()
  log.in.data.server()
}

connect.dataset.3 <- function()
{
  log.out.data.server()
  source("connection_to_datasets/login_details.R")
  init.dataset.3()
  log.in.data.server()
}

connect.testing.dataset.factor_levels <- function()
{
  log.out.data.server()
  source("connection_to_datasets/login_details.R")
  init.testing.dataset.factor_levels()
  log.in.data.server()
}

connect.testing.dataset.factor_levels.1 <- function()
{
  log.out.data.server()
  source("connection_to_datasets/login_details.R")
  init.testing.dataset.factor_levels.1()
  log.in.data.server()
}

disconnect.all.datasets <- function()
{
    log.out.data.server()
}

disconnect.dataset.1 <- function()
{
    log.out.data.server()
}

disconnect.dataset.2 <- function()
{
    log.out.data.server()
}

disconnect.dataset.3 <- function()
{
    log.out.data.server()
}

disconnect.testing.dataset.factor_levels <- function()
{
  log.out.data.server()
}

disconnect.testing.dataset.factor_levels.1 <- function()
{
  log.out.data.server()
}
```
