## Supplementary material for "dsSurvival: Privacy preserving survival models for federated individual patient meta-analysis in DataSHIELD": Client side dsSurvivalClient code: login_details.R

```
#This script determine all the login information to the servers used for the testing. 
#The first time, you will need to edit the server_ip_address. The latter is required to set 
#access to the data on the virtual machines. 


source("connection_to_datasets/init_local_settings.R")
init.ip.address()
  ds.test_env <- new.env()
  # this option helps DSI to find the connection objects by looking in the right environment
  options(datashield.env=ds.test_env)
  ds.test_env$tolerance = 10^-6

  ds.test_env$server_ip_address = init.ip.address()
  
  #This TCP/IP address is required to test a connect to the server. 
  ds.test_env$ping_address <- paste("http://", ds.test_env$server_ip_address, ":8080", sep="" )
  
  ds.test_env$ip_address_1 <- ds.test_env$ping_address
  ds.test_env$ip_address_2 <- ds.test_env$ping_address
  ds.test_env$ip_address_3 <- ds.test_env$ping_address

  ds.test_env$user_1 <- getOption("opal.user", "administrator")
  ds.test_env$user_2 <- getOption("opal.user", "administrator")
  ds.test_env$user_3 <- getOption("opal.user", "administrator")

  ds.test_env$password_1 <- getOption("opal.password", "datashield_test&")
  ds.test_env$password_2 <- getOption("opal.password", "datashield_test&")
  ds.test_env$password_3 <- getOption("opal.password", "datashield_test&")

  # switch to "DSLiteDriver" to test with DSLite  
  # ds.test_env$driver <- "DSLiteDriver"
  ds.test_env$driver <- "OpalDriver"
  ds.test_env$secure_login_details = TRUE
```
