## Supplementary material for "dsSurvival: Privacy preserving survival models for federated individual patient meta-analysis in DataSHIELD": Client side dsSurvivalClient code: ds_expect_variables.R

```
#-------------------------------------------------------------------------------
# Copyright (c) 2019-2020 University of Newcastle upon Tyne. All rights reserved.
#
# This program and the accompanying materials
# are made available under the terms of the GNU Public License v3.0.
#
# You should have received a copy of the GNU General Public License
# along with this program.  If not, see <http://www.gnu.org/licenses/>.
#-------------------------------------------------------------------------------

#
# Tests
#

ds_expect_variables <- function(expected.variables)
{
    studies.current.varables <- ds.ls()

    for (study.current.varables in studies.current.varables) {
        expect_setequal(study.current.varables$objects.found, expected.variables)
    }
}
```
