## Supplementary material for "dsSurvival: Privacy preserving survival models for federated individual patient meta-analysis in DataSHIELD": Client side dsSurvivalClient code: setup.R

```
#-------------------------------------------------------------------------------
# Copyright (c) 2019-2020 University of Newcastle upon Tyne. All rights reserved.
#
# This program and the accompanying materials
# are made available under the terms of the GNU Public License v3.0.
#
# You should have received a copy of the GNU General Public License
# along with this program.  If not, see <http://www.gnu.org/licenses/>.
#-------------------------------------------------------------------------------
#
# Datashield test suite set up
#

context("setup - start")

library(lme4)
library(e1071)
library(DescTools)

library(DSOpal)
library(DSLite)

library(devtools)
library(testthat)
library(dsBase)
library(dsBaseClient)
require('DSI')
require('DSOpal')


source("dstest_functions/ds_expect_variables.R")
source("connection_to_datasets/login_details.R")
source("connection_to_datasets/init_testing_datasets.R")
source("connection_to_datasets/init_studies_datasets.R")
source("connection_to_datasets/init_discordant_datasets.R")

context("setup - done")
```
