## Supplementary material for "dsSurvival: Privacy preserving survival models for federated individual patient meta-analysis in DataSHIELD": Client side dsSurvivalClient code: test-_-vm-test.R

```
#-------------------------------------------------------------------------------
# Copyright (c) 2019-2020 University of Newcastle upon Tyne. All rights reserved.
#
# This program and the accompanying materials
# are made available under the terms of the GNU Public License v3.0.
#
# You should have received a copy of the GNU General Public License
# along with this program.  If not, see <http://www.gnu.org/licenses/>.
#-------------------------------------------------------------------------------

#
# Set up
#

context("vm-test::_::setup")

init.testing.datasets()

#
# Tests
#

#connect to a server
context("vm-test::_::tests::vm")

test_that("The virtual machine is loaded. ",
{
    response <- httr::HEAD(url=ds.test_env$ping_address, config=config(timeout=5))
    expect_equal(http_status(response)$reason, "OK")
})

#
# Shutdown
#

context("vm-test::_::shutdown")

#
# Done
#

context("vm-test::_::done")
```
