## Supplementary material for "dsSurvival: Privacy preserving survival models for federated individual patient meta-analysis in DataSHIELD": Client side dsSurvivalClient code: test-datachk-SURVIVAL_EXPAND_WITH_MISSING.R

```
#-------------------------------------------------------------------------------
# Copyright (c) 2019-2020 University of Newcastle upon Tyne. All rights reserved.
#
# This program and the accompanying materials
# are made available under the terms of the GNU Public License v3.0.
#
# You should have received a copy of the GNU General Public License
# along with this program.  If not, see <http://www.gnu.org/licenses/>.
#-------------------------------------------------------------------------------

#
# Set up
#

context("SURVIVAL_EXPAND_WITH_MISSING::datachk::setup")

connect.studies.dataset.survival(list('id', 'study.id', 'time.id', 'starttime', 'endtime', 'survtime', 'cens', 'age.60', 'female', 'noise.56', 'pm10.16', 'bmi.26'))

test_that("setup", {
    ds_expect_variables(c("D"))
})

#
# Tests
#

context("SURVIVAL_EXPAND_WITH_MISSING::datachk")
test_that("Check SURVIVAL_EXPAND_WITH_MISSING dataset", {
    res.class <- ds.class(x='D')
    expect_length(res.class, 3)
    expect_length(res.class$survival1, 1)
    expect_equal(res.class$survival1, "data.frame")
    expect_length(res.class$survival2, 1)
    expect_equal(res.class$survival2, "data.frame")
    expect_length(res.class$survival3, 1)
    expect_equal(res.class$survival3, "data.frame")

    res.length <- ds.length(x='D')
    expect_length(res.length, 4)
    expect_length(res.length$`length of D in survival1`, 1)
    expect_equal(res.length$`length of D in survival1`, 12)
    expect_length(res.length$`length of D in survival2`, 1)
    expect_equal(res.length$`length of D in survival2`, 12)
    expect_length(res.length$`length of D in survival3`, 1)
    expect_equal(res.length$`length of D in survival3`, 12)
    expect_equal(res.length$`total length of D in all studies combined`, 36)

    res.colnames <- ds.colnames(x='D')
    expect_length(res.colnames, 3)
    expect_length(res.colnames$survival1, 12)
    expect_equal(res.colnames$survival1, c('id', 'study.id', 'time.id', 'starttime', 'endtime', 'survtime', 'cens', 'age.60', 'female', 'noise.56', 'pm10.16', 'bmi.26'))
    expect_length(res.colnames$survival2, 12)
    expect_equal(res.colnames$survival2, c('id', 'study.id', 'time.id', 'starttime', 'endtime', 'survtime', 'cens', 'age.60', 'female', 'noise.56', 'pm10.16', 'bmi.26'))
    expect_length(res.colnames$survival3, 12)
    expect_equal(res.colnames$survival3, c('id', 'study.id', 'time.id', 'starttime', 'endtime', 'survtime', 'cens', 'age.60', 'female', 'noise.56', 'pm10.16', 'bmi.26'))

    res.class.id <- ds.class(x='D$id')
    expect_length(res.class.id, 3)
    expect_length(res.class.id$survival1, 1)
    expect_equal(res.class.id$survival1, "integer")
    expect_length(res.class.id$survival2, 1)
    expect_equal(res.class.id$survival2, "integer")
    expect_length(res.class.id$survival3, 1)
    expect_equal(res.class.id$survival3, "integer")

    res.length.id <- ds.length(x='D$id')
    expect_length(res.length.id, 4)
    expect_length(res.length.id$`length of D$id in survival1`, 1)
    expect_equal(res.length.id$`length of D$id in survival1`, 2060)
    expect_length(res.length.id$`length of D$id in survival2`, 1)
    expect_equal(res.length.id$`length of D$id in survival2`, 1640)
    expect_length(res.length.id$`length of D$id in survival3`, 1)
    expect_equal(res.length.id$`length of D$id in survival3`, 2688)
    expect_length(res.length.id$`total length of D$id in all studies combined`, 1)
    expect_equal(res.length.id$`total length of D$id in all studies combined`, 6388)

    res.class.study.id <- ds.class(x='D$study.id')
    expect_length(res.class.study.id, 3)
    expect_length(res.class.study.id$survival1, 1)
    expect_equal(res.class.study.id$survival1, "integer")
    expect_length(res.class.study.id$survival2, 1)
    expect_equal(res.class.study.id$survival2, "integer")
    expect_length(res.class.study.id$survival3, 1)
    expect_equal(res.class.study.id$survival3, "integer")

    res.length.study.id <- ds.length(x='D$study.id')
    expect_length(res.length.study.id, 4)
    expect_length(res.length.study.id$`length of D$study.id in survival1`, 1)
    expect_equal(res.length.study.id$`length of D$study.id in survival1`, 2060)
    expect_length(res.length.study.id$`length of D$study.id in survival2`, 1)
    expect_equal(res.length.study.id$`length of D$study.id in survival2`, 1640)
    expect_length(res.length.study.id$`length of D$study.id in survival3`, 1)
    expect_equal(res.length.study.id$`length of D$study.id in survival3`, 2688)
    expect_length(res.length.study.id$`total length of D$study.id in all studies combined`, 1)
    expect_equal(res.length.study.id$`total length of D$study.id in all studies combined`, 6388)

    res.class.starttime <- ds.class(x='D$starttime')
    expect_length(res.class.starttime, 3)
    expect_length(res.class.starttime$survival1, 1)
    expect_equal(res.class.starttime$survival1, "numeric")
    expect_length(res.class.starttime$survival2, 1)
    expect_equal(res.class.starttime$survival2, "numeric")
    expect_length(res.class.starttime$survival3, 1)
    expect_equal(res.class.starttime$survival3, "numeric")

    res.length.starttime <- ds.length(x='D$starttime')
    expect_length(res.length.starttime, 4)
    expect_length(res.length.starttime$`length of D$starttime in survival1`, 1)
    expect_equal(res.length.starttime$`length of D$starttime in survival1`, 2060)
    expect_length(res.length.starttime$`length of D$starttime in survival2`, 1)
    expect_equal(res.length.starttime$`length of D$starttime in survival2`, 1640)
    expect_length(res.length.starttime$`length of D$starttime in survival3`, 1)
    expect_equal(res.length.starttime$`length of D$starttime in survival3`, 2688)
    expect_length(res.length.starttime$`total length of D$starttime in all studies combined`, 1)
    expect_equal(res.length.starttime$`total length of D$starttime in all studies combined`, 6388)

    res.class.endtime <- ds.class(x='D$endtime')
    expect_length(res.class.endtime, 3)
    expect_length(res.class.endtime$survival1, 1)
    expect_equal(res.class.endtime$survival1, "numeric")
    expect_length(res.class.endtime$survival2, 1)
    expect_equal(res.class.endtime$survival2, "numeric")
    expect_length(res.class.endtime$survival3, 1)
    expect_equal(res.class.endtime$survival3, "numeric")

    res.length.endtime <- ds.length(x='D$endtime')
    expect_length(res.length.endtime, 4)
    expect_length(res.length.endtime$`length of D$endtime in survival1`, 1)
    expect_equal(res.length.endtime$`length of D$endtime in survival1`, 2060)
    expect_length(res.length.endtime$`length of D$endtime in survival2`, 1)
    expect_equal(res.length.endtime$`length of D$endtime in survival2`, 1640)
    expect_length(res.length.endtime$`length of D$endtime in survival3`, 1)
    expect_equal(res.length.endtime$`length of D$endtime in survival3`, 2688)
    expect_length(res.length.endtime$`total length of D$endtime in all studies combined`, 1)
    expect_equal(res.length.endtime$`total length of D$endtime in all studies combined`, 6388)

    res.class.survtime <- ds.class(x='D$survtime')
    expect_length(res.class.survtime, 3)
    expect_length(res.class.survtime$survival1, 1)
    expect_equal(res.class.survtime$survival1, "numeric")
    expect_length(res.class.survtime$survival2, 1)
    expect_equal(res.class.survtime$survival2, "numeric")
    expect_length(res.class.survtime$survival3, 1)
    expect_equal(res.class.survtime$survival3, "numeric")

    res.length.survtime <- ds.length(x='D$survtime')
    expect_length(res.length.survtime, 4)
    expect_length(res.length.survtime$`length of D$survtime in survival1`, 1)
    expect_equal(res.length.survtime$`length of D$survtime in survival1`, 2060)
    expect_length(res.length.survtime$`length of D$survtime in survival2`, 1)
    expect_equal(res.length.survtime$`length of D$survtime in survival2`, 1640)
    expect_length(res.length.survtime$`length of D$survtime in survival3`, 1)
    expect_equal(res.length.survtime$`length of D$survtime in survival3`, 2688)
    expect_length(res.length.survtime$`total length of D$survtime in all studies combined`, 1)
    expect_equal(res.length.survtime$`total length of D$survtime in all studies combined`, 6388)

    res.class.cens <- ds.class(x='D$cens')
    expect_length(res.class.cens, 3)
    expect_length(res.class.cens$survival1, 1)
    expect_equal(res.class.cens$survival1, "factor")
    expect_length(res.class.cens$survival2, 1)
    expect_equal(res.class.cens$survival2, "factor")
    expect_length(res.class.cens$survival3, 1)
    expect_equal(res.class.cens$survival3, "factor")

    res.length.cens <- ds.length(x='D$cens')
    expect_length(res.length.cens, 4)
    expect_length(res.length.cens$`length of D$cens in survival1`, 1)
    expect_equal(res.length.cens$`length of D$cens in survival1`, 2060)
    expect_length(res.length.cens$`length of D$cens in survival2`, 1)
    expect_equal(res.length.cens$`length of D$cens in survival2`, 1640)
    expect_length(res.length.cens$`length of D$cens in survival3`, 1)
    expect_equal(res.length.cens$`length of D$cens in survival3`, 2688)
    expect_length(res.length.cens$`total length of D$cens in all studies combined`, 1)
    expect_equal(res.length.cens$`total length of D$cens in all studies combined`, 6388)

    res.class.age.60 <- ds.class(x='D$age.60')
    expect_length(res.class.age.60, 3)
    expect_length(res.class.age.60$survival1, 1)
    expect_equal(res.class.age.60$survival1, "numeric")
    expect_length(res.class.age.60$survival2, 1)
    expect_equal(res.class.age.60$survival2, "numeric")
    expect_length(res.class.age.60$survival3, 1)
    expect_equal(res.class.age.60$survival3, "numeric")

    res.length.age.60 <- ds.length(x='D$age.60')
    expect_length(res.length.age.60, 4)
    expect_length(res.length.age.60$`length of D$age.60 in survival1`, 1)
    expect_equal(res.length.age.60$`length of D$age.60 in survival1`, 2060)
    expect_length(res.length.age.60$`length of D$age.60 in survival2`, 1)
    expect_equal(res.length.age.60$`length of D$age.60 in survival2`, 1640)
    expect_length(res.length.age.60$`length of D$age.60 in survival3`, 1)
    expect_equal(res.length.age.60$`length of D$age.60 in survival3`, 2688)
    expect_length(res.length.age.60$`total length of D$age.60 in all studies combined`, 1)
    expect_equal(res.length.age.60$`total length of D$age.60 in all studies combined`, 6388)

    res.class.female <- ds.class(x='D$female')
    expect_length(res.class.female, 3)
    expect_length(res.class.female$survival1, 1)
    expect_equal(res.class.female$survival1, "factor")
    expect_length(res.class.female$survival2, 1)
    expect_equal(res.class.female$survival2, "factor")
    expect_length(res.class.female$survival3, 1)
    expect_equal(res.class.female$survival3, "factor")

    res.length.female <- ds.length(x='D$female')
    expect_length(res.length.female, 4)
    expect_length(res.length.female$`length of D$female in survival1`, 1)
    expect_equal(res.length.female$`length of D$female in survival1`, 2060)
    expect_length(res.length.female$`length of D$female in survival2`, 1)
    expect_equal(res.length.female$`length of D$female in survival2`, 1640)
    expect_length(res.length.female$`length of D$female in survival3`, 1)
    expect_equal(res.length.female$`length of D$female in survival3`, 2688)
    expect_length(res.length.female$`total length of D$female in all studies combined`, 1)
    expect_equal(res.length.female$`total length of D$female in all studies combined`, 6388)

    res.class.noise.56 <- ds.class(x='D$noise.56')
    expect_length(res.class.noise.56, 3)
    expect_length(res.class.noise.56$survival1, 1)
    expect_equal(res.class.noise.56$survival1, "numeric")
    expect_length(res.class.noise.56$survival2, 1)
    expect_equal(res.class.noise.56$survival2, "numeric")
    expect_length(res.class.noise.56$survival3, 1)
    expect_equal(res.class.noise.56$survival3, "numeric")

    res.length.noise.56 <- ds.length(x='D$noise.56')
    expect_length(res.length.noise.56, 4)
    expect_length(res.length.noise.56$`length of D$noise.56 in survival1`, 1)
    expect_equal(res.length.noise.56$`length of D$noise.56 in survival1`, 2060)
    expect_length(res.length.noise.56$`length of D$noise.56 in survival2`, 1)
    expect_equal(res.length.noise.56$`length of D$noise.56 in survival2`, 1640)
    expect_length(res.length.noise.56$`length of D$noise.56 in survival3`, 1)
    expect_equal(res.length.noise.56$`length of D$noise.56 in survival3`, 2688)
    expect_length(res.length.noise.56$`total length of D$noise.56 in all studies combined`, 1)
    expect_equal(res.length.noise.56$`total length of D$noise.56 in all studies combined`, 6388)

    res.class.pm10.16 <- ds.class(x='D$pm10.16')
    expect_length(res.class.pm10.16, 3)
    expect_length(res.class.pm10.16$survival1, 1)
    expect_equal(res.class.pm10.16$survival1, "numeric")
    expect_length(res.class.pm10.16$survival2, 1)
    expect_equal(res.class.pm10.16$survival2, "numeric")
    expect_length(res.class.pm10.16$survival3, 1)
    expect_equal(res.class.pm10.16$survival3, "numeric")

    res.length.pm10.16 <- ds.length(x='D$pm10.16')
    expect_length(res.length.pm10.16, 4)
    expect_length(res.length.pm10.16$`length of D$pm10.16 in survival1`, 1)
    expect_equal(res.length.pm10.16$`length of D$pm10.16 in survival1`, 2060)
    expect_length(res.length.pm10.16$`length of D$pm10.16 in survival2`, 1)
    expect_equal(res.length.pm10.16$`length of D$pm10.16 in survival2`, 1640)
    expect_length(res.length.pm10.16$`length of D$pm10.16 in survival3`, 1)
    expect_equal(res.length.pm10.16$`length of D$pm10.16 in survival3`, 2688)
    expect_length(res.length.pm10.16$`total length of D$pm10.16 in all studies combined`, 1)
    expect_equal(res.length.pm10.16$`total length of D$pm10.16 in all studies combined`, 6388)

    res.class.bmi.26 <- ds.class(x='D$bmi.26')
    expect_length(res.class.bmi.26, 3)
    expect_length(res.class.bmi.26$survival1, 1)
    expect_equal(res.class.bmi.26$survival1, "numeric")
    expect_length(res.class.bmi.26$survival2, 1)
    expect_equal(res.class.bmi.26$survival2, "numeric")
    expect_length(res.class.bmi.26$survival3, 1)
    expect_equal(res.class.bmi.26$survival3, "numeric")

    res.length.bmi.26 <- ds.length(x='D$bmi.26')
    expect_length(res.length.bmi.26, 4)
    expect_length(res.length.bmi.26$`length of D$bmi.26 in survival1`, 1)
    expect_equal(res.length.bmi.26$`length of D$bmi.26 in survival1`, 2060)
    expect_length(res.length.bmi.26$`length of D$bmi.26 in survival2`, 1)
    expect_equal(res.length.bmi.26$`length of D$bmi.26 in survival2`, 1640)
    expect_length(res.length.bmi.26$`length of D$bmi.26 in survival3`, 1)
    expect_equal(res.length.bmi.26$`length of D$bmi.26 in survival3`, 2688)
    expect_length(res.length.bmi.26$`total length of D$bmi.26 in all studies combined`, 1)
    expect_equal(res.length.bmi.26$`total length of D$bmi.26 in all studies combined`, 6388)
})

#
# Tear down
#

context("SURVIVAL_EXPAND_WITH_MISSING::datachk::shutdown")

test_that("shutdown", {
    ds_expect_variables(c("D"))
})

disconnect.studies.dataset.survival()

#
# Done
#

context("SURVIVAL_EXPAND_WITH_MISSING::datachk::done")
```
