## Supplementary material for "dsSurvival: Privacy preserving survival models for federated individual patient meta-analysis in DataSHIELD": Client side dsSurvivalClient code: test-smk-ds.coxphSLMA.R

```
#-------------------------------------------------------------------------------
# Copyright (c) 2019-2020 University of Newcastle upon Tyne. All rights reserved.
#
# This program and the accompanying materials
# are made available under the terms of the GNU Public License v3.0.
#
# You should have received a copy of the GNU General Public License
# along with this program.  If not, see <http://www.gnu.org/licenses/>.
#-------------------------------------------------------------------------------

#
# Set up
#

context("ds.coxphSLMA::smk::setup")

# connect.studies.dataset.cnsim(list("LAB_TSC"))
# load survival expand no missing data
connect.studies.dataset.survival_nomissing(list("cens","survtime","time.id","female","age.60","starttime","endtime"))

test_that("setup", {
    ds_expect_variables(c("D"))
})

#connect.studies.dataset.survival(list("D"))
##init.studies.dataset.survival(list("D"))
#init.studies.dataset.survival_nomissing(list("D"))
##connect.studies.dataset.dasim(c("SURVTIME"))

######################################
# add server side survival variables
######################################
# add survival related server side variables like SURVTIME, etc.
#   need to convert these to numeric and create server side
#   variables
ls_object <- add_server_side_var_survival()
# snure that objects have been added
print(ls_object)

#
# Tests
#

              
context("ds.coxphSLMA::smk")
test_that("simple error,wrong formula", {
    
    #try( cox_object <- ds.coxph.SLMA(formula = 'survival::Surv(time=SURVTIME,event=EVENT)~D$age.60')#, dataName = 'D') 
    #, silent = FALSE)
    
    dsSurvivalClient::ds.Surv(time='STARTTIME', time2='ENDTIME', event = 'EVENT', objectname='surv_object', type='counting')
    #try(
    cox_object <- dsSurvivalClient::ds.coxph.SLMA(formula = 'surv_object~AGE')#, dataName = 'D')
    #, silent=FALSE)
    # print(cox_object$study1$call)
    # print("coeff from simple model")
    # print(cox_object$study1$coefficients[1,1])
    
    # print( datashield.errors() )
    
    # summary(cox_object)
    
    # expect_error( as.character(  ds.coxph.SLMA(formula = 'survival::Surv(time=SURVTIME,event=EVENT)~D$age', dataName = 'D')   ) )
    
    # wrong formula
    expect_error( as.character(  dsSurvivalClient::ds.coxph.SLMA(formula = 'survival::Surv(time=SURVTIME,event=EVENT)=D$age', dataName = 'D')   ) )
    
})


context("ds.coxphSLMA::smk")
test_that("simple equal test, checking coefficients", {
    
    #try( cox_object <- ds.coxph.SLMA(formula = 'survival::Surv(time=SURVTIME,event=EVENT)~D$age.60')#, dataName = 'D') 
    #, silent = FALSE)
    
    dsSurvivalClient::ds.Surv(time='STARTTIME', time2='ENDTIME', event = 'EVENT', objectname='surv_object', type='counting')
    #try(
    # cox_object <- ds.coxph.SLMA(formula = 'surv_object~AGE')#, dataName = 'D')
    #, silent=FALSE)
    # print(cox_object$study1$call)
    # print("coeff from simple model")
    # print(cox_object$study1$coefficients[1,1])
    
    # print( datashield.errors() )
    
    # summary(cox_object)
    
    coxph_model_full <- dsSurvivalClient::ds.coxph.SLMA(formula = 'surv_object~AGE')
    cat("Model coeff")
    cat(coxph_model_full$survival1$coefficients[1])
    # print(summary(coxph_model_full))
    # print(coxph_model_full$survival1)
    
    expect_equal(coxph_model_full$survival1$coefficients[1], 0.0387, tolerance = 0.0001)
    
    #print(ds.ls())
    
})


context("ds.coxphSLMA::smk")
test_that("simple summary of survival object, checking message", {
    
    try(surv_object <- dsSurvivalClient::ds.Surv(time='STARTTIME', time2='ENDTIME', event = 'EVENT', objectname='surv_object', type='counting')
    ,silent=FALSE)
    print(ds.ls())
    # coxph_model_full <- dsBaseClient::ds.coxph.SLMA(formula = 'surv_object~AGE')
    
    print(datashield.errors())
    # print(ds.summary(x = 'surv_object'))
    expect_match(as.character(ds.summary(x='surv_object')), 'Mean', ignore.case = TRUE)
    
    
})


context("ds.coxphSLMA::smk")
test_that("simple summary of survival object, checking message non-existent object error", {
    
    surv_object <- dsSurvivalClient::ds.Surv(time='STARTTIME', time2='ENDTIME', event = 'EVENT', objectname='surv_object', type='counting')
    
    coxph_model_full <- dsSurvivalClient::ds.coxph.SLMA(formula = 'surv_object~AGE')
    
    # print(ds.summary(x = 'hello'))
    # expect_match(as.character(ds.summary(x='surv_object')), 'not defined', ignore.case = TRUE)
    expect_error(as.character(ds.summary(x='hello')) )
    
    
})


context("ds.coxphSLMA::smk")
test_that("summary of Cox model, error since only summary of survival object allowed", {
    
    surv_object <- dsSurvivalClient::ds.Surv(time='STARTTIME', time2='ENDTIME', event = 'EVENT', objectname='surv_object', type='counting')
    
    coxph_model_full <- dsSurvivalClient::ds.coxph.SLMA(formula = 'surv_object~AGE')
    
    expect_error(as.character(ds.summary(x='coxph_model_full')) )
    
    
})


#
# Done
#

context("ds.coxphSLMA::smk::shutdown")

#test_that("shutdown", {
#    ds_expect_variables(c("D"))
#})

disconnect.studies.dataset.cnsim()
disconnect.studies.dataset.survival()

context("ds.coxphSLMA::smk::done")
```
