## Supplementary material for "dsSurvival: Privacy preserving survival models for federated individual patient meta-analysis in DataSHIELD": Client side dsSurvivalClient code: test-smk-ds.summary.R

```
#-------------------------------------------------------------------------------
# Copyright (c) 2019-2020 University of Newcastle upon Tyne. All rights reserved.
#
# This program and the accompanying materials
# are made available under the terms of the GNU Public License v3.0.
#
# You should have received a copy of the GNU General Public License
# along with this program.  If not, see <http://www.gnu.org/licenses/>.
#-------------------------------------------------------------------------------

#
# Set up
#

context("ds.summary::smk::setup")

connect.studies.dataset.cnsim(list('LAB_TSC', 'LAB_TRIG', 'LAB_HDL', 'LAB_GLUC_ADJUSTED', 'PM_BMI_CONTINUOUS', 'DIS_CVA', 'MEDI_LPD', 'DIS_DIAB', 'DIS_AMI', 'GENDER', 'PM_BMI_CATEGORICAL'))

test_that("setup", {
    ds_expect_variables(c("D"))
})

#
# Tests
#

context("ds.summary::smk::summary of a numerical variable")
res <- ds.summary(x='D$LAB_TSC')
test_that("summary_numerical_variable", {
  expect_equal(res$sim1$class, "numeric")
  expect_equal(res$sim2$length, 3088)
  expect_equal(res$sim3$`quantiles & mean`[[4]], 5.786)
})

context("ds.summary::smk::summary of a character variable")
ds.asCharacter(x='D$GENDER', newobj="a_character")
res <- ds.summary(x='a_character')
test_that("summary_character_variable", {
  expect_equal(res$sim1$class, "character")
  expect_equal(res$sim2$length, 3088)
})

context("ds.summary::smk::summary of a factor variable")
ds.asFactor('D$PM_BMI_CATEGORICAL', newobj="a_factor")
res <- ds.summary(x='a_factor')
test_that("summary_factor_variable", {
  expect_length(res, 3)
  expect_length(res$sim1, 6)
  expect_equal(res$sim1$class, "factor")
  expect_equal(res$sim1$length, 2163)
  expect_equal(res$sim1$categories, c("1", "2", "3"))
  expect_equal(res$sim1$`count of '1'`, 641)
  expect_equal(res$sim1$`count of '2'`, 816)
  expect_equal(res$sim1$`count of '3'`, 609)
  expect_length(res$sim2, 6)
  expect_equal(res$sim2$class, "factor")
  expect_equal(res$sim2$length, 3088)
  expect_equal(res$sim2$categories, c("1", "2", "3"))
  expect_equal(res$sim2$`count of '1'`, 899)
  expect_equal(res$sim2$`count of '2'`, 1173)
  expect_equal(res$sim2$`count of '3'`, 866)
  expect_length(res$sim3, 6)
  expect_equal(res$sim3$class, "factor")
  expect_equal(res$sim3$length, 4128)
  expect_equal(res$sim3$categories, c("1", "2", "3"))
  expect_equal(res$sim3$`count of '1'`, 1213)
  expect_equal(res$sim3$`count of '2'`, 1556)
  expect_equal(res$sim3$`count of '3'`, 1154)
})

context("ds.summary::smk::summary of a list variable")
ds.asList(x.name='D$PM_BMI_CATEGORICAL', newobj="a_list")
res <- ds.summary(x='a_list')
test_that("summary_list_variable", {
  expect_length(res, 3)
  expect_length(res$sim1, 2)
  expect_equal(res$sim1$class, "list")
  expect_equal(res$sim1$length, 2163)
  expect_length(res$sim2, 2)
  expect_equal(res$sim2$class, "list")
  expect_equal(res$sim2$length, 3088)
  expect_length(res$sim3, 2)
  expect_equal(res$sim3$class, "list")
  expect_equal(res$sim3$length, 4128)
})

context("ds.summary::smk::summary of a data frame")
res <- ds.summary(x='D')
test_that("summary_data_frame", {
  expect_equal(res$sim1$class, "data.frame")
  expect_equal(res$sim2$`number of rows`, 3088)
  expect_equal(res$sim2$`number of columns`, 11)
  expect_equal(res$sim3$`variables held`[[11]], "PM_BMI_CATEGORICAL")
  expect_equal(res$sim1$`variables held`[[2]], "LAB_TRIG")
})

#
# Tear down
#

context("ds.summary::smk::teardown")

test_that("shutdown", {
    ds_expect_variables(c("D", "a_character", "a_factor", "a_list"))
})

disconnect.studies.dataset.cnsim()

context("ds.summary::smk::done")
```
