## Supplementary material for "dsSurvival: Privacy preserving survival models for federated individual patient meta-analysis in DataSHIELD": Client side dsSurvivalClient code: development_plan.html

Meta-analysis of survival models in the DataSHIELD platform


### Meta-analysis of survival models in the DataSHIELD platform

###### Soumya Banerjee, Tom Bishop and DataSHIELD technical team

###### 23 June 2021

- Summary
- Survival analysis in DataSHIELD
- Installation
- Computational workflow
- Creating server-side variables for survival analysis
- Create survival object and call ds.coxph.SLMA()
- Summary of survival objects
- Diagnostics for Cox proportional hazards models
- Meta-analyze hazard ratios
- Plotting of privacy-preserving survival curves
- Acknowledgements
- References

### Summary

This is a document that outlines a vignette for implementing survival models and meta-analyzing hazard ratios in the DataSHIELD platform.

### Survival analysis in DataSHIELD

We outline code for implementing survival models and meta-analysis of hazard ratios in DataSHIELD.

All code is available here:

- https://github.com/neelsoumya/dsSurvival
- https://github.com/neelsoumya/dsSurvivalClient
- https://github.com/neelsoumya/dsBase
- https://github.com/neelsoumya/dsBaseClient

### Installation

Install R Studio and the development environment as described below:

- https://data2knowledge.atlassian.net/wiki/spaces/DSDEV/pages/12943461/Getting+started

Then install the virtual machines as described below:

- https://data2knowledge.atlassian.net/wiki/spaces/DSDEV/pages/931069953/Installation+Training+Hub-+DataSHIELD+v6

Install the necessary packages by running the following commands in R Studio:

```
install.packages('devtools')

library(devtools)

devtools::install_github('neelsoumya/dsBaseClient')
    
devtools::install_github('neelsoumya/dsBase')

devtools::install_github('neelsoumya/dsSurvivalClient')
```

Install dsBase (neelsoumya/dsBase main branch) and dsSurvival (neelsoumya/dsSurvival main branch) on the Opal virtual machine.

### Computational workflow

The computational steps are outlined below. The first step is connecting to the server and loading the survival data. We assume that the reader is familiar with these details.

```
library(knitr)
library(rmarkdown)
library(tinytex)
library(survival)
library(metafor)
library(ggplot2)
library(survminer)
library(dsSurvivalClient)
require('DSI')
require('DSOpal')
require('dsBaseClient')

builder <- DSI::newDSLoginBuilder()

builder$append(server = "study1", 
               url = "http://192.168.56.100:8080/", 
               user = "administrator", password = "datashield_test&", 
               table = "SURVIVAL.EXPAND_NO_MISSING1", driver = "OpalDriver")
builder$append(server = "study2", 
               url = "http://192.168.56.100:8080/", 
               user = "administrator", password = "datashield_test&", 
               table = "SURVIVAL.EXPAND_NO_MISSING2", driver = "OpalDriver")
builder$append(server = "study3", 
               url = "http://192.168.56.100:8080/", 
               user = "administrator", password = "datashield_test&", 
               table = "SURVIVAL.EXPAND_NO_MISSING3", driver = "OpalDriver")

logindata <- builder$build()

connections <- DSI::datashield.login(logins = logindata, assign = TRUE, symbol = "D")
```

### Creating server-side variables for survival analysis

We now outline some steps for analysing survival data.

- make sure that the outcome variable is numeric

```
ds.asNumeric(x.name = "D$cens",
             newobj = "EVENT",
             datasources = connections)

ds.asNumeric(x.name = "D$survtime",
             newobj = "SURVTIME",
             datasources = connections)
```

- convert time id variable to a factor

```
ds.asFactor(input.var.name = "D$time.id",
            newobj = "TID",
            datasources = connections)
```

- create in the server-side the log(survtime) variable

```
ds.log(x = "D$survtime",
       newobj = "log.surv",
       datasources = connections)
```

- create start time variable

```
ds.asNumeric(x.name = "D$starttime",
             newobj = "STARTTIME",
             datasources = connections)

ds.asNumeric(x.name = "D$endtime",
             newobj = "ENDTIME",
             datasources = connections)
```

### Create survival object and call ds.coxph.SLMA()

- use constructed Surv object in *ds.coxph.SLMA()*

```
dsSurvivalClient::ds.Surv(time='STARTTIME', time2='ENDTIME', 
                      event = 'EVENT', objectname='surv_object',
                      type='counting')

coxph_model_full <- dsSurvivalClient::ds.coxph.SLMA(formula = 'surv_object~D$age+D$female')
```

- use direct inline call to *survival::Surv()*

```
dsSurvivalClient::ds.coxph.SLMA(formula = 'survival::Surv(time=SURVTIME,event=EVENT)~D$age+D$female', 
                                dataName = 'D', 
                                datasources = connections)
```

- call with *survival::strata()*

```
coxph_model_strata <- dsSurvivalClient::ds.coxph.SLMA(formula = 'surv_object~D$age + 
                          survival::strata(D$female)')

summary(coxph_model_strata)
```

### Summary of survival objects

We can also summarize a server-side object of type *survival::Surv()* using a call to *ds.coxphSummary()*. This will provide a non-disclosive summary of the server-side object. An example call is shown below:

```
dsSurvivalClient::ds.coxphSummary(x = 'coxph_serverside')
```

### Diagnostics for Cox proportional hazards models

We have also created functions to test for the assumptions of Cox proportional hazards models.

```
dsSurvivalClient::ds.coxphSLMAassign(formula = 'surv_object~D$age+D$female',
                            objectname = 'coxph_serverside')

dsSurvivalClient::ds.cox.zphSLMA(fit = 'coxph_serverside')

dsSurvivalClient::ds.coxphSummary(x = 'coxph_serverside')
```

A diagnostic summary is shown below.

```
## surv_object~D$age+D$female
```

```
## NULL
```

```
## $study1
##          chisq df    p
## D$age    1.022  1 0.31
## D$female 0.364  1 0.55
## GLOBAL   1.239  2 0.54
## 
## $study2
##              chisq df    p
## D$age    -1389.472  1 1.00
## D$female     0.591  1 0.44
## GLOBAL    -857.492  2 1.00
## 
## $study3
##          chisq df       p
## D$age    15.27  1 9.3e-05
## D$female  8.04  1  0.0046
## GLOBAL   23.31  2 8.7e-06
```

```
## $study1
## Call:
## survival::coxph(formula = formula, data = dataTable, weights = weights, 
##     ties = ties, singular.ok = singular.ok, model = model, x = x, 
##     y = y)
## 
##   n= 2060, number of events= 426 
## 
##                coef exp(coef)  se(coef)      z Pr(>|z|)    
## D$age      0.041609  1.042487  0.003498 11.894  < 2e-16 ***
## D$female1 -0.660002  0.516850  0.099481 -6.634 3.26e-11 ***
## ---
## Signif. codes:  0 '***' 0.001 '**' 0.01 '*' 0.05 '.' 0.1 ' ' 1
## 
##           exp(coef) exp(-coef) lower .95 upper .95
## D$age        1.0425     0.9592    1.0354    1.0497
## D$female1    0.5169     1.9348    0.4253    0.6281
## 
## Concordance= 0.676  (se = 0.014 )
## Likelihood ratio test= 170.7  on 2 df,   p=<2e-16
## Wald test            = 168.2  on 2 df,   p=<2e-16
## Score (logrank) test = 166.3  on 2 df,   p=<2e-16
## 
## 
## $study2
## Call:
## survival::coxph(formula = formula, data = dataTable, weights = weights, 
##     ties = ties, singular.ok = singular.ok, model = model, x = x, 
##     y = y)
## 
##   n= 1640, number of events= 300 
## 
##               coef exp(coef) se(coef)      z Pr(>|z|)    
## D$age      0.04067   1.04151  0.00416  9.776  < 2e-16 ***
## D$female1 -0.62756   0.53389  0.11767 -5.333 9.66e-08 ***
## ---
## Signif. codes:  0 '***' 0.001 '**' 0.01 '*' 0.05 '.' 0.1 ' ' 1
## 
##           exp(coef) exp(-coef) lower .95 upper .95
## D$age        1.0415     0.9601    1.0331    1.0500
## D$female1    0.5339     1.8730    0.4239    0.6724
## 
## Concordance= 0.674  (se = 0.017 )
## Likelihood ratio test= 117.8  on 2 df,   p=<2e-16
## Wald test            = 115.2  on 2 df,   p=<2e-16
## Score (logrank) test = 116.4  on 2 df,   p=<2e-16
## 
## 
## $study3
## Call:
## survival::coxph(formula = formula, data = dataTable, weights = weights, 
##     ties = ties, singular.ok = singular.ok, model = model, x = x, 
##     y = y)
## 
##   n= 2688, number of events= 578 
## 
##                coef exp(coef)  se(coef)      z Pr(>|z|)    
## D$age      0.042145  1.043045  0.003086 13.655  < 2e-16 ***
## D$female1 -0.599238  0.549230  0.084305 -7.108 1.18e-12 ***
## ---
## Signif. codes:  0 '***' 0.001 '**' 0.01 '*' 0.05 '.' 0.1 ' ' 1
## 
##           exp(coef) exp(-coef) lower .95 upper .95
## D$age        1.0430     0.9587    1.0368    1.0494
## D$female1    0.5492     1.8207    0.4656    0.6479
## 
## Concordance= 0.699  (se = 0.011 )
## Likelihood ratio test= 227.9  on 2 df,   p=<2e-16
## Wald test            = 228.4  on 2 df,   p=<2e-16
## Score (logrank) test = 229.4  on 2 df,   p=<2e-16
```

### Meta-analyze hazard ratios

We now outline how the hazard ratios from the survival models are meta-analyzed. We use the *metafor* package for meta-analysis. We show the summary of an example meta-analysis and a forest plot below. The forest plot shows a basic example of meta-analyzed hazard ratios from a survival model (analyzed in dsSurvivalClient).

The log-hazard ratios and their standard errors from each study can be found after running *ds.coxphSLMA()*

The hazard ratios can then be meta-analyzed:

```
input_logHR = c(coxph_model_full$study1$coefficients[1,2], 
        coxph_model_full$study2$coefficients[1,2], 
        coxph_model_full$study3$coefficients[1,2])

input_se    = c(coxph_model_full$study1$coefficients[1,3], 
        coxph_model_full$study2$coefficients[1,3], 
        coxph_model_full$study3$coefficients[1,3])

metafor::rma(log_hazard_ratio, sei = se_hazard_ratio, method = 'REML')
```

A summary of this meta-analyzed model is shown below.

```
## 
## Random-Effects Model (k = 3; tau^2 estimator: REML)
## 
##   logLik  deviance       AIC       BIC      AICc 
##   9.3824  -18.7648  -14.7648  -17.3785   -2.7648   
## 
## tau^2 (estimated amount of total heterogeneity): 0 (SE = 0.0000)
## tau (square root of estimated tau^2 value):      0
## I^2 (total heterogeneity / total variability):   0.00%
## H^2 (total variability / sampling variability):  1.00
## 
## Test for Heterogeneity:
## Q(df = 2) = 0.0880, p-val = 0.9569
## 
## Model Results:
## 
## estimate      se      zval    pval   ci.lb   ci.ub 
##   1.0425  0.0020  515.4456  <.0001  1.0385  1.0465  *** 
## 
## ---
## Signif. codes:  0 '***' 0.001 '**' 0.01 '*' 0.05 '.' 0.1 ' ' 1
```

We now show a forest plot with the meta-analyzed hazard ratios. The hazard ratios come from the dsSurvivalClient function *ds.coxphSLMA()*. The hazard ratios are meta-analyzed using the *metafor* package.

Example forest plot of meta-analyzed hazard ratios.

### Plotting of privacy-preserving survival curves

We also plot privacy preserving survival curves. Please note that is work in progress and is only available on a separate development branch. There will be a full release in v1.1.0.

```
dsSurvivalClient::ds.survfit(formula='surv_object~1', objectname='survfit_object')

dsSurvivalClient::ds.plotsurvfit(formula = 'survfit_object')
```

```
## NULL
```

Privacy preserving survival curves.

```
## $study1
## Call: survfit(formula = formula)
## 
## records       n  events  median 0.95LCL 0.95UCL 
## 2060.00  886.00  426.00    5.29    3.25    7.17 
## 
## $study2
## Call: survfit(formula = formula)
## 
## records       n  events  median 0.95LCL 0.95UCL 
## 1640.00  659.00  300.00    6.83    4.76    9.04 
## 
## $study3
## Call: survfit(formula = formula)
## 
## records       n  events  median 0.95LCL 0.95UCL 
## 2688.00 1167.00  578.00    4.30    2.58    6.36
```

### Acknowledgements

We acknowledge the help and support of the DataSHIELD technical team. We are especially grateful to Yannick Marcon, Paul Burton, Demetris Avraam, Stuart Wheater, Patricia Ryser-Welch, Xavier Escriba, Juan Gonzalez and Wolfgang Vichtbauer for fruitful discussions and feedback.

### References

- https://github.com/datashield
- http://www.metafor-project.org
- https://github.com/neelsoumya/dsBase
- https://github.com/neelsoumya/dsBaseClient
- https://github.com/neelsoumya/dsSurvival
- https://github.com/neelsoumya/dsSurvivalClient
- https://github.com/neelsoumya/datashield\_testing\_basic
