## Supplementary material for "dsSurvival: Privacy preserving survival models for federated individual patient meta-analysis in DataSHIELD": Client side dsSurvivalClient code: development_plan.rmd

````
---
title: "Meta-analysis of survival models in the DataSHIELD platform"
author: "Soumya Banerjee, Tom Bishop and DataSHIELD technical team "
date: "23 June 2021"
output:
  pdf_document:
    toc: yes
    number_sections: yes
    fig_caption: yes
  html_document:
    toc: yes
    df_print: paged
urlcolor: blue
---

```{r setup, include=FALSE}
knitr::opts_chunk$set(echo=TRUE)
```

# Summary

This is a document that outlines a vignette for implementing survival models
and meta-analyzing hazard ratios in the DataSHIELD platform.

# Survival analysis in DataSHIELD

We outline code for implementing survival models and meta-analysis
of hazard ratios in DataSHIELD.

All code is available here:

   * https://github.com/neelsoumya/dsSurvival

   * https://github.com/neelsoumya/dsSurvivalClient 

   * https://github.com/neelsoumya/dsBase

   * https://github.com/neelsoumya/dsBaseClient
    
   
# Installation   

Install R Studio and the development environment as described below:

   * https://data2knowledge.atlassian.net/wiki/spaces/DSDEV/pages/12943461/Getting+started


Then install the virtual machines as described below:

   * https://data2knowledge.atlassian.net/wiki/spaces/DSDEV/pages/931069953/Installation+Training+Hub-+DataSHIELD+v6


Install the necessary packages by running the following commands in R Studio:


```

install.packages('devtools')

library(devtools)

devtools::install_github('neelsoumya/dsBaseClient')
	
devtools::install_github('neelsoumya/dsBase')

devtools::install_github('neelsoumya/dsSurvivalClient')

```

Install dsBase (neelsoumya/dsBase main branch) and dsSurvival (neelsoumya/dsSurvival main branch) on the Opal virtual machine.


# Computational workflow

The computational steps are outlined below. 
The first step is connecting to the server and loading the survival data.
We assume that the reader is familiar with these details.

```

library(knitr)
library(rmarkdown)
library(tinytex)
library(survival)
library(metafor)
library(ggplot2)
library(survminer)
library(dsSurvivalClient)
require('DSI')
require('DSOpal')
require('dsBaseClient')

builder <- DSI::newDSLoginBuilder()

builder$append(server = "study1", 
               url = "http://192.168.56.100:8080/", 
               user = "administrator", password = "datashield_test&", 
               table = "SURVIVAL.EXPAND_NO_MISSING1", driver = "OpalDriver")
builder$append(server = "study2", 
               url = "http://192.168.56.100:8080/", 
               user = "administrator", password = "datashield_test&", 
               table = "SURVIVAL.EXPAND_NO_MISSING2", driver = "OpalDriver")
builder$append(server = "study3", 
               url = "http://192.168.56.100:8080/", 
               user = "administrator", password = "datashield_test&", 
               table = "SURVIVAL.EXPAND_NO_MISSING3", driver = "OpalDriver")

logindata <- builder$build()

connections <- DSI::datashield.login(logins = logindata, assign = TRUE, symbol = "D") 


```


```{r, include = FALSE}

####################
# Load library
####################
library(knitr)
library(rmarkdown)
library(tinytex)
library(survival)
library(metafor)
library(ggplot2)
library(survminer)
#library(dsSurvival)
library(dsSurvivalClient)
require('DSI')
require('DSOpal')
require('dsBaseClient')

#######################
# Get data
#######################
builder <- DSI::newDSLoginBuilder()

# TODO: use open server
# builder$append(server="server1", url="https://opal-sandbox.mrc-epid.cam.ac.uk",
#                user="dsuser", password="password", 
#               table = "SURVIVAL.EXPAND_NO_MISSING1")

# builder$append(server="server1", url="https://opal-sandbox.mrc-epid.cam.ac.uk",
#               user="dsuser", password="password", 
#               table = "SURVIVAL.EXPAND_NO_MISSING2")
	       
# builder$append(server="server1", url="https://opal-sandbox.mrc-epid.cam.ac.uk",
#               user="dsuser", password="password", 
#               table = "SURVIVAL.EXPAND_NO_MISSING3")	       


builder$append(server = "study1", 
               url = "http://192.168.56.100:8080/", 
               user = "administrator", password = "datashield_test&", 
               table = "SURVIVAL.EXPAND_NO_MISSING1", driver = "OpalDriver")
builder$append(server = "study2", 
               url = "http://192.168.56.100:8080/", 
               user = "administrator", password = "datashield_test&", 
               table = "SURVIVAL.EXPAND_NO_MISSING2", driver = "OpalDriver")
builder$append(server = "study3", 
               url = "http://192.168.56.100:8080/", 
               user = "administrator", password = "datashield_test&", 
               table = "SURVIVAL.EXPAND_NO_MISSING3", driver = "OpalDriver")

logindata <- builder$build()

##############
# login
##############

# Log onto the remote Opal training servers
connections <- DSI::datashield.login(logins = logindata, assign = TRUE, symbol = "D") 

```

# Creating server-side variables for survival analysis

We now outline some steps for analysing survival data.

* make sure that the outcome variable is numeric 

```
ds.asNumeric(x.name = "D$cens",
             newobj = "EVENT",
             datasources = connections)

ds.asNumeric(x.name = "D$survtime",
             newobj = "SURVTIME",
             datasources = connections)
```

*  convert time id variable to a factor 

```

ds.asFactor(input.var.name = "D$time.id",
            newobj = "TID",
            datasources = connections)

```

*  create in the server-side the log(survtime) variable

```

ds.log(x = "D$survtime",
       newobj = "log.surv",
       datasources = connections)

```

*  create start time variable

```

ds.asNumeric(x.name = "D$starttime",
             newobj = "STARTTIME",
             datasources = connections)

ds.asNumeric(x.name = "D$endtime",
             newobj = "ENDTIME",
             datasources = connections)

```


```{r, include=FALSE}

# make sure that the outcome is numeric 
ds.asNumeric(x.name = "D$cens",
             newobj = "EVENT",
             datasources = connections)

ds.asNumeric(x.name = "D$survtime",
             newobj = "SURVTIME",
             datasources = connections)

# convert time id variable to a factor 
ds.asFactor(input.var.name = "D$time.id",
            newobj = "TID",
            datasources = connections)

# create in the server-side the log(survtime) variable
ds.log(x = "D$survtime",
       newobj = "log.surv",
       datasources = connections)

# create start time variable
ds.asNumeric(x.name = "D$starttime",
             newobj = "STARTTIME",
             datasources = connections)

# create end time variable
ds.asNumeric(x.name = "D$endtime",
             newobj = "ENDTIME",
             datasources = connections)


# check which variables exist
dsBaseClient::ds.ls()

# call new function modify NAMESPACE and DESCRIPTION
# call coxph server side
# client side function is here:
# https://github.com/neelsoumya/dsBaseClient/blob/absolute_newbie_client/R/ds.coxph.SLMA.R
# server side function is here:
# https://github.com/neelsoumya/dsBase/blob/absolute_newbie/R/coxphSLMADS.R
```


# Create survival object and call ds.coxph.SLMA()


* use constructed Surv object in *ds.coxph.SLMA()*

```
dsSurvivalClient::ds.Surv(time='STARTTIME', time2='ENDTIME', 
                      event = 'EVENT', objectname='surv_object',
                      type='counting')

coxph_model_full <- dsSurvivalClient::ds.coxph.SLMA(formula = 'surv_object~D$age+D$female')

```

* use direct inline call to *survival::Surv()*

```

dsSurvivalClient::ds.coxph.SLMA(formula = 'survival::Surv(time=SURVTIME,event=EVENT)~D$age+D$female', 
                                dataName = 'D', 
                                datasources = connections)

```

* call with *survival::strata()*

```

coxph_model_strata <- dsSurvivalClient::ds.coxph.SLMA(formula = 'surv_object~D$age + 
                          survival::strata(D$female)')

summary(coxph_model_strata)

```


```{r, include=FALSE}

# 1. use constructed surv object in coxph
dsSurvivalClient::ds.Surv(time='STARTTIME', time2='ENDTIME', event = 'EVENT', objectname='surv_object', type='counting')
coxph_model_full <- dsSurvivalClient::ds.coxph.SLMA(formula = 'surv_object~D$age+D$female')

# 2. use direct inline call
dsSurvivalClient::ds.coxph.SLMA(formula = 'survival::Surv(time=SURVTIME,event=EVENT)~D$age+D$female', dataName = 'D', datasources = connections)

# 3. Call with strata()
coxph_model_strata <- dsSurvivalClient::ds.coxph.SLMA(formula = 'surv_object~D$age+survival::strata(D$female)')
summary(coxph_model_strata)

```

# Summary of survival objects

We can also summarize a server-side object of type *survival::Surv()*
using a call to *ds.coxphSummary()*. This will provide a non-disclosive
summary of the server-side object. An example call is shown below:


```

dsSurvivalClient::ds.coxphSummary(x = 'coxph_serverside')

```


```{r, include=FALSE}

#################################
# summary of coxphSLMA
#################################
# TODO:
# dsBaseClient::ds.summary(x = 'surv_object')
# dsBaseClient::ds.class(x = 'surv_object')
# dsBaseClient::ds.mean(x='surv_object')


#################################
# TODO: Plot survival curves
#################################
# fit <- survival::survfit(formula = 'surv_object~D$age+D$female', data = 'D')
# need ds.survfit() and survfitDS()
# fit_model <- ds.survfit(coxph_model[1])
# plot(fit_model)

# TODO:
# plot(survfit_km, fun="cloglog")

# TODO: 
# ggplot like functionality see other functions

# In dsBaseClient::
# ds.survfit()
#         datashield.aggregate("survfitDS", ....)
#          return (the fit model)

# In dsBase::
# survfitDS(coxph_model)  
#               fit_model <- survival::survfit(coxph_model, newdata = 'D')
#               return (fit_model)

# TODO: dsSurvivalClient::ds.survfit(formula='surv_object~1', objectname='survfit_object')
# verify that object has been created
dsBaseClient::ds.ls()

# ERROR summary of survfit not allowed
# dsBaseClient::ds.summary(x='survfit_object')
# dsBaseClient::ds.mean(x='survfit_object')

```


# Diagnostics for Cox proportional hazards models

We have also created functions to test for the assumptions of 
Cox proportional hazards models.

```
dsSurvivalClient::ds.coxphSLMAassign(formula = 'surv_object~D$age+D$female',
                            objectname = 'coxph_serverside')

dsSurvivalClient::ds.cox.zphSLMA(fit = 'coxph_serverside')

dsSurvivalClient::ds.coxphSummary(x = 'coxph_serverside')

```

A diagnostic summary is shown below. 

```{r, echo=FALSE}

dsSurvivalClient::ds.coxphSLMAassign(formula = 'surv_object~D$age+D$female', objectname = 'coxph_serverside')

dsSurvivalClient::ds.cox.zphSLMA(fit = 'coxph_serverside')

dsSurvivalClient::ds.coxphSummary(x = 'coxph_serverside')

```


# Meta-analyze hazard ratios

We now outline how the hazard ratios from the survival models are meta-analyzed. 
We use the *metafor* package for meta-analysis. We show the summary of an 
example meta-analysis and a forest plot below. The forest plot shows a basic
example of meta-analyzed hazard ratios from a survival model (analyzed in
dsSurvivalClient).

The log-hazard ratios and their standard errors from each study can be
found after running *ds.coxphSLMA()*

The hazard ratios can then be meta-analyzed:

```

input_logHR = c(coxph_model_full$study1$coefficients[1,2], 
		coxph_model_full$study2$coefficients[1,2], 
		coxph_model_full$study3$coefficients[1,2])

input_se    = c(coxph_model_full$study1$coefficients[1,3], 
		coxph_model_full$study2$coefficients[1,3], 
		coxph_model_full$study3$coefficients[1,3])

metafor::rma(log_hazard_ratio, sei = se_hazard_ratio, method = 'REML')

```

A summary of this meta-analyzed model is shown below.


```{r, echo=FALSE}

# TODO: for each study
for (i_temp_counter in c(1:length(coxph_model_full)))
{
      
}

# list of hazard ratios for first parameter (age) over 3 studies 
input_logHR = c(coxph_model_full$study1$coefficients[1,2], coxph_model_full$study2$coefficients[1,2], coxph_model_full$study3$coefficients[1,2])
input_se    = c(coxph_model_full$study1$coefficients[1,3], coxph_model_full$study2$coefficients[1,3], coxph_model_full$study3$coefficients[1,3])

meta_model <- metafor::rma(input_logHR, sei = input_se, method = 'REML')
summary(meta_model)

#######################################################
# forest plots of final meta-analyzed hazard ratios
#######################################################
```

We now show a forest plot with the meta-analyzed hazard ratios. The hazard ratios come
from the dsSurvivalClient function *ds.coxphSLMA()*. The hazard ratios are meta-analyzed
using the *metafor* package.

```{r fig.cap="Example forest plot of meta-analyzed hazard ratios.", echo=FALSE}

metafor::forest.rma(x = meta_model)

```    

# Plotting of privacy-preserving survival curves

We also plot privacy preserving survival curves. Please note that is work in progress and is only available on a separate development branch. There will be a full release in v1.1.0.

```
dsSurvivalClient::ds.survfit(formula='surv_object~1', objectname='survfit_object')

dsSurvivalClient::ds.plotsurvfit(formula = 'survfit_object')

```


```{r, echo = FALSE}


dsSurvivalClient::ds.survfit(formula='surv_object~1', objectname='survfit_object')


```

```{r fig.cap="Privacy preserving survival curves.", echo = FALSE}

dsSurvivalClient::ds.plotsurvfit(formula = 'survfit_object')

```

\newpage
   
```{r, include=FALSE}

#############################################
# disconnect
#############################################
DSI::datashield.logout(conns = connections)

```


# Acknowledgements

We acknowledge the help and support of the DataSHIELD technical team.
We are especially grateful to Yannick Marcon, Paul Burton, Demetris Avraam, Stuart Wheater, Patricia Ryser-Welch, Xavier Escriba, Juan Gonzalez and Wolfgang Vichtbauer for fruitful discussions and feedback.


# References

  * https://github.com/datashield
  
  * http://www.metafor-project.org
  
  * https://github.com/neelsoumya/dsBase

  * https://github.com/neelsoumya/dsBaseClient

  * https://github.com/neelsoumya/dsSurvival

  * https://github.com/neelsoumya/dsSurvivalClient
  
  * https://github.com/neelsoumya/datashield_testing_basic
  
  
````
