## Supplementary figures and images for "dsSurvival: Privacy preserving survival models for federated individual patient meta-analysis in DataSHIELD"

### Capture_VM_install_screenshot.PNG

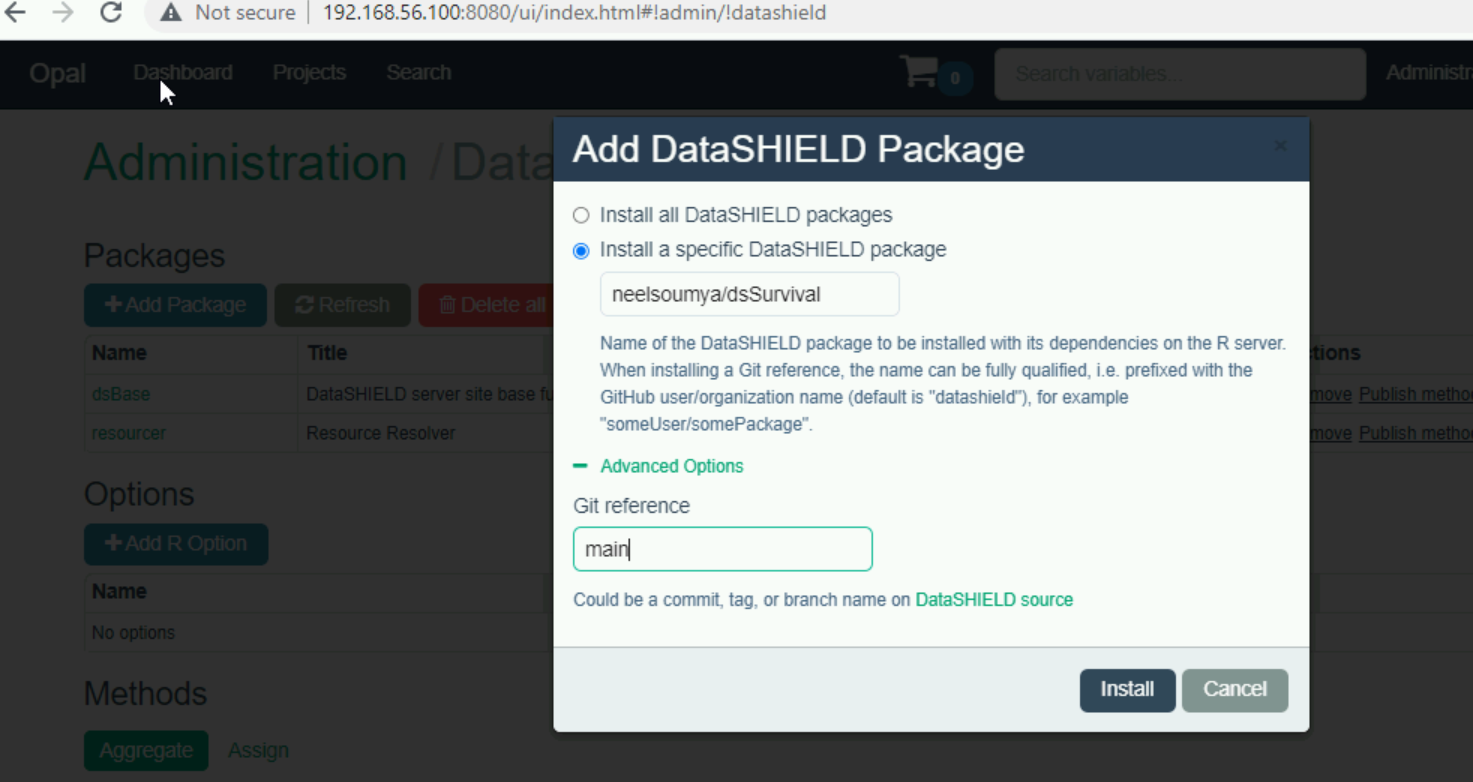

### hex-dsSurvival.png

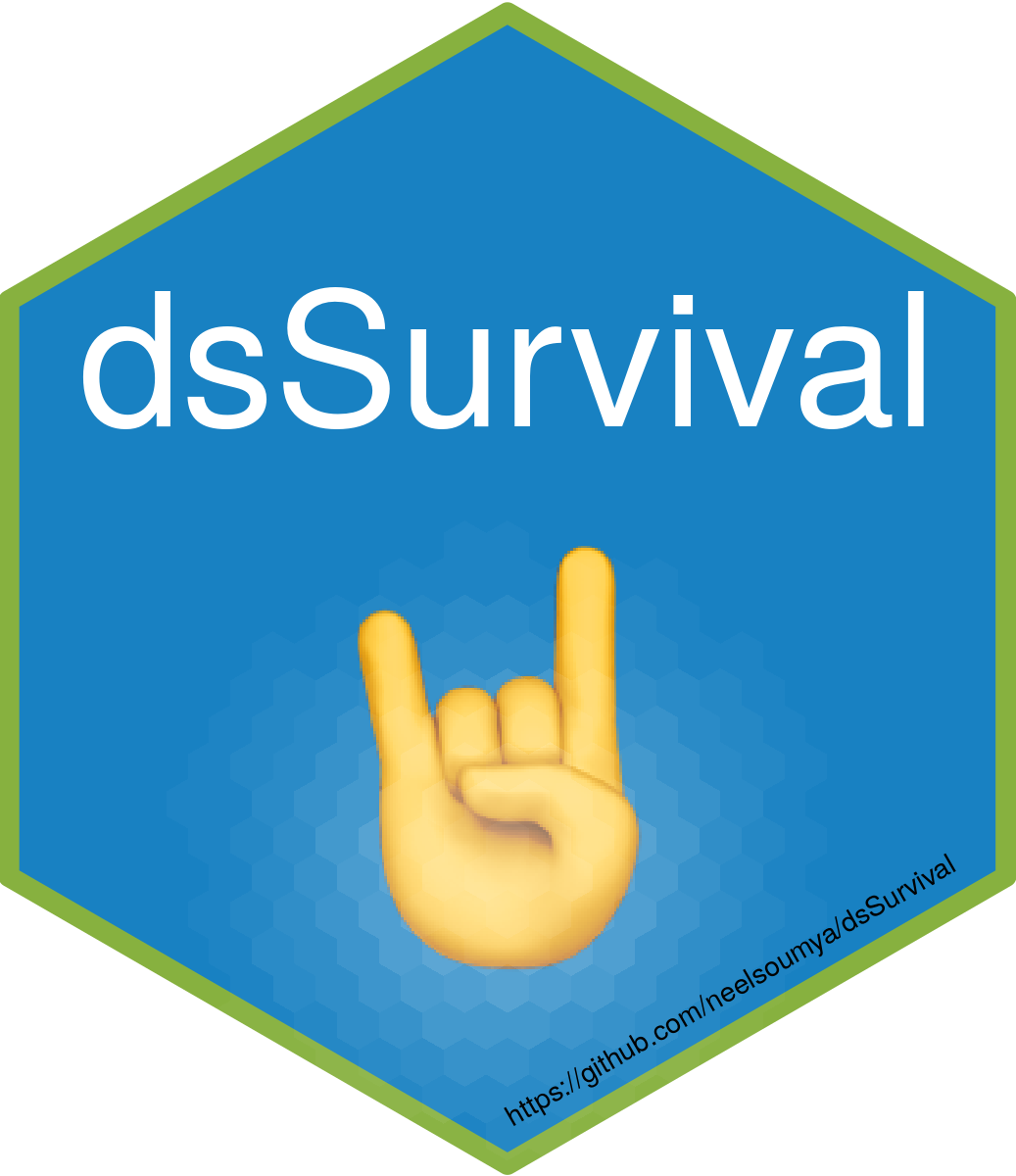

### nice-fig-1.png

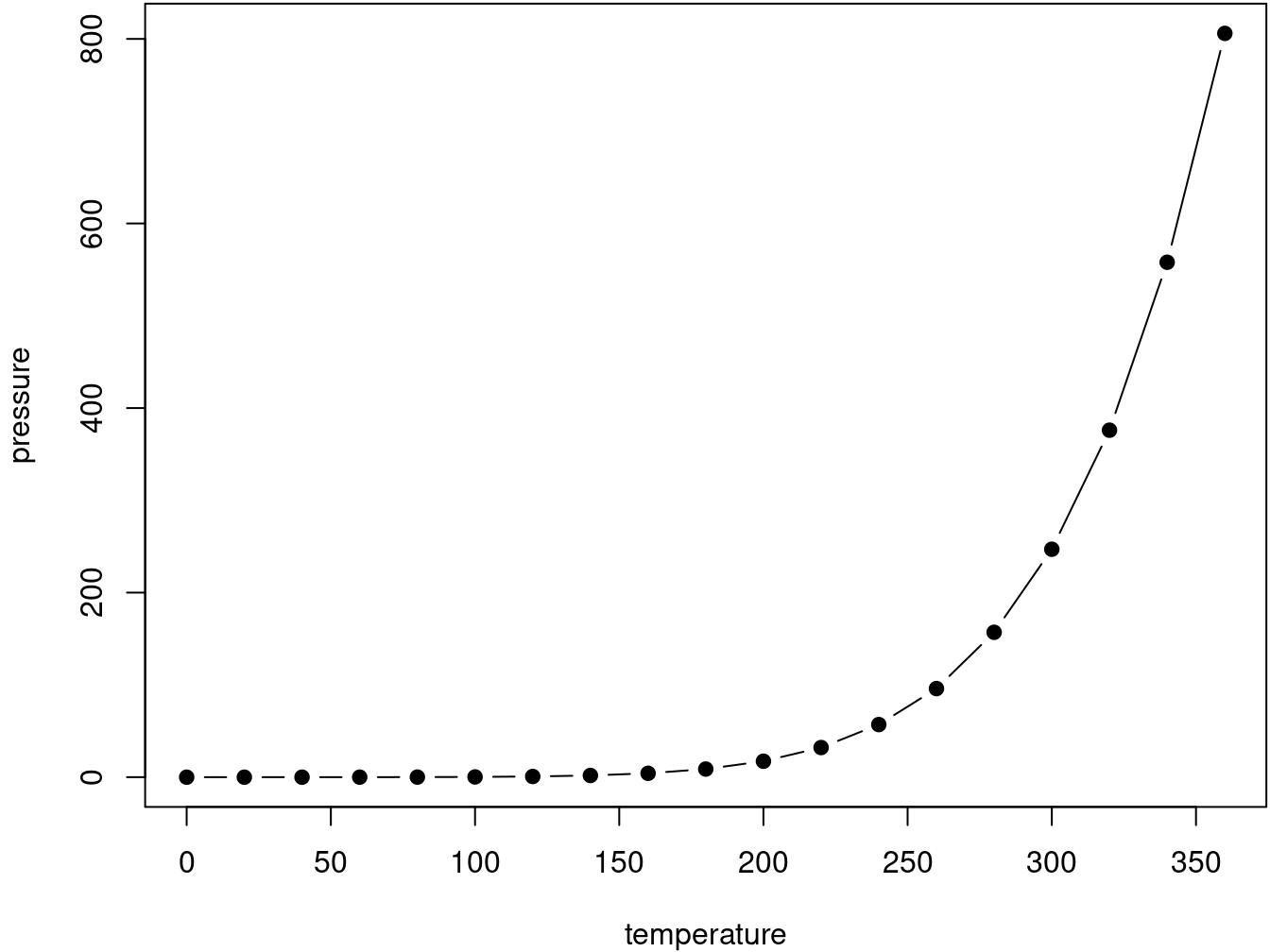

### screenshot_survival_models.png

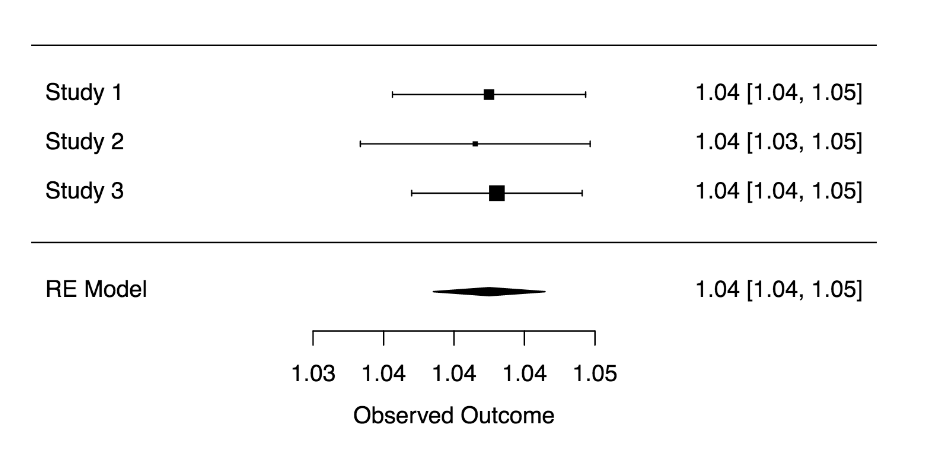

### unnamed-chunk-7-1.png

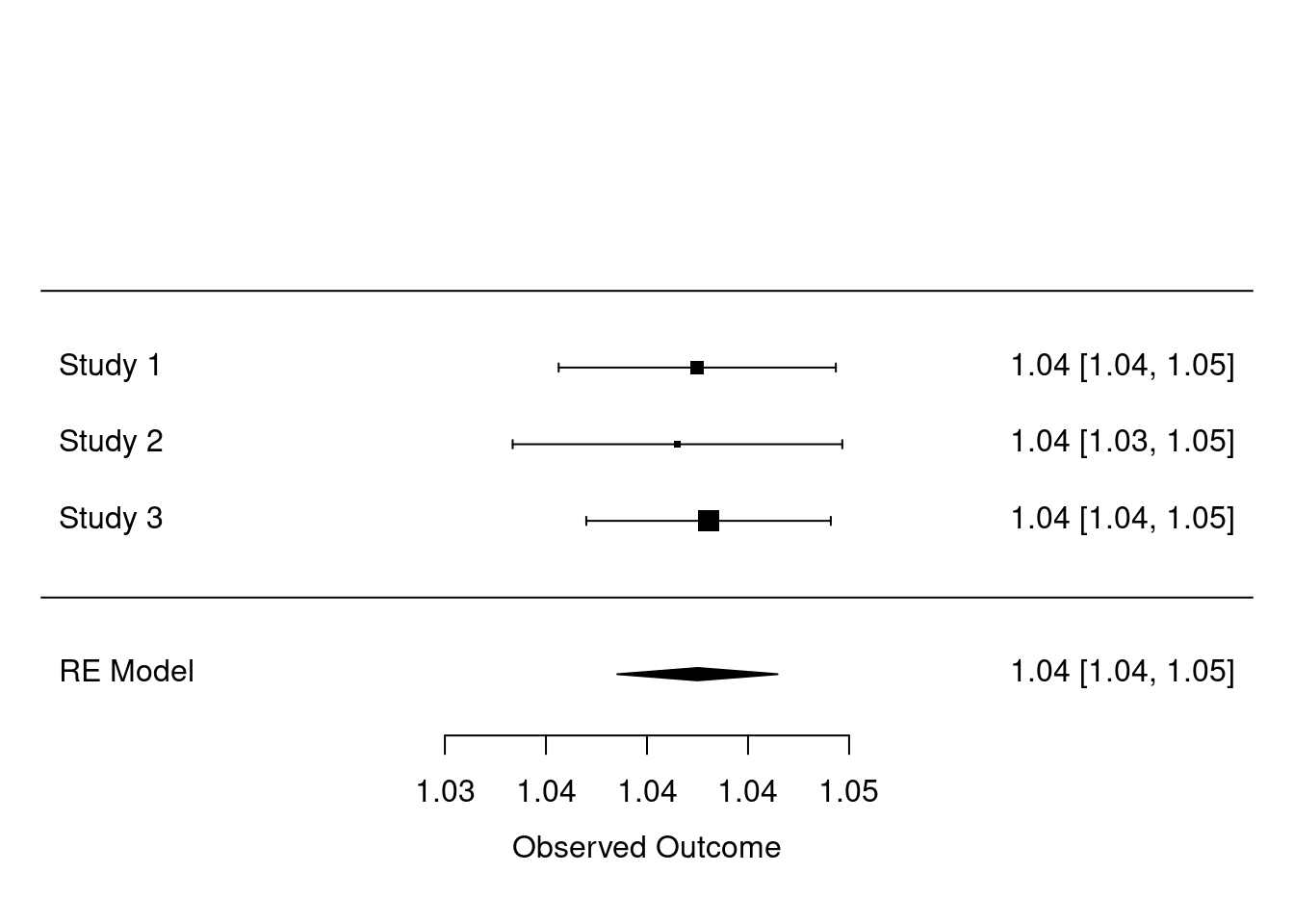

### unnamed-chunk-18-1.png

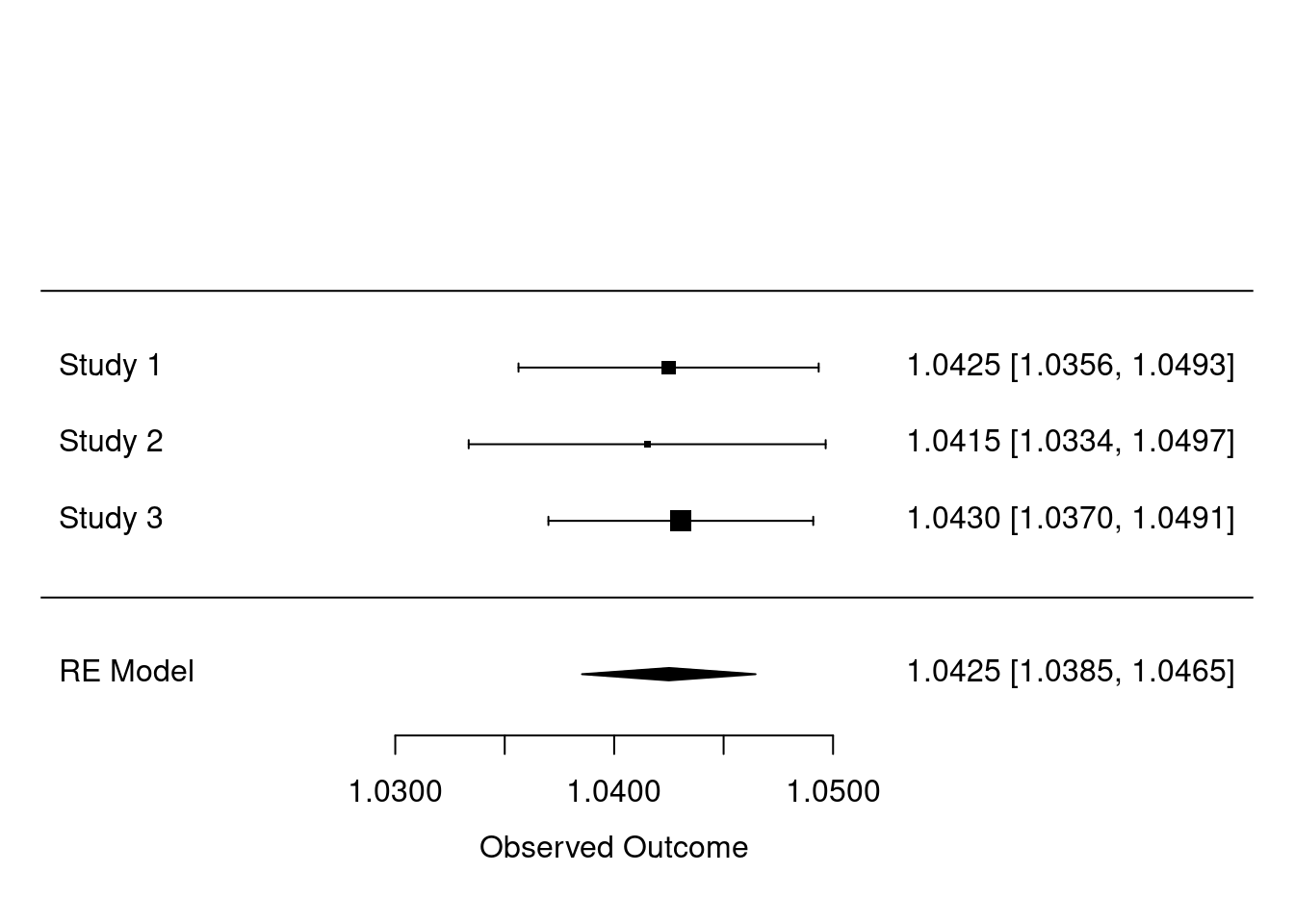

### unnamed-chunk-21-1.png

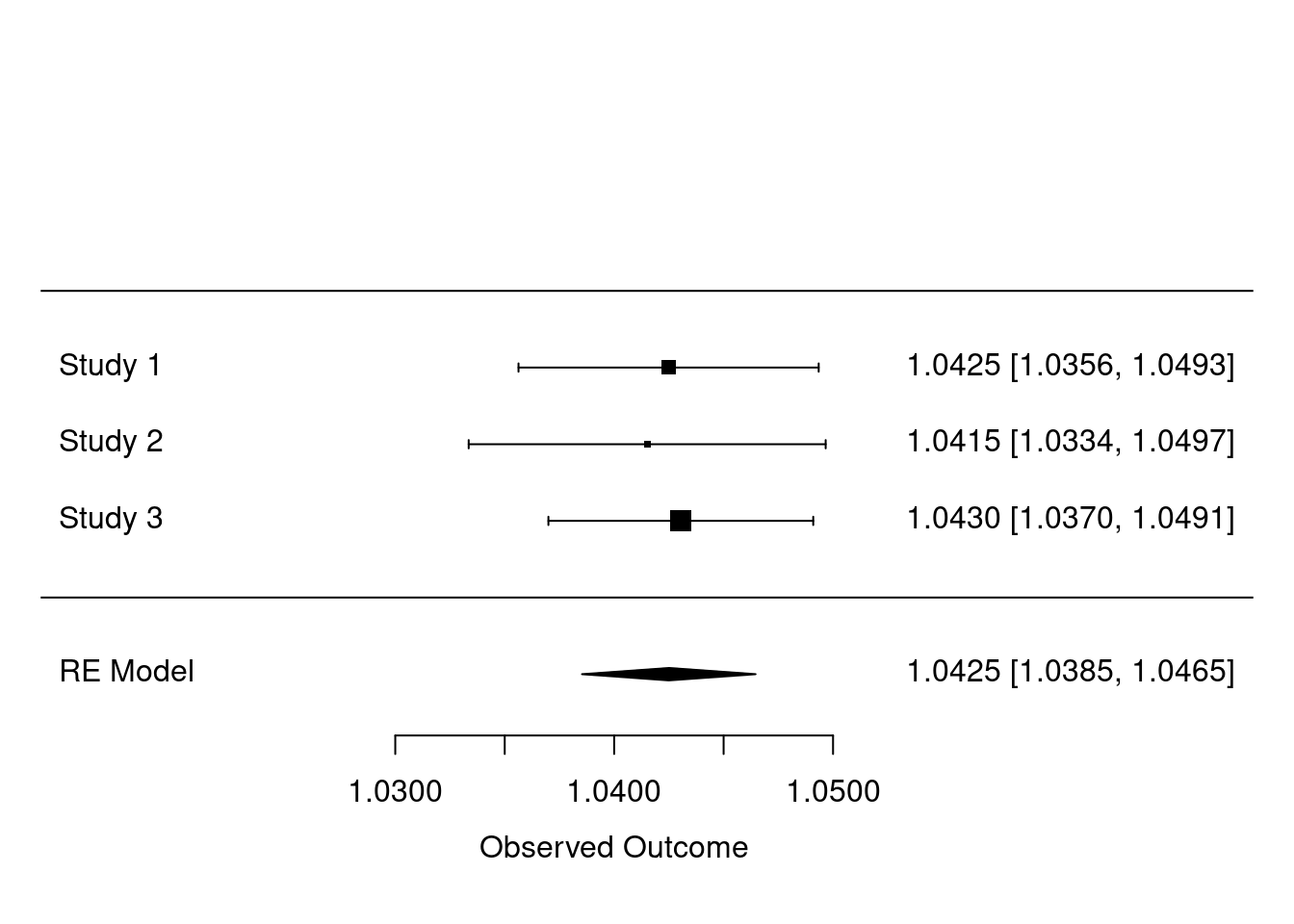
