## Supplementary material for "dsSurvival: Privacy preserving survival models for federated individual patient meta-analysis in DataSHIELD": Server side dsSurvival code: dsSurvival_1.0.0.pdf

### Package ‘dsSurvival’

July 2, 2021

**Title** DataSHIELD server side base functions for survival functions

**Description** DataSHIELD server side base functions for building survival models.

**Version** 1.0.0

**Author** Soumya Banerjee, Demetris Avraam, Xavier Escriba Montagut, Juan Gonzalez, Paul Burton and Tom R P Bishop <>

**Maintainer** Soumya Banerjee, Demetris Avraam, Xavier Escriba Montagut, Juan Gonzalez, Paul Burton and Tom R P Bishop <>

**License** GPL-3

**Depends** R (¿= 3.5.0)

**Imports** RANN,
stringr,
survival,
ggplot2,
dplyr,
reshape2,
dsBase

**RoxygenNote** 7.1.1

#### R topics documented:

`cox.zphSLMADS` *Tests the proportional hazards assumption of a Cox proportional hazards model that has been fit and saved serverside.*

#### Description

Tests the proportional hazards assumption of a Cox proportional hazards that has been fit and saved on the server side environment.

#### Usage

```
cox.zphSLMADS(
  fit = NULL,
  transform = "km",
  terms = TRUE,
  singledf = FALSE,
  global = TRUE
)
```

#### Arguments

| | |
|---|---|
| `fit` | character string specifying name of fit Cox proportional hazards model saved in the server-side. |
| `transform` | character string specifying how the survival times should be transformed before the test is performed. Possible values are "km", "rank", "identity" or a function of one argument. |
| `terms` | logical if TRUE, do a test for each term in the model rather than for each separate covariate. For a factor variable with k levels, for instance, this would lead to a k-1 degree of freedom test. The plot for such variables will be a single curve evaluating the linear predictor over time. |
| `singledf` | logical use a single degree of freedom test for terms that have multiple coefficients, i.e., the test that corresponds most closely to the plot. If terms=FALSE this argument has no effect. |
| `global` | logical should a global chi-square test be done, in addition to the per-variable or per-term tests tests. |

#### Details

Serverside aggregate function cox.zphSLMADS called by clientside function. ds.cox.zphSLMA. returns diagnostics for the test of proportional hazards assumptions from a Cox proportional hazards model. This request is not disclosive as it only returns summary statistics. For further details see help for ds.cox.zphSLMA function.

#### Value

diagnostics for the Cox proportional hazards from the server side environment.

##### Author(s)

Soumya Banerjee and Tom Bishop (2020).

---

`coxphSLMAassignDS` *Performs survival analysis using the Cox proportional hazards model at the serverside environment.*

---

##### Description

Performs survival analysis using the Cox proportional hazards models and stores the model on the server side environment.

##### Usage

```
coxphSLMAassignDS(
  formula = NULL,
  dataName = NULL,
  weights = NULL,
  init = NULL,
  ties = "efron",
  singular.ok = TRUE,
  model = FALSE,
  x = FALSE,
  y = TRUE,
  control = NULL
)
```

##### Arguments

| | |
|---|---|
| `formula` | either NULL or a character string (potentially including '*' wildcards) specifying a formula. |
| `dataName` | character string of name of data frame |
| `weights` | vector of case weights |
| `init` | vector of initial values of the iteration |
| `ties` | character string specifying the method for tie handling. The Efron approximation is used as the default. Other options are 'breslow' and 'exact'. |
| `singular.ok` | Logical value indicating how to handle collinearity in the model matrix. Default is TRUE. If TRUE, the program will automatically skip over columns of the X matrix that are linear combinations of earlier columns. In this case the coefficients of such columns will be NA and the variance matrix will contain zeros. |
| `model` | logical value. If TRUE, the model frame is returned in component model. |
| `x` | logical value. If TRUE, the x matrix is returned in component x. |
| `y` | logical value. If TRUE, the response vector is returned in component y. |
| `control` | object of type survival::coxph.control() specifying iteration limit and other control options. Default is survival::coxph.control() |

##### Details

Serverside assign function coxphSLMAassignDS called by clientside function. ds.coxphSLMAassign. stores the Cox proportional hazards in the server side environment This request is not disclosive as it only returns a string. For further details see help for ds.coxphSLMAassign function.

##### Value

the Cox proportional hazards from the server side environment from the server side environment.

##### Author(s)

Soumya Banerjee and Tom Bishop (2020).

---

#### coxphSLMADS — *Performs survival analysis using the Cox proportional hazards model at the serverside environment.*

---

##### Description

returns a summary of the Cox proportional hazards from the server side environment.

##### Usage

```
coxphSLMADS(
  formula = NULL,
  dataName = NULL,
  weights = NULL,
  init = NULL,
  ties = "efron",
  singular.ok = TRUE,
  model = FALSE,
  x = FALSE,
  y = TRUE,
  control = NULL
)
```

##### Arguments

| | |
|---|---|
| `formula` | either NULL or a character string (potentially including '*' wildcards) specifying a formula. |
| `dataName` | character string of name of data frame |
| `weights` | vector of case weights |
| `init` | vector of initial values of the iteration |
| `ties` | character string specifying the method for tie handling. The Efron approximation is used as the default. Other options are 'breslow' and 'exact'. |

| | |
|---|---|
| `singular.ok` | Logical value indicating how to handle collinearity in the model matrix. Default is TRUE. If TRUE, the program will automatically skip over columns of the X matrix that are linear combinations of earlier columns. In this case the coefficients of such columns will be NA and the variance matrix will contain zeros. |
| `model` | logical value. If TRUE, the model frame is returned in component model. |
| `x` | logical value. If TRUE, the x matrix is returned in component x. |
| `y` | logical value. If TRUE, the response vector is returned in component y. |
| `control` | object of type survival::coxph.control() specifying iteration limit and other control options. Default is survival::coxph.control() |

##### Details

Serverside aggregate function coxphSLMADS called by clientside function. ds.coxphSLMA. returns a summary of the Cox proportional hazards from the server side environment from the server side environment. This request is not disclosive as it only returns a string. For further details see help for ds.coxphSLMA function.

##### Value

a summary of the Cox proportional hazards from the server side environment from the server side environment.

##### Author(s)

Soumya Banerjee and Tom Bishop (2020).

---

#### `coxphSummaryDS` *Returns the summary of a Cox proportional hazards model that has been fit and saved serverside.*

---

##### Description

This function returns the summary of a Cox proportional hazards that has been fit and saved on the server side environment.

##### Usage

```
coxphSummaryDS(x = NULL)
```

##### Arguments

| | |
|---|---|
| `x` | character string specifying name of fit Cox proportional hazards model saved in the server-side. |

#### Details

Serverside aggregate function coxphSummaryDS called by clientside function. ds.coxphSummary. returns the summary from a Cox proportional hazards model. This request is not disclosive as it only returns summary statistics. For further details see help for ds.coxphSummary function.

#### Value

summary of the Cox proportional hazards from the server side environment.

#### Author(s)

Soumya Banerjee and Tom Bishop (2020).

---

### `listDisclosureSettingsDS` *listDisclosureSettingsDS*

---

#### Description

This serverside function is an aggregate function that is called by the ds.listDisclosureSettings

#### Usage

```
listDisclosureSettingsDS()
```

#### Details

For more details see the extensive header for ds.listDisclosureSettings

#### Author(s)

Paul Burton, Demetris Avraam for DataSHIELD Development Team

---

### `plotsurvfitDS` *Performs plotting of survival analysis curves.*

---

#### Description

returns a privacy preserving survival curve.

##### Usage

```
plotsurvfitDS(
  formula = NULL,
  dataName = NULL,
  method_anonymization = 2,
  noise = 0.03,
  knn = 20
)
```

##### Arguments

| | |
|---|---|
| `formula` | a character string which has the name of server-side survfit() object. This should be created using a call to ds.survfit() |
| `dataName` | character string of name of data frame |
| `method_anonymization` | an integer. Method of anonymization to be used (1: deterministic, 2: probabilistic). Default value is 2. |
| `noise` | an integer. fraction of noise (between 0 and 1) to be added to original data. Noise is added as a percentage of original value. This is used for probabilistic anonymization. Default value is 0.03 |
| `knn` | an integer. Number of nearest neighbours to be used for k nearest neighbours algoithm (for determinstic anonymization). Default value is 20. |

##### Details

Serverside aggregate function plotsurvfitDS called by clientside function. ds.plotsurvfit. returns a privacy preserving survival curve from the server side environment. This request is not disclosive as it is randomized. For further details see help for ds.plotsurvfit function.

##### Value

a privacy preserving survival curve from the server side environment.

##### Author(s)

Soumya Banerjee, Demetris Avraam, Paul Burton and Tom R P Bishop (2021).

---

#### `summarySurvDS` *Returns summary of survival object.*

---

##### Description

returns a summary of the survival Surv() object from the server side environment.

##### Usage

```
summarySurvDS(object = NULL)
```

##### Arguments

| | |
|---|---|
| `object` | name of server-side survival object. |

##### Details

Serverside aggregate function coxphSLMADS called by clientside function ds.summary. returns a list which is summary of the survival Surv() object. The list has the summary of the time and event parameter in the survival object. This request is not disclosive. For further details see help for ds.summary function.

##### Value

a list which is a summary of server-side survival model.

##### Author(s)

Soumya Banerjee and Tom Bishop (2021).

---

`SurvDS` *Creates a survival object for survival analysis using the Cox proportional hazards model at the serverside environment*

---

##### Description

returns a summary of the Cox proportional hazards from the server side environment.

##### Usage

```
SurvDS(time = NULL, time2 = NULL, event = NULL, type = NULL, origin = NULL)
```

##### Arguments

| | |
|---|---|
| `time` | name of start time or follow-up time parameter to be passed to Surv(). Should be a character string. |
| `time2` | name of stop time parameter to be passed to Surv(). Should be a character string. |
| `event` | name of event parameter to be passed to Surv() Should be character string. |
| `type` | character string specifying the type of censoring. Possible values are "right", "left", "counting", "interval", "interval2", or "mstate" |
| `origin` | numeric, used for counting process data and is the hazard function origin. The origin parameter is used with time-dependent strata in order to align the subjects properly when they cross over from one strata to another. This parameter has rarely proven useful. |

##### Details

Serverside assign function SurvDS called by clientside function. ds.Surv. returns a Survival object for use in Cox proportional hazards from the server side environment from the server side environment. This request is not disclosive as it only returns a string. For further details see help for ds.Surv function.

##### Value

a survival::Surv() object from the server side environment.

##### Author(s)

Soumya Banerjee and Tom Bishop (2021).

---

#### survfitDS — *Creates a survival survfit object for survival analysis at the serverside environment. This is to be used for eventually plotting survival models. A survival curve is based on a tabulation of the number at risk and number of events at each unique death time.*

---

##### Description

creates a survfit survival object in the server side environment.

##### Usage

```
survfitDS(formula = NULL)
```

##### Arguments

| | |
|---|---|
| `formula` | this is the formula to be passed to survfit(). Should be a character string. |

##### Details

Serverside assign function survfitDS called by clientside function. ds.survfit. creates a survfit survival object in the server side environment This request is not disclosive. For further details see help for ds.survfit function.

##### Value

creates a survfit survival object in the server side environment.

##### Author(s)

Soumya Banerjee and Tom Bishop (2020).
