## Supplementary material for "dsSurvival: Privacy preserving survival models for federated individual patient meta-analysis in DataSHIELD": Server side dsSurvival code: cox.zphSLMADS.R

```
#' 
#' @title Tests the proportional hazards assumption of a Cox proportional
#'	 hazards model that has been fit and saved serverside.
#' @description Tests the proportional hazards assumption of a 
#'	Cox proportional hazards that has been fit and saved on the
#'	server side environment.
#' @details Serverside aggregate function {cox.zphSLMADS} called by clientside function.
#' {ds.cox.zphSLMA}.
#' returns diagnostics for the test of proportional hazards assumptions
#'	from a Cox proportional hazards model.
#' This request is not disclosive as it only returns summary statistics.
#' For further details see help for {ds.cox.zphSLMA} function.
#' @param fit character string specifying name of fit Cox proportional 
#'	hazards model saved in the server-side.
#' @param transform character string specifying how the survival times should be transformed
#' 	before the test is performed. Possible values are "km", "rank", "identity" or a
#' 	function of one argument.
#' @param terms logical if TRUE, do a test for each term in the model rather than for each
#'	separate covariate. For a factor variable with k levels, for instance, this would lead
#'	to a k-1 degree of freedom test. The plot for such variables will be a single curve
#'	evaluating the linear predictor over time.
#' @param singledf logical use a single degree of freedom test for terms that have multiple 
#'	coefficients, i.e., the test that corresponds most closely to the plot. If terms=FALSE
#'	this argument has no effect.
#' @param global logical should a global chi-square test be done, in addition to the per-variable
#'	or per-term tests tests. 
#' @return diagnostics for the Cox proportional hazards from the server side environment.
#' @author Soumya Banerjee and Tom Bishop (2020).
#' @export
cox.zphSLMADS<-function(fit = NULL,
			transform = 'km',
			terms = TRUE,
			singledf = FALSE,
			global = TRUE
                       )
{
      
      errorMessage <- "No errors"
      
      #########################################################################
      # DataSHIELD MODULE: CAPTURE THE nfilter SETTINGS                       #
      thr <- listDisclosureSettingsDS()                                       #
      #nfilter.tab<-as.numeric(thr$nfilter.tab)                               #
      #nfilter.glm<-as.numeric(thr$nfilter.glm)                               #
      #nfilter.subset<-as.numeric(thr$nfilter.subset)                         #
      nfilter.string <- as.numeric(thr$nfilter.string)                        #
      nfilter.tab    <- as.numeric(thr$nfilter.tab)                           #
      nfilter.glm    <- as.numeric(thr$nfilter.glm)                           #
      #nfilter.stringShort<-as.numeric(thr$nfilter.stringShort)               #
      #nfilter.kNN<-as.numeric(thr$nfilter.kNN)                               #
      #datashield.privacyLevel<-as.numeric(thr$datashield.privacyLevel)       #
      #########################################################################
            
      # check if name of fit model is set
      if (is.null(fit))
      {
         stop("The name of the server-side fit Cox prportional hazards model must be set for use in survival::cox.zph()", call.=FALSE)
      } 	
	

      # evaluate the model fit parameter in parent environment
      fit_model <- eval(parse(text=fit), envir = parent.frame())

  
      ########################################
      # construct call to survival::cox.zph()
      ########################################
      coxzph_serverside <- survival::cox.zph(fit = fit_model,
					     transform = transform,
					     terms = terms,
					     singledf = singledf,
					     global = global
					    )      

      ###########################
      # disclosure checks
      ###########################
      # check if model oversaturated
      
      return(coxzph_serverside)
}
#AGGREGATE FUNCTION
# cox.zphSLMADS
```
