## Supplementary material for "dsSurvival: Privacy preserving survival models for federated individual patient meta-analysis in DataSHIELD": Server side dsSurvival code: coxphSLMAassignDS.R

```
#' 
#' @title Performs survival analysis using the Cox proportional hazards model at the serverside environment.
#' @description Performs survival analysis using the Cox proportional hazards models and stores the model on the server side environment.
#' @details Serverside assign function {coxphSLMAassignDS} called by clientside function.
#' {ds.coxphSLMAassign}.
#' stores the Cox proportional hazards in the server side environment
#' This request is not disclosive as it only returns a string.
#' For further details see help for {ds.coxphSLMAassign} function.
#' @param formula either NULL or a character string (potentially including '*'
#' wildcards) specifying a formula.
#' @param dataName character string of name of data frame
#' @param weights vector of case weights
#' @param init vector of initial values of the iteration
#' @param ties character string specifying the method for tie handling.
#'          The Efron approximation is used as the default. Other options are
#'          'breslow' and 'exact'.
#' @param singular.ok Logical value indicating how to handle collinearity in the model matrix.
#'        Default is TRUE. If TRUE, the program will automatically skip over columns of the 
#'        X matrix that are linear combinations of earlier columns. In this case the coefficients
#'        of such columns will be NA and the variance matrix will contain zeros.
#' @param model logical value. If TRUE, the model frame is returned in component model. 
#' @param x logical value. If TRUE, the x matrix is returned in component x.
#' @param y logical value. If TRUE, the response vector is returned in component y.
#' @param control object of type survival::coxph.control() specifying iteration limit and other
#'        control options. Default is survival::coxph.control()
#' @return the Cox proportional hazards from the server side environment from the server side environment.
#' @author Soumya Banerjee and Tom Bishop (2020).
#' @export
coxphSLMAassignDS<-function(formula = NULL,
                            dataName = NULL,
                            weights = NULL,
                            init = NULL,
                            ties = 'efron',
                            singular.ok = TRUE,
                            model = FALSE,
                            x = FALSE,
                            y = TRUE,
                            control = NULL
                           )
{
      
      errorMessage <- "No errors"
      
      #########################################################################
      # DataSHIELD MODULE: CAPTURE THE nfilter SETTINGS                       #
      thr <- listDisclosureSettingsDS()                                       #
      #nfilter.tab<-as.numeric(thr$nfilter.tab)                               #
      #nfilter.glm<-as.numeric(thr$nfilter.glm)                               #
      #nfilter.subset<-as.numeric(thr$nfilter.subset)                         #
      nfilter.string <- as.numeric(thr$nfilter.string)                        #
      nfilter.tab    <- as.numeric(thr$nfilter.tab)                           #
      nfilter.glm    <- as.numeric(thr$nfilter.glm)                           #
      #nfilter.stringShort<-as.numeric(thr$nfilter.stringShort)               #
      #nfilter.kNN<-as.numeric(thr$nfilter.kNN)                               #
      #datashield.privacyLevel<-as.numeric(thr$datashield.privacyLevel)       #
      #########################################################################
      
      # get the value of the 'data' and 'weights' parameters provided as character on the client side
      if(is.null(dataName))
      {
         dataTable <- NULL 
      }
      else
      {
         dataTable <- eval(parse(text=dataName), envir = parent.frame())
      }
      
      # check if formula is set
      if (is.null(formula))
      {
         stop("The formula must be set for use in survival::coxph()", call.=FALSE)
      } 

      ####################################################################	
      # Logic for parsing formula: since this need to be passed
      ####################################################################	
      # Put pipes back into formula
      #formula = as.formula(paste(formula,collapse="|"))
      formula <- Reduce(paste, deparse(formula))
      formula <- gsub("sssss", "survival::Surv(", formula, fixed = TRUE)
      formula <- gsub("lll", "=", formula, fixed = TRUE)
      formula <- gsub("xxx", "|", formula, fixed = TRUE)
      formula <- gsub("yyy", "(", formula, fixed = TRUE)
      formula <- gsub("zzz", ")", formula, fixed = TRUE)
      formula <- gsub("ppp", "/", formula, fixed = TRUE)
      formula <- gsub("qqq", ":", formula, fixed = TRUE)
      formula <- gsub("rrr", ",", formula, fixed = TRUE)

      # convert back to formula
      formula <- stats::as.formula(formula)
      
      
      #    formulatext <- Reduce(paste, deparse(formula))
      #    originalFormula <- formulatext
      #   
      # # Convert formula string into separate variable names split by |
      #   formulatext <- gsub(" ", "", formulatext, fixed=TRUE)
      #   formulatext <- gsub("~", "|", formulatext, fixed=TRUE)
      #   formulatext <- gsub("+", "|", formulatext, fixed=TRUE)
      #   formulatext <- gsub("*", "|", formulatext, fixed=TRUE)
      #   formulatext <- gsub("||", "|", formulatext, fixed=TRUE)
      # 
      #    formula2use <- stats::as.formula(paste0(Reduce(paste, deparse(originalFormula))), env = parent.frame()) # here we need the formula as a 'call' object
      
      # formula2use <- formula
      # mod.glm.ds <- stats::glm(formula2use, family=family, x=TRUE, control=stats::glm.control(maxit=1), contrasts=NULL, data=dataTable)
      #cxph_serverside <- survival::coxph(formula = survival::Surv(time = survtime, event = cens) ~  female,
      #                                   data = dataTable)
      
      #cxph_serverside <- survival::coxph(formula = survival::Surv(time = SURVTIME, event = EVENT) ~  1,
      #                                   data = dataTable)
      
      #cxph_serverside <- survival::coxph(formula = survival::Surv(time = SURVTIME, event = EVENT) ~  D$female,
      #                                   data = dataTable)
      
      ########################################
      # reconstruct control object
      ########################################
      if (is.null(control))
      {
            # if the value is null, then substitute default values which is 
            #   survival::coxph.control()
            control <- survival::coxph.control()
      }
      else
      {
            # reconstruct after passing this through parser
            ####################################################################	
            # Logic for parsing formula: since this need to be passed
            ####################################################################	
           
            # Put pipes back into formula
            #formula = as.formula(paste(formula,collapse="|"))
	    
            control <- Reduce(paste, deparse(control))
	    # remove the extra ~ bbbb passed here
	    #	this ~ bbbb needs to be passed because
	    #   everything needs to be passed as formula
	    #	and an expression of the form a ~ b is required	  
	    control <- gsub("~bbbb", "", control, fixed = TRUE)   
            control <- gsub("aaaaa", "survival::coxph.control(", control, fixed =  TRUE)
   	    control <- gsub("xxx", "|", control, fixed = TRUE)
   	    control <- gsub("yyy", "(", control, fixed = TRUE)
   	    control <- gsub("zzz", ")", control, fixed = TRUE)
	    control <- gsub("ppp", "/", control, fixed = TRUE)
	    control <- gsub("qqq", ":", control, fixed = TRUE)
	    control <- gsub("rrr", ",", control, fixed = TRUE)
	    #control <- gsub("", " ",    control, fixed = TRUE)
	    control <- gsub("lll", "=", control, fixed = TRUE)
            
            # use eval to construct an object of type survival::coxph.control()
            # control <- eval(parse(text=control), envir = parent.frame())
        
      }  
  
      ########################################
      # construct call to survival::coxph()
      ########################################
      # if init is NULL, then do not call coxph with init parameter
      if (!is.null(init))
      {
              cxph_serverside <- survival::coxph(formula = formula,
                                                 data = dataTable,
                                                 weights = weights,
                                                 init = init,
                                                 ties = ties,
                                                 singular.ok = singular.ok,
                                                 model = model,
                                                 x = x,
                                                 y = y#,
                                                 #control = eval(parse(text=as.character(control)))
                                                )
      }
      else
      {
              cxph_serverside <- survival::coxph(formula = formula,
                                                 data = dataTable,
                                                 weights = weights,
                                                 ties = ties,
                                                 singular.ok = singular.ok,
                                                 model = model,
                                                 x = x,
                                                 y = y#,
                                                 #control = eval(parse(text=as.character(control)))
                                                 )
      }
      
      ###########################
      # disclosure checks
      ###########################
      # check if model oversaturated
      num_parameters  <- length(cxph_serverside$coefficients)
      num_data_points <- cxph_serverside$n
      
      # if number of parameters greater than 0.2 * number of data points, then error
      if(num_parameters > (nfilter.glm * num_data_points) )
      {
            #glm.saturation.invalid<-1
            #errorMessage.gos<-paste0("ERROR: Model is oversaturated (too many model parameters relative to sample size)",
            #                 "leading to a possible risk of disclosure - please simplify model. With ",
            #                 num.p," parameters and nfilter.glm = ",round(nfilter.glm,4)," you need ",
            #                 round((num.p/nfilter.glm),0)," observations")
            return("ERROR: Model is oversaturated (too many model parameters relative to sample size)")
      }
      

      return(cxph_serverside)
}
#ASSIGN FUNCTION
# coxphSLMAassignDS
```
