## Supplementary material for "dsSurvival: Privacy preserving survival models for federated individual patient meta-analysis in DataSHIELD": Server side dsSurvival code: coxphSummaryDS.R

```
#' 
#' @title Returns the summary of a Cox proportional
#'	 hazards model that has been fit and saved serverside.
#' @description This function returns the summary of a 
#'	Cox proportional hazards that has been fit and saved on the
#'	server side environment.
#' @details Serverside aggregate function {coxphSummaryDS} called by clientside function.
#' {ds.coxphSummary}.
#' returns the summary from a Cox proportional hazards model.
#' This request is not disclosive as it only returns summary statistics.
#' For further details see help for {ds.coxphSummary} function.
#' @param x character string specifying name of fit Cox proportional 
#'	hazards model saved in the server-side.
#' @return summary of the Cox proportional hazards 
#'	from the server side environment.
#' @author Soumya Banerjee and Tom R. P. Bishop (2020).
#' @export
coxphSummaryDS<-function(x = NULL
                        )
{
      
      errorMessage <- "No errors"
      
      #########################################################################
      # DataSHIELD MODULE: CAPTURE THE nfilter SETTINGS                       #
      thr <- listDisclosureSettingsDS()                                       #
      #nfilter.tab<-as.numeric(thr$nfilter.tab)                               #
      #nfilter.glm<-as.numeric(thr$nfilter.glm)                               #
      #nfilter.subset<-as.numeric(thr$nfilter.subset)                         #
      nfilter.string <- as.numeric(thr$nfilter.string)                        #
      nfilter.tab    <- as.numeric(thr$nfilter.tab)                           #
      nfilter.glm    <- as.numeric(thr$nfilter.glm)                           #
      #nfilter.stringShort<-as.numeric(thr$nfilter.stringShort)               #
      #nfilter.kNN<-as.numeric(thr$nfilter.kNN)                               #
      #datashield.privacyLevel<-as.numeric(thr$datashield.privacyLevel)       #
      #########################################################################
            
      # check if name of fit model is set
      if (is.null(x))
      {
         stop("The name of the server-side fit Cox proportional hazards model must be set", call.=FALSE)
      } 	
	

      # evaluate the model fit parameter in parent environment
      fit_model <- eval(parse(text=x), envir = parent.frame())

  
      ###########################
      # disclosure checks
      ###########################
      # check if model oversaturated
      
      return(summary(fit_model))
}
#AGGREGATE FUNCTION
# coxphSummaryDS
```
