## Supplementary material for "dsSurvival: Privacy preserving survival models for federated individual patient meta-analysis in DataSHIELD": Server side dsSurvival code: listDisclosureSettingsDS.R

```
#'
#' @title listDisclosureSettingsDS
#' @description This serverside function is an aggregate function that is called by the
#' ds.listDisclosureSettings
#' @details For more details see the extensive header for ds.listDisclosureSettings
#' @author Paul Burton, Demetris Avraam for DataSHIELD Development Team
#' @export
#'
listDisclosureSettingsDS <- function(){

  nf.tab <- getOption("nfilter.tab")
  if (is.null(nf.tab))
    nf.tab <- getOption("default.nfilter.tab")
  nf.subset <- getOption("nfilter.subset")
  if (is.null(nf.subset))
    nf.subset <- getOption("default.nfilter.subset")
  nf.glm <- getOption("nfilter.glm")
  if (is.null(nf.glm))
    nf.glm <- getOption("default.nfilter.glm")
  nf.string <- getOption("nfilter.string")
  if (is.null(nf.string))
    nf.string <- getOption("default.nfilter.string")
  nf.stringShort <- getOption("nfilter.stringShort")
  if (is.null(nf.stringShort))
    nf.stringShort <- getOption("default.nfilter.stringShort")
  nf.kNN <- getOption("nfilter.kNN")
  if (is.null(nf.kNN))
    nf.kNN <- getOption("default.nfilter.kNN")
  nf.levels <- getOption("nfilter.levels")
  if (is.null(nf.levels))
    nf.levels <- getOption("default.nfilter.levels")
  nf.noise <- getOption("nfilter.noise")
  if (is.null(nf.noise))
    nf.noise <- getOption("default.nfilter.noise")
  nfilter.privacy.old <- getOption("datashield.privacyLevel")

  return(list(nfilter.tab=nf.tab,nfilter.subset=nf.subset,nfilter.glm=nf.glm,nfilter.string=nf.string,
              nfilter.stringShort=nf.stringShort,nfilter.kNN=nf.kNN,nfilter.levels=nf.levels,
              nfilter.noise=nf.noise,nfilter.privacy.old=nfilter.privacy.old))
}
#AGGREGATE FUNCTION
# listDisclosureSettingsDS
```
