## Supplementary material for "dsSurvival: Privacy preserving survival models for federated individual patient meta-analysis in DataSHIELD": Server side dsSurvival code: plotsurvfitDS.R

```
#' 
#' @title Performs plotting of survival analysis curves.
#' @description returns a privacy preserving survival curve.
#' @details Serverside aggregate function {plotsurvfitDS} called by clientside function.
#' {ds.plotsurvfit}.
#' returns a privacy preserving survival curve from the server side environment.
#' This request is not disclosive as it is randomized.
#' For further details see help for {ds.plotsurvfit} function.
#' @param formula a character string which has the name of server-side survfit() object.
#'		This should be created using a call to ds.survfit()
#' @param dataName character string of name of data frame
#' @param method_anonymization an integer. Method of anonymization to be used (1: deterministic, 2: probabilistic). Default value is 2.
#' @param noise an integer. fraction of noise (between 0 and 1) to be added to original data. 
#'     Noise is added as a percentage of original value. 
#'     This is used for probabilistic anonymization.
#'     Default value is 0.03 
#' @param knn an integer. Number of nearest neighbours to be used for k nearest neighbours algoithm (for determinstic anonymization). 
#'     Default value is 20.
#' @return a privacy preserving survival curve from the server side environment.
#' @author Soumya Banerjee, Demetris Avraam, Paul Burton and Tom RP Bishop (2021).
#' @export
plotsurvfitDS<-function(formula = NULL,
                        dataName = NULL,
			method_anonymization = 2,
			noise = 0.03,
			knn = 20
                       )
{
  
  errorMessage <- "No errors"
  
  #########################################################################
  # DataSHIELD MODULE: CAPTURE THE nfilter SETTINGS                       #
  thr <- listDisclosureSettingsDS()                                       #
  #nfilter.tab<-as.numeric(thr$nfilter.tab)                               #
  #nfilter.glm<-as.numeric(thr$nfilter.glm)                               #
  #nfilter.subset<-as.numeric(thr$nfilter.subset)                         #
  nfilter.string <- as.numeric(thr$nfilter.string)                        #
  nfilter.tab    <- as.numeric(thr$nfilter.tab)                           #
  nfilter.glm    <- as.numeric(thr$nfilter.glm)                           #
  nfilter.noise  <- as.numeric(thr$nfilter.noise)                         #
  #nfilter.stringShort<-as.numeric(thr$nfilter.stringShort)               #
  #nfilter.kNN<-as.numeric(thr$nfilter.kNN)                               #
  #datashield.privacyLevel<-as.numeric(thr$datashield.privacyLevel)       #
  #########################################################################
  
  # get the value of the 'data' and 'weights' parameters provided as character on the client side
  if(is.null(dataName))
  {
    dataTable <- NULL 
  }
  else
  {
    dataTable <- eval(parse(text=dataName), envir = parent.frame())
  }
  
  # check if formula is set
  if (is.null(formula))
  {
    stop("The formula must be set for use in survival::coxph()", call.=FALSE)
  }
  

  # get survfit model
  survfit_model_variable = eval(parse(text=formula), envir = parent.frame())
  
  # method_anonymization = 2
  # noise = 0.03 # 0.0003 # 0.03 0.26
  # knn <- 20	
  
  ##############################################################
  # if probabilistic anonymization then generate and add noise	
  ##############################################################
  if (method_anonymization == 2)
  {	      
    
    # set study specific seed
    seed <- getOption("datashield.seed")
    if (is.null(seed))
    {
      stop("plotsurvfitDS requires a seed to be set and requires 'datashield.seed' R option to operate", call.=FALSE)
    }
    
    # if there is a seed, then set it
    set.seed(seed)	
    
    # TODO: check if percentage of noise greater than threshold
    # if(noise < nfilter.noise)
    # {
    #	  stop(paste0("'noise' must be greater than or equal to ", nfilter.noise), call.=FALSE)
    # }
    # else
    # {
    #	  percentage <- noise
    # }
    
    percentage <- noise
    
    ##########################################
    # Approach 1: add noise before plotting
    #
    # add noise to:
    # surv (i.e. proportion surviving)
    # time (times at which events occur, ie when the proportion changes)
    # this is for the y axis
    # and for time on x axis
    ##########################################
    for ( i_temp_counter_inner in c(2:length(survfit_model_variable$surv)) )
    {
	      # current value, upper, lower at this index
	      # value_temp <- survfit_model_variable$surv[i_temp_counter_inner]
	      # upper_temp <- survfit_model_variable$upper[i_temp_counter_inner]
	      # lower_temp <- survfit_model_variable$lower[i_temp_counter_inner]

	      # previous value, upper, lower
	      # prev_value_temp <- survfit_model_variable$surv[i_temp_counter_inner - 1]
	      # prev_upper_temp <- survfit_model_variable$upper[i_temp_counter_inner - 1]
	      # prev_lower_temp <- survfit_model_variable$lower[i_temp_counter_inner - 1]

	      # add some noise 
	      # can make noise a percentage of previous OR current value
	      # # delta_noise <- abs(stats::rnorm(n = 1, mean = value_temp, sd = percentage * value_temp))
	      # delta_noise <- stats::rnorm(n = 1, mean = 0, sd = percentage)

	      # SUBTRACT this noise from the PREVIOUS VALUE if it does not cause problems with monotonicity

	      # value_noise = value_temp - delta_noise
	      # upper_noise = upper_temp - delta_noise
	      # lower_noise = lower_temp - delta_noise

	      #if (prev_value_temp >= value_noise)
	      #{
	      #  survfit_model_variable$surv[i_temp_counter_inner] <- value_noise
	      #  survfit_model_variable$upper[i_temp_counter_inner] <- upper_noise
	      #  survfit_model_variable$lower[i_temp_counter_inner] <- lower_noise
	      #}
	      #else
	      #{
	      #  survfit_model_variable$surv[i_temp_counter_inner] = prev_value_temp
	      #  survfit_model_variable$upper[i_temp_counter_inner] = prev_upper_temp
	      #  survfit_model_variable$lower[i_temp_counter_inner] = prev_lower_temp
	      #}

	      # survfit_model_variable$mono[i_temp_counter_inner] = prev_value_temp - survfit_model_variable$surv[i_temp_counter_inner]

	      # new noise for x axis
	      # delta_noise <- stats::rnorm(n = 1, mean = 0, sd = percentage)
	      # survfit_model_variable$time[i_temp_counter_inner] <- survfit_model_variable$time[i_temp_counter_inner] - delta_noise


	      # generate noise absolute value so alsways positive
	      # delta_noise_time <- abs( stats::rnorm(n = 1, mean = 0, sd = percentage) )
	      # do not take absolute value but have it reject values if not monotonically increasing
	      delta_noise_time <- stats::rnorm(n = 1, mean = 0, sd = percentage)	    
	      # survfit_model_variable$time[i_temp_counter_inner] <- survfit_model_variable$time[i_temp_counter_inner] + delta_noise

	      # previous value time
	      prev_value_temp_time <- survfit_model_variable$time[i_temp_counter_inner - 1]
	      # current value temp time
	      curr_value_temp_time <- survfit_model_variable$time[i_temp_counter_inner]

	      # proposed value of time with noise added as a proportion and added on top of whatever value exists
	      value_noise_time_proposed = curr_value_temp_time + (curr_value_temp_time*delta_noise_time)

	      if (prev_value_temp_time <= value_noise_time_proposed)
	      {
		  # if previous value time is less then proposed value then ok 
		  # since time is supposed to be increasing

		  # set this value to current time
		  survfit_model_variable$time[i_temp_counter_inner] = value_noise_time_proposed

	      }
	      else
	      {
		   # set to previous time value
		   survfit_model_variable$time[i_temp_counter_inner] = prev_value_temp_time
	      }	    
	    
	    
	    
    }
    # end for loop	  
    
  }
  
  ######################################################################
  # Approach 2: 
  # deterministic anonymization by Demetris Avraam and Paul Burton
  ######################################################################
  if (method_anonymization == 1)
  {
        
        # TODO: make this a parameter and check if knn is not less than a threshold (see plothistogramDS) uncomment lines in beginning og function and below	  
	# if (knn < nfilter.kNN)
        # {
        #      stop(paste0("'knn' must be greater than or equal to ", nfilter.kNN), call.=FALSE)  # nfilter.noise
        # }	  
	  
        # Step 1: Standardise the variable
        time.standardised <- (survfit_model_variable$time-mean(survfit_model_variable$time))/stats::sd(survfit_model_variable$time)

        # Step 2: Find the k-1 nearest neighbours of each data point
        nearest <- RANN::nn2(time.standardised, k = knn)

        # Step 3: Calculate the centroid of each n nearest data points
        time.centroid <- matrix()
        for (i in 1:length(survfit_model_variable$time))
	{
             time.centroid[i] <- mean(time.standardised[nearest$nn.idx[i,1:knn]])
        }

        # Step 4: Calculate the scaling factor
        time.scalingFactor <- stats::sd(time.standardised)/stats::sd(time.centroid)

        # Step 5: Apply the scaling factor to the centroids
        time.masked <- time.centroid * time.scalingFactor

        # Step 6: Shift the centroids back to the actual position and scale of the original data
        SURVTIME_anon <- (time.masked * stats::sd(survfit_model_variable$time)) + mean(survfit_model_variable$time)

        # modify time in survfit object (instead of original time)
        survfit_model_variable$time <- SURVTIME_anon
	  
	  
  }  
    

  # return modified survfit object
  return(survfit_model_variable)

}
#AGGREGATE FUNCTION
# plotsurvfitDS
```
