## Supplementary material for "dsSurvival: Privacy preserving survival models for federated individual patient meta-analysis in DataSHIELD": Server side dsSurvival code: summarySurvDS.R

```
#' 
#' @title Returns summary of survival object.
#' @description returns a summary of the survival Surv() object from the server side environment.
#' @details Serverside aggregate function {coxphSLMADS} called by clientside function 
#' {ds.summary}.
#' returns a list which is summary of the survival Surv() object. The list has the summary of the time
#'   and event parameter in the survival object.
#' This request is not disclosive.
#' For further details see help for {ds.summary} function.
#' @param object name of server-side survival object.
#' @return a list which is a summary of server-side survival model.
#' @author Soumya Banerjee and Tom Bishop (2021).
#' @export
summarySurvDS<-function(object=NULL)
{
      
      errorMessage <- "No errors"
      
      #########################################################################
      # DataSHIELD MODULE: CAPTURE THE nfilter SETTINGS                       #
      thr <- listDisclosureSettingsDS()                                       #
      #nfilter.tab<-as.numeric(thr$nfilter.tab)                               #
      #nfilter.glm<-as.numeric(thr$nfilter.glm)                               #
      #nfilter.subset<-as.numeric(thr$nfilter.subset)                         #
      nfilter.string <- as.numeric(thr$nfilter.string)                        #
      nfilter.tab    <- as.numeric(thr$nfilter.tab)                           #
      nfilter.glm    <- as.numeric(thr$nfilter.glm)                           #
      #nfilter.stringShort<-as.numeric(thr$nfilter.stringShort)               #
      #nfilter.kNN<-as.numeric(thr$nfilter.kNN)                               #
      #datashield.privacyLevel<-as.numeric(thr$datashield.privacyLevel)       #
      #########################################################################
      
      # get the value of the 'data' and 'weights' parameters provided as character on the client side
      #if(is.null(dataName)){
      #   dataTable <- NULL 
      #}else{
      #   dataTable <- eval(parse(text=dataName), envir = parent.frame())
      #}
      
      if (is.null(object))
      {
            stop("The input object must be a survival::Surv object", call.=FALSE)
      }
      
      ###########################
      # disclosure checks
      ###########################
      # check if model oversaturated
      #num_parameters  <- length(cxph_serverside$coefficients)
      #num_data_points <- cxph_serverside$n
      
      #if(num_parameters > (nfilter.glm * num_data_points) )
      #{
      #      #glm.saturation.invalid<-1
      #      #errorMessage.gos<-paste0("ERROR: Model is oversaturated (too many model parameters relative to sample size)",
      #      #                 "leading to a possible risk of disclosure - please simplify model. With ",
      #      #                 num.p," parameters and nfilter.glm = ",round(nfilter.glm,4)," you need ",
      #      #                 round((num.p/nfilter.glm),0)," observations")
      #      return("ERROR: Model is oversaturated (too many model parameters relative to sample size)")
      #}

      
      surv_object <- object
      summary_surv_object_time   <- dsBase::quantileMeanDS(xvect = surv_object[,1])
      summary_surv_object_status <- dsBase::quantileMeanDS(xvect = surv_object[,2])
      
      return_object <- list("time"=summary_surv_object_time, "event"=summary_surv_object_status)
      
      return(return_object)
      
      
}
#AGGREGATE FUNCTION
# summarySurvDS
```
