## Supplementary material for "dsSurvival: Privacy preserving survival models for federated individual patient meta-analysis in DataSHIELD": Server side dsSurvival code: SurvDS.R

```
#' 
#' @title Creates a survival object for survival analysis using the Cox proportional hazards model at the serverside environment
#' @description returns a summary of the Cox proportional hazards from the server side environment.
#' @details Serverside assign function {SurvDS} called by clientside function.
#' {ds.Surv}.
#' returns a Survival object for use in Cox proportional hazards from the server side environment from the server side environment.
#' This request is not disclosive as it only returns a string.
#' For further details see help for {ds.Surv} function.
#' @param time name of start time or follow-up time parameter to be passed to Surv(). 
#'      Should be a character string.
#' @param time2 name of stop time parameter to be passed to Surv(). 
#'      Should be a character string.
#' @param event name of event parameter to be passed to Surv()
#'      Should be character string.
#' @param type character string specifying the type of censoring. Possible values are "right", "left",
#'	"counting", "interval", "interval2", or "mstate"
#' @param origin numeric, used for counting process data and is the hazard function origin.
#'	The origin parameter is used with time-dependent strata in order to align the subjects
#'	properly when they cross over from one strata to another. This parameter has rarely
#'	proven useful.
#' @return a survival::Surv() object from the server side environment.
#' @author Soumya Banerjee and Tom Bishop (2021).
#' @export
SurvDS<-function(time = NULL,
                 time2 = NULL,
                 event = NULL,
                 type = NULL, #c('right', 'left', 'interval', 'counting', 'interval2', 'mstate'),
		 origin = NULL)
{
      #########################################################################
      # DataSHIELD MODULE: CAPTURE THE nfilter SETTINGS                       #
      thr <- listDisclosureSettingsDS()                                       #
      #nfilter.tab<-as.numeric(thr$nfilter.tab)                               #
      #nfilter.glm<-as.numeric(thr$nfilter.glm)                               #
      #nfilter.subset<-as.numeric(thr$nfilter.subset)                         #
      nfilter.string <- as.numeric(thr$nfilter.string)                        #
      nfilter.tab    <- as.numeric(thr$nfilter.tab)                           #
      nfilter.glm    <- as.numeric(thr$nfilter.glm)                           #
      #nfilter.stringShort<-as.numeric(thr$nfilter.stringShort)               #
      #nfilter.kNN<-as.numeric(thr$nfilter.kNN)                               #
      #datashield.privacyLevel<-as.numeric(thr$datashield.privacyLevel)       #
      #########################################################################
      
    
      #################################
      # check type of all parameters
      #################################
      # check type for time parameter
      class_time <- dsBase::classDS(x=time)
      if ( !('numeric' %in% class_time) & !('integer' %in% class_time) )
      {
            stop('Start time parameter or follow-up time parameter (time) must be numeric or integer.', call.=FALSE)
      }
      
      # check type for time2 parameter
      class_time2 <- dsBase::classDS(x=time2)
      # if time2 is not NULL, only then check if it is numeric or integer
      #     this is because time2 is an optional parameter
      #     for use in interval censored data
      if (!is.null(time2))
      {      
            if( !('numeric' %in% class_time2) & !('integer' %in% class_time2) )
            {
                  stop('Stop time parameter (time2) must be numeric or integer.', call.=FALSE)
            }
      }
      
      # check type for event parameter
      class_event <- dsBase::classDS(x=event)
      if ( !('numeric' %in% class_event) & !('integer' %in% class_event) & !('factor' %in% class_event) )
      {
            stop('Event parameter (event) must be numeric or integer or factor.', call.=FALSE)
      }
      
      #########################################################################	
      # evaluate all these parameters in the parent frame in the call stack	
      #########################################################################	
      if (!is.null(time))
      {
	      time_param = eval(parse(text=time), envir = parent.frame())
      }
      else
      {
	      time_param = NULL
      } 	      
      
      if (!is.null(time2))
      {
	      time2_param = eval(parse(text=time2), envir = parent.frame())
      }
      else
      {
	      time2_param = NULL
      }	      
	
      if (!is.null(event))
      {
	      event_param = eval(parse(text=event), envir = parent.frame())
      }
      else
      {
	      event_param = NULL
      }	      
	      
	
      # evaluate this
      # surv_object <- eval(parse(text=str_command), envir = parent.frame())
 
      # evaluate this expression	
      if (!is.null(time2_param))
      {	  
	   # if type is NULL, then do not pass it. Surv() has internal logic to derive type
	   # 	from the values of the other parameters   
	   if (!is.null(type))
	   {	   
           	surv_object <- survival::Surv(time = time_param, event = event_param, time2 = time2_param, type = type, origin = origin)
	   }
	   else
	   {
	        surv_object <- survival::Surv(time = time_param, event = event_param, time2 = time2_param, origin = origin)	   
	   }
      }
      else
      {
	   # if type is NULL, then do not pass it. Surv() has internal logic to derive type
	   #	from the values of the other parameters  
	   if (!is.null(type))
           {
	   	surv_object <- survival::Surv(time = time_param, event = event_param, type = type, origin = origin)
	   }
	   else
	   {
		surv_object <- survival::Surv(time = time_param, event = event_param, origin = origin)   
	   }	   
      }	      
	
      
      # surv_object <- eval(parse(text='survival::Surv(time = SURVTIME, event = EVENT)'), envir = parent.frame())
      
      return(surv_object)
}
#ASSIGN FUNCTION
# SurvDS
```
