## Supplementary material for "dsSurvival: Privacy preserving survival models for federated individual patient meta-analysis in DataSHIELD": Server side dsSurvival code: survfitDS.R

```
#' 
#' @title Creates a survival survfit object for survival analysis at the serverside environment. 
#'   This is to be used for eventually plotting survival models.
#'   A survival curve is based on a tabulation of the number at risk
#'   and number of events at each unique death time.
#' @description creates a survfit survival object in the server side environment.
#' @details Serverside assign function {survfitDS} called by clientside function.
#' {ds.survfit}.
#' creates a survfit survival object in the server side environment
#' This request is not disclosive.
#' For further details see help for {ds.survfit} function.
#' @param formula this is the formula to be passed to survfit(). 
#'      Should be a character string.
#' @return creates a survfit survival object in the server side environment.
#' @author Soumya Banerjee and Tom Bishop (2020).
#' @export
survfitDS<-function(formula=NULL)
{
      #########################################################################
      # DataSHIELD MODULE: CAPTURE THE nfilter SETTINGS                       #
      thr <- listDisclosureSettingsDS()                                       #
      #nfilter.tab<-as.numeric(thr$nfilter.tab)                               #
      #nfilter.glm<-as.numeric(thr$nfilter.glm)                               #
      #nfilter.subset<-as.numeric(thr$nfilter.subset)                         #
      nfilter.string <- as.numeric(thr$nfilter.string)                        #
      nfilter.tab    <- as.numeric(thr$nfilter.tab)                           #
      nfilter.glm    <- as.numeric(thr$nfilter.glm)                           #
      #nfilter.stringShort<-as.numeric(thr$nfilter.stringShort)               #
      #nfilter.kNN<-as.numeric(thr$nfilter.kNN)                               #
      #datashield.privacyLevel<-as.numeric(thr$datashield.privacyLevel)       #
      #########################################################################
      
    
      # construct a call to Surv function with these parameters
      
      #time  = "SURVTIME"
      #event = "EVENT"
      ## surv_object <- survival::Surv(time = SURVTIME, event = EVENT)
      #str_command = paste0('survival::Surv(time = ', time)
      #str_command = paste0(str_command, ', event = ') 
      #str_command = paste0(str_command, event)
      #str_command = paste0(str_command, ')')
      
      # evaluate this
      #surv_object <- eval(parse(text=str_command), envir = parent.frame())
      
      # TODO: convert special characters in formula here
      # TODO: use formula here and remove later
      # formula = "surv_object ~ 1"
      
      # check if formula is provided
      if (is.null(formula))
      {
            stop("The input must have a non-empty formula to be used in survival::survfit()", call.=FALSE)
      }

      #####################################################################	
      # Logic for reconstructing formula: since this need to be passed
      #     to parser, we need to remove special symbols
      #     On the server-side function (survfitDS) this needs
      #     to be reconstructed
      #     formula as text, then split at pipes to avoid triggering parser
      #####################################################################
      # formula = as.formula(paste(formula,collapse="|"))
      formula <- Reduce(paste, deparse(formula))
      formula <- gsub("sssss", "survival::Surv(", formula, fixed = TRUE)
      formula <- gsub("lll", "=", formula, fixed = TRUE)
      formula <- gsub("xxx", "|", formula, fixed = TRUE)
      formula <- gsub("yyy", "(", formula, fixed = TRUE)
      formula <- gsub("zzz", ")", formula, fixed = TRUE)
      formula <- gsub("ppp", "/", formula, fixed = TRUE)
      formula <- gsub("qqq", ":", formula, fixed = TRUE)
      formula <- gsub("rrr", ",", formula, fixed = TRUE)

      # convert to type formula
      formula <- stats::as.formula(formula)
            
      # call survival:: survfit
      # TODO: inject random noise
      survfit_object <- survival::survfit(formula)
            
      return(survfit_object)
}
#ASSIGN FUNCTION
# survfitDS
```
