## Supplementary material for "dsSurvival: Privacy preserving survival models for federated individual patient meta-analysis in DataSHIELD": Server side dsSurvival code: set_disclosure_settings.R

```
#-------------------------------------------------------------------------------
 2 # Copyright (c) 2019-2020 University of Newcastle upon Tyne. All rights reserved.
 3 #
 4 # This program and the accompanying materials
 5 # are made available under the terms of the GNU Public License v3.0.
 6 #
 7 # You should have received a copy of the GNU General Public License
 8 # along with this program.  If not, see <http://www.gnu.org/licenses/>.
 9 #-------------------------------------------------------------------------------
10 #
11 # DataSHIELD disclosure settings
12 #

set.standard.disclosure.settings <- function() {
    options(datashield.privacyLevel = "5")
    options(default.nfilter.glm = "0.33")
    options(default.nfilter.kNN = "3")
    options(default.nfilter.string = "80")
    options(default.nfilter.subset = "3")
    options(default.nfilter.stringShort = "20")
    options(default.nfilter.tab = "3")
    options(default.nfilter.noise = "0.25")
    options(default.nfilter.levels = "0.33")
}
```
