## Supplementary material for "dsSurvival: Privacy preserving survival models for federated individual patient meta-analysis in DataSHIELD": Server side dsSurvival code: set_random_seed_settings.R

```
#-------------------------------------------------------------------------------
 2 # Copyright (c) 2019-2020 University of Newcastle upon Tyne. All rights reserved.
 3 #
 4 # This program and the accompanying materials
 5 # are made available under the terms of the GNU Public License v3.0.
 6 #
 7 # You should have received a copy of the GNU General Public License
 8 # along with this program.  If not, see <http://www.gnu.org/licenses/>.
 9 #-------------------------------------------------------------------------------
10 #
11 # DataSHIELD random settings
12 #

set.random.seed.setting <- function(seed) {
    options(datashield.seed = seed)
}
```
