## Supplementary material for "dsSurvival: Privacy preserving survival models for federated individual patient meta-analysis in DataSHIELD": Server side dsSurvival code: setup.R

```
#-------------------------------------------------------------------------------
# Copyright (c) 2019-2020 University of Newcastle upon Tyne. All rights reserved.
#
# This program and the accompanying materials
# are made available under the terms of the GNU Public License v3.0.
#
# You should have received a copy of the GNU General Public License
# along with this program.  If not, see <http://www.gnu.org/licenses/>.
#-------------------------------------------------------------------------------
#
# Datashield test suite set up
#

context("setup - start")

library(RANN)
library(nlme)
library(stringr)
library(lme4)

library(devtools)
library(testthat)
library(dsBase)
library(dsBaseClient)
require('DSI')
require('DSOpal')


source("disclosure/set_disclosure_settings.R")
source("random/set_random_seed_settings.R")

context("setup - done")
```
