## Supplementary material for "dsSurvival: Privacy preserving survival models for federated individual patient meta-analysis in DataSHIELD": Server side dsSurvival code: test-smk-listDisclosureSettingsDS.R

```
#-------------------------------------------------------------------------------
# Copyright (c) 2019-2020 University of Newcastle upon Tyne. All rights reserved.
#
# This program and the accompanying materials
# are made available under the terms of the GNU Public License v3.0.
#
# You should have received a copy of the GNU General Public License
# along with this program.  If not, see <http://www.gnu.org/licenses/>.
#-------------------------------------------------------------------------------

#
# Set up
#

context("listDisclosureSettingsDS::smk::setup")

set.standard.disclosure.settings()

#
# Tests
#

context("listDisclosureSettingsDS::smk")
test_that("simple listDisclosureSettingsDS", {
    res <- listDisclosureSettingsDS()

    expect_equal(class(res), "list")
    expect_length(res, 9)

    res.names <- names(res)
    expect_length(res.names, 9)

    expect_true("nfilter.tab" %in% res.names)
    expect_true("nfilter.subset" %in% res.names)
    expect_true("nfilter.glm" %in% res.names)
    expect_true("nfilter.string" %in% res.names)
    expect_true("nfilter.stringShort" %in% res.names)
    expect_true("nfilter.kNN" %in% res.names)
    expect_true("nfilter.levels" %in% res.names)
    expect_true("nfilter.noise" %in% res.names)
    expect_true("nfilter.privacy.old" %in% res.names)

    expect_equal(class(res$nfilter.tab), "character")
    expect_equal(class(res$nfilter.subset), "character")
    expect_equal(class(res$nfilter.glm), "character")
    expect_equal(class(res$nfilter.string), "character")
    expect_equal(class(res$nfilter.stringShort), "character")
    expect_equal(class(res$nfilter.kNN), "character")
    expect_equal(class(res$nfilter.levels), "character")
    expect_equal(class(res$nfilter.noise), "character")
    expect_equal(class(res$nfilter.privacy.old), "character")

    expect_equal(res$nfilter.tab, "3")
    expect_equal(res$nfilter.subset, "3")
    expect_equal(res$nfilter.glm, "0.33")
    expect_equal(res$nfilter.string, "80")
    expect_equal(res$nfilter.stringShort, "20")
    expect_equal(res$nfilter.kNN, "3")
    expect_equal(res$nfilter.levels, "0.33")
    expect_equal(res$nfilter.noise, "0.25")
    expect_equal(res$nfilter.privacy.old, "5")
})

#
# Done
#

context("listDisclosureSettingsDS::smk::shutdown")

context("listDisclosureSettingsDS::smk::done")
```
