## Supplementary material for "dsSurvival: Privacy preserving survival models for federated individual patient meta-analysis in DataSHIELD": Vignette with example code on how to use dsSurvival

We also plot privacy preserving survival curves. Please note that is work in progress and is only available on a separate development branch. There will be a full release in v1.1.0.

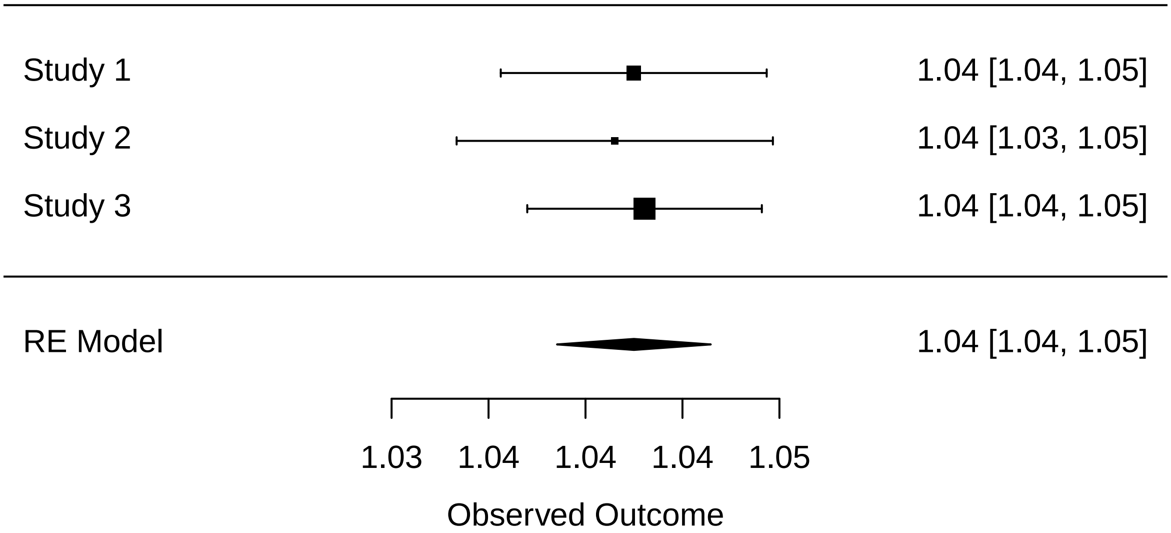


Figure 1: Example forest plot of meta-analyzed hazard ratios.

```
dsSurvivalClient::ds.survfit(formula='surv_object~1', objectname='survfit_object')

dsSurvivalClient::ds.plotsurvfit(formula = 'survfit_object')

## NULL
```

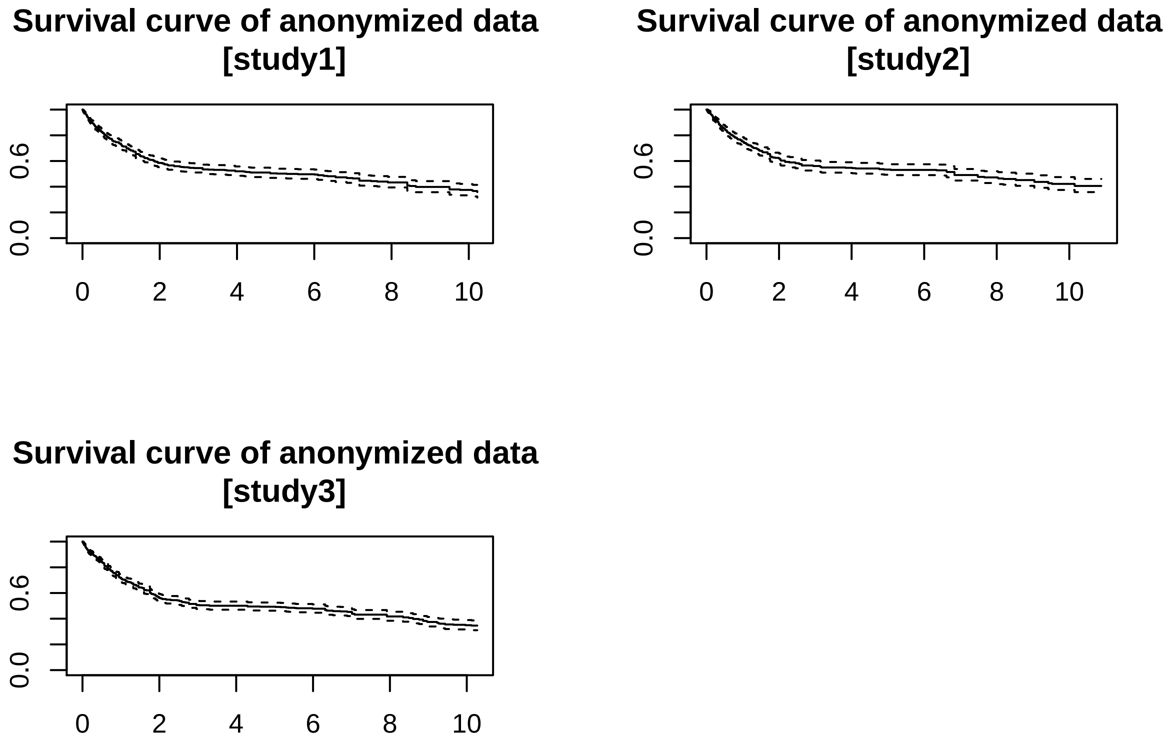


Figure 2: Privacy preserving survival curves.

```
## $study1
## Call: survfit(formula = formula)
##
## records       n  events  median 0.95LCL 0.95UCL
## 2060.00  886.00  426.00    5.29    3.25    7.17
##
## $study2
## Call: survfit(formula = formula)
##
## records       n  events  median 0.95LCL 0.95UCL
## 1640.00  659.00  300.00    6.83    4.76    9.04
##
## $study3
## Call: survfit(formula = formula)
##
## records       n  events  median 0.95LCL 0.95UCL
## 2688.00 1167.00  578.00    4.30    2.58    6.36
```

### 11 Acknowledgements

We acknowledge the help and support of the DataSHIELD technical team. We are especially grateful to Yannick Marcon, Paul Burton, Demetris Avraam, Stuart Wheater, Patricia Ryser-Welch, Xavier Escriba, Juan Gonzalez and Wolfgang Vichtbauer for fruitful discussions and feedback.

### 12 References

- https://github.com/datashield
- http://www.metafor-project.org
- https://github.com/neelsoumya/dsBase
- https://github.com/neelsoumya/dsBaseClient
- https://github.com/neelsoumya/dsSurvival
- https://github.com/neelsoumya/dsSurvivalClient
- https://github.com/neelsoumya/datashield_testing_basic
