## Supplementary material for "dsSurvival: Privacy preserving survival models for federated individual patient meta-analysis in DataSHIELD": Bookdown and documentation for using dsSurvival: _output.yml

```
bookdown::gitbook:
  css: style.css
  config:
    toc:
      before: |
        <li><a href="./">A Minimal Book Example</a></li>
      after: |
        <li><a href="https://github.com/rstudio/bookdown" target="blank">Published with bookdown</a></li>
    download: ["pdf", "epub"]
bookdown::pdf_book:
  includes:
    in_header: preamble.tex
  latex_engine: xelatex
  citation_package: natbib
  keep_tex: yes
bookdown::epub_book: default
```
