## Supplementary material for "dsSurvival: Privacy preserving survival models for federated individual patient meta-analysis in DataSHIELD": Bookdown and documentation for using dsSurvival: better_forestplot.R

```
# code to generate better plots by Thodoris 

library(DSI)
library(DSOpal) 
library(devtools)
library(dsBaseClient)
library(meta)
library(dplyr)
library(pheatmap)
library(gridExtra)
library(dsSurvivalClient)


rm(list=ls())

builder <- DSI::newDSLoginBuilder()
#CLUSTER simulated data
builder$append(server = "study1", 
               url = "http://192.168.56.100:8080/", 
               user = "administrator", password = "datashield_test&", 
               table = "CLUSTER.CLUSTER_SLO1", driver = "OpalDriver")
builder$append(server = "study2", 
               url = "http://192.168.56.100:8080/", 
               user = "administrator", password = "datashield_test&", 
               table = "CLUSTER.CLUSTER_SLO2", driver = "OpalDriver")
builder$append(server = "study3",
               url = "http://192.168.56.100:8080/", 
               user = "administrator", password = "datashield_test&", 
               table = "CLUSTER.CLUSTER_SLO3", driver = "OpalDriver")
builder$append(server = "study4",
               url = "http://192.168.56.100:8080/", 
               user = "administrator", password = "datashield_test&", 
               table = "CLUSTER.CLUSTER_INT1", driver = "OpalDriver")
builder$append(server = "study5",
               url = "http://192.168.56.100:8080/", 
               user = "administrator", password = "datashield_test&", 
               table = "CLUSTER.CLUSTER_INT2", driver = "OpalDriver")
builder$append(server = "study6",
               url = "http://192.168.56.100:8080/", 
               user = "administrator", password = "datashield_test&", 
               table = "CLUSTER.CLUSTER_INT3", driver = "OpalDriver")

logindata <- builder$build()

# Log onto the remote Opal training servers
connections <- DSI::datashield.login( logins = logindata
                                    , assign = TRUE
                                    , symbol = "D") 
#Example without heterogeneity
OR <- ds.glmSLMA(formula = "diabetes ~ Male",
                 dataName="D",
                 family="binomial",
                 newobj="summaryOR",
                 combine.with.metafor=T)

OR$SLMA.pooled.ests.matrix[2,]


#Example with heterogeneity 
MD <- ds.glmSLMA(formula = "incid_rate ~ trtGrp",
                 dataName="D",
                 family="gaussian",
                 newobj="summaryMD",
                 combine.with.metafor=T)

MD$SLMA.pooled.ests.matrix[2,]

MD$betamatrix.valid[2,]

MD$sematrix.valid[2,]

studylabels <- c("center1","center2","center3","center4","center5","center6")

#Variables should be formatted as a matrix
effects <- matrix(c(MD$betamatrix.valid[2,]
                   ,MD$sematrix.valid[2,])
                   ,dimnames = list(studylabels,c("TE","seTE"))
                   ,nrow=length(studylabels)
                   ,ncol=2)

mtotal <- metagen(TE=TE, seTE=seTE, data=as.data.frame(effects)
                  , method.tau = 'REML', prediction=T, hakn=F)

meta::forest(mtotal, xlim = c(12,16))

meta::baujat(mtotal)

heatPlot <- function (x, method.tau = 'REML'){
  bootstrap <- function(x) {
    #x needs a label column
    nrs <- nrow(x)
    res <- Reduce(function(ag, xr){
      ag[[rownames(x)[xr]]] <- x[-xr,]
      return(ag)
    }, 1:nrs, list() )
    res$total <- x
    return(res)
  }
  effs <- bootstrap(x)
  res <- lapply(effs, function(ef){
    mt <- metagen(TE=ef[,"TE"], seTE=ef[,"seTE"]
                  , method.tau = method.tau, prediction=T)
    return(list(I2=mt$I2, tau=mt$tau, Q=mt$Q))
  })
  labels <- rownames(as.matrix(res))
  out <- matrix(unlist(res)
               ,ncol = length(labels)
               ,dimnames = list(c("I2","tau","Q"),labels))
  
  plist <- list(pheatmap(t(out[1,]), labels_row=expression(I^2),
                         show_colnames=F, 
                         silent=T,
                         display_numbers = T,
                         cluster_cols = F,cluster_rows=F,
                         color = colorRampPalette(c("firebrick3", "yellow", "blue"))(150)
                         )$gtable
               ,pheatmap(t(out[2,]),
                         color = colorRampPalette(c("firebrick3", "yellow", "blue"))(150),
                         labels_row=expression(tau), show_colnames=F,silent=T,display_numbers = T,cluster_cols = F,cluster_rows=F)$gtable
               ,pheatmap(t(out[3,]), 
                         color = colorRampPalette(c("firebrick3", "yellow", "blue"))(150),
                         labels_row="Q", display_numbers = T,cluster_cols = F,silent=T,cluster_rows=F)$gtable
  )
  do.call("grid.arrange", c(plist, ncol=1))
  return(out)
}

heatPlot(effects)
```
