## Supplementary material for "dsSurvival: Privacy preserving survival models for federated individual patient meta-analysis in DataSHIELD": Bookdown and documentation for using dsSurvival: book.bib

@Book{xie2015,
  title = {Dynamic Documents with {R} and knitr},
  author = {Yihui Xie},
  publisher = {Chapman and Hall/CRC},
  address = {Boca Raton, Florida},
  year = {2015},
  edition = {2nd},
  note = {ISBN 978-1498716963},
  url = {http://yihui.org/knitr/},
}

@misc{Banerjeef,
author = {Banerjee, Soumya and Bishop, Tom},
doi = {10.5281/zenodo.4806588},
mendeley-groups = {cam{\_}project,My{\_}PAPERS,Software},
title = {{neelsoumya/dsSurvivalClient: Survival functions (client side) for DataSHIELD}},
url = {https://github.com/neelsoumya/dsSurvivalClient},
year = {2021}
}

@software{soumya_banerjee_2021_4917552,
  author       = {Soumya Banerjee and
                  Tom R. P. Bishop},
  title        = {{neelsoumya/dsSurvival: v1.0.0 Survival models in 
                   DataSHIELD}},
  month        = jun,
  year         = 2021,
  publisher    = {Zenodo},
  version      = {v1.0.0},
  doi          = {10.5281/zenodo.4917552},
  url          = {https://doi.org/10.5281/zenodo.4806871}
}

@article{Viechtbauer2010,
abstract = {Epigenetic biomarkers of aging (the “epigenetic clock”) have the potential to address puzzling findings surrounding mortality rates and incidence of cardio-metabolic disease such as: (1) women consistently exhibiting lower mortality than men despite having higher levels of morbidity; (2) racial/ethnic groups having different mortality rates even after adjusting for socioeconomic differences; (3) the black/white mortality cross-over effect in late adulthood; and (4) Hispanics in the United States having a longer life expectancy than Caucasians despite having a higher burden of traditional cardio-metabolic risk factors. We analyzed blood, saliva, and brain samples from seven different racial/ethnic groups. We assessed the intrinsic epigenetic age acceleration of blood (independent of blood cell counts) and the extrinsic epigenetic aging rates of blood (dependent on blood cell counts and tracks the age of the immune system). In blood, Hispanics and Tsimane Amerindians have lower intrinsic but higher extrinsic epigenetic aging rates than Caucasians. African-Americans have lower extrinsic epigenetic aging rates than Caucasians and Hispanics but no differences were found for the intrinsic measure. Men have higher epigenetic aging rates than women in blood, saliva, and brain tissue. Epigenetic aging rates are significantly associated with sex, race/ethnicity, and to a lesser extent with CHD risk factors, but not with incident CHD outcomes. These results may help elucidate lower than expected mortality rates observed in Hispanics, older African-Americans, and women.},
archivePrefix = {arXiv},
arxivId = {arXiv:1501.0228},
author = {Viechtbauer, Wolfgang},
doi = {10.18637/jss.v036.i03},
eprint = {arXiv:1501.0228},
isbn = {1548-7660},
issn = {1548-7660},
journal = {J. Stat. Softw.},
mendeley-groups = {Periphery{\_}project,cam{\_}project},
month = {aug},
number = {3},
pages = {1--48},
pmid = {18291371},
title = {{Conducting Meta-Analyses in R with the metafor Package}},
url = {http://www.jstatsoft.org/v36/i03/},
volume = {36},
year = {2010}
}


@article{Viechtbauer,
author = {Viechtbauer, Wolfgang},
mendeley-groups = {Periphery{\_}project,Machine Learning,cam{\_}project},
publisher = {Comprehensive R Archive Network (CRAN)},
title = {{Meta-Analysis Package for R. Package ‘metafor'. R package version 1.6-1.}},
url = {https://cran.r-project.org/web/packages/metafor/index.html http://cran.r-project.org/web/packages/metafor/index.html},
year = {2011}
}

@article{Banerjee2022,
abstract = {Achieving sufficient statistical power in a survival analysis usually requires large amounts of data from different sites. Sensitivity of individual-level data, ethical and practical considerations regarding data sharing across institutions could be a potential challenge for achieving this added power. Hence we implemented a federated meta-analysis approach of survival models in DataSHIELD, where only anonymous aggregated data are shared across institutions, while simultaneously allowing for exploratory, interactive modelling. In this case, meta-analysis techniques to combine analysis results from each site are a solution, but a manual analysis workflow hinders exploration. Thus, the aim is to provide a framework for performing meta-analysis of Cox regression models across institutions without manual analysis steps for the data providers. We introduce a package (dsSurvival) which allows privacy preserving meta-analysis of survival models, including the calculation of hazard ratios. Our tool can be of great use in biomedical research where there is a need for building survival models and there are privacy concerns about sharing data. A tutorial in bookdown format with code, diagnostics, plots and synthetic data is available here: https://neelsoumya.github.io/dsSurvivalbookdown/ All code is available from the following repositories: https://github.com/neelsoumya/dsSurvivalClient/ https://github.com/neelsoumya/dsSurvival/ {\#}{\#}{\#} Competing Interest Statement The authors have declared no competing interest.},
author = {Banerjee, Soumya and Sofack, Ghislain and Papakonstantinou, Thodoris and Avraam, Demetris and Burton, Paul and Z{\"{o}}ller, Daniela and Bishop, Tom RP},
doi = {10.1101/2022.01.04.471418},
journal = {bioRxiv},
mendeley-groups = {cam{\_}project,My{\_}PAPERS},
month = {jan},
pages = {2022.01.04.471418},
publisher = {Cold Spring Harbor Laboratory},
title = {{dsSurvival: Privacy preserving survival models for federated individual patient meta-analysis in DataSHIELD}},
year = {2022}
}
