## Supplementary material for "dsSurvival: Privacy preserving survival models for federated individual patient meta-analysis in DataSHIELD": Bookdown and documentation for using dsSurvival: combined_vignette_MRCserver.pdf

*2021-11-30*

### Chapter 1 Prerequisites

This is a bookdown with executable code demonstrating how to use the dsSurvival package to create privacy preserving survival models in DataSHIELD. dsSurvival builds privacy preserving survival models.

DataSHIELD is a platform for federated analysis of private data. DataSHIELD has a client-server architecture and this package has a client side and server side component.

The client side package is called dsSurvivalClient:

https://github.com/neelsoumya/dsSurvivalClient

The server side package is called dsSurvival:

https://github.com/neelsoumya/dsSurvival

Please install R. Then install R Studio

https://www.rstudio.com/products/rstudio/download/preview/

We assume that the reader is familiar with R and DataSHIELD.

We outline some of the other prerequisites that need to be installed below:

```
install.packages('devtools')

library(devtools)

devtools::install_github('neelsoumya/dsSurvivalClient')

devtools::install_github('datashield/dsBaseClient@6.1.1')

install.packages('rmarkdown')

install.packages('knitr')

install.packages('tinytex')

install.packages('metafor')

install.packages('DSOpal')

install.packages('DSI')

install.packages('opalr')
```

### Chapter 2 Introduction #

This is a document that outlines a vignette for implementing privacy preserving survival models and meta-analyzing hazard ratios in the DataSHIELD platform.

We used the **bookdown** package (Xie 2021), R Markdown and **knitr** (Xie 2015) for this document. Our package **dsSurvival** (Banerjee and Bishop 2021a)(Banerjee and Bishop 2021b) uses the **metafor** package for meta-analysis (Viechtbauer 2010).

#### 2.1 Survival models

Survival models are used extensively in healthcare. Previously building survival models in DataSHIELD involved building piecewise exponential regression models. This is an approximation and involves having to define appropriate time buckets. A lack of familiarity with this approach also makes people suspicious.

The scope of our package implementation is restricted to being study-level meta-analysis (SLMA) rather than full likelihood.

### Chapter 3 Computational workflow

The computational steps are outlined below. The first step is connecting to the server and loading the survival data.

```
library(knitr)
library(rmarkdown)
library(tinytex)
library(survival)
library(metafor)
library(ggplot2)
library(dsSurvivalClient)
require('DSI')
require('DSOpal')
require('dsBaseClient')

builder <- DSI::newDSLoginBuilder()

builder$append(server="server1", url="https://opal-sandbox.mrc-epid.cam.ac.uk",
                user="dsuser", password="P@ssw0rd",
               table = "SURVIVAL.EXPAND_NO_MISSING1")

builder$append(server="server2", url="https://opal-sandbox.mrc-epid.cam.ac.uk",
               user="dsuser", password="P@ssw0rd",
               table = "SURVIVAL.EXPAND_NO_MISSING2")

builder$append(server="server3", url="https://opal-sandbox.mrc-epid.cam.ac.uk",
               user="dsuser", password="P@ssw0rd",
               table = "SURVIVAL.EXPAND_NO_MISSING3")

logindata <- builder$build()

connections <- DSI::datashield.login(logins = logindata, assign = TRUE, symbol = "D")
```

#### 3.1 Creating server-side variables for survival analysis

We now outline the steps for analysing survival data.

We show this using synthetic data. There are 3 data sets that are held on the same server but can be considered to be on separate servers/sites.

The **cens** variable has the event information and the **survtime** variable has the time information. There is also age and gender information in the variables named **age** and **female**, respectively.

We will look at how age and gender affect survival time and then meta-analyze the hazard ratios from the survival model.

- make sure that the outcome variable is numeric

```
ds.asNumeric(x.name = "D$cens",
             newobj = "EVENT",
             datasources = connections)

ds.asNumeric(x.name = "D$survtime",
             newobj = "SURVTIME",
             datasources = connections)
```

- convert time id variable to a factor

```
ds.asFactor(input.var.name = "D$time.id",
            newobj = "TID",
            datasources = connections)
```

- create in the server-side the log(survtime) variable

```
ds.log(x = "D$survtime",
       newobj = "log.surv",
       datasources = connections)
```

- create start time variable

```
ds.asNumeric(x.name = "D$starttime",
             newobj = "STARTTIME",
             datasources = connections)
ds.asNumeric(x.name = "D$endtime",
             newobj = "ENDTIME",
             datasources = connections)
```

#### 3.2 Create survival object and call ds.coxph.SLMA()

There are two options to generate the survival object. You can generate it separately or in line.

If a survival object is generated separately, it is stored on the server and can be used later in an assign function ( ds.coxphSLMAassign() ). The motivation for creating the model on the server side is inspired from the ds.glmassign functions. This allows the survival model to be stored on the server and can be used later for diagnostics.

- use constructed Surv object in *ds.coxph.SLMA()*

```
dsSurvivalClient::ds.Surv(time='STARTTIME', time2='ENDTIME',
                          event = 'EVENT', objectname='surv_object',
                          type='counting')

coxph_model_full <- dsSurvivalClient::ds.coxph.SLMA(formula = 'surv_object~D$age+D$femal
```

- use direct inline call to *survival::Surv()*

```
dsSurvivalClient::ds.coxph.SLMA(formula = 'survival::Surv(time=SURVTIME,event=EVENT)~D$a
                                dataName = 'D',
                                datasources = connections)
```

- call with *survival::strata()*

The strata() option allows us to relax some of the proportional hazards assumptions. It allows fitting of a separate baseline hazard function within each strata.

```
coxph_model_strata <- dsSurvivalClient::ds.coxph.SLMA(formula = 'surv_object~D$age +
                                    survival::strata(D$female)')
summary(coxph_model_strata)
```

#### 3.3 Summary of survival objects

We can also summarize a server-side object of type *survival::Surv()* using a call to *ds.coxphSummary()*. This will provide a non-disclosive summary of the server-side object. The server-side survival object can be created using ds.coxphSLMAassign(). An example call is shown below:

```
dsSurvivalClient::ds.coxphSummary(x = 'coxph_serverside')
```

#### 3.4 Diagnostics for Cox proportional hazards models

We have also created functions to test for the assumptions of Cox proportional hazards models. This requires a call to the function ds.cox.zphSLMA. Before the call, a server-side object has to be created using the assign function ds.coxphSLMAassign().

All the function calls are shown below:

```
dsSurvivalClient::ds.coxphSLMAassign(formula = 'surv_object~D$age+D$female',
                                    objectname = 'coxph_serverside')

dsSurvivalClient::ds.cox.zphSLMA(fit = 'coxph_serverside')

dsSurvivalClient::ds.coxphSummary(x = 'coxph_serverside')
```

These diagnostics can allow an analyst to determine if the proportional hazards assumption in Cox proportional hazards models is satisfied. If the p-values shown below are greater than 0.05 for any co-variate, then the proportional hazards assumption is correct for that co-variate.

If the proportional hazards assumptions are violated (p-values less than 0.05), then the analyst will have to modify the model. Modifications may include introducing strata or using time-dependent covariates. Please see the links below for more information on this:

- https://stats.stackexchange.com/questions/317336/interpreting-r-coxph-cox-zph
- https://stats.stackexchange.com/questions/144923/extended-cox-model-and-cox-zph/238964#238964

A diagnostic summary is shown below.

```
## surv_object~D$age+D$female
```

```
## NULL
```

```
##
  Aggregating study2 (cox.zphSLMADS("coxph_serverside", "km", TRUE, FALSE, TRUE)) [==>--
  Aggregating study3 (cox.zphSLMADS("coxph_serverside", "km", TRUE, FALSE, TRUE)) [===>-
  Aggregated (cox.zphSLMADS("coxph_serverside", "km", TRUE, FALSE, TRUE)) [=============
```

```
## $study1
##          chisq df    p
## D$age    1.022  1 0.31
## D$female 0.364  1 0.55
## GLOBAL   1.239  2 0.54
##
## $study2
##              chisq df    p
## D$age    -1389.472  1 1.00
## D$female     0.591  1 0.44
## GLOBAL    -857.492  2 1.00
##
## $study3
##          chisq df       p
## D$age    15.27  1 9.3e-05
## D$female  8.04  1  0.0046
## GLOBAL   23.31  2 8.7e-06
```

```
##
  Aggregating study3 (coxphSummaryDS("coxph_serverside")) [====================>------
  Aggregated (coxphSummaryDS("coxph_serverside")) [====================================
```

```
## $study1
## Call:
## survival::coxph(formula = formula, data = dataTable, weights = weights,
##     ties = ties, singular.ok = singular.ok, model = model, x = x,
##     y = y)
##
##   n= 2060, number of events= 426
##
##                coef exp(coef)  se(coef)      z
## D$age      0.041609  1.042487  0.003498 11.894
## D$female1 -0.660002  0.516850  0.099481 -6.634
##           Pr(>|z|)
## D$age      < 2e-16 ***
## D$female1 3.26e-11 ***
## ---
## Signif. codes:
## 0 '***' 0.001 '**' 0.01 '*' 0.05 '.' 0.1 ' ' 1
##
##           exp(coef) exp(-coef) lower .95 upper .95
## D$age        1.0425     0.9592    1.0354    1.0497
## D$female1    0.5169     1.9348    0.4253    0.6281
##
## Concordance= 0.676  (se = 0.014 )
## Likelihood ratio test= 170.7  on 2 df,   p=<2e-16
## Wald test            = 168.2  on 2 df,   p=<2e-16
## Score (logrank) test = 166.3  on 2 df,   p=<2e-16
##
##
## $study2
## Call:
## survival::coxph(formula = formula, data = dataTable, weights = weights,
##     ties = ties, singular.ok = singular.ok, model = model, x = x,
##     y = y)
##
##   n= 1640, number of events= 300
##
##               coef exp(coef) se(coef)      z
## D$age      0.04067   1.04151  0.00416  9.776
```

```
## D$female1 -0.62756   0.53389  0.11767 -5.333
##           Pr(>|z|)
## D$age      < 2e-16 ***
## D$female1 9.66e-08 ***
## ---
## Signif. codes:
## 0 '***' 0.001 '**' 0.01 '*' 0.05 '.' 0.1 ' ' 1
##
##           exp(coef) exp(-coef) lower .95 upper .95
## D$age        1.0415     0.9601    1.0331    1.0500
## D$female1    0.5339     1.8730    0.4239    0.6724
##
## Concordance= 0.674  (se = 0.017 )
## Likelihood ratio test= 117.8  on 2 df,   p=<2e-16
## Wald test            = 115.2  on 2 df,   p=<2e-16
## Score (logrank) test = 116.4  on 2 df,   p=<2e-16
##
##
## $study3
## Call:
## survival::coxph(formula = formula, data = dataTable, weights = weights,
##     ties = ties, singular.ok = singular.ok, model = model, x = x,
##     y = y)
##
##   n= 2688, number of events= 578
##
##                coef exp(coef)  se(coef)      z
## D$age      0.042145  1.043045  0.003086 13.655
## D$female1 -0.599238  0.549230  0.084305 -7.108
##           Pr(>|z|)
## D$age      < 2e-16 ***
## D$female1 1.18e-12 ***
## ---
## Signif. codes:
## 0 '***' 0.001 '**' 0.01 '*' 0.05 '.' 0.1 ' ' 1
##
##           exp(coef) exp(-coef) lower .95 upper .95
## D$age        1.0430     0.9587    1.0368    1.0494
```

```
## D$female1    0.5492     1.8207    0.4656    0.6479
##
## Concordance= 0.699  (se = 0.011 )
## Likelihood ratio test= 227.9  on 2 df,   p=<2e-16
## Wald test            = 228.4  on 2 df,   p=<2e-16
## Score (logrank) test = 229.4  on 2 df,   p=<2e-16
```

#### 3.5 Meta-analyze hazard ratios

We now outline how the hazard ratios from the survival models are meta-analyzed. We use the *metafor* package for meta-analysis. We show the summary of an example meta-analysis and a forest plot below. The forest plot shows a basic example of meta-analyzed hazard ratios from a survival model (analyzed in dsSurvivalClient).

The log-hazard ratios and their standard errors from each study can be found after running *ds.coxphSLMA()*

The hazard ratios can then be meta-analyzed by running the commands shown below. These commands get the hazard ratios correspondng to age in the survival model.

```
input_logHR = c(coxph_model_full$study1$coefficients[1,2],
        coxph_model_full$study2$coefficients[1,2],
        coxph_model_full$study3$coefficients[1,2])

input_se    = c(coxph_model_full$study1$coefficients[1,3],
        coxph_model_full$study2$coefficients[1,3],
        coxph_model_full$study3$coefficients[1,3])

metafor::rma(input_logHR, sei = input_se, method = 'REML')
```

A summary of this meta-analyzed model is shown below.

```
##
## Random-Effects Model (k = 3; tau^2 estimator: REML)
##
##   logLik  deviance       AIC       BIC      AICc
##   9.3824  -18.7648  -14.7648  -17.3785   -2.7648
##
## tau^2 (estimated amount of total heterogeneity): 0 (SE = 0.0000)
## tau (square root of estimated tau^2 value):      0
## I^2 (total heterogeneity / total variability):   0.00%
## H^2 (total variability / sampling variability):  1.00
##
## Test for Heterogeneity:
## Q(df = 2) = 0.0880, p-val = 0.9569
##
## Model Results:
##
## estimate      se      zval    pval   ci.lb   ci.ub
##   1.0425  0.0020  515.4456  <.0001  1.0385  1.0465  ***
##
## ---
## Signif. codes:  0 '***' 0.001 '**' 0.01 '*' 0.05 '.' 0.1 ' ' 1
```

We now show a forest plot with the meta-analyzed hazard ratios. The hazard ratios come from the dsSurvivalClient function *ds.coxphSLMA()*. The plot shows the coefficients for age in the survival model. The command is shown below.

```
metafor::forest.rma(x = meta_model, digits = 4)
```

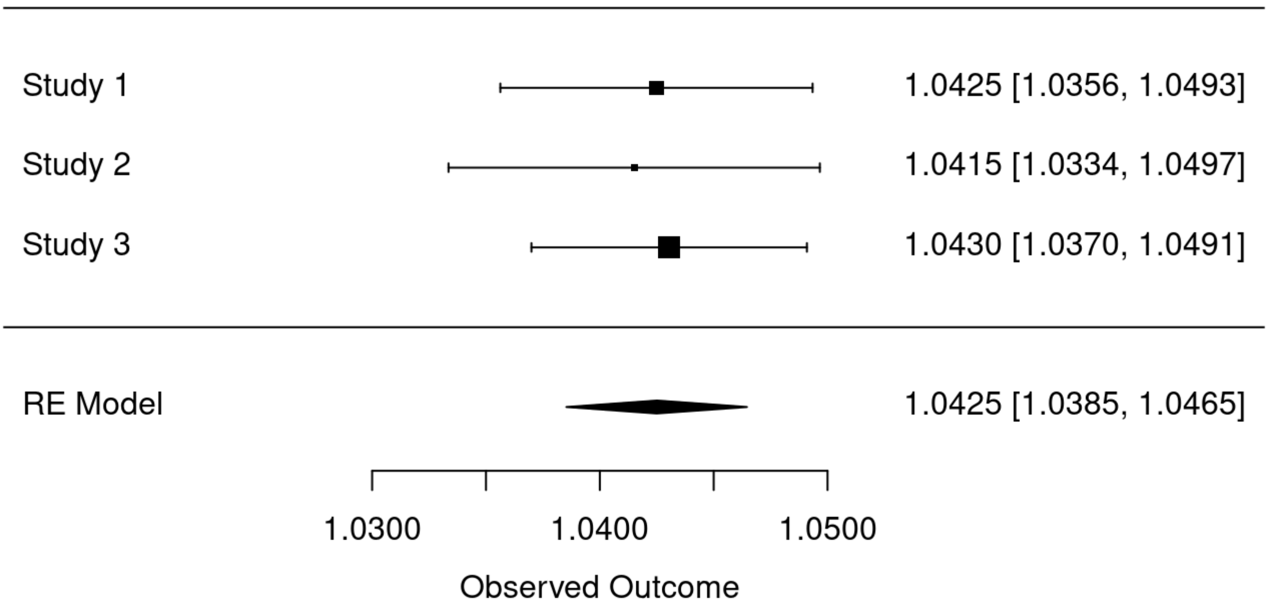


Figure 3.1: Example forest plot of meta-analyzed hazard ratios.

Finally, once you have finished your analysis, you can disconnect from the server(s) using the following command:

```
DSI::datashield.logout(conns = connections)
```

- https://github.com/datashield
- http://www.metafor-project.org
- https://github.com/neelsoumya/dsSurvival
- https://github.com/neelsoumya/dsSurvivalClient

### Chapter 4 Summary

This bookdown shows how to build privacy preserving survival models using dsSurvival in DataSHIELD. You can read more at:

- https://github.com/neelsoumya/dsSurvivalClient
- https://github.com/neelsoumya/dsSurvival
- https://neelsoumya.github.io/dsSurvival_bookdown/
- https://github.com/datashield
