## Supplementary material for "dsSurvival: Privacy preserving survival models for federated individual patient meta-analysis in DataSHIELD": Bookdown and documentation for using dsSurvival: 404.html

Page not found | dsSurvival: Privacy preserving survival models in DataSHIELD


- A Minimal Book Example
- **1** Prerequisites
- **2** Introduction
  - **2.1** Survival models
- **3** Computational workflow
  - **3.1** Creating server-side variables for survival analysis
  - **3.2** Create survival object and call ds.coxph.SLMA()
  - **3.3** Diagnostics for Cox proportional hazards models
  - **3.4** Summary of survival objects
  - **3.5** Meta-analyze hazard ratios
- **4** Summary
- References
- Published with bookdown

### dsSurvival: Privacy preserving survival models in DataSHIELD

### Page not found

The page you requested cannot be found (perhaps it was moved or renamed).

You may want to try searching to find the page's new location, or use
the table of contents to find the page you are looking for.
