## Supplementary material for "dsSurvival: Privacy preserving survival models for federated individual patient meta-analysis in DataSHIELD": Bookdown and documentation for using dsSurvival: applications.html

Chapter 12 Applications | dsSurvival: Survival models in DataSHIELD


- A Minimal Book Example
- **1** Prerequisites
- **2** Introduction
- **3** Computational workflow
- **4** Creating server-side variables for survival analysis
- **5** Create survival object and call ds.coxph.SLMA()
- **6** Summary of survival objects
- **7** Diagnostics for Cox proportional hazards models
- **8** Meta-analyze hazard ratios
- **9** Literature
- **10** Methods
- **11** Dealing with preserving privacy and disclosure checks
  - **11.1** Disclosure related to oversaturated models
  - **11.2** Summary of Cox models with no individual data
- **12** Applications
  - **12.1** Example one
  - **12.2** Example two
- **13** Final Words
- References
- Published with bookdown

### dsSurvival: Survival models in DataSHIELD

### Chapter 12 Applications

Some *significant* applications are demonstrated in this chapter.

#### 12.1 Example one

#### 12.2 Example two
