## Supplementary material for "dsSurvival: Privacy preserving survival models for federated individual patient meta-analysis in DataSHIELD": Bookdown and documentation for using dsSurvival: computational-workflow.html

```
## surv_object~D$age+D$female
```

```
## 
   [-------------------------------------------------------------------------------------]   0% / 0s
  Checking study1 (coxph_serverside <- coxphSLMAassignDS(formula = surv_object ~ D$age + D$female...
  Checking study2 (coxph_serverside <- coxphSLMAassignDS(formula = surv_object ~ D$age + D$female...
  Checking study3 (coxph_serverside <- coxphSLMAassignDS(formula = surv_object ~ D$age + D$female...
  Waiting...  (coxph_serverside <- coxphSLMAassignDS(formula = surv_object ~ D$age + D$female, NU...
  Checking study1 (coxph_serverside <- coxphSLMAassignDS(formula = surv_object ~ D$age + D$female...
  Checking study2 (coxph_serverside <- coxphSLMAassignDS(formula = surv_object ~ D$age + D$female...
  Checking study3 (coxph_serverside <- coxphSLMAassignDS(formula = surv_object ~ D$age + D$female...
  Waiting...  (coxph_serverside <- coxphSLMAassignDS(formula = surv_object ~ D$age + D$female, NU...
  Checking study1 (coxph_serverside <- coxphSLMAassignDS(formula = surv_object ~ D$age + D$female...
  Finalizing assignment study1 (coxph_serverside <- coxphSLMAassignDS(formula = surv_object ~ D$a...
  Checking study2 (coxph_serverside <- coxphSLMAassignDS(formula = surv_object ~ D$age + D$female...
  Finalizing assignment study2 (coxph_serverside <- coxphSLMAassignDS(formula = surv_object ~ D$a...
  Checking study3 (coxph_serverside <- coxphSLMAassignDS(formula = surv_object ~ D$age + D$female...
  Finalizing assignment study3 (coxph_serverside <- coxphSLMAassignDS(formula = surv_object ~ D$a...
  Assigned expr. (coxph_serverside <- coxphSLMAassignDS(formula = surv_object ~ D$age + D$female,...
```

```
## NULL
```

```
## 
   [-------------------------------------------------------------------------------------]   0% / 0s
  Checking study1 (cox.zphSLMADS("coxph_serverside", "km", TRUE, FALSE, TRUE)) [---------]   0% / 0s
  Checking study2 (cox.zphSLMADS("coxph_serverside", "km", TRUE, FALSE, TRUE)) [---------]   0% / 0s
  Checking study3 (cox.zphSLMADS("coxph_serverside", "km", TRUE, FALSE, TRUE)) [---------]   0% / 0s
  Waiting...  (cox.zphSLMADS("coxph_serverside", "km", TRUE, FALSE, TRUE)) [-------------]   0% / 0s
  Checking study1 (cox.zphSLMADS("coxph_serverside", "km", TRUE, FALSE, TRUE)) [---------]   0% / 0s
  Checking study2 (cox.zphSLMADS("coxph_serverside", "km", TRUE, FALSE, TRUE)) [---------]   0% / 0s
  Checking study3 (cox.zphSLMADS("coxph_serverside", "km", TRUE, FALSE, TRUE)) [---------]   0% / 0s
  Waiting...  (cox.zphSLMADS("coxph_serverside", "km", TRUE, FALSE, TRUE)) [-------------]   0% / 0s
  Checking study1 (cox.zphSLMADS("coxph_serverside", "km", TRUE, FALSE, TRUE)) [---------]   0% / 0s
  Getting aggregate study1 (cox.zphSLMADS("coxph_serverside", "km", TRUE, FALSE, TRUE)) []  25% / 0s
  Checking study2 (cox.zphSLMADS("coxph_serverside", "km", TRUE, FALSE, TRUE)) [=>-------]  25% / 0s
  Getting aggregate study2 (cox.zphSLMADS("coxph_serverside", "km", TRUE, FALSE, TRUE)) []  50% / 0s
  Checking study3 (cox.zphSLMADS("coxph_serverside", "km", TRUE, FALSE, TRUE)) [===>-----]  50% / 0s
  Getting aggregate study3 (cox.zphSLMADS("coxph_serverside", "km", TRUE, FALSE, TRUE)) []  75% / 0s
  Aggregated (cox.zphSLMADS("coxph_serverside", "km", TRUE, FALSE, TRUE)) [==============] 100% / 0s
```

```
## $study1
##          chisq df    p
## D$age    1.022  1 0.31
## D$female 0.364  1 0.55
## GLOBAL   1.239  2 0.54
## 
## $study2
##             chisq df    p
## D$age    -254.681  1 1.00
## D$female    0.991  1 0.32
## GLOBAL   -222.531  2 1.00
## 
## $study3
##          chisq df       p
## D$age    15.27  1 9.3e-05
## D$female  8.04  1  0.0046
## GLOBAL   23.31  2 8.7e-06
```

```
## 
   [-------------------------------------------------------------------------------------]   0% / 0s
  Checking study1 (coxphSummaryDS("coxph_serverside")) [---------------------------------]   0% / 0s
  Checking study2 (coxphSummaryDS("coxph_serverside")) [---------------------------------]   0% / 0s
  Checking study3 (coxphSummaryDS("coxph_serverside")) [---------------------------------]   0% / 0s
  Waiting...  (coxphSummaryDS("coxph_serverside")) [-------------------------------------]   0% / 0s
  Checking study1 (coxphSummaryDS("coxph_serverside")) [---------------------------------]   0% / 0s
  Checking study2 (coxphSummaryDS("coxph_serverside")) [---------------------------------]   0% / 0s
  Checking study3 (coxphSummaryDS("coxph_serverside")) [---------------------------------]   0% / 0s
  Waiting...  (coxphSummaryDS("coxph_serverside")) [-------------------------------------]   0% / 0s
  Checking study1 (coxphSummaryDS("coxph_serverside")) [---------------------------------]   0% / 0s
  Getting aggregate study1 (coxphSummaryDS("coxph_serverside")) [=====>------------------]  25% / 0s
  Checking study2 (coxphSummaryDS("coxph_serverside")) [=======>-------------------------]  25% / 0s
  Getting aggregate study2 (coxphSummaryDS("coxph_serverside")) [===========>------------]  50% / 0s
  Checking study3 (coxphSummaryDS("coxph_serverside")) [===============>-----------------]  50% / 0s
  Getting aggregate study3 (coxphSummaryDS("coxph_serverside")) [=================>------]  75% / 0s
  Aggregated (coxphSummaryDS("coxph_serverside")) [======================================] 100% / 0s
```

```
## $study1
## Call:
## survival::coxph(formula = formula, data = dataTable, weights = weights, 
##     ties = ties, singular.ok = singular.ok, model = model, x = x, 
##     y = y)
## 
##   n= 2060, number of events= 426 
## 
##                coef exp(coef)  se(coef)      z
## D$age      0.041609  1.042487  0.003498 11.894
## D$female1 -0.660002  0.516850  0.099481 -6.634
##           Pr(>|z|)    
## D$age      < 2e-16 ***
## D$female1 3.26e-11 ***
## ---
## Signif. codes:  
## 0 '***' 0.001 '**' 0.01 '*' 0.05 '.' 0.1 ' ' 1
## 
##           exp(coef) exp(-coef) lower .95 upper .95
## D$age        1.0425     0.9592    1.0354    1.0497
## D$female1    0.5169     1.9348    0.4253    0.6281
## 
## Concordance= 0.676  (se = 0.014 )
## Likelihood ratio test= 170.7  on 2 df,   p=<2e-16
## Wald test            = 168.2  on 2 df,   p=<2e-16
## Score (logrank) test = 166.3  on 2 df,   p=<2e-16
## 
## 
## $study2
## Call:
## survival::coxph(formula = formula, data = dataTable, weights = weights, 
##     ties = ties, singular.ok = singular.ok, model = model, x = x, 
##     y = y)
## 
##   n= 1640, number of events= 300 
## 
##               coef exp(coef) se(coef)      z Pr(>|z|)
## D$age      0.04067   1.04151  0.00416  9.776  < 2e-16
## D$female1 -0.62756   0.53389  0.11767 -5.333 9.66e-08
##              
## D$age     ***
## D$female1 ***
## ---
## Signif. codes:  
## 0 '***' 0.001 '**' 0.01 '*' 0.05 '.' 0.1 ' ' 1
## 
##           exp(coef) exp(-coef) lower .95 upper .95
## D$age        1.0415     0.9601    1.0331    1.0500
## D$female1    0.5339     1.8730    0.4239    0.6724
## 
## Concordance= 0.674  (se = 0.017 )
## Likelihood ratio test= 117.8  on 2 df,   p=<2e-16
## Wald test            = 115.2  on 2 df,   p=<2e-16
## Score (logrank) test = 116.4  on 2 df,   p=<2e-16
## 
## 
## $study3
## Call:
## survival::coxph(formula = formula, data = dataTable, weights = weights, 
##     ties = ties, singular.ok = singular.ok, model = model, x = x, 
##     y = y)
## 
##   n= 2688, number of events= 578 
## 
##                coef exp(coef)  se(coef)      z
## D$age      0.042145  1.043045  0.003086 13.655
## D$female1 -0.599238  0.549230  0.084305 -7.108
##           Pr(>|z|)    
## D$age      < 2e-16 ***
## D$female1 1.18e-12 ***
## ---
## Signif. codes:  
## 0 '***' 0.001 '**' 0.01 '*' 0.05 '.' 0.1 ' ' 1
## 
##           exp(coef) exp(-coef) lower .95 upper .95
## D$age        1.0430     0.9587    1.0368    1.0494
## D$female1    0.5492     1.8207    0.4656    0.6479
## 
## Concordance= 0.699  (se = 0.011 )
## Likelihood ratio test= 227.9  on 2 df,   p=<2e-16
## Wald test            = 228.4  on 2 df,   p=<2e-16
## Score (logrank) test = 229.4  on 2 df,   p=<2e-16
```

### 3.4 Summary of survival objects

We can also summarize a server-side object of type *survival::Surv()*
using a call to *ds.coxphSummary()*. This will provide a non-disclosive
summary of the server-side object. The server-side survival object can be created using ds.coxphSLMAassign().
An example call is shown below:

```
dsSurvivalClient::ds.coxphSummary(x = 'coxph_serverside')
```

### 3.5 Meta-analyze hazard ratios

We now outline how the hazard ratios from the survival models are meta-analyzed.
We use the *metafor* package for meta-analysis. We show the summary of an
example meta-analysis and a forest plot below. The forest plot shows a basic
example of meta-analyzed hazard ratios from a survival model (analyzed in dsSurvivalClient).

The log-hazard ratios and their standard errors from each study can be
found after running *ds.coxphSLMA()*

The hazard ratios can then be meta-analyzed by running the commands shown below.
These commands get the hazard ratios correspondng to age in the survival model.

```
input_logHR = c(coxph_model_full$server1$coefficients[1,2], 
        coxph_model_full$server2$coefficients[1,2], 
        coxph_model_full$server3$coefficients[1,2])
        
input_se    = c(coxph_model_full$server1$coefficients[1,3], 
        coxph_model_full$server2$coefficients[1,3], 
        coxph_model_full$server3$coefficients[1,3])
        
meta_model <- metafor::rma(input_logHR, sei = input_se, method = 'REML')
```

A summary of this meta-analyzed model is shown below.

```
## 
## Random-Effects Model (k = 3; tau^2 estimator: REML)
## 
##   logLik  deviance       AIC       BIC      AICc 
##   9.3824  -18.7648  -14.7648  -17.3785   -2.7648   
## 
## tau^2 (estimated amount of total heterogeneity): 0 (SE = 0.0000)
## tau (square root of estimated tau^2 value):      0
## I^2 (total heterogeneity / total variability):   0.00%
## H^2 (total variability / sampling variability):  1.00
## 
## Test for Heterogeneity:
## Q(df = 2) = 0.0880, p-val = 0.9569
## 
## Model Results:
## 
## estimate      se      zval    pval   ci.lb   ci.ub 
##   1.0425  0.0020  515.4456  <.0001  1.0385  1.0465  *** 
## 
## ---
## Signif. codes:  0 '***' 0.001 '**' 0.01 '*' 0.05 '.' 0.1 ' ' 1
```

We now show a forest plot with the meta-analyzed hazard ratios. The hazard ratios come
from the dsSurvivalClient function *ds.coxphSLMA()*. The plot shows the coefficients for age in the survival model.
The command is shown below.

```
metafor::forest.rma(x = meta_model, digits = 4)
```

Figure 3.1: Example forest plot of meta-analyzed hazard ratios.

Finally, once you have finished your analysis, you can disconnect from the server(s) using the following command:

```
DSI::datashield.logout(conns = connections)
```

- https://github.com/datashield
- http://www.metafor-project.org
- https://github.com/neelsoumya/dsSurvival
- https://github.com/neelsoumya/dsSurvivalClient
