## Supplementary material for "dsSurvival: Privacy preserving survival models for federated individual patient meta-analysis in DataSHIELD": Bookdown and documentation for using dsSurvival: dealing-with-preserving-privacy-and-disclosure-checks.html

Chapter 11 Dealing with preserving privacy and disclosure checks | dsSurvival: Survival models in DataSHIELD


- A Minimal Book Example
- **1** Prerequisites
- **2** Introduction
- **3** Computational workflow
- **4** Creating server-side variables for survival analysis
- **5** Create survival object and call ds.coxph.SLMA()
- **6** Summary of survival objects
- **7** Diagnostics for Cox proportional hazards models
- **8** Meta-analyze hazard ratios
- **9** Literature
- **10** Methods
- **11** Dealing with preserving privacy and disclosure checks
  - **11.1** Disclosure related to oversaturated models
  - **11.2** Summary of Cox models with no individual data
- **12** Applications
  - **12.1** Example one
  - **12.2** Example two
- **13** Final Words
- References
- Published with bookdown

### dsSurvival: Survival models in DataSHIELD

### Chapter 11 Dealing with preserving privacy and disclosure checks

Disclosure checks are an integral part of DataSHIELD. We outline some of the checks we have built in to ensure privacy of individual level data. We also outline some issues that need to be discussed.

#### 11.1 Disclosure related to oversaturated models

We disallow any Cox models where the number of parameters are greater than a fraction (set to 0.2) of the number of data points. The number of data points is the number of entries (for all patients) in the survival data.

#### 11.2 Summary of Cox models with no individual data

We also present privacy preserving summaries of Cox models and survival objects. These functions reveal no individual level data and present only quantiles.
