## Supplementary material for "dsSurvival: Privacy preserving survival models for federated individual patient meta-analysis in DataSHIELD": Bookdown and documentation for using dsSurvival: diagnostics-for-cox-proportional-hazards-models.html

Chapter 7 Diagnostics for Cox proportional hazards models | dsSurvival: Privacy preserving survival models in DataSHIELD


- A Minimal Book Example
- **1** Prerequisites
- **2** Introduction
- **3** Computational workflow
- **4** Creating server-side variables for survival analysis
- **5** Create survival object and call ds.coxph.SLMA()
- **6** Summary of survival objects
- **7** Diagnostics for Cox proportional hazards models
- **8** Meta-analyze hazard ratios
- **9** Summary
- References
- Published with bookdown

### dsSurvival: Privacy preserving survival models in DataSHIELD

### Chapter 7 Diagnostics for Cox proportional hazards models

We have also created functions to test for the assumptions of
Cox proportional hazards models.

```
dsSurvivalClient::ds.coxphSLMAassign(formula = 'surv_object~D$age+D$female',
                            objectname = 'coxph_serverside')
                
dsSurvivalClient::ds.cox.zphSLMA(fit = 'coxph_serverside')

dsSurvivalClient::ds.coxphSummary(x = 'coxph_serverside')
```

A diagnostic summary is shown below.

```
## surv_object~D$age+D$female
```

```
## 
  Assigning study2 (coxph_serverside <- coxphSLMAassignDS(formula = surv_object ~ D$age + D$femal...
  Assigning study3 (coxph_serverside <- coxphSLMAassignDS(formula = surv_object ~ D$age + D$femal...
  Assigned expr. (coxph_serverside <- coxphSLMAassignDS(formula = surv_object ~ D$age + D$female,...
```

```
## NULL
```

```
## 
  Aggregating study2 (cox.zphSLMADS("coxph_serverside", "km", TRUE, FALSE, TRUE)) [==>---]  50% / 2s
  Aggregating study3 (cox.zphSLMADS("coxph_serverside", "km", TRUE, FALSE, TRUE)) [===>--]  75% / 4s
  Aggregated (cox.zphSLMADS("coxph_serverside", "km", TRUE, FALSE, TRUE)) [==============] 100% / 6s
```

```
## $study1
##          chisq df    p
## D$age    1.022  1 0.31
## D$female 0.364  1 0.55
## GLOBAL   1.239  2 0.54
## 
## $study2
##              chisq df    p
## D$age    -1389.472  1 1.00
## D$female     0.591  1 0.44
## GLOBAL    -857.492  2 1.00
## 
## $study3
##          chisq df       p
## D$age    15.27  1 9.3e-05
## D$female  8.04  1  0.0046
## GLOBAL   23.31  2 8.7e-06
```

```
## 
  Aggregating study2 (coxphSummaryDS("coxph_serverside")) [==============>---------------]  50% / 2s
  Aggregating study3 (coxphSummaryDS("coxph_serverside")) [=====================>--------]  75% / 4s
  Aggregated (coxphSummaryDS("coxph_serverside")) [======================================] 100% / 6s
```

```
## $study1
## Call:
## survival::coxph(formula = formula, data = dataTable, weights = weights, 
##     ties = ties, singular.ok = singular.ok, model = model, x = x, 
##     y = y)
## 
##   n= 2060, number of events= 426 
## 
##                coef exp(coef)  se(coef)      z
## D$age      0.041609  1.042487  0.003498 11.894
## D$female1 -0.660002  0.516850  0.099481 -6.634
##           Pr(>|z|)    
## D$age      < 2e-16 ***
## D$female1 3.26e-11 ***
## ---
## Signif. codes:  
## 0 '***' 0.001 '**' 0.01 '*' 0.05 '.' 0.1 ' ' 1
## 
##           exp(coef) exp(-coef) lower .95 upper .95
## D$age        1.0425     0.9592    1.0354    1.0497
## D$female1    0.5169     1.9348    0.4253    0.6281
## 
## Concordance= 0.676  (se = 0.014 )
## Likelihood ratio test= 170.7  on 2 df,   p=<2e-16
## Wald test            = 168.2  on 2 df,   p=<2e-16
## Score (logrank) test = 166.3  on 2 df,   p=<2e-16
## 
## 
## $study2
## Call:
## survival::coxph(formula = formula, data = dataTable, weights = weights, 
##     ties = ties, singular.ok = singular.ok, model = model, x = x, 
##     y = y)
## 
##   n= 1640, number of events= 300 
## 
##               coef exp(coef) se(coef)      z
## D$age      0.04067   1.04151  0.00416  9.776
## D$female1 -0.62756   0.53389  0.11767 -5.333
##           Pr(>|z|)    
## D$age      < 2e-16 ***
## D$female1 9.66e-08 ***
## ---
## Signif. codes:  
## 0 '***' 0.001 '**' 0.01 '*' 0.05 '.' 0.1 ' ' 1
## 
##           exp(coef) exp(-coef) lower .95 upper .95
## D$age        1.0415     0.9601    1.0331    1.0500
## D$female1    0.5339     1.8730    0.4239    0.6724
## 
## Concordance= 0.674  (se = 0.017 )
## Likelihood ratio test= 117.8  on 2 df,   p=<2e-16
## Wald test            = 115.2  on 2 df,   p=<2e-16
## Score (logrank) test = 116.4  on 2 df,   p=<2e-16
## 
## 
## $study3
## Call:
## survival::coxph(formula = formula, data = dataTable, weights = weights, 
##     ties = ties, singular.ok = singular.ok, model = model, x = x, 
##     y = y)
## 
##   n= 2688, number of events= 578 
## 
##                coef exp(coef)  se(coef)      z
## D$age      0.042145  1.043045  0.003086 13.655
## D$female1 -0.599238  0.549230  0.084305 -7.108
##           Pr(>|z|)    
## D$age      < 2e-16 ***
## D$female1 1.18e-12 ***
## ---
## Signif. codes:  
## 0 '***' 0.001 '**' 0.01 '*' 0.05 '.' 0.1 ' ' 1
## 
##           exp(coef) exp(-coef) lower .95 upper .95
## D$age        1.0430     0.9587    1.0368    1.0494
## D$female1    0.5492     1.8207    0.4656    0.6479
## 
## Concordance= 0.699  (se = 0.011 )
## Likelihood ratio test= 227.9  on 2 df,   p=<2e-16
## Wald test            = 228.4  on 2 df,   p=<2e-16
## Score (logrank) test = 229.4  on 2 df,   p=<2e-16
```
