## Supplementary material for "dsSurvival: Privacy preserving survival models for federated individual patient meta-analysis in DataSHIELD": Bookdown and documentation for using dsSurvival: index.md

## Welcome to GitHub Pages

You can use the [editor on GitHub](https://github.com/neelsoumya/dsSurvival_bookdown/edit/main/docs/index.md) to maintain and preview the content for your website in Markdown files.

Whenever you commit to this repository, GitHub Pages will run [Jekyll](https://jekyllrb.com/) to rebuild the pages in your site, from the content in your Markdown files.

### Markdown

Markdown is a lightweight and easy-to-use syntax for styling your writing. It includes conventions for

```markdown
Syntax highlighted code block

# Header 1
## Header 2
### Header 3

- Bulleted
- List

1. Numbered
2. List

**Bold** and _Italic_ and `Code` text

[Link](url) and ![Image](src)
```

For more details see [GitHub Flavored Markdown](https://guides.github.com/features/mastering-markdown/).

### Jekyll Themes

Your Pages site will use the layout and styles from the Jekyll theme you have selected in your [repository settings](https://github.com/neelsoumya/dsSurvival_bookdown/settings/pages). The name of this theme is saved in the Jekyll `_config.yml` configuration file.

### Support or Contact

Having trouble with Pages? Check out our [documentation](https://docs.github.com/categories/github-pages-basics/) or [contact support](https://support.github.com/contact) and we’ll help you sort it out.
