## Supplementary material for "dsSurvival: Privacy preserving survival models for federated individual patient meta-analysis in DataSHIELD": Bookdown and documentation for using dsSurvival: empty-anchor.js

```
// Hide empty <a> tag within highlighted CodeBlock for screen reader accessibility (see https://github.com/jgm/pandoc/issues/6352#issuecomment-626106786) -->
// v0.0.1
// Written by JooYoung Seo and Atsushi Yasumoto on June 1st, 2020.

document.addEventListener('DOMContentLoaded', function() {
  const codeList = document.getElementsByClassName("sourceCode");
  for (var i = 0; i < codeList.length; i++) {
    var linkList = codeList[i].getElementsByTagName('a');
    for (var j = 0; j < linkList.length; j++) {
      if (linkList[j].innerHTML === "") {
        linkList[j].setAttribute('aria-hidden', 'true');
      }
    }
  }
});
```
