## Supplementary material for "dsSurvival: Privacy preserving survival models for federated individual patient meta-analysis in DataSHIELD": Bookdown and documentation for using dsSurvival: anchor-sections.js

```
// Anchor sections v1.0 written by Atsushi Yasumoto on Oct 3rd, 2020.
document.addEventListener('DOMContentLoaded', function() {
  // Do nothing if AnchorJS is used
  if (typeof window.anchors === 'object' && anchors.hasOwnProperty('hasAnchorJSLink')) {
    return;
  }

  const h = document.querySelectorAll('h1, h2, h3, h4, h5, h6');

  // Do nothing if sections are already anchored
  if (Array.from(h).some(x => x.classList.contains('hasAnchor'))) {
    return null;
  }

  // Use section id when pandoc runs with --section-divs
  const section_id = function(x) {
    return ((x.classList.contains('section') || (x.tagName === 'SECTION'))
            ? x.id : '');
  };

  // Add anchors
  h.forEach(function(x) {
    const id = x.id || section_id(x.parentElement);
    if (id === '' || x.matches(':empty')) {
      return null;
    }
    let anchor = document.createElement('a');
    anchor.href = '#' + id;
    anchor.classList = ['anchor-section'];
    x.classList.add('hasAnchor');
    x.appendChild(anchor);
  });
});
```
