## Supplementary material for "dsSurvival: Privacy preserving survival models for federated individual patient meta-analysis in DataSHIELD": Bookdown and documentation for using dsSurvival: app.min.js

```
(function e(t,n,r){function s(o,u){if(!n[o]){if(!t[o]){var a=typeof require=="function"&&require;if(!u&&a)return a(o,!0);if(i)return i(o,!0);var f=new Error("Cannot find module '"+o+"'");throw f.code="MODULE_NOT_FOUND",f}var l=n[o]={exports:{}};t[o][0].call(l.exports,function(e){var n=t[o][1][e];return s(n?n:e)},l,l.exports,e,t,n,r)}return n[o].exports}var i=typeof require=="function"&&require;for(var o=0;o<r.length;o++)s(r[o]);return s})({1:[function(require,module,exports){if(typeof module==="object"&&typeof module.exports==="object"){module.exports=jQuery}},{}],2:[function(require,module,exports){(function(global){(function(){var undefined;var VERSION="3.10.1";var BIND_FLAG=1,BIND_KEY_FLAG=2,CURRY_BOUND_FLAG=4,CURRY_FLAG=8,CURRY_RIGHT_FLAG=16,PARTIAL_FLAG=32,PARTIAL_RIGHT_FLAG=64,ARY_FLAG=128,REARG_FLAG=256;var DEFAULT_TRUNC_LENGTH=30,DEFAULT_TRUNC_OMISSION="...";var HOT_COUNT=150,HOT_SPAN=16;var LARGE_ARRAY_SIZE=200;var LAZY_FILTER_FLAG=1,LAZY_MAP_FLAG=2;var FUNC_ERROR_TEXT="Expected a function";var PLACEHOLDER="__lodash_placeholder__";var argsTag="[object Arguments]",arrayTag="[object Array]",boolTag="[object Boolean]",dateTag="[object Date]",errorTag="[object Error]",funcTag="[object Function]",mapTag="[object Map]",numberTag="[object Number]",objectTag="[object Object]",regexpTag="[object RegExp]",setTag="[object Set]",stringTag="[object String]",weakMapTag="[object WeakMap]";var arrayBufferTag="[object ArrayBuffer]",float32Tag="[object Float32Array]",float64Tag="[object Float64Array]",int8Tag="[object Int8Array]",int16Tag="[object Int16Array]",int32Tag="[object Int32Array]",uint8Tag="[object Uint8Array]",uint8ClampedTag="[object Uint8ClampedArray]",uint16Tag="[object Uint16Array]",uint32Tag="[object Uint32Array]";var reEmptyStringLeading=/\b__p \+= '';/g,reEmptyStringMiddle=/\b(__p \+=) '' \+/g,reEmptyStringTrailing=/(__e\(.*?\)|\b__t\)) \+\n'';/g;var reEscapedHtml=/&(?:amp|lt|gt|quot|#39|#96);/g,reUnescapedHtml=/[&<>"'`]/g,reHasEscapedHtml=RegExp(reEscapedHtml.source),reHasUnescapedHtml=RegExp(reUnescapedHtml.source);var reEscape=/<%-([\s\S]+?)%>/g,reEvaluate=/<%([\s\S]+?)%>/g,reInterpolate=/<%=([\s\S]+?)%>/g;var reIsDeepProp=/\.|\[(?:[^[\]]*|(["'])(?:(?!\1)[^\n\\]|\\.)*?\1)\]/,reIsPlainProp=/^\w*$/,rePropName=/[^.[\]]+|\[(?:(-?\d+(?:\.\d+)?)|(["'])((?:(?!\2)[^\n\\]|\\.)*?)\2)\]/g;var reRegExpChars=/^[:!,]|[\\^$.*+?()[\]{}|\/]|(^[0-9a-fA-Fnrtuvx])|([\n\r\u2028\u2029])/g,reHasRegExpChars=RegExp(reRegExpChars.source);var reComboMark=/[\u0300-\u036f\ufe20-\ufe23]/g;var reEscapeChar=/\\(\\)?/g;var reEsTemplate=/\$\{([^\\}]*(?:\\.[^\\}]*)*)\}/g;var reFlags=/\w*$/;var reHasHexPrefix=/^0[xX]/;var reIsHostCtor=/^\[object .+?Constructor\]$/;var reIsUint=/^\d+$/;var reLatin1=/[\xc0-\xd6\xd8-\xde\xdf-\xf6\xf8-\xff]/g;var reNoMatch=/($^)/;var reUnescapedString=/['\n\r\u2028\u2029\\]/g;var reWords=function(){var upper="[A-Z\\xc0-\\xd6\\xd8-\\xde]",lower="[a-z\\xdf-\\xf6\\xf8-\\xff]+";return RegExp(upper+"+(?="+upper+lower+")|"+upper+"?"+lower+"|"+upper+"+|[0-9]+","g")}();var contextProps=["Array","ArrayBuffer","Date","Error","Float32Array","Float64Array","Function","Int8Array","Int16Array","Int32Array","Math","Number","Object","RegExp","Set","String","_","clearTimeout","isFinite","parseFloat","parseInt","setTimeout","TypeError","Uint8Array","Uint8ClampedArray","Uint16Array","Uint32Array","WeakMap"];var templateCounter=-1;var typedArrayTags={};typedArrayTags[float32Tag]=typedArrayTags[float64Tag]=typedArrayTags[int8Tag]=typedArrayTags[int16Tag]=typedArrayTags[int32Tag]=typedArrayTags[uint8Tag]=typedArrayTags[uint8ClampedTag]=typedArrayTags[uint16Tag]=typedArrayTags[uint32Tag]=true;typedArrayTags[argsTag]=typedArrayTags[arrayTag]=typedArrayTags[arrayBufferTag]=typedArrayTags[boolTag]=typedArrayTags[dateTag]=typedArrayTags[errorTag]=typedArrayTags[funcTag]=typedArrayTags[mapTag]=typedArrayTags[numberTag]=typedArrayTags[objectTag]=typedArrayTags[regexpTag]=typedArrayTags[setTag]=typedArrayTags[stringTag]=typedArrayTags[weakMapTag]=false;var cloneableTags={};cloneableTags[argsTag]=cloneableTags[arrayTag]=cloneableTags[arrayBufferTag]=cloneableTags[boolTag]=cloneableTags[dateTag]=cloneableTags[float32Tag]=cloneableTags[float64Tag]=cloneableTags[int8Tag]=cloneableTags[int16Tag]=cloneableTags[int32Tag]=cloneableTags[numberTag]=cloneableTags[objectTag]=cloneableTags[regexpTag]=cloneableTags[stringTag]=cloneableTags[uint8Tag]=cloneableTags[uint8ClampedTag]=cloneableTags[uint16Tag]=cloneableTags[uint32Tag]=true;cloneableTags[errorTag]=cloneableTags[funcTag]=cloneableTags[mapTag]=cloneableTags[setTag]=cloneableTags[weakMapTag]=false;var deburredLetters={"À":"A","Á":"A","Â":"A","Ã":"A","Ä":"A","Å":"A","à":"a","á":"a","â":"a","ã":"a","ä":"a","å":"a","Ç":"C","ç":"c","Ð":"D","ð":"d","È":"E","É":"E","Ê":"E","Ë":"E","è":"e","é":"e","ê":"e","ë":"e","Ì":"I","Í":"I","Î":"I","Ï":"I","ì":"i","í":"i","î":"i","ï":"i","Ñ":"N","ñ":"n","Ò":"O","Ó":"O","Ô":"O","Õ":"O","Ö":"O","Ø":"O","ò":"o","ó":"o","ô":"o","õ":"o","ö":"o","ø":"o","Ù":"U","Ú":"U","Û":"U","Ü":"U","ù":"u","ú":"u","û":"u","ü":"u","Ý":"Y","ý":"y","ÿ":"y","Æ":"Ae","æ":"ae","Þ":"Th","þ":"th","ß":"ss"};var htmlEscapes={"&":"&amp;","<":"&lt;",">":"&gt;",'"':"&quot;","'":"&#39;","`":"&#96;"};var htmlUnescapes={"&amp;":"&","&lt;":"<","&gt;":">","&quot;":'"',"&#39;":"'","&#96;":"`"};var objectTypes={function:true,object:true};var regexpEscapes={0:"x30",1:"x31",2:"x32",3:"x33",4:"x34",5:"x35",6:"x36",7:"x37",8:"x38",9:"x39",A:"x41",B:"x42",C:"x43",D:"x44",E:"x45",F:"x46",a:"x61",b:"x62",c:"x63",d:"x64",e:"x65",f:"x66",n:"x6e",r:"x72",t:"x74",u:"x75",v:"x76",x:"x78"};var stringEscapes={"\\":"\\","'":"'","\n":"n","\r":"r","\u2028":"u2028","\u2029":"u2029"};var freeExports=objectTypes[typeof exports]&&exports&&!exports.nodeType&&exports;var freeModule=objectTypes[typeof module]&&module&&!module.nodeType&&module;var freeGlobal=freeExports&&freeModule&&typeof global=="object"&&global&&global.Object&&global;var freeSelf=objectTypes[typeof self]&&self&&self.Object&&self;var freeWindow=objectTypes[typeof window]&&window&&window.Object&&window;var moduleExports=freeModule&&freeModule.exports===freeExports&&freeExports;var root=freeGlobal||freeWindow!==(this&&this.window)&&freeWindow||freeSelf||this;function baseCompareAscending(value,other){if(value!==other){var valIsNull=value===null,valIsUndef=value===undefined,valIsReflexive=value===value;var othIsNull=other===null,othIsUndef=other===undefined,othIsReflexive=other===other;if(value>other&&!othIsNull||!valIsReflexive||valIsNull&&!othIsUndef&&othIsReflexive||valIsUndef&&othIsReflexive){return 1}if(value<other&&!valIsNull||!othIsReflexive||othIsNull&&!valIsUndef&&valIsReflexive||othIsUndef&&valIsReflexive){return-1}}return 0}function baseFindIndex(array,predicate,fromRight){var length=array.length,index=fromRight?length:-1;while(fromRight?index--:++index<length){if(predicate(array[index],index,array)){return index}}return-1}function baseIndexOf(array,value,fromIndex){if(value!==value){return indexOfNaN(array,fromIndex)}var index=fromIndex-1,length=array.length;while(++index<length){if(array[index]===value){return index}}return-1}function baseIsFunction(value){return typeof value=="function"||false}function baseToString(value){return value==null?"":value+""}function charsLeftIndex(string,chars){var index=-1,length=string.length;while(++index<length&&chars.indexOf(string.charAt(index))>-1){}return index}function charsRightIndex(string,chars){var index=string.length;while(index--&&chars.indexOf(string.charAt(index))>-1){}return index}function compareAscending(object,other){return baseCompareAscending(object.criteria,other.criteria)||object.index-other.index}function compareMultiple(object,other,orders){var index=-1,objCriteria=object.criteria,othCriteria=other.criteria,length=objCriteria.length,ordersLength=orders.length;while(++index<length){var result=baseCompareAscending(objCriteria[index],othCriteria[index]);if(result){if(index>=ordersLength){return result}var order=orders[index];return result*(order==="asc"||order===true?1:-1)}}return object.index-other.index}function deburrLetter(letter){return deburredLetters[letter]}function escapeHtmlChar(chr){return htmlEscapes[chr]}function escapeRegExpChar(chr,leadingChar,whitespaceChar){if(leadingChar){chr=regexpEscapes[chr]}else if(whitespaceChar){chr=stringEscapes[chr]}return"\\"+chr}function escapeStringChar(chr){return"\\"+stringEscapes[chr]}function indexOfNaN(array,fromIndex,fromRight){var length=array.length,index=fromIndex+(fromRight?0:-1);while(fromRight?index--:++index<length){var other=array[index];if(other!==other){return index}}return-1}function isObjectLike(value){return!!value&&typeof value=="object"}function isSpace(charCode){return charCode<=160&&(charCode>=9&&charCode<=13)||charCode==32||charCode==160||charCode==5760||charCode==6158||charCode>=8192&&(charCode<=8202||charCode==8232||charCode==8233||charCode==8239||charCode==8287||charCode==12288||charCode==65279)}function replaceHolders(array,placeholder){var index=-1,length=array.length,resIndex=-1,result=[];while(++index<length){if(array[index]===placeholder){array[index]=PLACEHOLDER;result[++resIndex]=index}}return result}function sortedUniq(array,iteratee){var seen,index=-1,length=array.length,resIndex=-1,result=[];while(++index<length){var value=array[index],computed=iteratee?iteratee(value,index,array):value;if(!index||seen!==computed){seen=computed;result[++resIndex]=value}}return result}function trimmedLeftIndex(string){var index=-1,length=string.length;while(++index<length&&isSpace(string.charCodeAt(index))){}return index}function trimmedRightIndex(string){var index=string.length;while(index--&&isSpace(string.charCodeAt(index))){}return index}function unescapeHtmlChar(chr){return htmlUnescapes[chr]}function runInContext(context){context=context?_.defaults(root.Object(),context,_.pick(root,contextProps)):root;var Array=context.Array,Date=context.Date,Error=context.Error,Function=context.Function,Math=context.Math,Number=context.Number,Object=context.Object,RegExp=context.RegExp,String=context.String,TypeError=context.TypeError;var arrayProto=Array.prototype,objectProto=Object.prototype,stringProto=String.prototype;var fnToString=Function.prototype.toString;var hasOwnProperty=objectProto.hasOwnProperty;var idCounter=0;var objToString=objectProto.toString;var oldDash=root._;var reIsNative=RegExp("^"+fnToString.call(hasOwnProperty).replace(/[\\^$.*+?()[\]{}|]/g,"\\$&").replace(/hasOwnProperty|(function).*?(?=\\\()| for .+?(?=\\\])/g,"$1.*?")+"$");var ArrayBuffer=context.ArrayBuffer,clearTimeout=context.clearTimeout,parseFloat=context.parseFloat,pow=Math.pow,propertyIsEnumerable=objectProto.propertyIsEnumerable,Set=getNative(context,"Set"),setTimeout=context.setTimeout,splice=arrayProto.splice,Uint8Array=context.Uint8Array,WeakMap=getNative(context,"WeakMap");var nativeCeil=Math.ceil,nativeCreate=getNative(Object,"create"),nativeFloor=Math.floor,nativeIsArray=getNative(Array,"isArray"),nativeIsFinite=context.isFinite,nativeKeys=getNative(Object,"keys"),nativeMax=Math.max,nativeMin=Math.min,nativeNow=getNative(Date,"now"),nativeParseInt=context.parseInt,nativeRandom=Math.random;var NEGATIVE_INFINITY=Number.NEGATIVE_INFINITY,POSITIVE_INFINITY=Number.POSITIVE_INFINITY;var MAX_ARRAY_LENGTH=4294967295,MAX_ARRAY_INDEX=MAX_ARRAY_LENGTH-1,HALF_MAX_ARRAY_LENGTH=MAX_ARRAY_LENGTH>>>1;var MAX_SAFE_INTEGER=9007199254740991;var metaMap=WeakMap&&new WeakMap;var realNames={};function lodash(value){if(isObjectLike(value)&&!isArray(value)&&!(value instanceof LazyWrapper)){if(value instanceof LodashWrapper){return value}if(hasOwnProperty.call(value,"__chain__")&&hasOwnProperty.call(value,"__wrapped__")){return wrapperClone(value)}}return new LodashWrapper(value)}function baseLodash(){}function LodashWrapper(value,chainAll,actions){this.__wrapped__=value;this.__actions__=actions||[];this.__chain__=!!chainAll}var support=lodash.support={};lodash.templateSettings={escape:reEscape,evaluate:reEvaluate,interpolate:reInterpolate,variable:"",imports:{_:lodash}};function LazyWrapper(value){this.__wrapped__=value;this.__actions__=[];this.__dir__=1;this.__filtered__=false;this.__iteratees__=[];this.__takeCount__=POSITIVE_INFINITY;this.__views__=[]}function lazyClone(){var result=new LazyWrapper(this.__wrapped__);result.__actions__=arrayCopy(this.__actions__);result.__dir__=this.__dir__;result.__filtered__=this.__filtered__;result.__iteratees__=arrayCopy(this.__iteratees__);result.__takeCount__=this.__takeCount__;result.__views__=arrayCopy(this.__views__);return result}function lazyReverse(){if(this.__filtered__){var result=new LazyWrapper(this);result.__dir__=-1;result.__filtered__=true}else{result=this.clone();result.__dir__*=-1}return result}function lazyValue(){var array=this.__wrapped__.value(),dir=this.__dir__,isArr=isArray(array),isRight=dir<0,arrLength=isArr?array.length:0,view=getView(0,arrLength,this.__views__),start=view.start,end=view.end,length=end-start,index=isRight?end:start-1,iteratees=this.__iteratees__,iterLength=iteratees.length,resIndex=0,takeCount=nativeMin(length,this.__takeCount__);if(!isArr||arrLength<LARGE_ARRAY_SIZE||arrLength==length&&takeCount==length){return baseWrapperValue(isRight&&isArr?array.reverse():array,this.__actions__)}var result=[];outer:while(length--&&resIndex<takeCount){index+=dir;var iterIndex=-1,value=array[index];while(++iterIndex<iterLength){var data=iteratees[iterIndex],iteratee=data.iteratee,type=data.type,computed=iteratee(value);if(type==LAZY_MAP_FLAG){value=computed}else if(!computed){if(type==LAZY_FILTER_FLAG){continue outer}else{break outer}}}result[resIndex++]=value}return result}function MapCache(){this.__data__={}}function mapDelete(key){return this.has(key)&&delete this.__data__[key]}function mapGet(key){return key=="__proto__"?undefined:this.__data__[key]}function mapHas(key){return key!="__proto__"&&hasOwnProperty.call(this.__data__,key)}function mapSet(key,value){if(key!="__proto__"){this.__data__[key]=value}return this}function SetCache(values){var length=values?values.length:0;this.data={hash:nativeCreate(null),set:new Set};while(length--){this.push(values[length])}}function cacheIndexOf(cache,value){var data=cache.data,result=typeof value=="string"||isObject(value)?data.set.has(value):data.hash[value];return result?0:-1}function cachePush(value){var data=this.data;if(typeof value=="string"||isObject(value)){data.set.add(value)}else{data.hash[value]=true}}function arrayConcat(array,other){var index=-1,length=array.length,othIndex=-1,othLength=other.length,result=Array(length+othLength);while(++index<length){result[index]=array[index]}while(++othIndex<othLength){result[index++]=other[othIndex]}return result}function arrayCopy(source,array){var index=-1,length=source.length;array||(array=Array(length));while(++index<length){array[index]=source[index]}return array}function arrayEach(array,iteratee){var index=-1,length=array.length;while(++index<length){if(iteratee(array[index],index,array)===false){break}}return array}function arrayEachRight(array,iteratee){var length=array.length;while(length--){if(iteratee(array[length],length,array)===false){break}}return array}function arrayEvery(array,predicate){var index=-1,length=array.length;while(++index<length){if(!predicate(array[index],index,array)){return false}}return true}function arrayExtremum(array,iteratee,comparator,exValue){var index=-1,length=array.length,computed=exValue,result=computed;while(++index<length){var value=array[index],current=+iteratee(value);if(comparator(current,computed)){computed=current;result=value}}return result}function arrayFilter(array,predicate){var index=-1,length=array.length,resIndex=-1,result=[];while(++index<length){var value=array[index];if(predicate(value,index,array)){result[++resIndex]=value}}return result}function arrayMap(array,iteratee){var index=-1,length=array.length,result=Array(length);while(++index<length){result[index]=iteratee(array[index],index,array)}return result}function arrayPush(array,values){var index=-1,length=values.length,offset=array.length;while(++index<length){array[offset+index]=values[index]}return array}function arrayReduce(array,iteratee,accumulator,initFromArray){var index=-1,length=array.length;if(initFromArray&&length){accumulator=array[++index]}while(++index<length){accumulator=iteratee(accumulator,array[index],index,array)}return accumulator}function arrayReduceRight(array,iteratee,accumulator,initFromArray){var length=array.length;if(initFromArray&&length){accumulator=array[--length]}while(length--){accumulator=iteratee(accumulator,array[length],length,array)}return accumulator}function arraySome(array,predicate){var index=-1,length=array.length;while(++index<length){if(predicate(array[index],index,array)){return true}}return false}function arraySum(array,iteratee){var length=array.length,result=0;while(length--){result+=+iteratee(array[length])||0}return result}function assignDefaults(objectValue,sourceValue){return objectValue===undefined?sourceValue:objectValue}function assignOwnDefaults(objectValue,sourceValue,key,object){return objectValue===undefined||!hasOwnProperty.call(object,key)?sourceValue:objectValue}function assignWith(object,source,customizer){var index=-1,props=keys(source),length=props.length;while(++index<length){var key=props[index],value=object[key],result=customizer(value,source[key],key,object,source);if((result===result?result!==value:value===value)||value===undefined&&!(key in object)){object[key]=result}}return object}function baseAssign(object,source){return source==null?object:baseCopy(source,keys(source),object)}function baseAt(collection,props){var index=-1,isNil=collection==null,isArr=!isNil&&isArrayLike(collection),length=isArr?collection.length:0,propsLength=props.length,result=Array(propsLength);while(++index<propsLength){var key=props[index];if(isArr){result[index]=isIndex(key,length)?collection[key]:undefined}else{result[index]=isNil?undefined:collection[key]}}return result}function baseCopy(source,props,object){object||(object={});var index=-1,length=props.length;while(++index<length){var key=props[index];object[key]=source[key]}return object}function baseCallback(func,thisArg,argCount){var type=typeof func;if(type=="function"){return thisArg===undefined?func:bindCallback(func,thisArg,argCount)}if(func==null){return identity}if(type=="object"){return baseMatches(func)}return thisArg===undefined?property(func):baseMatchesProperty(func,thisArg)}function baseClone(value,isDeep,customizer,key,object,stackA,stackB){var result;if(customizer){result=object?customizer(value,key,object):customizer(value)}if(result!==undefined){return result}if(!isObject(value)){return value}var isArr=isArray(value);if(isArr){result=initCloneArray(value);if(!isDeep){return arrayCopy(value,result)}}else{var tag=objToString.call(value),isFunc=tag==funcTag;if(tag==objectTag||tag==argsTag||isFunc&&!object){result=initCloneObject(isFunc?{}:value);if(!isDeep){return baseAssign(result,value)}}else{return cloneableTags[tag]?initCloneByTag(value,tag,isDeep):object?value:{}}}stackA||(stackA=[]);stackB||(stackB=[]);var length=stackA.length;while(length--){if(stackA[length]==value){return stackB[length]}}stackA.push(value);stackB.push(result);(isArr?arrayEach:baseForOwn)(value,function(subValue,key){result[key]=baseClone(subValue,isDeep,customizer,key,value,stackA,stackB)});return result}var baseCreate=function(){function object(){}return function(prototype){if(isObject(prototype)){object.prototype=prototype;var result=new object;object.prototype=undefined}return result||{}}}();function baseDelay(func,wait,args){if(typeof func!="function"){throw new TypeError(FUNC_ERROR_TEXT)}return setTimeout(function(){func.apply(undefined,args)},wait)}function baseDifference(array,values){var length=array?array.length:0,result=[];if(!length){return result}var index=-1,indexOf=getIndexOf(),isCommon=indexOf==baseIndexOf,cache=isCommon&&values.length>=LARGE_ARRAY_SIZE?createCache(values):null,valuesLength=values.length;if(cache){indexOf=cacheIndexOf;isCommon=false;values=cache}outer:while(++index<length){var value=array[index];if(isCommon&&value===value){var valuesIndex=valuesLength;while(valuesIndex--){if(values[valuesIndex]===value){continue outer}}result.push(value)}else if(indexOf(values,value,0)<0){result.push(value)}}return result}var baseEach=createBaseEach(baseForOwn);var baseEachRight=createBaseEach(baseForOwnRight,true);function baseEvery(collection,predicate){var result=true;baseEach(collection,function(value,index,collection){result=!!predicate(value,index,collection);return result});return result}function baseExtremum(collection,iteratee,comparator,exValue){var computed=exValue,result=computed;baseEach(collection,function(value,index,collection){var current=+iteratee(value,index,collection);if(comparator(current,computed)||current===exValue&&current===result){computed=current;result=value}});return result}function baseFill(array,value,start,end){var length=array.length;start=start==null?0:+start||0;if(start<0){start=-start>length?0:length+start}end=end===undefined||end>length?length:+end||0;if(end<0){end+=length}length=start>end?0:end>>>0;start>>>=0;while(start<length){array[start++]=value}return array}function baseFilter(collection,predicate){var result=[];baseEach(collection,function(value,index,collection){if(predicate(value,index,collection)){result.push(value)}});return result}function baseFind(collection,predicate,eachFunc,retKey){var result;eachFunc(collection,function(value,key,collection){if(predicate(value,key,collection)){result=retKey?key:value;return false}});return result}function baseFlatten(array,isDeep,isStrict,result){result||(result=[]);var index=-1,length=array.length;while(++index<length){var value=array[index];if(isObjectLike(value)&&isArrayLike(value)&&(isStrict||isArray(value)||isArguments(value))){if(isDeep){baseFlatten(value,isDeep,isStrict,result)}else{arrayPush(result,value)}}else if(!isStrict){result[result.length]=value}}return result}var baseFor=createBaseFor();var baseForRight=createBaseFor(true);function baseForIn(object,iteratee){return baseFor(object,iteratee,keysIn)}function baseForOwn(object,iteratee){return baseFor(object,iteratee,keys)}function baseForOwnRight(object,iteratee){return baseForRight(object,iteratee,keys)}function baseFunctions(object,props){var index=-1,length=props.length,resIndex=-1,result=[];while(++index<length){var key=props[index];if(isFunction(object[key])){result[++resIndex]=key}}return result}function baseGet(object,path,pathKey){if(object==null){return}if(pathKey!==undefined&&pathKey in toObject(object)){path=[pathKey]}var index=0,length=path.length;while(object!=null&&index<length){object=object[path[index++]]}return index&&index==length?object:undefined}function baseIsEqual(value,other,customizer,isLoose,stackA,stackB){if(value===other){return true}if(value==null||other==null||!isObject(value)&&!isObjectLike(other)){return value!==value&&other!==other}return baseIsEqualDeep(value,other,baseIsEqual,customizer,isLoose,stackA,stackB)}function baseIsEqualDeep(object,other,equalFunc,customizer,isLoose,stackA,stackB){var objIsArr=isArray(object),othIsArr=isArray(other),objTag=arrayTag,othTag=arrayTag;if(!objIsArr){objTag=objToString.call(object);if(objTag==argsTag){objTag=objectTag}else if(objTag!=objectTag){objIsArr=isTypedArray(object)}}if(!othIsArr){othTag=objToString.call(other);if(othTag==argsTag){othTag=objectTag}else if(othTag!=objectTag){othIsArr=isTypedArray(other)}}var objIsObj=objTag==objectTag,othIsObj=othTag==objectTag,isSameTag=objTag==othTag;if(isSameTag&&!(objIsArr||objIsObj)){return equalByTag(object,other,objTag)}if(!isLoose){var objIsWrapped=objIsObj&&hasOwnProperty.call(object,"__wrapped__"),othIsWrapped=othIsObj&&hasOwnProperty.call(other,"__wrapped__");if(objIsWrapped||othIsWrapped){return equalFunc(objIsWrapped?object.value():object,othIsWrapped?other.value():other,customizer,isLoose,stackA,stackB)}}if(!isSameTag){return false}stackA||(stackA=[]);stackB||(stackB=[]);var length=stackA.length;while(length--){if(stackA[length]==object){return stackB[length]==other}}stackA.push(object);stackB.push(other);var result=(objIsArr?equalArrays:equalObjects)(object,other,equalFunc,customizer,isLoose,stackA,stackB);stackA.pop();stackB.pop();return result}function baseIsMatch(object,matchData,customizer){var index=matchData.length,length=index,noCustomizer=!customizer;if(object==null){return!length}object=toObject(object);while(index--){var data=matchData[index];if(noCustomizer&&data[2]?data[1]!==object[data[0]]:!(data[0]in object)){return false}}while(++index<length){data=matchData[index];var key=data[0],objValue=object[key],srcValue=data[1];if(noCustomizer&&data[2]){if(objValue===undefined&&!(key in object)){return false}}else{var result=customizer?customizer(objValue,srcValue,key):undefined;if(!(result===undefined?baseIsEqual(srcValue,objValue,customizer,true):result)){return false}}}return true}function baseMap(collection,iteratee){var index=-1,result=isArrayLike(collection)?Array(collection.length):[];baseEach(collection,function(value,key,collection){result[++index]=iteratee(value,key,collection)});return result}function baseMatches(source){var matchData=getMatchData(source);if(matchData.length==1&&matchData[0][2]){var key=matchData[0][0],value=matchData[0][1];return function(object){if(object==null){return false}return object[key]===value&&(value!==undefined||key in toObject(object))}}return function(object){return baseIsMatch(object,matchData)}}function baseMatchesProperty(path,srcValue){var isArr=isArray(path),isCommon=isKey(path)&&isStrictComparable(srcValue),pathKey=path+"";path=toPath(path);return function(object){if(object==null){return false}var key=pathKey;object=toObject(object);if((isArr||!isCommon)&&!(key in object)){object=path.length==1?object:baseGet(object,baseSlice(path,0,-1));if(object==null){return false}key=last(path);object=toObject(object)}return object[key]===srcValue?srcValue!==undefined||key in object:baseIsEqual(srcValue,object[key],undefined,true)}}function baseMerge(object,source,customizer,stackA,stackB){if(!isObject(object)){return object}var isSrcArr=isArrayLike(source)&&(isArray(source)||isTypedArray(source)),props=isSrcArr?undefined:keys(source);arrayEach(props||source,function(srcValue,key){if(props){key=srcValue;srcValue=source[key]}if(isObjectLike(srcValue)){stackA||(stackA=[]);stackB||(stackB=[]);baseMergeDeep(object,source,key,baseMerge,customizer,stackA,stackB)}else{var value=object[key],result=customizer?customizer(value,srcValue,key,object,source):undefined,isCommon=result===undefined;if(isCommon){result=srcValue}if((result!==undefined||isSrcArr&&!(key in object))&&(isCommon||(result===result?result!==value:value===value))){object[key]=result}}});return object}function baseMergeDeep(object,source,key,mergeFunc,customizer,stackA,stackB){var length=stackA.length,srcValue=source[key];while(length--){if(stackA[length]==srcValue){object[key]=stackB[length];return}}var value=object[key],result=customizer?customizer(value,srcValue,key,object,source):undefined,isCommon=result===undefined;if(isCommon){result=srcValue;if(isArrayLike(srcValue)&&(isArray(srcValue)||isTypedArray(srcValue))){result=isArray(value)?value:isArrayLike(value)?arrayCopy(value):[]}else if(isPlainObject(srcValue)||isArguments(srcValue)){result=isArguments(value)?toPlainObject(value):isPlainObject(value)?value:{}}else{isCommon=false}}stackA.push(srcValue);stackB.push(result);if(isCommon){object[key]=mergeFunc(result,srcValue,customizer,stackA,stackB)}else if(result===result?result!==value:value===value){object[key]=result}}function baseProperty(key){return function(object){return object==null?undefined:object[key]}}function basePropertyDeep(path){var pathKey=path+"";path=toPath(path);return function(object){return baseGet(object,path,pathKey)}}function basePullAt(array,indexes){var length=array?indexes.length:0;while(length--){var index=indexes[length];if(index!=previous&&isIndex(index)){var previous=index;splice.call(array,index,1)}}return array}function baseRandom(min,max){return min+nativeFloor(nativeRandom()*(max-min+1))}function baseReduce(collection,iteratee,accumulator,initFromCollection,eachFunc){eachFunc(collection,function(value,index,collection){accumulator=initFromCollection?(initFromCollection=false,value):iteratee(accumulator,value,index,collection)});return accumulator}var baseSetData=!metaMap?identity:function(func,data){metaMap.set(func,data);return func};function baseSlice(array,start,end){var index=-1,length=array.length;start=start==null?0:+start||0;if(start<0){start=-start>length?0:length+start}end=end===undefined||end>length?length:+end||0;if(end<0){end+=length}length=start>end?0:end-start>>>0;start>>>=0;var result=Array(length);while(++index<length){result[index]=array[index+start]}return result}function baseSome(collection,predicate){var result;baseEach(collection,function(value,index,collection){result=predicate(value,index,collection);return!result});return!!result}function baseSortBy(array,comparer){var length=array.length;array.sort(comparer);while(length--){array[length]=array[length].value}return array}function baseSortByOrder(collection,iteratees,orders){var callback=getCallback(),index=-1;iteratees=arrayMap(iteratees,function(iteratee){return callback(iteratee)});var result=baseMap(collection,function(value){var criteria=arrayMap(iteratees,function(iteratee){return iteratee(value)});return{criteria:criteria,index:++index,value:value}});return baseSortBy(result,function(object,other){return compareMultiple(object,other,orders)})}function baseSum(collection,iteratee){var result=0;baseEach(collection,function(value,index,collection){result+=+iteratee(value,index,collection)||0});return result}function baseUniq(array,iteratee){var index=-1,indexOf=getIndexOf(),length=array.length,isCommon=indexOf==baseIndexOf,isLarge=isCommon&&length>=LARGE_ARRAY_SIZE,seen=isLarge?createCache():null,result=[];if(seen){indexOf=cacheIndexOf;isCommon=false}else{isLarge=false;seen=iteratee?[]:result}outer:while(++index<length){var value=array[index],computed=iteratee?iteratee(value,index,array):value;if(isCommon&&value===value){var seenIndex=seen.length;while(seenIndex--){if(seen[seenIndex]===computed){continue outer}}if(iteratee){seen.push(computed)}result.push(value)}else if(indexOf(seen,computed,0)<0){if(iteratee||isLarge){seen.push(computed)}result.push(value)}}return result}function baseValues(object,props){var index=-1,length=props.length,result=Array(length);while(++index<length){result[index]=object[props[index]]}return result}function baseWhile(array,predicate,isDrop,fromRight){var length=array.length,index=fromRight?length:-1;while((fromRight?index--:++index<length)&&predicate(array[index],index,array)){}return isDrop?baseSlice(array,fromRight?0:index,fromRight?index+1:length):baseSlice(array,fromRight?index+1:0,fromRight?length:index)}function baseWrapperValue(value,actions){var result=value;if(result instanceof LazyWrapper){result=result.value()}var index=-1,length=actions.length;while(++index<length){var action=actions[index];result=action.func.apply(action.thisArg,arrayPush([result],action.args))}return result}function binaryIndex(array,value,retHighest){var low=0,high=array?array.length:low;if(typeof value=="number"&&value===value&&high<=HALF_MAX_ARRAY_LENGTH){while(low<high){var mid=low+high>>>1,computed=array[mid];if((retHighest?computed<=value:computed<value)&&computed!==null){low=mid+1}else{high=mid}}return high}return binaryIndexBy(array,value,identity,retHighest)}function binaryIndexBy(array,value,iteratee,retHighest){value=iteratee(value);var low=0,high=array?array.length:0,valIsNaN=value!==value,valIsNull=value===null,valIsUndef=value===undefined;while(low<high){var mid=nativeFloor((low+high)/2),computed=iteratee(array[mid]),isDef=computed!==undefined,isReflexive=computed===computed;if(valIsNaN){var setLow=isReflexive||retHighest}else if(valIsNull){setLow=isReflexive&&isDef&&(retHighest||computed!=null)}else if(valIsUndef){setLow=isReflexive&&(retHighest||isDef)}else if(computed==null){setLow=false}else{setLow=retHighest?computed<=value:computed<value}if(setLow){low=mid+1}else{high=mid}}return nativeMin(high,MAX_ARRAY_INDEX)}function bindCallback(func,thisArg,argCount){if(typeof func!="function"){return identity}if(thisArg===undefined){return func}switch(argCount){case 1:return function(value){return func.call(thisArg,value)};case 3:return function(value,index,collection){return func.call(thisArg,value,index,collection)};case 4:return function(accumulator,value,index,collection){return func.call(thisArg,accumulator,value,index,collection)};case 5:return function(value,other,key,object,source){return func.call(thisArg,value,other,key,object,source)}}return function(){return func.apply(thisArg,arguments)}}function bufferClone(buffer){var result=new ArrayBuffer(buffer.byteLength),view=new Uint8Array(result);view.set(new Uint8Array(buffer));return result}function composeArgs(args,partials,holders){var holdersLength=holders.length,argsIndex=-1,argsLength=nativeMax(args.length-holdersLength,0),leftIndex=-1,leftLength=partials.length,result=Array(leftLength+argsLength);while(++leftIndex<leftLength){result[leftIndex]=partials[leftIndex]}while(++argsIndex<holdersLength){result[holders[argsIndex]]=args[argsIndex]}while(argsLength--){result[leftIndex++]=args[argsIndex++]}return result}function composeArgsRight(args,partials,holders){var holdersIndex=-1,holdersLength=holders.length,argsIndex=-1,argsLength=nativeMax(args.length-holdersLength,0),rightIndex=-1,rightLength=partials.length,result=Array(argsLength+rightLength);while(++argsIndex<argsLength){result[argsIndex]=args[argsIndex]}var offset=argsIndex;while(++rightIndex<rightLength){result[offset+rightIndex]=partials[rightIndex]}while(++holdersIndex<holdersLength){result[offset+holders[holdersIndex]]=args[argsIndex++]}return result}function createAggregator(setter,initializer){return function(collection,iteratee,thisArg){var result=initializer?initializer():{};iteratee=getCallback(iteratee,thisArg,3);if(isArray(collection)){var index=-1,length=collection.length;while(++index<length){var value=collection[index];setter(result,value,iteratee(value,index,collection),collection)}}else{baseEach(collection,function(value,key,collection){setter(result,value,iteratee(value,key,collection),collection)})}return result}}function createAssigner(assigner){return restParam(function(object,sources){var index=-1,length=object==null?0:sources.length,customizer=length>2?sources[length-2]:undefined,guard=length>2?sources[2]:undefined,thisArg=length>1?sources[length-1]:undefined;if(typeof customizer=="function"){customizer=bindCallback(customizer,thisArg,5);length-=2}else{customizer=typeof thisArg=="function"?thisArg:undefined;length-=customizer?1:0}if(guard&&isIterateeCall(sources[0],sources[1],guard)){customizer=length<3?undefined:customizer;length=1}while(++index<length){var source=sources[index];if(source){assigner(object,source,customizer)}}return object})}function createBaseEach(eachFunc,fromRight){return function(collection,iteratee){var length=collection?getLength(collection):0;if(!isLength(length)){return eachFunc(collection,iteratee)}var index=fromRight?length:-1,iterable=toObject(collection);while(fromRight?index--:++index<length){if(iteratee(iterable[index],index,iterable)===false){break}}return collection}}function createBaseFor(fromRight){return function(object,iteratee,keysFunc){var iterable=toObject(object),props=keysFunc(object),length=props.length,index=fromRight?length:-1;while(fromRight?index--:++index<length){var key=props[index];if(iteratee(iterable[key],key,iterable)===false){break}}return object}}function createBindWrapper(func,thisArg){var Ctor=createCtorWrapper(func);function wrapper(){var fn=this&&this!==root&&this instanceof wrapper?Ctor:func;return fn.apply(thisArg,arguments)}return wrapper}function createCache(values){return nativeCreate&&Set?new SetCache(values):null}function createCompounder(callback){return function(string){var index=-1,array=words(deburr(string)),length=array.length,result="";while(++index<length){result=callback(result,array[index],index)}return result}}function createCtorWrapper(Ctor){return function(){var args=arguments;switch(args.length){case 0:return new Ctor;case 1:return new Ctor(args[0]);case 2:return new Ctor(args[0],args[1]);case 3:return new Ctor(args[0],args[1],args[2]);case 4:return new Ctor(args[0],args[1],args[2],args[3]);case 5:return new Ctor(args[0],args[1],args[2],args[3],args[4]);case 6:return new Ctor(args[0],args[1],args[2],args[3],args[4],args[5]);case 7:return new Ctor(args[0],args[1],args[2],args[3],args[4],args[5],args[6])}var thisBinding=baseCreate(Ctor.prototype),result=Ctor.apply(thisBinding,args);return isObject(result)?result:thisBinding}}function createCurry(flag){function curryFunc(func,arity,guard){if(guard&&isIterateeCall(func,arity,guard)){arity=undefined}var result=createWrapper(func,flag,undefined,undefined,undefined,undefined,undefined,arity);result.placeholder=curryFunc.placeholder;return result}return curryFunc}function createDefaults(assigner,customizer){return restParam(function(args){var object=args[0];if(object==null){return object}args.push(customizer);return assigner.apply(undefined,args)})}function createExtremum(comparator,exValue){return function(collection,iteratee,thisArg){if(thisArg&&isIterateeCall(collection,iteratee,thisArg)){iteratee=undefined}iteratee=getCallback(iteratee,thisArg,3);if(iteratee.length==1){collection=isArray(collection)?collection:toIterable(collection);var result=arrayExtremum(collection,iteratee,comparator,exValue);if(!(collection.length&&result===exValue)){return result}}return baseExtremum(collection,iteratee,comparator,exValue)}}function createFind(eachFunc,fromRight){return function(collection,predicate,thisArg){predicate=getCallback(predicate,thisArg,3);if(isArray(collection)){var index=baseFindIndex(collection,predicate,fromRight);return index>-1?collection[index]:undefined}return baseFind(collection,predicate,eachFunc)}}function createFindIndex(fromRight){return function(array,predicate,thisArg){if(!(array&&array.length)){return-1}predicate=getCallback(predicate,thisArg,3);return baseFindIndex(array,predicate,fromRight)}}function createFindKey(objectFunc){return function(object,predicate,thisArg){predicate=getCallback(predicate,thisArg,3);return baseFind(object,predicate,objectFunc,true)}}function createFlow(fromRight){return function(){var wrapper,length=arguments.length,index=fromRight?length:-1,leftIndex=0,funcs=Array(length);while(fromRight?index--:++index<length){var func=funcs[leftIndex++]=arguments[index];if(typeof func!="function"){throw new TypeError(FUNC_ERROR_TEXT)}if(!wrapper&&LodashWrapper.prototype.thru&&getFuncName(func)=="wrapper"){wrapper=new LodashWrapper([],true)}}index=wrapper?-1:length;while(++index<length){func=funcs[index];var funcName=getFuncName(func),data=funcName=="wrapper"?getData(func):undefined;if(data&&isLaziable(data[0])&&data[1]==(ARY_FLAG|CURRY_FLAG|PARTIAL_FLAG|REARG_FLAG)&&!data[4].length&&data[9]==1){wrapper=wrapper[getFuncName(data[0])].apply(wrapper,data[3])}else{wrapper=func.length==1&&isLaziable(func)?wrapper[funcName]():wrapper.thru(func)}}return function(){var args=arguments,value=args[0];if(wrapper&&args.length==1&&isArray(value)&&value.length>=LARGE_ARRAY_SIZE){return wrapper.plant(value).value()}var index=0,result=length?funcs[index].apply(this,args):value;while(++index<length){result=funcs[index].call(this,result)}return result}}}function createForEach(arrayFunc,eachFunc){return function(collection,iteratee,thisArg){return typeof iteratee=="function"&&thisArg===undefined&&isArray(collection)?arrayFunc(collection,iteratee):eachFunc(collection,bindCallback(iteratee,thisArg,3))}}function createForIn(objectFunc){return function(object,iteratee,thisArg){if(typeof iteratee!="function"||thisArg!==undefined){iteratee=bindCallback(iteratee,thisArg,3)}return objectFunc(object,iteratee,keysIn)}}function createForOwn(objectFunc){return function(object,iteratee,thisArg){if(typeof iteratee!="function"||thisArg!==undefined){iteratee=bindCallback(iteratee,thisArg,3)}return objectFunc(object,iteratee)}}function createObjectMapper(isMapKeys){return function(object,iteratee,thisArg){var result={};iteratee=getCallback(iteratee,thisArg,3);baseForOwn(object,function(value,key,object){var mapped=iteratee(value,key,object);key=isMapKeys?mapped:key;value=isMapKeys?value:mapped;result[key]=value});return result}}function createPadDir(fromRight){return function(string,length,chars){string=baseToString(string);return(fromRight?string:"")+createPadding(string,length,chars)+(fromRight?"":string)}}function createPartial(flag){var partialFunc=restParam(function(func,partials){var holders=replaceHolders(partials,partialFunc.placeholder);return createWrapper(func,flag,undefined,partials,holders)});return partialFunc}function createReduce(arrayFunc,eachFunc){return function(collection,iteratee,accumulator,thisArg){var initFromArray=arguments.length<3;return typeof iteratee=="function"&&thisArg===undefined&&isArray(collection)?arrayFunc(collection,iteratee,accumulator,initFromArray):baseReduce(collection,getCallback(iteratee,thisArg,4),accumulator,initFromArray,eachFunc)}}function createHybridWrapper(func,bitmask,thisArg,partials,holders,partialsRight,holdersRight,argPos,ary,arity){var isAry=bitmask&ARY_FLAG,isBind=bitmask&BIND_FLAG,isBindKey=bitmask&BIND_KEY_FLAG,isCurry=bitmask&CURRY_FLAG,isCurryBound=bitmask&CURRY_BOUND_FLAG,isCurryRight=bitmask&CURRY_RIGHT_FLAG,Ctor=isBindKey?undefined:createCtorWrapper(func);function wrapper(){var length=arguments.length,index=length,args=Array(length);while(index--){args[index]=arguments[index]}if(partials){args=composeArgs(args,partials,holders)}if(partialsRight){args=composeArgsRight(args,partialsRight,holdersRight)}if(isCurry||isCurryRight){var placeholder=wrapper.placeholder,argsHolders=replaceHolders(args,placeholder);length-=argsHolders.length;if(length<arity){var newArgPos=argPos?arrayCopy(argPos):undefined,newArity=nativeMax(arity-length,0),newsHolders=isCurry?argsHolders:undefined,newHoldersRight=isCurry?undefined:argsHolders,newPartials=isCurry?args:undefined,newPartialsRight=isCurry?undefined:args;bitmask|=isCurry?PARTIAL_FLAG:PARTIAL_RIGHT_FLAG;bitmask&=~(isCurry?PARTIAL_RIGHT_FLAG:PARTIAL_FLAG);if(!isCurryBound){bitmask&=~(BIND_FLAG|BIND_KEY_FLAG)}var newData=[func,bitmask,thisArg,newPartials,newsHolders,newPartialsRight,newHoldersRight,newArgPos,ary,newArity],result=createHybridWrapper.apply(undefined,newData);if(isLaziable(func)){setData(result,newData)}result.placeholder=placeholder;return result}}var thisBinding=isBind?thisArg:this,fn=isBindKey?thisBinding[func]:func;if(argPos){args=reorder(args,argPos)}if(isAry&&ary<args.length){args.length=ary}if(this&&this!==root&&this instanceof wrapper){fn=Ctor||createCtorWrapper(func)}return fn.apply(thisBinding,args)}return wrapper}function createPadding(string,length,chars){var strLength=string.length;length=+length;if(strLength>=length||!nativeIsFinite(length)){return""}var padLength=length-strLength;chars=chars==null?" ":chars+"";return repeat(chars,nativeCeil(padLength/chars.length)).slice(0,padLength)}function createPartialWrapper(func,bitmask,thisArg,partials){var isBind=bitmask&BIND_FLAG,Ctor=createCtorWrapper(func);function wrapper(){var argsIndex=-1,argsLength=arguments.length,leftIndex=-1,leftLength=partials.length,args=Array(leftLength+argsLength);while(++leftIndex<leftLength){args[leftIndex]=partials[leftIndex]}while(argsLength--){args[leftIndex++]=arguments[++argsIndex]}var fn=this&&this!==root&&this instanceof wrapper?Ctor:func;return fn.apply(isBind?thisArg:this,args)}return wrapper}function createRound(methodName){var func=Math[methodName];return function(number,precision){precision=precision===undefined?0:+precision||0;if(precision){precision=pow(10,precision);return func(number*precision)/precision}return func(number)}}function createSortedIndex(retHighest){return function(array,value,iteratee,thisArg){var callback=getCallback(iteratee);return iteratee==null&&callback===baseCallback?binaryIndex(array,value,retHighest):binaryIndexBy(array,value,callback(iteratee,thisArg,1),retHighest)}}function createWrapper(func,bitmask,thisArg,partials,holders,argPos,ary,arity){var isBindKey=bitmask&BIND_KEY_FLAG;if(!isBindKey&&typeof func!="function"){throw new TypeError(FUNC_ERROR_TEXT)}var length=partials?partials.length:0;if(!length){bitmask&=~(PARTIAL_FLAG|PARTIAL_RIGHT_FLAG);partials=holders=undefined}length-=holders?holders.length:0;if(bitmask&PARTIAL_RIGHT_FLAG){var partialsRight=partials,holdersRight=holders;partials=holders=undefined}var data=isBindKey?undefined:getData(func),newData=[func,bitmask,thisArg,partials,holders,partialsRight,holdersRight,argPos,ary,arity];if(data){mergeData(newData,data);bitmask=newData[1];arity=newData[9]}newData[9]=arity==null?isBindKey?0:func.length:nativeMax(arity-length,0)||0;if(bitmask==BIND_FLAG){var result=createBindWrapper(newData[0],newData[2])}else if((bitmask==PARTIAL_FLAG||bitmask==(BIND_FLAG|PARTIAL_FLAG))&&!newData[4].length){result=createPartialWrapper.apply(undefined,newData)}else{result=createHybridWrapper.apply(undefined,newData)}var setter=data?baseSetData:setData;return setter(result,newData)}function equalArrays(array,other,equalFunc,customizer,isLoose,stackA,stackB){var index=-1,arrLength=array.length,othLength=other.length;if(arrLength!=othLength&&!(isLoose&&othLength>arrLength)){return false}while(++index<arrLength){var arrValue=array[index],othValue=other[index],result=customizer?customizer(isLoose?othValue:arrValue,isLoose?arrValue:othValue,index):undefined;if(result!==undefined){if(result){continue}return false}if(isLoose){if(!arraySome(other,function(othValue){return arrValue===othValue||equalFunc(arrValue,othValue,customizer,isLoose,stackA,stackB)})){return false}}else if(!(arrValue===othValue||equalFunc(arrValue,othValue,customizer,isLoose,stackA,stackB))){return false}}return true}function equalByTag(object,other,tag){switch(tag){case boolTag:case dateTag:return+object==+other;case errorTag:return object.name==other.name&&object.message==other.message;case numberTag:return object!=+object?other!=+other:object==+other;case regexpTag:case stringTag:return object==other+""}return false}function equalObjects(object,other,equalFunc,customizer,isLoose,stackA,stackB){var objProps=keys(object),objLength=objProps.length,othProps=keys(other),othLength=othProps.length;if(objLength!=othLength&&!isLoose){return false}var index=objLength;while(index--){var key=objProps[index];if(!(isLoose?key in other:hasOwnProperty.call(other,key))){return false}}var skipCtor=isLoose;while(++index<objLength){key=objProps[index];var objValue=object[key],othValue=other[key],result=customizer?customizer(isLoose?othValue:objValue,isLoose?objValue:othValue,key):undefined;if(!(result===undefined?equalFunc(objValue,othValue,customizer,isLoose,stackA,stackB):result)){return false}skipCtor||(skipCtor=key=="constructor")}if(!skipCtor){var objCtor=object.constructor,othCtor=other.constructor;if(objCtor!=othCtor&&("constructor"in object&&"constructor"in other)&&!(typeof objCtor=="function"&&objCtor instanceof objCtor&&typeof othCtor=="function"&&othCtor instanceof othCtor)){return false}}return true}function getCallback(func,thisArg,argCount){var result=lodash.callback||callback;result=result===callback?baseCallback:result;return argCount?result(func,thisArg,argCount):result}var getData=!metaMap?noop:function(func){return metaMap.get(func)};function getFuncName(func){var result=func.name,array=realNames[result],length=array?array.length:0;while(length--){var data=array[length],otherFunc=data.func;if(otherFunc==null||otherFunc==func){return data.name}}return result}function getIndexOf(collection,target,fromIndex){var result=lodash.indexOf||indexOf;result=result===indexOf?baseIndexOf:result;return collection?result(collection,target,fromIndex):result}var getLength=baseProperty("length");function getMatchData(object){var result=pairs(object),length=result.length;while(length--){result[length][2]=isStrictComparable(result[length][1])}return result}function getNative(object,key){var value=object==null?undefined:object[key];return isNative(value)?value:undefined}function getView(start,end,transforms){var index=-1,length=transforms.length;while(++index<length){var data=transforms[index],size=data.size;switch(data.type){case"drop":start+=size;break;case"dropRight":end-=size;break;case"take":end=nativeMin(end,start+size);break;case"takeRight":start=nativeMax(start,end-size);break}}return{start:start,end:end}}function initCloneArray(array){var length=array.length,result=new array.constructor(length);if(length&&typeof array[0]=="string"&&hasOwnProperty.call(array,"index")){result.index=array.index;result.input=array.input}return result}function initCloneObject(object){var Ctor=object.constructor;if(!(typeof Ctor=="function"&&Ctor instanceof Ctor)){Ctor=Object}return new Ctor}function initCloneByTag(object,tag,isDeep){var Ctor=object.constructor;switch(tag){case arrayBufferTag:return bufferClone(object);case boolTag:case dateTag:return new Ctor(+object);case float32Tag:case float64Tag:case int8Tag:case int16Tag:case int32Tag:case uint8Tag:case uint8ClampedTag:case uint16Tag:case uint32Tag:var buffer=object.buffer;return new Ctor(isDeep?bufferClone(buffer):buffer,object.byteOffset,object.length);case numberTag:case stringTag:return new Ctor(object);case regexpTag:var result=new Ctor(object.source,reFlags.exec(object));result.lastIndex=object.lastIndex}return result}function invokePath(object,path,args){if(object!=null&&!isKey(path,object)){path=toPath(path);object=path.length==1?object:baseGet(object,baseSlice(path,0,-1));path=last(path)}var func=object==null?object:object[path];return func==null?undefined:func.apply(object,args)}function isArrayLike(value){return value!=null&&isLength(getLength(value))}function isIndex(value,length){value=typeof value=="number"||reIsUint.test(value)?+value:-1;length=length==null?MAX_SAFE_INTEGER:length;return value>-1&&value%1==0&&value<length}function isIterateeCall(value,index,object){if(!isObject(object)){return false}var type=typeof index;if(type=="number"?isArrayLike(object)&&isIndex(index,object.length):type=="string"&&index in object){var other=object[index];return value===value?value===other:other!==other}return false}function isKey(value,object){var type=typeof value;if(type=="string"&&reIsPlainProp.test(value)||type=="number"){return true}if(isArray(value)){return false}var result=!reIsDeepProp.test(value);return result||object!=null&&value in toObject(object)}function isLaziable(func){var funcName=getFuncName(func);if(!(funcName in LazyWrapper.prototype)){return false}var other=lodash[funcName];if(func===other){return true}var data=getData(other);return!!data&&func===data[0]}function isLength(value){return typeof value=="number"&&value>-1&&value%1==0&&value<=MAX_SAFE_INTEGER}function isStrictComparable(value){return value===value&&!isObject(value)}function mergeData(data,source){var bitmask=data[1],srcBitmask=source[1],newBitmask=bitmask|srcBitmask,isCommon=newBitmask<ARY_FLAG;var isCombo=srcBitmask==ARY_FLAG&&bitmask==CURRY_FLAG||srcBitmask==ARY_FLAG&&bitmask==REARG_FLAG&&data[7].length<=source[8]||srcBitmask==(ARY_FLAG|REARG_FLAG)&&bitmask==CURRY_FLAG;if(!(isCommon||isCombo)){return data}if(srcBitmask&BIND_FLAG){data[2]=source[2];newBitmask|=bitmask&BIND_FLAG?0:CURRY_BOUND_FLAG}var value=source[3];if(value){var partials=data[3];data[3]=partials?composeArgs(partials,value,source[4]):arrayCopy(value);data[4]=partials?replaceHolders(data[3],PLACEHOLDER):arrayCopy(source[4])}value=source[5];if(value){partials=data[5];data[5]=partials?composeArgsRight(partials,value,source[6]):arrayCopy(value);data[6]=partials?replaceHolders(data[5],PLACEHOLDER):arrayCopy(source[6])}value=source[7];if(value){data[7]=arrayCopy(value)}if(srcBitmask&ARY_FLAG){data[8]=data[8]==null?source[8]:nativeMin(data[8],source[8])}if(data[9]==null){data[9]=source[9]}data[0]=source[0];data[1]=newBitmask;return data}function mergeDefaults(objectValue,sourceValue){return objectValue===undefined?sourceValue:merge(objectValue,sourceValue,mergeDefaults)}function pickByArray(object,props){object=toObject(object);var index=-1,length=props.length,result={};while(++index<length){var key=props[index];if(key in object){result[key]=object[key]}}return result}function pickByCallback(object,predicate){var result={};baseForIn(object,function(value,key,object){if(predicate(value,key,object)){result[key]=value}});return result}function reorder(array,indexes){var arrLength=array.length,length=nativeMin(indexes.length,arrLength),oldArray=arrayCopy(array);while(length--){var index=indexes[length];array[length]=isIndex(index,arrLength)?oldArray[index]:undefined}return array}var setData=function(){var count=0,lastCalled=0;return function(key,value){var stamp=now(),remaining=HOT_SPAN-(stamp-lastCalled);lastCalled=stamp;if(remaining>0){if(++count>=HOT_COUNT){return key}}else{count=0}return baseSetData(key,value)}}();function shimKeys(object){var props=keysIn(object),propsLength=props.length,length=propsLength&&object.length;var allowIndexes=!!length&&isLength(length)&&(isArray(object)||isArguments(object));var index=-1,result=[];while(++index<propsLength){var key=props[index];if(allowIndexes&&isIndex(key,length)||hasOwnProperty.call(object,key)){result.push(key)}}return result}function toIterable(value){if(value==null){return[]}if(!isArrayLike(value)){return values(value)}return isObject(value)?value:Object(value)}function toObject(value){return isObject(value)?value:Object(value)}function toPath(value){if(isArray(value)){return value}var result=[];baseToString(value).replace(rePropName,function(match,number,quote,string){result.push(quote?string.replace(reEscapeChar,"$1"):number||match)});return result}function wrapperClone(wrapper){return wrapper instanceof LazyWrapper?wrapper.clone():new LodashWrapper(wrapper.__wrapped__,wrapper.__chain__,arrayCopy(wrapper.__actions__))}function chunk(array,size,guard){if(guard?isIterateeCall(array,size,guard):size==null){size=1}else{size=nativeMax(nativeFloor(size)||1,1)}var index=0,length=array?array.length:0,resIndex=-1,result=Array(nativeCeil(length/size));while(index<length){result[++resIndex]=baseSlice(array,index,index+=size)}return result}function compact(array){var index=-1,length=array?array.length:0,resIndex=-1,result=[];while(++index<length){var value=array[index];if(value){result[++resIndex]=value}}return result}var difference=restParam(function(array,values){return isObjectLike(array)&&isArrayLike(array)?baseDifference(array,baseFlatten(values,false,true)):[]});function drop(array,n,guard){var length=array?array.length:0;if(!length){return[]}if(guard?isIterateeCall(array,n,guard):n==null){n=1}return baseSlice(array,n<0?0:n)}function dropRight(array,n,guard){var length=array?array.length:0;if(!length){return[]}if(guard?isIterateeCall(array,n,guard):n==null){n=1}n=length-(+n||0);return baseSlice(array,0,n<0?0:n)}function dropRightWhile(array,predicate,thisArg){return array&&array.length?baseWhile(array,getCallback(predicate,thisArg,3),true,true):[]}function dropWhile(array,predicate,thisArg){return array&&array.length?baseWhile(array,getCallback(predicate,thisArg,3),true):[]}function fill(array,value,start,end){var length=array?array.length:0;if(!length){return[]}if(start&&typeof start!="number"&&isIterateeCall(array,value,start)){start=0;end=length}return baseFill(array,value,start,end)}var findIndex=createFindIndex();var findLastIndex=createFindIndex(true);function first(array){return array?array[0]:undefined}function flatten(array,isDeep,guard){var length=array?array.length:0;if(guard&&isIterateeCall(array,isDeep,guard)){isDeep=false}return length?baseFlatten(array,isDeep):[]}function flattenDeep(array){var length=array?array.length:0;return length?baseFlatten(array,true):[]}function indexOf(array,value,fromIndex){var length=array?array.length:0;if(!length){return-1}if(typeof fromIndex=="number"){fromIndex=fromIndex<0?nativeMax(length+fromIndex,0):fromIndex}else if(fromIndex){var index=binaryIndex(array,value);if(index<length&&(value===value?value===array[index]:array[index]!==array[index])){return index}return-1}return baseIndexOf(array,value,fromIndex||0)}function initial(array){return dropRight(array,1)}var intersection=restParam(function(arrays){var othLength=arrays.length,othIndex=othLength,caches=Array(length),indexOf=getIndexOf(),isCommon=indexOf==baseIndexOf,result=[];while(othIndex--){var value=arrays[othIndex]=isArrayLike(value=arrays[othIndex])?value:[];caches[othIndex]=isCommon&&value.length>=120?createCache(othIndex&&value):null}var array=arrays[0],index=-1,length=array?array.length:0,seen=caches[0];outer:while(++index<length){value=array[index];if((seen?cacheIndexOf(seen,value):indexOf(result,value,0))<0){var othIndex=othLength;while(--othIndex){var cache=caches[othIndex];if((cache?cacheIndexOf(cache,value):indexOf(arrays[othIndex],value,0))<0){continue outer}}if(seen){seen.push(value)}result.push(value)}}return result});function last(array){var length=array?array.length:0;return length?array[length-1]:undefined}function lastIndexOf(array,value,fromIndex){var length=array?array.length:0;if(!length){return-1}var index=length;if(typeof fromIndex=="number"){index=(fromIndex<0?nativeMax(length+fromIndex,0):nativeMin(fromIndex||0,length-1))+1}else if(fromIndex){index=binaryIndex(array,value,true)-1;var other=array[index];if(value===value?value===other:other!==other){return index}return-1}if(value!==value){return indexOfNaN(array,index,true)}while(index--){if(array[index]===value){return index}}return-1}function pull(){var args=arguments,array=args[0];if(!(array&&array.length)){return array}var index=0,indexOf=getIndexOf(),length=args.length;while(++index<length){var fromIndex=0,value=args[index];while((fromIndex=indexOf(array,value,fromIndex))>-1){splice.call(array,fromIndex,1)}}return array}var pullAt=restParam(function(array,indexes){indexes=baseFlatten(indexes);var result=baseAt(array,indexes);basePullAt(array,indexes.sort(baseCompareAscending));return result});function remove(array,predicate,thisArg){var result=[];if(!(array&&array.length)){return result}var index=-1,indexes=[],length=array.length;predicate=getCallback(predicate,thisArg,3);while(++index<length){var value=array[index];if(predicate(value,index,array)){result.push(value);indexes.push(index)}}basePullAt(array,indexes);return result}function rest(array){return drop(array,1)}function slice(array,start,end){var length=array?array.length:0;if(!length){return[]}if(end&&typeof end!="number"&&isIterateeCall(array,start,end)){start=0;end=length}return baseSlice(array,start,end)}var sortedIndex=createSortedIndex();var sortedLastIndex=createSortedIndex(true);function take(array,n,guard){var length=array?array.length:0;if(!length){return[]}if(guard?isIterateeCall(array,n,guard):n==null){n=1}return baseSlice(array,0,n<0?0:n)}function takeRight(array,n,guard){var length=array?array.length:0;if(!length){return[]}if(guard?isIterateeCall(array,n,guard):n==null){n=1}n=length-(+n||0);return baseSlice(array,n<0?0:n)}function takeRightWhile(array,predicate,thisArg){return array&&array.length?baseWhile(array,getCallback(predicate,thisArg,3),false,true):[]}function takeWhile(array,predicate,thisArg){return array&&array.length?baseWhile(array,getCallback(predicate,thisArg,3)):[]}var union=restParam(function(arrays){return baseUniq(baseFlatten(arrays,false,true))});function uniq(array,isSorted,iteratee,thisArg){var length=array?array.length:0;if(!length){return[]}if(isSorted!=null&&typeof isSorted!="boolean"){thisArg=iteratee;iteratee=isIterateeCall(array,isSorted,thisArg)?undefined:isSorted;isSorted=false}var callback=getCallback();if(!(iteratee==null&&callback===baseCallback)){iteratee=callback(iteratee,thisArg,3)}return isSorted&&getIndexOf()==baseIndexOf?sortedUniq(array,iteratee):baseUniq(array,iteratee)}function unzip(array){if(!(array&&array.length)){return[]}var index=-1,length=0;array=arrayFilter(array,function(group){if(isArrayLike(group)){length=nativeMax(group.length,length);return true}});var result=Array(length);while(++index<length){result[index]=arrayMap(array,baseProperty(index))}return result}function unzipWith(array,iteratee,thisArg){var length=array?array.length:0;if(!length){return[]}var result=unzip(array);if(iteratee==null){return result}iteratee=bindCallback(iteratee,thisArg,4);return arrayMap(result,function(group){return arrayReduce(group,iteratee,undefined,true)})}var without=restParam(function(array,values){return isArrayLike(array)?baseDifference(array,values):[]});function xor(){var index=-1,length=arguments.length;while(++index<length){var array=arguments[index];if(isArrayLike(array)){var result=result?arrayPush(baseDifference(result,array),baseDifference(array,result)):array}}return result?baseUniq(result):[]}var zip=restParam(unzip);function zipObject(props,values){var index=-1,length=props?props.length:0,result={};if(length&&!values&&!isArray(props[0])){values=[]}while(++index<length){var key=props[index];if(values){result[key]=values[index]}else if(key){result[key[0]]=key[1]}}return result}var zipWith=restParam(function(arrays){var length=arrays.length,iteratee=length>2?arrays[length-2]:undefined,thisArg=length>1?arrays[length-1]:undefined;if(length>2&&typeof iteratee=="function"){length-=2}else{iteratee=length>1&&typeof thisArg=="function"?(--length,thisArg):undefined;thisArg=undefined}arrays.length=length;return unzipWith(arrays,iteratee,thisArg)});function chain(value){var result=lodash(value);result.__chain__=true;return result}function tap(value,interceptor,thisArg){interceptor.call(thisArg,value);return value}function thru(value,interceptor,thisArg){return interceptor.call(thisArg,value)}function wrapperChain(){return chain(this)}function wrapperCommit(){return new LodashWrapper(this.value(),this.__chain__)}var wrapperConcat=restParam(function(values){values=baseFlatten(values);return this.thru(function(array){return arrayConcat(isArray(array)?array:[toObject(array)],values)})});function wrapperPlant(value){var result,parent=this;while(parent instanceof baseLodash){var clone=wrapperClone(parent);if(result){previous.__wrapped__=clone}else{result=clone}var previous=clone;parent=parent.__wrapped__}previous.__wrapped__=value;return result}function wrapperReverse(){var value=this.__wrapped__;var interceptor=function(value){return wrapped&&wrapped.__dir__<0?value:value.reverse()};if(value instanceof LazyWrapper){var wrapped=value;if(this.__actions__.length){wrapped=new LazyWrapper(this)}wrapped=wrapped.reverse();wrapped.__actions__.push({func:thru,args:[interceptor],thisArg:undefined});return new LodashWrapper(wrapped,this.__chain__)}return this.thru(interceptor)}function wrapperToString(){return this.value()+""}function wrapperValue(){return baseWrapperValue(this.__wrapped__,this.__actions__)}var at=restParam(function(collection,props){return baseAt(collection,baseFlatten(props))});var countBy=createAggregator(function(result,value,key){hasOwnProperty.call(result,key)?++result[key]:result[key]=1});function every(collection,predicate,thisArg){var func=isArray(collection)?arrayEvery:baseEvery;if(thisArg&&isIterateeCall(collection,predicate,thisArg)){predicate=undefined}if(typeof predicate!="function"||thisArg!==undefined){predicate=getCallback(predicate,thisArg,3)}return func(collection,predicate)}function filter(collection,predicate,thisArg){var func=isArray(collection)?arrayFilter:baseFilter;predicate=getCallback(predicate,thisArg,3);return func(collection,predicate)}var find=createFind(baseEach);var findLast=createFind(baseEachRight,true);function findWhere(collection,source){return find(collection,baseMatches(source))}var forEach=createForEach(arrayEach,baseEach);var forEachRight=createForEach(arrayEachRight,baseEachRight);var groupBy=createAggregator(function(result,value,key){if(hasOwnProperty.call(result,key)){result[key].push(value)}else{result[key]=[value]}});function includes(collection,target,fromIndex,guard){var length=collection?getLength(collection):0;if(!isLength(length)){collection=values(collection);length=collection.length}if(typeof fromIndex!="number"||guard&&isIterateeCall(target,fromIndex,guard)){fromIndex=0}else{fromIndex=fromIndex<0?nativeMax(length+fromIndex,0):fromIndex||0}return typeof collection=="string"||!isArray(collection)&&isString(collection)?fromIndex<=length&&collection.indexOf(target,fromIndex)>-1:!!length&&getIndexOf(collection,target,fromIndex)>-1}var indexBy=createAggregator(function(result,value,key){result[key]=value});var invoke=restParam(function(collection,path,args){var index=-1,isFunc=typeof path=="function",isProp=isKey(path),result=isArrayLike(collection)?Array(collection.length):[];baseEach(collection,function(value){var func=isFunc?path:isProp&&value!=null?value[path]:undefined;result[++index]=func?func.apply(value,args):invokePath(value,path,args)});return result});function map(collection,iteratee,thisArg){var func=isArray(collection)?arrayMap:baseMap;iteratee=getCallback(iteratee,thisArg,3);return func(collection,iteratee)}var partition=createAggregator(function(result,value,key){result[key?0:1].push(value)},function(){return[[],[]]});function pluck(collection,path){return map(collection,property(path))}var reduce=createReduce(arrayReduce,baseEach);var reduceRight=createReduce(arrayReduceRight,baseEachRight);function reject(collection,predicate,thisArg){var func=isArray(collection)?arrayFilter:baseFilter;predicate=getCallback(predicate,thisArg,3);return func(collection,function(value,index,collection){return!predicate(value,index,collection)})}function sample(collection,n,guard){if(guard?isIterateeCall(collection,n,guard):n==null){collection=toIterable(collection);var length=collection.length;return length>0?collection[baseRandom(0,length-1)]:undefined}var index=-1,result=toArray(collection),length=result.length,lastIndex=length-1;n=nativeMin(n<0?0:+n||0,length);while(++index<n){var rand=baseRandom(index,lastIndex),value=result[rand];result[rand]=result[index];result[index]=value}result.length=n;return result}function shuffle(collection){return sample(collection,POSITIVE_INFINITY)}function size(collection){var length=collection?getLength(collection):0;return isLength(length)?length:keys(collection).length}function some(collection,predicate,thisArg){var func=isArray(collection)?arraySome:baseSome;if(thisArg&&isIterateeCall(collection,predicate,thisArg)){predicate=undefined}if(typeof predicate!="function"||thisArg!==undefined){predicate=getCallback(predicate,thisArg,3)}return func(collection,predicate)}function sortBy(collection,iteratee,thisArg){if(collection==null){return[]}if(thisArg&&isIterateeCall(collection,iteratee,thisArg)){iteratee=undefined}var index=-1;iteratee=getCallback(iteratee,thisArg,3);var result=baseMap(collection,function(value,key,collection){return{criteria:iteratee(value,key,collection),index:++index,value:value}});return baseSortBy(result,compareAscending)}var sortByAll=restParam(function(collection,iteratees){if(collection==null){return[]}var guard=iteratees[2];if(guard&&isIterateeCall(iteratees[0],iteratees[1],guard)){iteratees.length=1}return baseSortByOrder(collection,baseFlatten(iteratees),[])});function sortByOrder(collection,iteratees,orders,guard){if(collection==null){return[]}if(guard&&isIterateeCall(iteratees,orders,guard)){orders=undefined}if(!isArray(iteratees)){iteratees=iteratees==null?[]:[iteratees]}if(!isArray(orders)){orders=orders==null?[]:[orders]}return baseSortByOrder(collection,iteratees,orders)}function where(collection,source){return filter(collection,baseMatches(source))}var now=nativeNow||function(){return(new Date).getTime()};function after(n,func){if(typeof func!="function"){if(typeof n=="function"){var temp=n;n=func;func=temp}else{throw new TypeError(FUNC_ERROR_TEXT)}}n=nativeIsFinite(n=+n)?n:0;return function(){if(--n<1){return func.apply(this,arguments)}}}function ary(func,n,guard){if(guard&&isIterateeCall(func,n,guard)){n=undefined}n=func&&n==null?func.length:nativeMax(+n||0,0);return createWrapper(func,ARY_FLAG,undefined,undefined,undefined,undefined,n)}function before(n,func){var result;if(typeof func!="function"){if(typeof n=="function"){var temp=n;n=func;func=temp}else{throw new TypeError(FUNC_ERROR_TEXT)}}return function(){if(--n>0){result=func.apply(this,arguments)}if(n<=1){func=undefined}return result}}var bind=restParam(function(func,thisArg,partials){var bitmask=BIND_FLAG;if(partials.length){var holders=replaceHolders(partials,bind.placeholder);bitmask|=PARTIAL_FLAG}return createWrapper(func,bitmask,thisArg,partials,holders)});var bindAll=restParam(function(object,methodNames){methodNames=methodNames.length?baseFlatten(methodNames):functions(object);var index=-1,length=methodNames.length;while(++index<length){var key=methodNames[index];object[key]=createWrapper(object[key],BIND_FLAG,object)}return object});var bindKey=restParam(function(object,key,partials){var bitmask=BIND_FLAG|BIND_KEY_FLAG;if(partials.length){var holders=replaceHolders(partials,bindKey.placeholder);bitmask|=PARTIAL_FLAG}return createWrapper(key,bitmask,object,partials,holders)});var curry=createCurry(CURRY_FLAG);var curryRight=createCurry(CURRY_RIGHT_FLAG);function debounce(func,wait,options){var args,maxTimeoutId,result,stamp,thisArg,timeoutId,trailingCall,lastCalled=0,maxWait=false,trailing=true;if(typeof func!="function"){throw new TypeError(FUNC_ERROR_TEXT)}wait=wait<0?0:+wait||0;if(options===true){var leading=true;trailing=false}else if(isObject(options)){leading=!!options.leading;maxWait="maxWait"in options&&nativeMax(+options.maxWait||0,wait);trailing="trailing"in options?!!options.trailing:trailing}function cancel(){if(timeoutId){clearTimeout(timeoutId)}if(maxTimeoutId){clearTimeout(maxTimeoutId)}lastCalled=0;maxTimeoutId=timeoutId=trailingCall=undefined}function complete(isCalled,id){if(id){clearTimeout(id)}maxTimeoutId=timeoutId=trailingCall=undefined;if(isCalled){lastCalled=now();result=func.apply(thisArg,args);if(!timeoutId&&!maxTimeoutId){args=thisArg=undefined}}}function delayed(){var remaining=wait-(now()-stamp);if(remaining<=0||remaining>wait){complete(trailingCall,maxTimeoutId)}else{timeoutId=setTimeout(delayed,remaining)}}function maxDelayed(){complete(trailing,timeoutId)}function debounced(){args=arguments;stamp=now();thisArg=this;trailingCall=trailing&&(timeoutId||!leading);if(maxWait===false){var leadingCall=leading&&!timeoutId}else{if(!maxTimeoutId&&!leading){lastCalled=stamp}var remaining=maxWait-(stamp-lastCalled),isCalled=remaining<=0||remaining>maxWait;if(isCalled){if(maxTimeoutId){maxTimeoutId=clearTimeout(maxTimeoutId)}lastCalled=stamp;result=func.apply(thisArg,args)}else if(!maxTimeoutId){maxTimeoutId=setTimeout(maxDelayed,remaining)}}if(isCalled&&timeoutId){timeoutId=clearTimeout(timeoutId)}else if(!timeoutId&&wait!==maxWait){timeoutId=setTimeout(delayed,wait)}if(leadingCall){isCalled=true;result=func.apply(thisArg,args)}if(isCalled&&!timeoutId&&!maxTimeoutId){args=thisArg=undefined}return result}debounced.cancel=cancel;return debounced}var defer=restParam(function(func,args){return baseDelay(func,1,args)});var delay=restParam(function(func,wait,args){return baseDelay(func,wait,args)});var flow=createFlow();var flowRight=createFlow(true);function memoize(func,resolver){if(typeof func!="function"||resolver&&typeof resolver!="function"){throw new TypeError(FUNC_ERROR_TEXT)}var memoized=function(){var args=arguments,key=resolver?resolver.apply(this,args):args[0],cache=memoized.cache;if(cache.has(key)){return cache.get(key)}var result=func.apply(this,args);memoized.cache=cache.set(key,result);return result};memoized.cache=new memoize.Cache;return memoized}var modArgs=restParam(function(func,transforms){transforms=baseFlatten(transforms);if(typeof func!="function"||!arrayEvery(transforms,baseIsFunction)){throw new TypeError(FUNC_ERROR_TEXT)}var length=transforms.length;return restParam(function(args){var index=nativeMin(args.length,length);while(index--){args[index]=transforms[index](args[index])}return func.apply(this,args)})});function negate(predicate){if(typeof predicate!="function"){throw new TypeError(FUNC_ERROR_TEXT)}return function(){return!predicate.apply(this,arguments)}}function once(func){return before(2,func)}var partial=createPartial(PARTIAL_FLAG);var partialRight=createPartial(PARTIAL_RIGHT_FLAG);var rearg=restParam(function(func,indexes){return createWrapper(func,REARG_FLAG,undefined,undefined,undefined,baseFlatten(indexes))});function restParam(func,start){if(typeof func!="function"){throw new TypeError(FUNC_ERROR_TEXT)}start=nativeMax(start===undefined?func.length-1:+start||0,0);return function(){var args=arguments,index=-1,length=nativeMax(args.length-start,0),rest=Array(length);while(++index<length){rest[index]=args[start+index]}switch(start){case 0:return func.call(this,rest);case 1:return func.call(this,args[0],rest);case 2:return func.call(this,args[0],args[1],rest)}var otherArgs=Array(start+1);index=-1;while(++index<start){otherArgs[index]=args[index]}otherArgs[start]=rest;return func.apply(this,otherArgs)}}function spread(func){if(typeof func!="function"){throw new TypeError(FUNC_ERROR_TEXT)}return function(array){return func.apply(this,array)}}function throttle(func,wait,options){var leading=true,trailing=true;if(typeof func!="function"){throw new TypeError(FUNC_ERROR_TEXT)}if(options===false){leading=false}else if(isObject(options)){leading="leading"in options?!!options.leading:leading;trailing="trailing"in options?!!options.trailing:trailing}return debounce(func,wait,{leading:leading,maxWait:+wait,trailing:trailing})}function wrap(value,wrapper){wrapper=wrapper==null?identity:wrapper;return createWrapper(wrapper,PARTIAL_FLAG,undefined,[value],[])}function clone(value,isDeep,customizer,thisArg){if(isDeep&&typeof isDeep!="boolean"&&isIterateeCall(value,isDeep,customizer)){isDeep=false}else if(typeof isDeep=="function"){thisArg=customizer;customizer=isDeep;isDeep=false}return typeof customizer=="function"?baseClone(value,isDeep,bindCallback(customizer,thisArg,1)):baseClone(value,isDeep)}function cloneDeep(value,customizer,thisArg){return typeof customizer=="function"?baseClone(value,true,bindCallback(customizer,thisArg,1)):baseClone(value,true)}function gt(value,other){return value>other}function gte(value,other){return value>=other}function isArguments(value){return isObjectLike(value)&&isArrayLike(value)&&hasOwnProperty.call(value,"callee")&&!propertyIsEnumerable.call(value,"callee")}var isArray=nativeIsArray||function(value){return isObjectLike(value)&&isLength(value.length)&&objToString.call(value)==arrayTag};function isBoolean(value){return value===true||value===false||isObjectLike(value)&&objToString.call(value)==boolTag}function isDate(value){return isObjectLike(value)&&objToString.call(value)==dateTag}function isElement(value){return!!value&&value.nodeType===1&&isObjectLike(value)&&!isPlainObject(value)}function isEmpty(value){if(value==null){return true}if(isArrayLike(value)&&(isArray(value)||isString(value)||isArguments(value)||isObjectLike(value)&&isFunction(value.splice))){return!value.length}return!keys(value).length}function isEqual(value,other,customizer,thisArg){customizer=typeof customizer=="function"?bindCallback(customizer,thisArg,3):undefined;var result=customizer?customizer(value,other):undefined;return result===undefined?baseIsEqual(value,other,customizer):!!result}function isError(value){return isObjectLike(value)&&typeof value.message=="string"&&objToString.call(value)==errorTag}function isFinite(value){return typeof value=="number"&&nativeIsFinite(value)}function isFunction(value){return isObject(value)&&objToString.call(value)==funcTag}function isObject(value){var type=typeof value;return!!value&&(type=="object"||type=="function")}function isMatch(object,source,customizer,thisArg){customizer=typeof customizer=="function"?bindCallback(customizer,thisArg,3):undefined;return baseIsMatch(object,getMatchData(source),customizer)}function isNaN(value){return isNumber(value)&&value!=+value}function isNative(value){if(value==null){return false}if(isFunction(value)){return reIsNative.test(fnToString.call(value))}return isObjectLike(value)&&reIsHostCtor.test(value)}function isNull(value){return value===null}function isNumber(value){return typeof value=="number"||isObjectLike(value)&&objToString.call(value)==numberTag}function isPlainObject(value){var Ctor;if(!(isObjectLike(value)&&objToString.call(value)==objectTag&&!isArguments(value))||!hasOwnProperty.call(value,"constructor")&&(Ctor=value.constructor,typeof Ctor=="function"&&!(Ctor instanceof Ctor))){return false}var result;baseForIn(value,function(subValue,key){result=key});return result===undefined||hasOwnProperty.call(value,result)}function isRegExp(value){return isObject(value)&&objToString.call(value)==regexpTag}function isString(value){return typeof value=="string"||isObjectLike(value)&&objToString.call(value)==stringTag}function isTypedArray(value){return isObjectLike(value)&&isLength(value.length)&&!!typedArrayTags[objToString.call(value)]}function isUndefined(value){return value===undefined}function lt(value,other){return value<other}function lte(value,other){return value<=other}function toArray(value){var length=value?getLength(value):0;if(!isLength(length)){return values(value)}if(!length){return[]}return arrayCopy(value)}function toPlainObject(value){return baseCopy(value,keysIn(value))}var merge=createAssigner(baseMerge);var assign=createAssigner(function(object,source,customizer){return customizer?assignWith(object,source,customizer):baseAssign(object,source)});function create(prototype,properties,guard){var result=baseCreate(prototype);if(guard&&isIterateeCall(prototype,properties,guard)){properties=undefined}return properties?baseAssign(result,properties):result}var defaults=createDefaults(assign,assignDefaults);var defaultsDeep=createDefaults(merge,mergeDefaults);var findKey=createFindKey(baseForOwn);var findLastKey=createFindKey(baseForOwnRight);var forIn=createForIn(baseFor);var forInRight=createForIn(baseForRight);var forOwn=createForOwn(baseForOwn);var forOwnRight=createForOwn(baseForOwnRight);function functions(object){return baseFunctions(object,keysIn(object))}function get(object,path,defaultValue){var result=object==null?undefined:baseGet(object,toPath(path),path+"");return result===undefined?defaultValue:result}function has(object,path){if(object==null){return false}var result=hasOwnProperty.call(object,path);if(!result&&!isKey(path)){path=toPath(path);object=path.length==1?object:baseGet(object,baseSlice(path,0,-1));if(object==null){return false}path=last(path);result=hasOwnProperty.call(object,path)}return result||isLength(object.length)&&isIndex(path,object.length)&&(isArray(object)||isArguments(object))}function invert(object,multiValue,guard){if(guard&&isIterateeCall(object,multiValue,guard)){multiValue=undefined}var index=-1,props=keys(object),length=props.length,result={};while(++index<length){var key=props[index],value=object[key];if(multiValue){if(hasOwnProperty.call(result,value)){result[value].push(key)}else{result[value]=[key]}}else{result[value]=key}}return result}var keys=!nativeKeys?shimKeys:function(object){var Ctor=object==null?undefined:object.constructor;if(typeof Ctor=="function"&&Ctor.prototype===object||typeof object!="function"&&isArrayLike(object)){return shimKeys(object)}return isObject(object)?nativeKeys(object):[]};function keysIn(object){if(object==null){return[]}if(!isObject(object)){object=Object(object)}var length=object.length;length=length&&isLength(length)&&(isArray(object)||isArguments(object))&&length||0;var Ctor=object.constructor,index=-1,isProto=typeof Ctor=="function"&&Ctor.prototype===object,result=Array(length),skipIndexes=length>0;while(++index<length){result[index]=index+""}for(var key in object){if(!(skipIndexes&&isIndex(key,length))&&!(key=="constructor"&&(isProto||!hasOwnProperty.call(object,key)))){result.push(key)}}return result}var mapKeys=createObjectMapper(true);var mapValues=createObjectMapper();var omit=restParam(function(object,props){if(object==null){return{}}if(typeof props[0]!="function"){var props=arrayMap(baseFlatten(props),String);return pickByArray(object,baseDifference(keysIn(object),props))}var predicate=bindCallback(props[0],props[1],3);return pickByCallback(object,function(value,key,object){return!predicate(value,key,object)})});function pairs(object){object=toObject(object);var index=-1,props=keys(object),length=props.length,result=Array(length);while(++index<length){var key=props[index];result[index]=[key,object[key]]}return result}var pick=restParam(function(object,props){if(object==null){return{}}return typeof props[0]=="function"?pickByCallback(object,bindCallback(props[0],props[1],3)):pickByArray(object,baseFlatten(props))});function result(object,path,defaultValue){var result=object==null?undefined:object[path];if(result===undefined){if(object!=null&&!isKey(path,object)){path=toPath(path);object=path.length==1?object:baseGet(object,baseSlice(path,0,-1));result=object==null?undefined:object[last(path)]}result=result===undefined?defaultValue:result}return isFunction(result)?result.call(object):result}function set(object,path,value){if(object==null){return object}var pathKey=path+"";path=object[pathKey]!=null||isKey(path,object)?[pathKey]:toPath(path);var index=-1,length=path.length,lastIndex=length-1,nested=object;while(nested!=null&&++index<length){var key=path[index];if(isObject(nested)){if(index==lastIndex){nested[key]=value}else if(nested[key]==null){nested[key]=isIndex(path[index+1])?[]:{}}}nested=nested[key]}return object}function transform(object,iteratee,accumulator,thisArg){var isArr=isArray(object)||isTypedArray(object);iteratee=getCallback(iteratee,thisArg,4);if(accumulator==null){if(isArr||isObject(object)){var Ctor=object.constructor;if(isArr){accumulator=isArray(object)?new Ctor:[]}else{accumulator=baseCreate(isFunction(Ctor)?Ctor.prototype:undefined)}}else{accumulator={}}}(isArr?arrayEach:baseForOwn)(object,function(value,index,object){return iteratee(accumulator,value,index,object)});return accumulator}function values(object){return baseValues(object,keys(object))}function valuesIn(object){return baseValues(object,keysIn(object))}function inRange(value,start,end){start=+start||0;if(end===undefined){end=start;start=0}else{end=+end||0}return value>=nativeMin(start,end)&&value<nativeMax(start,end)}function random(min,max,floating){if(floating&&isIterateeCall(min,max,floating)){max=floating=undefined}var noMin=min==null,noMax=max==null;if(floating==null){if(noMax&&typeof min=="boolean"){floating=min;min=1}else if(typeof max=="boolean"){floating=max;noMax=true}}if(noMin&&noMax){max=1;noMax=false}min=+min||0;if(noMax){max=min;min=0}else{max=+max||0}if(floating||min%1||max%1){var rand=nativeRandom();return nativeMin(min+rand*(max-min+parseFloat("1e-"+((rand+"").length-1))),max)}return baseRandom(min,max)}var camelCase=createCompounder(function(result,word,index){word=word.toLowerCase();return result+(index?word.charAt(0).toUpperCase()+word.slice(1):word)});function capitalize(string){string=baseToString(string);return string&&string.charAt(0).toUpperCase()+string.slice(1)}function deburr(string){string=baseToString(string);return string&&string.replace(reLatin1,deburrLetter).replace(reComboMark,"")}function endsWith(string,target,position){string=baseToString(string);target=target+"";var length=string.length;position=position===undefined?length:nativeMin(position<0?0:+position||0,length);position-=target.length;return position>=0&&string.indexOf(target,position)==position}function escape(string){string=baseToString(string);return string&&reHasUnescapedHtml.test(string)?string.replace(reUnescapedHtml,escapeHtmlChar):string}function escapeRegExp(string){string=baseToString(string);return string&&reHasRegExpChars.test(string)?string.replace(reRegExpChars,escapeRegExpChar):string||"(?:)"}var kebabCase=createCompounder(function(result,word,index){return result+(index?"-":"")+word.toLowerCase()});function pad(string,length,chars){string=baseToString(string);length=+length;var strLength=string.length;if(strLength>=length||!nativeIsFinite(length)){return string}var mid=(length-strLength)/2,leftLength=nativeFloor(mid),rightLength=nativeCeil(mid);chars=createPadding("",rightLength,chars);return chars.slice(0,leftLength)+string+chars}var padLeft=createPadDir();var padRight=createPadDir(true);function parseInt(string,radix,guard){if(guard?isIterateeCall(string,radix,guard):radix==null){radix=0}else if(radix){radix=+radix}string=trim(string);return nativeParseInt(string,radix||(reHasHexPrefix.test(string)?16:10))}function repeat(string,n){var result="";string=baseToString(string);n=+n;if(n<1||!string||!nativeIsFinite(n)){return result}do{if(n%2){result+=string}n=nativeFloor(n/2);string+=string}while(n);return result}var snakeCase=createCompounder(function(result,word,index){return result+(index?"_":"")+word.toLowerCase()});var startCase=createCompounder(function(result,word,index){return result+(index?" ":"")+(word.charAt(0).toUpperCase()+word.slice(1))});function startsWith(string,target,position){string=baseToString(string);position=position==null?0:nativeMin(position<0?0:+position||0,string.length);return string.lastIndexOf(target,position)==position}function template(string,options,otherOptions){var settings=lodash.templateSettings;if(otherOptions&&isIterateeCall(string,options,otherOptions)){options=otherOptions=undefined}string=baseToString(string);options=assignWith(baseAssign({},otherOptions||options),settings,assignOwnDefaults);var imports=assignWith(baseAssign({},options.imports),settings.imports,assignOwnDefaults),importsKeys=keys(imports),importsValues=baseValues(imports,importsKeys);var isEscaping,isEvaluating,index=0,interpolate=options.interpolate||reNoMatch,source="__p += '";var reDelimiters=RegExp((options.escape||reNoMatch).source+"|"+interpolate.source+"|"+(interpolate===reInterpolate?reEsTemplate:reNoMatch).source+"|"+(options.evaluate||reNoMatch).source+"|$","g");var sourceURL="//# sourceURL="+("sourceURL"in options?options.sourceURL:"lodash.templateSources["+ ++templateCounter+"]")+"\n";string.replace(reDelimiters,function(match,escapeValue,interpolateValue,esTemplateValue,evaluateValue,offset){interpolateValue||(interpolateValue=esTemplateValue);source+=string.slice(index,offset).replace(reUnescapedString,escapeStringChar);if(escapeValue){isEscaping=true;source+="' +\n__e("+escapeValue+") +\n'"}if(evaluateValue){isEvaluating=true;source+="';\n"+evaluateValue+";\n__p += '"}if(interpolateValue){source+="' +\n((__t = ("+interpolateValue+")) == null ? '' : __t) +\n'"}index=offset+match.length;return match});source+="';\n";var variable=options.variable;if(!variable){source="with (obj) {\n"+source+"\n}\n"}source=(isEvaluating?source.replace(reEmptyStringLeading,""):source).replace(reEmptyStringMiddle,"$1").replace(reEmptyStringTrailing,"$1;");source="function("+(variable||"obj")+") {\n"+(variable?"":"obj || (obj = {});\n")+"var __t, __p = ''"+(isEscaping?", __e = _.escape":"")+(isEvaluating?", __j = Array.prototype.join;\n"+"function print() { __p += __j.call(arguments, '') }\n":";\n")+source+"return __p\n}";var result=attempt(function(){return Function(importsKeys,sourceURL+"return "+source).apply(undefined,importsValues)});result.source=source;if(isError(result)){throw result}return result}function trim(string,chars,guard){var value=string;string=baseToString(string);if(!string){return string}if(guard?isIterateeCall(value,chars,guard):chars==null){return string.slice(trimmedLeftIndex(string),trimmedRightIndex(string)+1)}chars=chars+"";return string.slice(charsLeftIndex(string,chars),charsRightIndex(string,chars)+1)}function trimLeft(string,chars,guard){var value=string;string=baseToString(string);if(!string){return string}if(guard?isIterateeCall(value,chars,guard):chars==null){return string.slice(trimmedLeftIndex(string))}return string.slice(charsLeftIndex(string,chars+""))}function trimRight(string,chars,guard){var value=string;string=baseToString(string);if(!string){return string}if(guard?isIterateeCall(value,chars,guard):chars==null){return string.slice(0,trimmedRightIndex(string)+1)}return string.slice(0,charsRightIndex(string,chars+"")+1)}function trunc(string,options,guard){if(guard&&isIterateeCall(string,options,guard)){options=undefined}var length=DEFAULT_TRUNC_LENGTH,omission=DEFAULT_TRUNC_OMISSION;if(options!=null){if(isObject(options)){var separator="separator"in options?options.separator:separator;length="length"in options?+options.length||0:length;omission="omission"in options?baseToString(options.omission):omission}else{length=+options||0}}string=baseToString(string);if(length>=string.length){return string}var end=length-omission.length;if(end<1){return omission}var result=string.slice(0,end);if(separator==null){return result+omission}if(isRegExp(separator)){if(string.slice(end).search(separator)){var match,newEnd,substring=string.slice(0,end);if(!separator.global){separator=RegExp(separator.source,(reFlags.exec(separator)||"")+"g")}separator.lastIndex=0;while(match=separator.exec(substring)){newEnd=match.index}result=result.slice(0,newEnd==null?end:newEnd)}}else if(string.indexOf(separator,end)!=end){var index=result.lastIndexOf(separator);if(index>-1){result=result.slice(0,index)}}return result+omission}function unescape(string){string=baseToString(string);return string&&reHasEscapedHtml.test(string)?string.replace(reEscapedHtml,unescapeHtmlChar):string}function words(string,pattern,guard){if(guard&&isIterateeCall(string,pattern,guard)){pattern=undefined}string=baseToString(string);return string.match(pattern||reWords)||[]}var attempt=restParam(function(func,args){try{return func.apply(undefined,args)}catch(e){return isError(e)?e:new Error(e)}});function callback(func,thisArg,guard){if(guard&&isIterateeCall(func,thisArg,guard)){thisArg=undefined}return isObjectLike(func)?matches(func):baseCallback(func,thisArg)}function constant(value){return function(){return value}}function identity(value){return value}function matches(source){return baseMatches(baseClone(source,true))}function matchesProperty(path,srcValue){return baseMatchesProperty(path,baseClone(srcValue,true))}var method=restParam(function(path,args){return function(object){return invokePath(object,path,args)}});var methodOf=restParam(function(object,args){return function(path){return invokePath(object,path,args)}});function mixin(object,source,options){if(options==null){var isObj=isObject(source),props=isObj?keys(source):undefined,methodNames=props&&props.length?baseFunctions(source,props):undefined;if(!(methodNames?methodNames.length:isObj)){methodNames=false;options=source;source=object;object=this}}if(!methodNames){methodNames=baseFunctions(source,keys(source))}var chain=true,index=-1,isFunc=isFunction(object),length=methodNames.length;if(options===false){chain=false}else if(isObject(options)&&"chain"in options){chain=options.chain}while(++index<length){var methodName=methodNames[index],func=source[methodName];object[methodName]=func;if(isFunc){object.prototype[methodName]=function(func){return function(){var chainAll=this.__chain__;if(chain||chainAll){var result=object(this.__wrapped__),actions=result.__actions__=arrayCopy(this.__actions__);actions.push({func:func,args:arguments,thisArg:object});result.__chain__=chainAll;return result}return func.apply(object,arrayPush([this.value()],arguments))}}(func)}}return object}function noConflict(){root._=oldDash;return this}function noop(){}function property(path){return isKey(path)?baseProperty(path):basePropertyDeep(path)}function propertyOf(object){return function(path){return baseGet(object,toPath(path),path+"")}}function range(start,end,step){if(step&&isIterateeCall(start,end,step)){end=step=undefined}start=+start||0;step=step==null?1:+step||0;if(end==null){end=start;start=0}else{end=+end||0}var index=-1,length=nativeMax(nativeCeil((end-start)/(step||1)),0),result=Array(length);while(++index<length){result[index]=start;start+=step}return result}function times(n,iteratee,thisArg){n=nativeFloor(n);if(n<1||!nativeIsFinite(n)){return[]}var index=-1,result=Array(nativeMin(n,MAX_ARRAY_LENGTH));iteratee=bindCallback(iteratee,thisArg,1);while(++index<n){if(index<MAX_ARRAY_LENGTH){result[index]=iteratee(index)}else{iteratee(index)}}return result}function uniqueId(prefix){var id=++idCounter;return baseToString(prefix)+id}function add(augend,addend){return(+augend||0)+(+addend||0)}var ceil=createRound("ceil");var floor=createRound("floor");var max=createExtremum(gt,NEGATIVE_INFINITY);var min=createExtremum(lt,POSITIVE_INFINITY);var round=createRound("round");function sum(collection,iteratee,thisArg){if(thisArg&&isIterateeCall(collection,iteratee,thisArg)){iteratee=undefined}iteratee=getCallback(iteratee,thisArg,3);return iteratee.length==1?arraySum(isArray(collection)?collection:toIterable(collection),iteratee):baseSum(collection,iteratee)}lodash.prototype=baseLodash.prototype;LodashWrapper.prototype=baseCreate(baseLodash.prototype);LodashWrapper.prototype.constructor=LodashWrapper;LazyWrapper.prototype=baseCreate(baseLodash.prototype);LazyWrapper.prototype.constructor=LazyWrapper;MapCache.prototype["delete"]=mapDelete;MapCache.prototype.get=mapGet;MapCache.prototype.has=mapHas;MapCache.prototype.set=mapSet;SetCache.prototype.push=cachePush;memoize.Cache=MapCache;lodash.after=after;lodash.ary=ary;lodash.assign=assign;lodash.at=at;lodash.before=before;lodash.bind=bind;lodash.bindAll=bindAll;lodash.bindKey=bindKey;lodash.callback=callback;lodash.chain=chain;lodash.chunk=chunk;lodash.compact=compact;lodash.constant=constant;lodash.countBy=countBy;lodash.create=create;lodash.curry=curry;lodash.curryRight=curryRight;lodash.debounce=debounce;lodash.defaults=defaults;lodash.defaultsDeep=defaultsDeep;lodash.defer=defer;lodash.delay=delay;lodash.difference=difference;lodash.drop=drop;lodash.dropRight=dropRight;lodash.dropRightWhile=dropRightWhile;lodash.dropWhile=dropWhile;lodash.fill=fill;lodash.filter=filter;lodash.flatten=flatten;lodash.flattenDeep=flattenDeep;lodash.flow=flow;lodash.flowRight=flowRight;lodash.forEach=forEach;lodash.forEachRight=forEachRight;lodash.forIn=forIn;lodash.forInRight=forInRight;lodash.forOwn=forOwn;lodash.forOwnRight=forOwnRight;lodash.functions=functions;lodash.groupBy=groupBy;lodash.indexBy=indexBy;lodash.initial=initial;lodash.intersection=intersection;lodash.invert=invert;lodash.invoke=invoke;lodash.keys=keys;lodash.keysIn=keysIn;lodash.map=map;lodash.mapKeys=mapKeys;lodash.mapValues=mapValues;lodash.matches=matches;lodash.matchesProperty=matchesProperty;lodash.memoize=memoize;lodash.merge=merge;lodash.method=method;lodash.methodOf=methodOf;lodash.mixin=mixin;lodash.modArgs=modArgs;lodash.negate=negate;lodash.omit=omit;lodash.once=once;lodash.pairs=pairs;lodash.partial=partial;lodash.partialRight=partialRight;lodash.partition=partition;lodash.pick=pick;lodash.pluck=pluck;lodash.property=property;lodash.propertyOf=propertyOf;lodash.pull=pull;lodash.pullAt=pullAt;lodash.range=range;lodash.rearg=rearg;lodash.reject=reject;lodash.remove=remove;lodash.rest=rest;lodash.restParam=restParam;lodash.set=set;lodash.shuffle=shuffle;lodash.slice=slice;lodash.sortBy=sortBy;lodash.sortByAll=sortByAll;lodash.sortByOrder=sortByOrder;lodash.spread=spread;lodash.take=take;lodash.takeRight=takeRight;lodash.takeRightWhile=takeRightWhile;lodash.takeWhile=takeWhile;lodash.tap=tap;lodash.throttle=throttle;lodash.thru=thru;lodash.times=times;lodash.toArray=toArray;lodash.toPlainObject=toPlainObject;lodash.transform=transform;lodash.union=union;lodash.uniq=uniq;lodash.unzip=unzip;lodash.unzipWith=unzipWith;lodash.values=values;lodash.valuesIn=valuesIn;lodash.where=where;lodash.without=without;lodash.wrap=wrap;lodash.xor=xor;lodash.zip=zip;lodash.zipObject=zipObject;lodash.zipWith=zipWith;lodash.backflow=flowRight;lodash.collect=map;lodash.compose=flowRight;lodash.each=forEach;lodash.eachRight=forEachRight;lodash.extend=assign;lodash.iteratee=callback;lodash.methods=functions;lodash.object=zipObject;lodash.select=filter;lodash.tail=rest;lodash.unique=uniq;mixin(lodash,lodash);lodash.add=add;lodash.attempt=attempt;lodash.camelCase=camelCase;lodash.capitalize=capitalize;lodash.ceil=ceil;lodash.clone=clone;lodash.cloneDeep=cloneDeep;lodash.deburr=deburr;lodash.endsWith=endsWith;lodash.escape=escape;lodash.escapeRegExp=escapeRegExp;lodash.every=every;lodash.find=find;lodash.findIndex=findIndex;lodash.findKey=findKey;lodash.findLast=findLast;lodash.findLastIndex=findLastIndex;lodash.findLastKey=findLastKey;lodash.findWhere=findWhere;lodash.first=first;lodash.floor=floor;lodash.get=get;lodash.gt=gt;lodash.gte=gte;lodash.has=has;lodash.identity=identity;lodash.includes=includes;lodash.indexOf=indexOf;lodash.inRange=inRange;lodash.isArguments=isArguments;lodash.isArray=isArray;lodash.isBoolean=isBoolean;lodash.isDate=isDate;lodash.isElement=isElement;lodash.isEmpty=isEmpty;lodash.isEqual=isEqual;lodash.isError=isError;lodash.isFinite=isFinite;lodash.isFunction=isFunction;lodash.isMatch=isMatch;lodash.isNaN=isNaN;lodash.isNative=isNative;lodash.isNull=isNull;lodash.isNumber=isNumber;lodash.isObject=isObject;lodash.isPlainObject=isPlainObject;lodash.isRegExp=isRegExp;lodash.isString=isString;lodash.isTypedArray=isTypedArray;lodash.isUndefined=isUndefined;lodash.kebabCase=kebabCase;lodash.last=last;lodash.lastIndexOf=lastIndexOf;lodash.lt=lt;lodash.lte=lte;lodash.max=max;lodash.min=min;lodash.noConflict=noConflict;lodash.noop=noop;lodash.now=now;lodash.pad=pad;lodash.padLeft=padLeft;lodash.padRight=padRight;lodash.parseInt=parseInt;lodash.random=random;lodash.reduce=reduce;lodash.reduceRight=reduceRight;lodash.repeat=repeat;lodash.result=result;lodash.round=round;lodash.runInContext=runInContext;lodash.size=size;lodash.snakeCase=snakeCase;lodash.some=some;lodash.sortedIndex=sortedIndex;lodash.sortedLastIndex=sortedLastIndex;lodash.startCase=startCase;lodash.startsWith=startsWith;lodash.sum=sum;lodash.template=template;lodash.trim=trim;lodash.trimLeft=trimLeft;lodash.trimRight=trimRight;lodash.trunc=trunc;lodash.unescape=unescape;lodash.uniqueId=uniqueId;lodash.words=words;lodash.all=every;lodash.any=some;lodash.contains=includes;lodash.eq=isEqual;lodash.detect=find;lodash.foldl=reduce;lodash.foldr=reduceRight;lodash.head=first;lodash.include=includes;lodash.inject=reduce;mixin(lodash,function(){var source={};baseForOwn(lodash,function(func,methodName){if(!lodash.prototype[methodName]){source[methodName]=func}});return source}(),false);lodash.sample=sample;lodash.prototype.sample=function(n){if(!this.__chain__&&n==null){return sample(this.value())}return this.thru(function(value){return sample(value,n)})};lodash.VERSION=VERSION;arrayEach(["bind","bindKey","curry","curryRight","partial","partialRight"],function(methodName){lodash[methodName].placeholder=lodash});arrayEach(["drop","take"],function(methodName,index){LazyWrapper.prototype[methodName]=function(n){var filtered=this.__filtered__;if(filtered&&!index){return new LazyWrapper(this)}n=n==null?1:nativeMax(nativeFloor(n)||0,0);var result=this.clone();if(filtered){result.__takeCount__=nativeMin(result.__takeCount__,n)}else{result.__views__.push({size:n,type:methodName+(result.__dir__<0?"Right":"")})}return result};LazyWrapper.prototype[methodName+"Right"]=function(n){return this.reverse()[methodName](n).reverse()}});arrayEach(["filter","map","takeWhile"],function(methodName,index){var type=index+1,isFilter=type!=LAZY_MAP_FLAG;LazyWrapper.prototype[methodName]=function(iteratee,thisArg){var result=this.clone();result.__iteratees__.push({iteratee:getCallback(iteratee,thisArg,1),type:type});result.__filtered__=result.__filtered__||isFilter;return result}});arrayEach(["first","last"],function(methodName,index){var takeName="take"+(index?"Right":"");LazyWrapper.prototype[methodName]=function(){return this[takeName](1).value()[0]}});arrayEach(["initial","rest"],function(methodName,index){var dropName="drop"+(index?"":"Right");LazyWrapper.prototype[methodName]=function(){return this.__filtered__?new LazyWrapper(this):this[dropName](1)}});arrayEach(["pluck","where"],function(methodName,index){var operationName=index?"filter":"map",createCallback=index?baseMatches:property;LazyWrapper.prototype[methodName]=function(value){return this[operationName](createCallback(value))}});LazyWrapper.prototype.compact=function(){return this.filter(identity)};LazyWrapper.prototype.reject=function(predicate,thisArg){predicate=getCallback(predicate,thisArg,1);return this.filter(function(value){return!predicate(value)})};LazyWrapper.prototype.slice=function(start,end){start=start==null?0:+start||0;var result=this;if(result.__filtered__&&(start>0||end<0)){return new LazyWrapper(result)}if(start<0){result=result.takeRight(-start)}else if(start){result=result.drop(start)}if(end!==undefined){end=+end||0;result=end<0?result.dropRight(-end):result.take(end-start)}return result};LazyWrapper.prototype.takeRightWhile=function(predicate,thisArg){return this.reverse().takeWhile(predicate,thisArg).reverse()};LazyWrapper.prototype.toArray=function(){return this.take(POSITIVE_INFINITY)};baseForOwn(LazyWrapper.prototype,function(func,methodName){var checkIteratee=/^(?:filter|map|reject)|While$/.test(methodName),retUnwrapped=/^(?:first|last)$/.test(methodName),lodashFunc=lodash[retUnwrapped?"take"+(methodName=="last"?"Right":""):methodName];if(!lodashFunc){return}lodash.prototype[methodName]=function(){var args=retUnwrapped?[1]:arguments,chainAll=this.__chain__,value=this.__wrapped__,isHybrid=!!this.__actions__.length,isLazy=value instanceof LazyWrapper,iteratee=args[0],useLazy=isLazy||isArray(value);if(useLazy&&checkIteratee&&typeof iteratee=="function"&&iteratee.length!=1){isLazy=useLazy=false}var interceptor=function(value){return retUnwrapped&&chainAll?lodashFunc(value,1)[0]:lodashFunc.apply(undefined,arrayPush([value],args))};var action={func:thru,args:[interceptor],thisArg:undefined},onlyLazy=isLazy&&!isHybrid;if(retUnwrapped&&!chainAll){if(onlyLazy){value=value.clone();value.__actions__.push(action);return func.call(value)}return lodashFunc.call(undefined,this.value())[0]}if(!retUnwrapped&&useLazy){value=onlyLazy?value:new LazyWrapper(this);var result=func.apply(value,args);result.__actions__.push(action);return new LodashWrapper(result,chainAll)}return this.thru(interceptor)}});arrayEach(["join","pop","push","replace","shift","sort","splice","split","unshift"],function(methodName){var func=(/^(?:replace|split)$/.test(methodName)?stringProto:arrayProto)[methodName],chainName=/^(?:push|sort|unshift)$/.test(methodName)?"tap":"thru",retUnwrapped=/^(?:join|pop|replace|shift)$/.test(methodName);lodash.prototype[methodName]=function(){var args=arguments;if(retUnwrapped&&!this.__chain__){return func.apply(this.value(),args)}return this[chainName](function(value){return func.apply(value,args)})}});baseForOwn(LazyWrapper.prototype,function(func,methodName){var lodashFunc=lodash[methodName];if(lodashFunc){var key=lodashFunc.name,names=realNames[key]||(realNames[key]=[]);names.push({name:methodName,func:lodashFunc})}});realNames[createHybridWrapper(undefined,BIND_KEY_FLAG).name]=[{name:"wrapper",func:undefined}];LazyWrapper.prototype.clone=lazyClone;LazyWrapper.prototype.reverse=lazyReverse;LazyWrapper.prototype.value=lazyValue;lodash.prototype.chain=wrapperChain;lodash.prototype.commit=wrapperCommit;lodash.prototype.concat=wrapperConcat;lodash.prototype.plant=wrapperPlant;lodash.prototype.reverse=wrapperReverse;lodash.prototype.toString=wrapperToString;lodash.prototype.run=lodash.prototype.toJSON=lodash.prototype.valueOf=lodash.prototype.value=wrapperValue;lodash.prototype.collect=lodash.prototype.map;lodash.prototype.head=lodash.prototype.first;lodash.prototype.select=lodash.prototype.filter;lodash.prototype.tail=lodash.prototype.rest;return lodash}var _=runInContext();if(typeof define=="function"&&typeof define.amd=="object"&&define.amd){root._=_;define(function(){return _})}else if(freeExports&&freeModule){if(moduleExports){(freeModule.exports=_)._=_}else{freeExports._=_}}else{root._=_}}).call(this)}).call(this,typeof global!=="undefined"?global:typeof self!=="undefined"?self:typeof window!=="undefined"?window:{})},{}],3:[function(require,module,exports){(function(window,document,undefined){var _MAP={8:"backspace",9:"tab",13:"enter",16:"shift",17:"ctrl",18:"alt",20:"capslock",27:"esc",32:"space",33:"pageup",34:"pagedown",35:"end",36:"home",37:"left",38:"up",39:"right",40:"down",45:"ins",46:"del",91:"meta",93:"meta",224:"meta"};var _KEYCODE_MAP={106:"*",107:"+",109:"-",110:".",111:"/",186:";",187:"=",188:",",189:"-",190:".",191:"/",192:"`",219:"[",220:"\\",221:"]",222:"'"};var _SHIFT_MAP={"~":"`","!":"1","@":"2","#":"3",$:"4","%":"5","^":"6","&":"7","*":"8","(":"9",")":"0",_:"-","+":"=",":":";",'"':"'","<":",",">":".","?":"/","|":"\\"};var _SPECIAL_ALIASES={option:"alt",command:"meta",return:"enter",escape:"esc",plus:"+",mod:/Mac|iPod|iPhone|iPad/.test(navigator.platform)?"meta":"ctrl"};var _REVERSE_MAP;for(var i=1;i<20;++i){_MAP[111+i]="f"+i}for(i=0;i<=9;++i){_MAP[i+96]=i}function _addEvent(object,type,callback){if(object.addEventListener){object.addEventListener(type,callback,false);return}object.attachEvent("on"+type,callback)}function _characterFromEvent(e){if(e.type=="keypress"){var character=String.fromCharCode(e.which);if(!e.shiftKey){character=character.toLowerCase()}return character}if(_MAP[e.which]){return _MAP[e.which]}if(_KEYCODE_MAP[e.which]){return _KEYCODE_MAP[e.which]}return String.fromCharCode(e.which).toLowerCase()}function _modifiersMatch(modifiers1,modifiers2){return modifiers1.sort().join(",")===modifiers2.sort().join(",")}function _eventModifiers(e){var modifiers=[];if(e.shiftKey){modifiers.push("shift")}if(e.altKey){modifiers.push("alt")}if(e.ctrlKey){modifiers.push("ctrl")}if(e.metaKey){modifiers.push("meta")}return modifiers}function _preventDefault(e){if(e.preventDefault){e.preventDefault();return}e.returnValue=false}function _stopPropagation(e){if(e.stopPropagation){e.stopPropagation();return}e.cancelBubble=true}function _isModifier(key){return key=="shift"||key=="ctrl"||key=="alt"||key=="meta"}function _getReverseMap(){if(!_REVERSE_MAP){_REVERSE_MAP={};for(var key in _MAP){if(key>95&&key<112){continue}if(_MAP.hasOwnProperty(key)){_REVERSE_MAP[_MAP[key]]=key}}}return _REVERSE_MAP}function _pickBestAction(key,modifiers,action){if(!action){action=_getReverseMap()[key]?"keydown":"keypress"}if(action=="keypress"&&modifiers.length){action="keydown"}return action}function _keysFromString(combination){if(combination==="+"){return["+"]}combination=combination.replace(/\+{2}/g,"+plus");return combination.split("+")}function _getKeyInfo(combination,action){var keys;var key;var i;var modifiers=[];keys=_keysFromString(combination);for(i=0;i<keys.length;++i){key=keys[i];if(_SPECIAL_ALIASES[key]){key=_SPECIAL_ALIASES[key]}if(action&&action!="keypress"&&_SHIFT_MAP[key]){key=_SHIFT_MAP[key];modifiers.push("shift")}if(_isModifier(key)){modifiers.push(key)}}action=_pickBestAction(key,modifiers,action);return{key:key,modifiers:modifiers,action:action}}function _belongsTo(element,ancestor){if(element===null||element===document){return false}if(element===ancestor){return true}return _belongsTo(element.parentNode,ancestor)}function Mousetrap(targetElement){var self=this;targetElement=targetElement||document;if(!(self instanceof Mousetrap)){return new Mousetrap(targetElement)}self.target=targetElement;self._callbacks={};self._directMap={};var _sequenceLevels={};var _resetTimer;var _ignoreNextKeyup=false;var _ignoreNextKeypress=false;var _nextExpectedAction=false;function _resetSequences(doNotReset){doNotReset=doNotReset||{};var activeSequences=false,key;for(key in _sequenceLevels){if(doNotReset[key]){activeSequences=true;continue}_sequenceLevels[key]=0}if(!activeSequences){_nextExpectedAction=false}}function _getMatches(character,modifiers,e,sequenceName,combination,level){var i;var callback;var matches=[];var action=e.type;if(!self._callbacks[character]){return[]}if(action=="keyup"&&_isModifier(character)){modifiers=[character]}for(i=0;i<self._callbacks[character].length;++i){callback=self._callbacks[character][i];if(!sequenceName&&callback.seq&&_sequenceLevels[callback.seq]!=callback.level){continue}if(action!=callback.action){continue}if(action=="keypress"&&!e.metaKey&&!e.ctrlKey||_modifiersMatch(modifiers,callback.modifiers)){var deleteCombo=!sequenceName&&callback.combo==combination;var deleteSequence=sequenceName&&callback.seq==sequenceName&&callback.level==level;if(deleteCombo||deleteSequence){self._callbacks[character].splice(i,1)}matches.push(callback)}}return matches}function _fireCallback(callback,e,combo,sequence){if(self.stopCallback(e,e.target||e.srcElement,combo,sequence)){return}if(callback(e,combo)===false){_preventDefault(e);_stopPropagation(e)}}self._handleKey=function(character,modifiers,e){var callbacks=_getMatches(character,modifiers,e);var i;var doNotReset={};var maxLevel=0;var processedSequenceCallback=false;for(i=0;i<callbacks.length;++i){if(callbacks[i].seq){maxLevel=Math.max(maxLevel,callbacks[i].level)}}for(i=0;i<callbacks.length;++i){if(callbacks[i].seq){if(callbacks[i].level!=maxLevel){continue}processedSequenceCallback=true;doNotReset[callbacks[i].seq]=1;_fireCallback(callbacks[i].callback,e,callbacks[i].combo,callbacks[i].seq);continue}if(!processedSequenceCallback){_fireCallback(callbacks[i].callback,e,callbacks[i].combo)}}var ignoreThisKeypress=e.type=="keypress"&&_ignoreNextKeypress;if(e.type==_nextExpectedAction&&!_isModifier(character)&&!ignoreThisKeypress){_resetSequences(doNotReset)}_ignoreNextKeypress=processedSequenceCallback&&e.type=="keydown"};function _handleKeyEvent(e){if(typeof e.which!=="number"){e.which=e.keyCode}var character=_characterFromEvent(e);if(!character){return}if(e.type=="keyup"&&_ignoreNextKeyup===character){_ignoreNextKeyup=false;return}self.handleKey(character,_eventModifiers(e),e)}function _resetSequenceTimer(){clearTimeout(_resetTimer);_resetTimer=setTimeout(_resetSequences,1e3)}function _bindSequence(combo,keys,callback,action){_sequenceLevels[combo]=0;function _increaseSequence(nextAction){return function(){_nextExpectedAction=nextAction;++_sequenceLevels[combo];_resetSequenceTimer()}}function _callbackAndReset(e){_fireCallback(callback,e,combo);if(action!=="keyup"){_ignoreNextKeyup=_characterFromEvent(e)}setTimeout(_resetSequences,10)}for(var i=0;i<keys.length;++i){var isFinal=i+1===keys.length;var wrappedCallback=isFinal?_callbackAndReset:_increaseSequence(action||_getKeyInfo(keys[i+1]).action);_bindSingle(keys[i],wrappedCallback,action,combo,i)}}function _bindSingle(combination,callback,action,sequenceName,level){self._directMap[combination+":"+action]=callback;combination=combination.replace(/\s+/g," ");var sequence=combination.split(" ");var info;if(sequence.length>1){_bindSequence(combination,sequence,callback,action);return}info=_getKeyInfo(combination,action);self._callbacks[info.key]=self._callbacks[info.key]||[];_getMatches(info.key,info.modifiers,{type:info.action},sequenceName,combination,level);self._callbacks[info.key][sequenceName?"unshift":"push"]({callback:callback,modifiers:info.modifiers,action:info.action,seq:sequenceName,level:level,combo:combination})}self._bindMultiple=function(combinations,callback,action){for(var i=0;i<combinations.length;++i){_bindSingle(combinations[i],callback,action)}};_addEvent(targetElement,"keypress",_handleKeyEvent);_addEvent(targetElement,"keydown",_handleKeyEvent);_addEvent(targetElement,"keyup",_handleKeyEvent)}Mousetrap.prototype.bind=function(keys,callback,action){var self=this;keys=keys instanceof Array?keys:[keys];self._bindMultiple.call(self,keys,callback,action);return self};Mousetrap.prototype.unbind=function(keys,action){var self=this;return self.bind.call(self,keys,function(){},action)};Mousetrap.prototype.trigger=function(keys,action){var self=this;if(self._directMap[keys+":"+action]){self._directMap[keys+":"+action]({},keys)}return self};Mousetrap.prototype.reset=function(){var self=this;self._callbacks={};self._directMap={};return self};Mousetrap.prototype.stopCallback=function(e,element){var self=this;if((" "+element.className+" ").indexOf(" mousetrap ")>-1){return false}if(_belongsTo(element,self.target)){return false}return element.tagName=="INPUT"||element.tagName=="SELECT"||element.tagName=="TEXTAREA"||element.isContentEditable};Mousetrap.prototype.handleKey=function(){var self=this;return self._handleKey.apply(self,arguments)};Mousetrap.init=function(){var documentMousetrap=Mousetrap(document);for(var method in documentMousetrap){if(method.charAt(0)!=="_"){Mousetrap[method]=function(method){return function(){return documentMousetrap[method].apply(documentMousetrap,arguments)}}(method)}}};Mousetrap.init();window.Mousetrap=Mousetrap;if(typeof module!=="undefined"&&module.exports){module.exports=Mousetrap}if(typeof define==="function"&&define.amd){define(function(){return Mousetrap})}})(window,document)},{}],4:[function(require,module,exports){(function(process){function normalizeArray(parts,allowAboveRoot){var up=0;for(var i=parts.length-1;i>=0;i--){var last=parts[i];if(last==="."){parts.splice(i,1)}else if(last===".."){parts.splice(i,1);up++}else if(up){parts.splice(i,1);up--}}if(allowAboveRoot){for(;up--;up){parts.unshift("..")}}return parts}var splitPathRe=/^(\/?|)([\s\S]*?)((?:\.{1,2}|[^\/]+?|)(\.[^.\/]*|))(?:[\/]*)$/;var splitPath=function(filename){return splitPathRe.exec(filename).slice(1)};exports.resolve=function(){var resolvedPath="",resolvedAbsolute=false;for(var i=arguments.length-1;i>=-1&&!resolvedAbsolute;i--){var path=i>=0?arguments[i]:process.cwd();if(typeof path!=="string"){throw new TypeError("Arguments to path.resolve must be strings")}else if(!path){continue}resolvedPath=path+"/"+resolvedPath;resolvedAbsolute=path.charAt(0)==="/"}resolvedPath=normalizeArray(filter(resolvedPath.split("/"),function(p){return!!p}),!resolvedAbsolute).join("/");return(resolvedAbsolute?"/":"")+resolvedPath||"."};exports.normalize=function(path){var isAbsolute=exports.isAbsolute(path),trailingSlash=substr(path,-1)==="/";path=normalizeArray(filter(path.split("/"),function(p){return!!p}),!isAbsolute).join("/");if(!path&&!isAbsolute){path="."}if(path&&trailingSlash){path+="/"}return(isAbsolute?"/":"")+path};exports.isAbsolute=function(path){return path.charAt(0)==="/"};exports.join=function(){var paths=Array.prototype.slice.call(arguments,0);return exports.normalize(filter(paths,function(p,index){if(typeof p!=="string"){throw new TypeError("Arguments to path.join must be strings")}return p}).join("/"))};exports.relative=function(from,to){from=exports.resolve(from).substr(1);to=exports.resolve(to).substr(1);function trim(arr){var start=0;for(;start<arr.length;start++){if(arr[start]!=="")break}var end=arr.length-1;for(;end>=0;end--){if(arr[end]!=="")break}if(start>end)return[];return arr.slice(start,end-start+1)}var fromParts=trim(from.split("/"));var toParts=trim(to.split("/"));var length=Math.min(fromParts.length,toParts.length);var samePartsLength=length;for(var i=0;i<length;i++){if(fromParts[i]!==toParts[i]){samePartsLength=i;break}}var outputParts=[];for(var i=samePartsLength;i<fromParts.length;i++){outputParts.push("..")}outputParts=outputParts.concat(toParts.slice(samePartsLength));return outputParts.join("/")};exports.sep="/";exports.delimiter=":";exports.dirname=function(path){var result=splitPath(path),root=result[0],dir=result[1];if(!root&&!dir){return"."}if(dir){dir=dir.substr(0,dir.length-1)}return root+dir};exports.basename=function(path,ext){var f=splitPath(path)[2];if(ext&&f.substr(-1*ext.length)===ext){f=f.substr(0,f.length-ext.length)}return f};exports.extname=function(path){return splitPath(path)[3]};function filter(xs,f){if(xs.filter)return xs.filter(f);var res=[];for(var i=0;i<xs.length;i++){if(f(xs[i],i,xs))res.push(xs[i])}return res}var substr="ab".substr(-1)==="b"?function(str,start,len){return str.substr(start,len)}:function(str,start,len){if(start<0)start=str.length+start;return str.substr(start,len)}}).call(this,require("_process"))},{_process:5}],5:[function(require,module,exports){var process=module.exports={};var queue=[];var draining=false;var currentQueue;var queueIndex=-1;function cleanUpNextTick(){draining=false;if(currentQueue.length){queue=currentQueue.concat(queue)}else{queueIndex=-1}if(queue.length){drainQueue()}}function drainQueue(){if(draining){return}var timeout=setTimeout(cleanUpNextTick);draining=true;var len=queue.length;while(len){currentQueue=queue;queue=[];while(++queueIndex<len){if(currentQueue){currentQueue[queueIndex].run()}}queueIndex=-1;len=queue.length}currentQueue=null;draining=false;clearTimeout(timeout)}process.nextTick=function(fun){var args=new Array(arguments.length-1);if(arguments.length>1){for(var i=1;i<arguments.length;i++){args[i-1]=arguments[i]}}queue.push(new Item(fun,args));if(queue.length===1&&!draining){setTimeout(drainQueue,0)}};function Item(fun,array){this.fun=fun;this.array=array}Item.prototype.run=function(){this.fun.apply(null,this.array)};process.title="browser";process.browser=true;process.env={};process.argv=[];process.version="";process.versions={};function noop(){}process.on=noop;process.addListener=noop;process.once=noop;process.off=noop;process.removeListener=noop;process.removeAllListeners=noop;process.emit=noop;process.binding=function(name){throw new Error("process.binding is not supported")};process.cwd=function(){return"/"};process.chdir=function(dir){throw new Error("process.chdir is not supported")};process.umask=function(){return 0}},{}],6:[function(require,module,exports){(function(global){(function(root){var freeExports=typeof exports=="object"&&exports&&!exports.nodeType&&exports;var freeModule=typeof module=="object"&&module&&!module.nodeType&&module;var freeGlobal=typeof global=="object"&&global;if(freeGlobal.global===freeGlobal||freeGlobal.window===freeGlobal||freeGlobal.self===freeGlobal){root=freeGlobal}var punycode,maxInt=2147483647,base=36,tMin=1,tMax=26,skew=38,damp=700,initialBias=72,initialN=128,delimiter="-",regexPunycode=/^xn--/,regexNonASCII=/[^\x20-\x7E]/,regexSeparators=/[\x2E\u3002\uFF0E\uFF61]/g,errors={overflow:"Overflow: input needs wider integers to process","not-basic":"Illegal input >= 0x80 (not a basic code point)","invalid-input":"Invalid input"},baseMinusTMin=base-tMin,floor=Math.floor,stringFromCharCode=String.fromCharCode,key;function error(type){throw RangeError(errors[type])}function map(array,fn){var length=array.length;var result=[];while(length--){result[length]=fn(array[length])}return result}function mapDomain(string,fn){var parts=string.split("@");var result="";if(parts.length>1){result=parts[0]+"@";string=parts[1]}string=string.replace(regexSeparators,".");var labels=string.split(".");var encoded=map(labels,fn).join(".");return result+encoded}function ucs2decode(string){var output=[],counter=0,length=string.length,value,extra;while(counter<length){value=string.charCodeAt(counter++);if(value>=55296&&value<=56319&&counter<length){extra=string.charCodeAt(counter++);if((extra&64512)==56320){output.push(((value&1023)<<10)+(extra&1023)+65536)}else{output.push(value);counter--}}else{output.push(value)}}return output}function ucs2encode(array){return map(array,function(value){var output="";if(value>65535){value-=65536;output+=stringFromCharCode(value>>>10&1023|55296);value=56320|value&1023}output+=stringFromCharCode(value);return output}).join("")}function basicToDigit(codePoint){if(codePoint-48<10){return codePoint-22}if(codePoint-65<26){return codePoint-65}if(codePoint-97<26){return codePoint-97}return base}function digitToBasic(digit,flag){return digit+22+75*(digit<26)-((flag!=0)<<5)}function adapt(delta,numPoints,firstTime){var k=0;delta=firstTime?floor(delta/damp):delta>>1;delta+=floor(delta/numPoints);for(;delta>baseMinusTMin*tMax>>1;k+=base){delta=floor(delta/baseMinusTMin)}return floor(k+(baseMinusTMin+1)*delta/(delta+skew))}function decode(input){var output=[],inputLength=input.length,out,i=0,n=initialN,bias=initialBias,basic,j,index,oldi,w,k,digit,t,baseMinusT;basic=input.lastIndexOf(delimiter);if(basic<0){basic=0}for(j=0;j<basic;++j){if(input.charCodeAt(j)>=128){error("not-basic")}output.push(input.charCodeAt(j))}for(index=basic>0?basic+1:0;index<inputLength;){for(oldi=i,w=1,k=base;;k+=base){if(index>=inputLength){error("invalid-input")}digit=basicToDigit(input.charCodeAt(index++));if(digit>=base||digit>floor((maxInt-i)/w)){error("overflow")}i+=digit*w;t=k<=bias?tMin:k>=bias+tMax?tMax:k-bias;if(digit<t){break}baseMinusT=base-t;if(w>floor(maxInt/baseMinusT)){error("overflow")}w*=baseMinusT}out=output.length+1;bias=adapt(i-oldi,out,oldi==0);if(floor(i/out)>maxInt-n){error("overflow")}n+=floor(i/out);i%=out;output.splice(i++,0,n)}return ucs2encode(output)}function encode(input){var n,delta,handledCPCount,basicLength,bias,j,m,q,k,t,currentValue,output=[],inputLength,handledCPCountPlusOne,baseMinusT,qMinusT;input=ucs2decode(input);inputLength=input.length;n=initialN;delta=0;bias=initialBias;for(j=0;j<inputLength;++j){currentValue=input[j];if(currentValue<128){output.push(stringFromCharCode(currentValue))}}handledCPCount=basicLength=output.length;if(basicLength){output.push(delimiter)}while(handledCPCount<inputLength){for(m=maxInt,j=0;j<inputLength;++j){currentValue=input[j];if(currentValue>=n&&currentValue<m){m=currentValue}}handledCPCountPlusOne=handledCPCount+1;if(m-n>floor((maxInt-delta)/handledCPCountPlusOne)){error("overflow")}delta+=(m-n)*handledCPCountPlusOne;n=m;for(j=0;j<inputLength;++j){currentValue=input[j];if(currentValue<n&&++delta>maxInt){error("overflow")}if(currentValue==n){for(q=delta,k=base;;k+=base){t=k<=bias?tMin:k>=bias+tMax?tMax:k-bias;if(q<t){break}qMinusT=q-t;baseMinusT=base-t;output.push(stringFromCharCode(digitToBasic(t+qMinusT%baseMinusT,0)));q=floor(qMinusT/baseMinusT)}output.push(stringFromCharCode(digitToBasic(q,0)));bias=adapt(delta,handledCPCountPlusOne,handledCPCount==basicLength);delta=0;++handledCPCount}}++delta;++n}return output.join("")}function toUnicode(input){return mapDomain(input,function(string){return regexPunycode.test(string)?decode(string.slice(4).toLowerCase()):string})}function toASCII(input){return mapDomain(input,function(string){return regexNonASCII.test(string)?"xn--"+encode(string):string})}punycode={version:"1.3.2",ucs2:{decode:ucs2decode,encode:ucs2encode},decode:decode,encode:encode,toASCII:toASCII,toUnicode:toUnicode};if(typeof define=="function"&&typeof define.amd=="object"&&define.amd){define("punycode",function(){return punycode})}else if(freeExports&&freeModule){if(module.exports==freeExports){freeModule.exports=punycode}else{for(key in punycode){punycode.hasOwnProperty(key)&&(freeExports[key]=punycode[key])}}}else{root.punycode=punycode}})(this)}).call(this,typeof global!=="undefined"?global:typeof self!=="undefined"?self:typeof window!=="undefined"?window:{})},{}],7:[function(require,module,exports){"use strict";function hasOwnProperty(obj,prop){return Object.prototype.hasOwnProperty.call(obj,prop)}module.exports=function(qs,sep,eq,options){sep=sep||"&";eq=eq||"=";var obj={};if(typeof qs!=="string"||qs.length===0){return obj}var regexp=/\+/g;qs=qs.split(sep);var maxKeys=1e3;if(options&&typeof options.maxKeys==="number"){maxKeys=options.maxKeys}var len=qs.length;if(maxKeys>0&&len>maxKeys){len=maxKeys}for(var i=0;i<len;++i){var x=qs[i].replace(regexp,"%20"),idx=x.indexOf(eq),kstr,vstr,k,v;if(idx>=0){kstr=x.substr(0,idx);vstr=x.substr(idx+1)}else{kstr=x;vstr=""}k=decodeURIComponent(kstr);v=decodeURIComponent(vstr);if(!hasOwnProperty(obj,k)){obj[k]=v}else if(isArray(obj[k])){obj[k].push(v)}else{obj[k]=[obj[k],v]}}return obj};var isArray=Array.isArray||function(xs){return Object.prototype.toString.call(xs)==="[object Array]"}},{}],8:[function(require,module,exports){"use strict";var stringifyPrimitive=function(v){switch(typeof v){case"string":return v;case"boolean":return v?"true":"false";case"number":return isFinite(v)?v:"";default:return""}};module.exports=function(obj,sep,eq,name){sep=sep||"&";eq=eq||"=";if(obj===null){obj=undefined}if(typeof obj==="object"){return map(objectKeys(obj),function(k){var ks=encodeURIComponent(stringifyPrimitive(k))+eq;if(isArray(obj[k])){return map(obj[k],function(v){return ks+encodeURIComponent(stringifyPrimitive(v))}).join(sep)}else{return ks+encodeURIComponent(stringifyPrimitive(obj[k]))}}).join(sep)}if(!name)return"";return encodeURIComponent(stringifyPrimitive(name))+eq+encodeURIComponent(stringifyPrimitive(obj))};var isArray=Array.isArray||function(xs){return Object.prototype.toString.call(xs)==="[object Array]"};function map(xs,f){if(xs.map)return xs.map(f);var res=[];for(var i=0;i<xs.length;i++){res.push(f(xs[i],i))}return res}var objectKeys=Object.keys||function(obj){var res=[];for(var key in obj){if(Object.prototype.hasOwnProperty.call(obj,key))res.push(key)}return res}},{}],9:[function(require,module,exports){"use strict";exports.decode=exports.parse=require("./decode");exports.encode=exports.stringify=require("./encode")},{"./decode":7,"./encode":8}],10:[function(require,module,exports){var punycode=require("punycode");exports.parse=urlParse;exports.resolve=urlResolve;exports.resolveObject=urlResolveObject;exports.format=urlFormat;exports.Url=Url;function Url(){this.protocol=null;this.slashes=null;this.auth=null;this.host=null;this.port=null;this.hostname=null;this.hash=null;this.search=null;this.query=null;this.pathname=null;this.path=null;this.href=null}var protocolPattern=/^([a-z0-9.+-]+:)/i,portPattern=/:[0-9]*$/,delims=["<",">",'"',"`"," ","\r","\n","\t"],unwise=["{","}","|","\\","^","`"].concat(delims),autoEscape=["'"].concat(unwise),nonHostChars=["%","/","?",";","#"].concat(autoEscape),hostEndingChars=["/","?","#"],hostnameMaxLen=255,hostnamePartPattern=/^[a-z0-9A-Z_-]{0,63}$/,hostnamePartStart=/^([a-z0-9A-Z_-]{0,63})(.*)$/,unsafeProtocol={javascript:true,"javascript:":true},hostlessProtocol={javascript:true,"javascript:":true},slashedProtocol={http:true,https:true,ftp:true,gopher:true,file:true,"http:":true,"https:":true,"ftp:":true,"gopher:":true,"file:":true},querystring=require("querystring");function urlParse(url,parseQueryString,slashesDenoteHost){if(url&&isObject(url)&&url instanceof Url)return url;var u=new Url;u.parse(url,parseQueryString,slashesDenoteHost);return u}Url.prototype.parse=function(url,parseQueryString,slashesDenoteHost){if(!isString(url)){throw new TypeError("Parameter 'url' must be a string, not "+typeof url)}var rest=url;rest=rest.trim();var proto=protocolPattern.exec(rest);if(proto){proto=proto[0];var lowerProto=proto.toLowerCase();this.protocol=lowerProto;rest=rest.substr(proto.length)}if(slashesDenoteHost||proto||rest.match(/^\/\/[^@\/]+@[^@\/]+/)){var slashes=rest.substr(0,2)==="//";if(slashes&&!(proto&&hostlessProtocol[proto])){rest=rest.substr(2);this.slashes=true}}if(!hostlessProtocol[proto]&&(slashes||proto&&!slashedProtocol[proto])){var hostEnd=-1;for(var i=0;i<hostEndingChars.length;i++){var hec=rest.indexOf(hostEndingChars[i]);if(hec!==-1&&(hostEnd===-1||hec<hostEnd))hostEnd=hec}var auth,atSign;if(hostEnd===-1){atSign=rest.lastIndexOf("@")}else{atSign=rest.lastIndexOf("@",hostEnd)}if(atSign!==-1){auth=rest.slice(0,atSign);rest=rest.slice(atSign+1);this.auth=decodeURIComponent(auth)}hostEnd=-1;for(var i=0;i<nonHostChars.length;i++){var hec=rest.indexOf(nonHostChars[i]);if(hec!==-1&&(hostEnd===-1||hec<hostEnd))hostEnd=hec}if(hostEnd===-1)hostEnd=rest.length;this.host=rest.slice(0,hostEnd);rest=rest.slice(hostEnd);this.parseHost();this.hostname=this.hostname||"";var ipv6Hostname=this.hostname[0]==="["&&this.hostname[this.hostname.length-1]==="]";if(!ipv6Hostname){var hostparts=this.hostname.split(/\./);for(var i=0,l=hostparts.length;i<l;i++){var part=hostparts[i];if(!part)continue;if(!part.match(hostnamePartPattern)){var newpart="";for(var j=0,k=part.length;j<k;j++){if(part.charCodeAt(j)>127){newpart+="x"}else{newpart+=part[j]}}if(!newpart.match(hostnamePartPattern)){var validParts=hostparts.slice(0,i);var notHost=hostparts.slice(i+1);var bit=part.match(hostnamePartStart);if(bit){validParts.push(bit[1]);notHost.unshift(bit[2])}if(notHost.length){rest="/"+notHost.join(".")+rest}this.hostname=validParts.join(".");break}}}}if(this.hostname.length>hostnameMaxLen){this.hostname=""}else{this.hostname=this.hostname.toLowerCase()}if(!ipv6Hostname){var domainArray=this.hostname.split(".");var newOut=[];for(var i=0;i<domainArray.length;++i){var s=domainArray[i];newOut.push(s.match(/[^A-Za-z0-9_-]/)?"xn--"+punycode.encode(s):s)}this.hostname=newOut.join(".")}var p=this.port?":"+this.port:"";var h=this.hostname||"";this.host=h+p;this.href+=this.host;if(ipv6Hostname){this.hostname=this.hostname.substr(1,this.hostname.length-2);if(rest[0]!=="/"){rest="/"+rest}}}if(!unsafeProtocol[lowerProto]){for(var i=0,l=autoEscape.length;i<l;i++){var ae=autoEscape[i];var esc=encodeURIComponent(ae);if(esc===ae){esc=escape(ae)}rest=rest.split(ae).join(esc)}}var hash=rest.indexOf("#");if(hash!==-1){this.hash=rest.substr(hash);rest=rest.slice(0,hash)}var qm=rest.indexOf("?");if(qm!==-1){this.search=rest.substr(qm);this.query=rest.substr(qm+1);if(parseQueryString){this.query=querystring.parse(this.query)}rest=rest.slice(0,qm)}else if(parseQueryString){this.search="";this.query={}}if(rest)this.pathname=rest;if(slashedProtocol[lowerProto]&&this.hostname&&!this.pathname){this.pathname="/"}if(this.pathname||this.search){var p=this.pathname||"";var s=this.search||"";this.path=p+s}this.href=this.format();return this};function urlFormat(obj){if(isString(obj))obj=urlParse(obj);if(!(obj instanceof Url))return Url.prototype.format.call(obj);return obj.format()}Url.prototype.format=function(){var auth=this.auth||"";if(auth){auth=encodeURIComponent(auth);auth=auth.replace(/%3A/i,":");auth+="@"}var protocol=this.protocol||"",pathname=this.pathname||"",hash=this.hash||"",host=false,query="";if(this.host){host=auth+this.host}else if(this.hostname){host=auth+(this.hostname.indexOf(":")===-1?this.hostname:"["+this.hostname+"]");if(this.port){host+=":"+this.port}}if(this.query&&isObject(this.query)&&Object.keys(this.query).length){query=querystring.stringify(this.query)}var search=this.search||query&&"?"+query||"";if(protocol&&protocol.substr(-1)!==":")protocol+=":";if(this.slashes||(!protocol||slashedProtocol[protocol])&&host!==false){host="//"+(host||"");if(pathname&&pathname.charAt(0)!=="/")pathname="/"+pathname}else if(!host){host=""}if(hash&&hash.charAt(0)!=="#")hash="#"+hash;if(search&&search.charAt(0)!=="?")search="?"+search;pathname=pathname.replace(/[?#]/g,function(match){return encodeURIComponent(match)});search=search.replace("#","%23");return protocol+host+pathname+search+hash};function urlResolve(source,relative){return urlParse(source,false,true).resolve(relative)}Url.prototype.resolve=function(relative){return this.resolveObject(urlParse(relative,false,true)).format()};function urlResolveObject(source,relative){if(!source)return relative;return urlParse(source,false,true).resolveObject(relative)}Url.prototype.resolveObject=function(relative){if(isString(relative)){var rel=new Url;rel.parse(relative,false,true);relative=rel}var result=new Url;Object.keys(this).forEach(function(k){result[k]=this[k]},this);result.hash=relative.hash;if(relative.href===""){result.href=result.format();return result}if(relative.slashes&&!relative.protocol){Object.keys(relative).forEach(function(k){if(k!=="protocol")result[k]=relative[k]});if(slashedProtocol[result.protocol]&&result.hostname&&!result.pathname){result.path=result.pathname="/"}result.href=result.format();return result}if(relative.protocol&&relative.protocol!==result.protocol){if(!slashedProtocol[relative.protocol]){Object.keys(relative).forEach(function(k){result[k]=relative[k]});result.href=result.format();return result}result.protocol=relative.protocol;if(!relative.host&&!hostlessProtocol[relative.protocol]){var relPath=(relative.pathname||"").split("/");while(relPath.length&&!(relative.host=relPath.shift()));if(!relative.host)relative.host="";if(!relative.hostname)relative.hostname="";if(relPath[0]!=="")relPath.unshift("");if(relPath.length<2)relPath.unshift("");result.pathname=relPath.join("/")}else{result.pathname=relative.pathname}result.search=relative.search;result.query=relative.query;result.host=relative.host||"";result.auth=relative.auth;result.hostname=relative.hostname||relative.host;result.port=relative.port;if(result.pathname||result.search){var p=result.pathname||"";var s=result.search||"";result.path=p+s}result.slashes=result.slashes||relative.slashes;result.href=result.format();return result}var isSourceAbs=result.pathname&&result.pathname.charAt(0)==="/",isRelAbs=relative.host||relative.pathname&&relative.pathname.charAt(0)==="/",mustEndAbs=isRelAbs||isSourceAbs||result.host&&relative.pathname,removeAllDots=mustEndAbs,srcPath=result.pathname&&result.pathname.split("/")||[],relPath=relative.pathname&&relative.pathname.split("/")||[],psychotic=result.protocol&&!slashedProtocol[result.protocol];if(psychotic){result.hostname="";result.port=null;if(result.host){if(srcPath[0]==="")srcPath[0]=result.host;else srcPath.unshift(result.host)}result.host="";if(relative.protocol){relative.hostname=null;relative.port=null;if(relative.host){if(relPath[0]==="")relPath[0]=relative.host;else relPath.unshift(relative.host)}relative.host=null}mustEndAbs=mustEndAbs&&(relPath[0]===""||srcPath[0]==="")}if(isRelAbs){result.host=relative.host||relative.host===""?relative.host:result.host;result.hostname=relative.hostname||relative.hostname===""?relative.hostname:result.hostname;result.search=relative.search;result.query=relative.query;srcPath=relPath}else if(relPath.length){if(!srcPath)srcPath=[];srcPath.pop();srcPath=srcPath.concat(relPath);result.search=relative.search;result.query=relative.query}else if(!isNullOrUndefined(relative.search)){if(psychotic){result.hostname=result.host=srcPath.shift();var authInHost=result.host&&result.host.indexOf("@")>0?result.host.split("@"):false;if(authInHost){result.auth=authInHost.shift();result.host=result.hostname=authInHost.shift()}}result.search=relative.search;result.query=relative.query;if(!isNull(result.pathname)||!isNull(result.search)){result.path=(result.pathname?result.pathname:"")+(result.search?result.search:"")}result.href=result.format();return result}if(!srcPath.length){result.pathname=null;if(result.search){result.path="/"+result.search}else{result.path=null}result.href=result.format();return result}var last=srcPath.slice(-1)[0];var hasTrailingSlash=(result.host||relative.host)&&(last==="."||last==="..")||last==="";var up=0;for(var i=srcPath.length;i>=0;i--){last=srcPath[i];if(last=="."){srcPath.splice(i,1)}else if(last===".."){srcPath.splice(i,1);up++}else if(up){srcPath.splice(i,1);up--}}if(!mustEndAbs&&!removeAllDots){for(;up--;up){srcPath.unshift("..")}}if(mustEndAbs&&srcPath[0]!==""&&(!srcPath[0]||srcPath[0].charAt(0)!=="/")){srcPath.unshift("")}if(hasTrailingSlash&&srcPath.join("/").substr(-1)!=="/"){srcPath.push("")}var isAbsolute=srcPath[0]===""||srcPath[0]&&srcPath[0].charAt(0)==="/";if(psychotic){result.hostname=result.host=isAbsolute?"":srcPath.length?srcPath.shift():"";var authInHost=result.host&&result.host.indexOf("@")>0?result.host.split("@"):false;if(authInHost){result.auth=authInHost.shift();result.host=result.hostname=authInHost.shift()}}mustEndAbs=mustEndAbs||result.host&&srcPath.length;if(mustEndAbs&&!isAbsolute){srcPath.unshift("")}if(!srcPath.length){result.pathname=null;result.path=null}else{result.pathname=srcPath.join("/")}if(!isNull(result.pathname)||!isNull(result.search)){result.path=(result.pathname?result.pathname:"")+(result.search?result.search:"")}result.auth=relative.auth||result.auth;result.slashes=result.slashes||relative.slashes;result.href=result.format();return result};Url.prototype.parseHost=function(){var host=this.host;var port=portPattern.exec(host);if(port){port=port[0];if(port!==":"){this.port=port.substr(1)}host=host.substr(0,host.length-port.length)}if(host)this.hostname=host};function isString(arg){return typeof arg==="string"}function isObject(arg){return typeof arg==="object"&&arg!==null}function isNull(arg){return arg===null}function isNullOrUndefined(arg){return arg==null}},{punycode:6,querystring:9}],11:[function(require,module,exports){var $=require("jquery");function toggleDropdown(e){var $dropdown=$(e.currentTarget).parent().find(".dropdown-menu");$dropdown.toggleClass("open");e.stopPropagation();e.preventDefault()}function closeDropdown(e){$(".dropdown-menu").removeClass("open")}function init(){$(document).on("click",".toggle-dropdown",toggleDropdown);$(document).on("click",".dropdown-menu",function(e){e.stopPropagation()});$(document).on("click",closeDropdown)}module.exports={init:init}},{jquery:1}],12:[function(require,module,exports){var $=require("jquery");module.exports=$({})},{jquery:1}],13:[function(require,module,exports){var $=require("jquery");var _=require("lodash");var storage=require("./storage");var dropdown=require("./dropdown");var events=require("./events");var state=require("./state");var keyboard=require("./keyboard");var navigation=require("./navigation");var sidebar=require("./sidebar");var toolbar=require("./toolbar");function start(config){sidebar.init();keyboard.init();dropdown.init();navigation.init();toolbar.createButton({index:0,icon:"fa fa-align-justify",label:"Toggle Sidebar",onClick:function(e){e.preventDefault();sidebar.toggle()}});events.trigger("start",config);navigation.notify()}var gitbook={start:start,events:events,state:state,toolbar:toolbar,sidebar:sidebar,storage:storage,keyboard:keyboard};var MODULES={gitbook:gitbook,jquery:$,lodash:_};window.gitbook=gitbook;window.$=$;window.jQuery=$;gitbook.require=function(mods,fn){mods=_.map(mods,function(mod){mod=mod.toLowerCase();if(!MODULES[mod]){throw new Error("GitBook module "+mod+" doesn't exist")}return MODULES[mod]});fn.apply(null,mods)};module.exports={}},{"./dropdown":11,"./events":12,"./keyboard":14,"./navigation":16,"./sidebar":18,"./state":19,"./storage":20,"./toolbar":21,jquery:1,lodash:2}],14:[function(require,module,exports){var Mousetrap=require("mousetrap");var navigation=require("./navigation");var sidebar=require("./sidebar");function bindShortcut(keys,fn){Mousetrap.bind(keys,function(e){fn();return false})}function init(){bindShortcut(["right"],function(e){navigation.goNext()});bindShortcut(["left"],function(e){navigation.goPrev()});bindShortcut(["s"],function(e){sidebar.toggle()})}module.exports={init:init,bind:bindShortcut}},{"./navigation":16,"./sidebar":18,mousetrap:3}],15:[function(require,module,exports){var state=require("./state");function showLoading(p){state.$book.addClass("is-loading");p.always(function(){state.$book.removeClass("is-loading")});return p}module.exports={show:showLoading}},{"./state":19}],16:[function(require,module,exports){var $=require("jquery");var url=require("url");var events=require("./events");var state=require("./state");var loading=require("./loading");var usePushState=typeof history.pushState!=="undefined";function handleNavigation(relativeUrl,push){var uri=url.resolve(window.location.pathname,relativeUrl);notifyPageChange();location.href=relativeUrl;return}function updateNavigationPosition(){var bodyInnerWidth,pageWrapperWidth;bodyInnerWidth=parseInt($(".body-inner").css("width"),10);pageWrapperWidth=parseInt($(".page-wrapper").css("width"),10);$(".navigation-next").css("margin-right",bodyInnerWidth-pageWrapperWidth+"px")}function notifyPageChange(){events.trigger("page.change")}function preparePage(notify){var $bookBody=$(".book-body");var $bookInner=$bookBody.find(".body-inner");var $pageWrapper=$bookInner.find(".page-wrapper");updateNavigationPosition();$bookInner.scrollTop(0);$bookBody.scrollTop(0);if(notify!==false)notifyPageChange()}function isLeftClickEvent(e){return e.button===0}function isModifiedEvent(e){return!!(e.metaKey||e.altKey||e.ctrlKey||e.shiftKey)}function handlePagination(e){if(isModifiedEvent(e)||!isLeftClickEvent(e)){return}e.stopPropagation();e.preventDefault();var url=$(this).attr("href");if(url)handleNavigation(url,true)}function goNext(){var url=$(".navigation-next").attr("href");if(url)handleNavigation(url,true)}function goPrev(){var url=$(".navigation-prev").attr("href");if(url)handleNavigation(url,true)}function init(){$.ajaxSetup({});if(location.protocol!=="file:"){history.replaceState({path:window.location.href},"")}window.onpopstate=function(event){if(event.state===null){return}return handleNavigation(event.state.path,false)};$(document).on("click",".navigation-prev",handlePagination);$(document).on("click",".navigation-next",handlePagination);$(document).on("click",".summary [data-path] a",handlePagination);$(window).resize(updateNavigationPosition);preparePage(false)}module.exports={init:init,goNext:goNext,goPrev:goPrev,notify:notifyPageChange}},{"./events":12,"./loading":15,"./state":19,jquery:1,url:10}],17:[function(require,module,exports){module.exports={isMobile:function(){return document.body.clientWidth<=600}}},{}],18:[function(require,module,exports){var $=require("jquery");var _=require("lodash");var storage=require("./storage");var platform=require("./platform");var state=require("./state");function toggleSidebar(_state,animation){if(state!=null&&isOpen()==_state)return;if(animation==null)animation=true;state.$book.toggleClass("without-animation",!animation);state.$book.toggleClass("with-summary",_state);storage.set("sidebar",isOpen())}function isOpen(){return state.$book.hasClass("with-summary")}function init(){if(platform.isMobile()){toggleSidebar(false,false)}else{toggleSidebar(storage.get("sidebar",true),false)}$(document).on("click",".book-summary li.chapter a",function(e){if(platform.isMobile())toggleSidebar(false,false)})}function filterSummary(paths){var $summary=$(".book-summary");$summary.find("li").each(function(){var path=$(this).data("path");var st=paths==null||_.contains(paths,path);$(this).toggle(st);if(st)$(this).parents("li").show()})}module.exports={init:init,isOpen:isOpen,toggle:toggleSidebar,filter:filterSummary}},{"./platform":17,"./state":19,"./storage":20,jquery:1,lodash:2}],19:[function(require,module,exports){var $=require("jquery");var url=require("url");var path=require("path");var state={};state.update=function(dom){var $book=$(dom.find(".book"));state.$book=$book;state.level=$book.data("level");state.basePath=$book.data("basepath");state.innerLanguage=$book.data("innerlanguage");state.revision=$book.data("revision");state.filepath=$book.data("filepath");state.chapterTitle=$book.data("chapter-title");state.root=url.resolve(location.protocol+"//"+location.host,path.dirname(path.resolve(location.pathname.replace(/\/$/,"/index.html"),state.basePath))).replace(/\/?$/,"/");state.bookRoot=state.innerLanguage?url.resolve(state.root,".."):state.root};state.update($);module.exports=state},{jquery:1,path:4,url:10}],20:[function(require,module,exports){var baseKey="";module.exports={setBaseKey:function(key){baseKey=key},set:function(key,value){key=baseKey+":"+key;try{sessionStorage[key]=JSON.stringify(value)}catch(e){}},get:function(key,def){key=baseKey+":"+key;if(sessionStorage[key]===undefined)return def;try{var v=JSON.parse(sessionStorage[key]);return v==null?def:v}catch(err){return sessionStorage[key]||def}},remove:function(key){key=baseKey+":"+key;sessionStorage.removeItem(key)}}},{}],21:[function(require,module,exports){var $=require("jquery");var _=require("lodash");var events=require("./events");var buttons=[];function insertAt(parent,selector,index,element){var lastIndex=parent.children(selector).length;if(index<0){index=Math.max(0,lastIndex+1+index)}parent.append(element);if(index<lastIndex){parent.children(selector).eq(index).before(parent.children(selector).last())}}function defaultOnClick(e){e.preventDefault()}function createDropdownMenu(dropdown){var $menu=$("<div>",{class:"dropdown-menu",html:'<div class="dropdown-caret"><span class="caret-outer"></span><span class="caret-inner"></span></div>'});if(_.isString(dropdown)){$menu.append(dropdown)}else{var groups=_.map(dropdown,function(group){if(_.isArray(group))return group;else return[group]});_.each(groups,function(group){var $group=$("<div>",{class:"buttons"});var sizeClass="size-"+group.length;_.each(group,function(btn){btn=_.defaults(btn||{},{text:"",className:"",onClick:defaultOnClick});var $btn=$("<button>",{class:"button "+sizeClass+" "+btn.className,text:btn.text});$btn.click(btn.onClick);$group.append($btn)});$menu.append($group)})}return $menu}function createButton(opts){opts=_.defaults(opts||{},{label:"",icon:"",text:"",position:"left",className:"",onClick:defaultOnClick,dropdown:null,index:null});buttons.push(opts);updateButton(opts)}function updateButton(opts){var $result;var $toolbar=$(".book-header");var $title=$toolbar.find("h1");var positionClass="pull-"+opts.position;var $btn=$("<a>",{class:"btn",text:opts.text?" "+opts.text:"","aria-label":opts.label,title:opts.label,href:"#"});$btn.click(opts.onClick);if(opts.icon){$("<i>",{class:opts.icon}).prependTo($btn)}if(opts.dropdown){var $container=$("<div>",{class:"dropdown "+positionClass+" "+opts.className});$btn.addClass("toggle-dropdown");$container.append($btn);var $menu=createDropdownMenu(opts.dropdown);$menu.addClass("dropdown-"+(opts.position=="right"?"left":"right"));$container.append($menu);$result=$container}else{$btn.addClass(positionClass);$btn.addClass(opts.className);$result=$btn}$result.addClass("js-toolbar-action");if(_.isNumber(opts.index)&&opts.index>=0){insertAt($toolbar,".btn, .dropdown, h1",opts.index,$result)}else{$result.insertBefore($title)}}module.exports={createButton:createButton}},{"./events":12,jquery:1,lodash:2}]},{},[13]);
```
