## Supplementary material for "dsSurvival: Privacy preserving survival models for federated individual patient meta-analysis in DataSHIELD": Bookdown and documentation for using dsSurvival: clipboard.min.js

```
/*!
 * clipboard.js v2.0.4
 * https://zenorocha.github.io/clipboard.js
 *
 * Licensed MIT © Zeno Rocha
 */
!function(t,e){"object"==typeof exports&&"object"==typeof module?module.exports=e():"function"==typeof define&&define.amd?define([],e):"object"==typeof exports?exports.ClipboardJS=e():t.ClipboardJS=e()}(this,function(){return function(n){var o={};function r(t){if(o[t])return o[t].exports;var e=o[t]={i:t,l:!1,exports:{}};return n[t].call(e.exports,e,e.exports,r),e.l=!0,e.exports}return r.m=n,r.c=o,r.d=function(t,e,n){r.o(t,e)||Object.defineProperty(t,e,{enumerable:!0,get:n})},r.r=function(t){"undefined"!=typeof Symbol&&Symbol.toStringTag&&Object.defineProperty(t,Symbol.toStringTag,{value:"Module"}),Object.defineProperty(t,"__esModule",{value:!0})},r.t=function(e,t){if(1&t&&(e=r(e)),8&t)return e;if(4&t&&"object"==typeof e&&e&&e.__esModule)return e;var n=Object.create(null);if(r.r(n),Object.defineProperty(n,"default",{enumerable:!0,value:e}),2&t&&"string"!=typeof e)for(var o in e)r.d(n,o,function(t){return e[t]}.bind(null,o));return n},r.n=function(t){var e=t&&t.__esModule?function(){return t.default}:function(){return t};return r.d(e,"a",e),e},r.o=function(t,e){return Object.prototype.hasOwnProperty.call(t,e)},r.p="",r(r.s=0)}([function(t,e,n){"use strict";var r="function"==typeof Symbol&&"symbol"==typeof Symbol.iterator?function(t){return typeof t}:function(t){return t&&"function"==typeof Symbol&&t.constructor===Symbol&&t!==Symbol.prototype?"symbol":typeof t},i=function(){function o(t,e){for(var n=0;n<e.length;n++){var o=e[n];o.enumerable=o.enumerable||!1,o.configurable=!0,"value"in o&&(o.writable=!0),Object.defineProperty(t,o.key,o)}}return function(t,e,n){return e&&o(t.prototype,e),n&&o(t,n),t}}(),a=o(n(1)),c=o(n(3)),u=o(n(4));function o(t){return t&&t.__esModule?t:{default:t}}var l=function(t){function o(t,e){!function(t,e){if(!(t instanceof e))throw new TypeError("Cannot call a class as a function")}(this,o);var n=function(t,e){if(!t)throw new ReferenceError("this hasn't been initialised - super() hasn't been called");return!e||"object"!=typeof e&&"function"!=typeof e?t:e}(this,(o.__proto__||Object.getPrototypeOf(o)).call(this));return n.resolveOptions(e),n.listenClick(t),n}return function(t,e){if("function"!=typeof e&&null!==e)throw new TypeError("Super expression must either be null or a function, not "+typeof e);t.prototype=Object.create(e&&e.prototype,{constructor:{value:t,enumerable:!1,writable:!0,configurable:!0}}),e&&(Object.setPrototypeOf?Object.setPrototypeOf(t,e):t.__proto__=e)}(o,c.default),i(o,[{key:"resolveOptions",value:function(){var t=0<arguments.length&&void 0!==arguments[0]?arguments[0]:{};this.action="function"==typeof t.action?t.action:this.defaultAction,this.target="function"==typeof t.target?t.target:this.defaultTarget,this.text="function"==typeof t.text?t.text:this.defaultText,this.container="object"===r(t.container)?t.container:document.body}},{key:"listenClick",value:function(t){var e=this;this.listener=(0,u.default)(t,"click",function(t){return e.onClick(t)})}},{key:"onClick",value:function(t){var e=t.delegateTarget||t.currentTarget;this.clipboardAction&&(this.clipboardAction=null),this.clipboardAction=new a.default({action:this.action(e),target:this.target(e),text:this.text(e),container:this.container,trigger:e,emitter:this})}},{key:"defaultAction",value:function(t){return s("action",t)}},{key:"defaultTarget",value:function(t){var e=s("target",t);if(e)return document.querySelector(e)}},{key:"defaultText",value:function(t){return s("text",t)}},{key:"destroy",value:function(){this.listener.destroy(),this.clipboardAction&&(this.clipboardAction.destroy(),this.clipboardAction=null)}}],[{key:"isSupported",value:function(){var t=0<arguments.length&&void 0!==arguments[0]?arguments[0]:["copy","cut"],e="string"==typeof t?[t]:t,n=!!document.queryCommandSupported;return e.forEach(function(t){n=n&&!!document.queryCommandSupported(t)}),n}}]),o}();function s(t,e){var n="data-clipboard-"+t;if(e.hasAttribute(n))return e.getAttribute(n)}t.exports=l},function(t,e,n){"use strict";var o,r="function"==typeof Symbol&&"symbol"==typeof Symbol.iterator?function(t){return typeof t}:function(t){return t&&"function"==typeof Symbol&&t.constructor===Symbol&&t!==Symbol.prototype?"symbol":typeof t},i=function(){function o(t,e){for(var n=0;n<e.length;n++){var o=e[n];o.enumerable=o.enumerable||!1,o.configurable=!0,"value"in o&&(o.writable=!0),Object.defineProperty(t,o.key,o)}}return function(t,e,n){return e&&o(t.prototype,e),n&&o(t,n),t}}(),a=n(2),c=(o=a)&&o.__esModule?o:{default:o};var u=function(){function e(t){!function(t,e){if(!(t instanceof e))throw new TypeError("Cannot call a class as a function")}(this,e),this.resolveOptions(t),this.initSelection()}return i(e,[{key:"resolveOptions",value:function(){var t=0<arguments.length&&void 0!==arguments[0]?arguments[0]:{};this.action=t.action,this.container=t.container,this.emitter=t.emitter,this.target=t.target,this.text=t.text,this.trigger=t.trigger,this.selectedText=""}},{key:"initSelection",value:function(){this.text?this.selectFake():this.target&&this.selectTarget()}},{key:"selectFake",value:function(){var t=this,e="rtl"==document.documentElement.getAttribute("dir");this.removeFake(),this.fakeHandlerCallback=function(){return t.removeFake()},this.fakeHandler=this.container.addEventListener("click",this.fakeHandlerCallback)||!0,this.fakeElem=document.createElement("textarea"),this.fakeElem.style.fontSize="12pt",this.fakeElem.style.border="0",this.fakeElem.style.padding="0",this.fakeElem.style.margin="0",this.fakeElem.style.position="absolute",this.fakeElem.style[e?"right":"left"]="-9999px";var n=window.pageYOffset||document.documentElement.scrollTop;this.fakeElem.style.top=n+"px",this.fakeElem.setAttribute("readonly",""),this.fakeElem.value=this.text,this.container.appendChild(this.fakeElem),this.selectedText=(0,c.default)(this.fakeElem),this.copyText()}},{key:"removeFake",value:function(){this.fakeHandler&&(this.container.removeEventListener("click",this.fakeHandlerCallback),this.fakeHandler=null,this.fakeHandlerCallback=null),this.fakeElem&&(this.container.removeChild(this.fakeElem),this.fakeElem=null)}},{key:"selectTarget",value:function(){this.selectedText=(0,c.default)(this.target),this.copyText()}},{key:"copyText",value:function(){var e=void 0;try{e=document.execCommand(this.action)}catch(t){e=!1}this.handleResult(e)}},{key:"handleResult",value:function(t){this.emitter.emit(t?"success":"error",{action:this.action,text:this.selectedText,trigger:this.trigger,clearSelection:this.clearSelection.bind(this)})}},{key:"clearSelection",value:function(){this.trigger&&this.trigger.focus(),window.getSelection().removeAllRanges()}},{key:"destroy",value:function(){this.removeFake()}},{key:"action",set:function(){var t=0<arguments.length&&void 0!==arguments[0]?arguments[0]:"copy";if(this._action=t,"copy"!==this._action&&"cut"!==this._action)throw new Error('Invalid "action" value, use either "copy" or "cut"')},get:function(){return this._action}},{key:"target",set:function(t){if(void 0!==t){if(!t||"object"!==(void 0===t?"undefined":r(t))||1!==t.nodeType)throw new Error('Invalid "target" value, use a valid Element');if("copy"===this.action&&t.hasAttribute("disabled"))throw new Error('Invalid "target" attribute. Please use "readonly" instead of "disabled" attribute');if("cut"===this.action&&(t.hasAttribute("readonly")||t.hasAttribute("disabled")))throw new Error('Invalid "target" attribute. You can\'t cut text from elements with "readonly" or "disabled" attributes');this._target=t}},get:function(){return this._target}}]),e}();t.exports=u},function(t,e){t.exports=function(t){var e;if("SELECT"===t.nodeName)t.focus(),e=t.value;else if("INPUT"===t.nodeName||"TEXTAREA"===t.nodeName){var n=t.hasAttribute("readonly");n||t.setAttribute("readonly",""),t.select(),t.setSelectionRange(0,t.value.length),n||t.removeAttribute("readonly"),e=t.value}else{t.hasAttribute("contenteditable")&&t.focus();var o=window.getSelection(),r=document.createRange();r.selectNodeContents(t),o.removeAllRanges(),o.addRange(r),e=o.toString()}return e}},function(t,e){function n(){}n.prototype={on:function(t,e,n){var o=this.e||(this.e={});return(o[t]||(o[t]=[])).push({fn:e,ctx:n}),this},once:function(t,e,n){var o=this;function r(){o.off(t,r),e.apply(n,arguments)}return r._=e,this.on(t,r,n)},emit:function(t){for(var e=[].slice.call(arguments,1),n=((this.e||(this.e={}))[t]||[]).slice(),o=0,r=n.length;o<r;o++)n[o].fn.apply(n[o].ctx,e);return this},off:function(t,e){var n=this.e||(this.e={}),o=n[t],r=[];if(o&&e)for(var i=0,a=o.length;i<a;i++)o[i].fn!==e&&o[i].fn._!==e&&r.push(o[i]);return r.length?n[t]=r:delete n[t],this}},t.exports=n},function(t,e,n){var d=n(5),h=n(6);t.exports=function(t,e,n){if(!t&&!e&&!n)throw new Error("Missing required arguments");if(!d.string(e))throw new TypeError("Second argument must be a String");if(!d.fn(n))throw new TypeError("Third argument must be a Function");if(d.node(t))return s=e,f=n,(l=t).addEventListener(s,f),{destroy:function(){l.removeEventListener(s,f)}};if(d.nodeList(t))return a=t,c=e,u=n,Array.prototype.forEach.call(a,function(t){t.addEventListener(c,u)}),{destroy:function(){Array.prototype.forEach.call(a,function(t){t.removeEventListener(c,u)})}};if(d.string(t))return o=t,r=e,i=n,h(document.body,o,r,i);throw new TypeError("First argument must be a String, HTMLElement, HTMLCollection, or NodeList");var o,r,i,a,c,u,l,s,f}},function(t,n){n.node=function(t){return void 0!==t&&t instanceof HTMLElement&&1===t.nodeType},n.nodeList=function(t){var e=Object.prototype.toString.call(t);return void 0!==t&&("[object NodeList]"===e||"[object HTMLCollection]"===e)&&"length"in t&&(0===t.length||n.node(t[0]))},n.string=function(t){return"string"==typeof t||t instanceof String},n.fn=function(t){return"[object Function]"===Object.prototype.toString.call(t)}},function(t,e,n){var a=n(7);function i(t,e,n,o,r){var i=function(e,n,t,o){return function(t){t.delegateTarget=a(t.target,n),t.delegateTarget&&o.call(e,t)}}.apply(this,arguments);return t.addEventListener(n,i,r),{destroy:function(){t.removeEventListener(n,i,r)}}}t.exports=function(t,e,n,o,r){return"function"==typeof t.addEventListener?i.apply(null,arguments):"function"==typeof n?i.bind(null,document).apply(null,arguments):("string"==typeof t&&(t=document.querySelectorAll(t)),Array.prototype.map.call(t,function(t){return i(t,e,n,o,r)}))}},function(t,e){if("undefined"!=typeof Element&&!Element.prototype.matches){var n=Element.prototype;n.matches=n.matchesSelector||n.mozMatchesSelector||n.msMatchesSelector||n.oMatchesSelector||n.webkitMatchesSelector}t.exports=function(t,e){for(;t&&9!==t.nodeType;){if("function"==typeof t.matches&&t.matches(e))return t;t=t.parentNode}}}])});
```
