## Supplementary material for "dsSurvival: Privacy preserving survival models for federated individual patient meta-analysis in DataSHIELD": Bookdown and documentation for using dsSurvival: jquery.highlight.js

```
gitbook.require(["jQuery"], function(jQuery) {

/*
 * jQuery Highlight plugin
 *
 * Based on highlight v3 by Johann Burkard
 * http://johannburkard.de/blog/programming/javascript/highlight-javascript-text-higlighting-jquery-plugin.html
 *
 * Code a little bit refactored and cleaned (in my humble opinion).
 * Most important changes:
 *  - has an option to highlight only entire words (wordsOnly - false by default),
 *  - has an option to be case sensitive (caseSensitive - false by default)
 *  - highlight element tag and class names can be specified in options
 *
 * Copyright (c) 2009 Bartek Szopka
 *
 * Licensed under MIT license.
 *
 */

jQuery.extend({
    highlight: function (node, re, nodeName, className) {
        if (node.nodeType === 3) {
            var match = node.data.match(re);
            if (match) {
                var highlight = document.createElement(nodeName || 'span');
                highlight.className = className || 'highlight';
                var wordNode = node.splitText(match.index);
                wordNode.splitText(match[0].length);
                var wordClone = wordNode.cloneNode(true);
                highlight.appendChild(wordClone);
                wordNode.parentNode.replaceChild(highlight, wordNode);
                return 1; //skip added node in parent
            }
        } else if ((node.nodeType === 1 && node.childNodes) && // only element nodes that have children
                !/(script|style)/i.test(node.tagName) && // ignore script and style nodes
                !(node.tagName === nodeName.toUpperCase() && node.className === className)) { // skip if already highlighted
            for (var i = 0; i < node.childNodes.length; i++) {
                i += jQuery.highlight(node.childNodes[i], re, nodeName, className);
            }
        }
        return 0;
    }
});

jQuery.fn.unhighlight = function (options) {
    var settings = { className: 'highlight', element: 'span' };
    jQuery.extend(settings, options);

    return this.find(settings.element + "." + settings.className).each(function () {
        var parent = this.parentNode;
        parent.replaceChild(this.firstChild, this);
        parent.normalize();
    }).end();
};

jQuery.fn.highlight = function (words, options) {
    var settings = { className: 'highlight', element: 'span', caseSensitive: false, wordsOnly: false };
    jQuery.extend(settings, options);

    if (words.constructor === String) {
        words = [words];
        // also match 'foo-bar' if search for 'foo bar'
        if (/\s/.test(words[0])) words.push(words[0].replace(/\s+/, '-'));
    }
    words = jQuery.grep(words, function(word, i){
      return word !== '';
    });
    words = jQuery.map(words, function(word, i) {
      return word.replace(/[-[\]{}()*+?.,\\^$|#\s]/g, "\\$&");
    });
    if (words.length === 0) { return this; }

    var flag = settings.caseSensitive ? "" : "i";
    var pattern = "(" + words.join("|") + ")";
    if (settings.wordsOnly) {
        pattern = "\\b" + pattern + "\\b";
    }
    var re = new RegExp(pattern, flag);

    return this.each(function () {
        jQuery.highlight(this, re, settings.element, settings.className);
    });
};

});
```
