## Supplementary material for "dsSurvival: Privacy preserving survival models for federated individual patient meta-analysis in DataSHIELD": Bookdown and documentation for using dsSurvival: lunr.js

```
/**
 * lunr - http://lunrjs.com - A bit like Solr, but much smaller and not as bright - 0.5.12
 * Copyright (C) 2015 Oliver Nightingale
 * MIT Licensed
 * @license
 */
!function(){var t=function(e){var n=new t.Index;return n.pipeline.add(t.trimmer,t.stopWordFilter,t.stemmer),e&&e.call(n,n),n};t.version="0.5.12",t.utils={},t.utils.warn=function(t){return function(e){t.console&&console.warn&&console.warn(e)}}(this),t.EventEmitter=function(){this.events={}},t.EventEmitter.prototype.addListener=function(){var t=Array.prototype.slice.call(arguments),e=t.pop(),n=t;if("function"!=typeof e)throw new TypeError("last argument must be a function");n.forEach(function(t){this.hasHandler(t)||(this.events[t]=[]),this.events[t].push(e)},this)},t.EventEmitter.prototype.removeListener=function(t,e){if(this.hasHandler(t)){var n=this.events[t].indexOf(e);this.events[t].splice(n,1),this.events[t].length||delete this.events[t]}},t.EventEmitter.prototype.emit=function(t){if(this.hasHandler(t)){var e=Array.prototype.slice.call(arguments,1);this.events[t].forEach(function(t){t.apply(void 0,e)})}},t.EventEmitter.prototype.hasHandler=function(t){return t in this.events},t.tokenizer=function(t){return arguments.length&&null!=t&&void 0!=t?Array.isArray(t)?t.map(function(t){return t.toLowerCase()}):t.toString().trim().toLowerCase().split(/[\s\-\/]+/):[]},t.Pipeline=function(){this._stack=[]},t.Pipeline.registeredFunctions={},t.Pipeline.registerFunction=function(e,n){n in this.registeredFunctions&&t.utils.warn("Overwriting existing registered function: "+n),e.label=n,t.Pipeline.registeredFunctions[e.label]=e},t.Pipeline.warnIfFunctionNotRegistered=function(e){var n=e.label&&e.label in this.registeredFunctions;n||t.utils.warn("Function is not registered with pipeline. This may cause problems when serialising the index.\n",e)},t.Pipeline.load=function(e){var n=new t.Pipeline;return e.forEach(function(e){var i=t.Pipeline.registeredFunctions[e];if(!i)throw new Error("Cannot load un-registered function: "+e);n.add(i)}),n},t.Pipeline.prototype.add=function(){var e=Array.prototype.slice.call(arguments);e.forEach(function(e){t.Pipeline.warnIfFunctionNotRegistered(e),this._stack.push(e)},this)},t.Pipeline.prototype.after=function(e,n){t.Pipeline.warnIfFunctionNotRegistered(n);var i=this._stack.indexOf(e);if(-1==i)throw new Error("Cannot find existingFn");i+=1,this._stack.splice(i,0,n)},t.Pipeline.prototype.before=function(e,n){t.Pipeline.warnIfFunctionNotRegistered(n);var i=this._stack.indexOf(e);if(-1==i)throw new Error("Cannot find existingFn");this._stack.splice(i,0,n)},t.Pipeline.prototype.remove=function(t){var e=this._stack.indexOf(t);-1!=e&&this._stack.splice(e,1)},t.Pipeline.prototype.run=function(t){for(var e=[],n=t.length,i=this._stack.length,o=0;n>o;o++){for(var r=t[o],s=0;i>s&&(r=this._stack[s](r,o,t),void 0!==r);s++);void 0!==r&&e.push(r)}return e},t.Pipeline.prototype.reset=function(){this._stack=[]},t.Pipeline.prototype.toJSON=function(){return this._stack.map(function(e){return t.Pipeline.warnIfFunctionNotRegistered(e),e.label})},t.Vector=function(){this._magnitude=null,this.list=void 0,this.length=0},t.Vector.Node=function(t,e,n){this.idx=t,this.val=e,this.next=n},t.Vector.prototype.insert=function(e,n){this._magnitude=void 0;var i=this.list;if(!i)return this.list=new t.Vector.Node(e,n,i),this.length++;if(e<i.idx)return this.list=new t.Vector.Node(e,n,i),this.length++;for(var o=i,r=i.next;void 0!=r;){if(e<r.idx)return o.next=new t.Vector.Node(e,n,r),this.length++;o=r,r=r.next}return o.next=new t.Vector.Node(e,n,r),this.length++},t.Vector.prototype.magnitude=function(){if(this._magnitude)return this._magnitude;for(var t,e=this.list,n=0;e;)t=e.val,n+=t*t,e=e.next;return this._magnitude=Math.sqrt(n)},t.Vector.prototype.dot=function(t){for(var e=this.list,n=t.list,i=0;e&&n;)e.idx<n.idx?e=e.next:e.idx>n.idx?n=n.next:(i+=e.val*n.val,e=e.next,n=n.next);return i},t.Vector.prototype.similarity=function(t){return this.dot(t)/(this.magnitude()*t.magnitude())},t.SortedSet=function(){this.length=0,this.elements=[]},t.SortedSet.load=function(t){var e=new this;return e.elements=t,e.length=t.length,e},t.SortedSet.prototype.add=function(){var t,e;for(t=0;t<arguments.length;t++)e=arguments[t],~this.indexOf(e)||this.elements.splice(this.locationFor(e),0,e);this.length=this.elements.length},t.SortedSet.prototype.toArray=function(){return this.elements.slice()},t.SortedSet.prototype.map=function(t,e){return this.elements.map(t,e)},t.SortedSet.prototype.forEach=function(t,e){return this.elements.forEach(t,e)},t.SortedSet.prototype.indexOf=function(t){for(var e=0,n=this.elements.length,i=n-e,o=e+Math.floor(i/2),r=this.elements[o];i>1;){if(r===t)return o;t>r&&(e=o),r>t&&(n=o),i=n-e,o=e+Math.floor(i/2),r=this.elements[o]}return r===t?o:-1},t.SortedSet.prototype.locationFor=function(t){for(var e=0,n=this.elements.length,i=n-e,o=e+Math.floor(i/2),r=this.elements[o];i>1;)t>r&&(e=o),r>t&&(n=o),i=n-e,o=e+Math.floor(i/2),r=this.elements[o];return r>t?o:t>r?o+1:void 0},t.SortedSet.prototype.intersect=function(e){for(var n=new t.SortedSet,i=0,o=0,r=this.length,s=e.length,a=this.elements,h=e.elements;;){if(i>r-1||o>s-1)break;a[i]!==h[o]?a[i]<h[o]?i++:a[i]>h[o]&&o++:(n.add(a[i]),i++,o++)}return n},t.SortedSet.prototype.clone=function(){var e=new t.SortedSet;return e.elements=this.toArray(),e.length=e.elements.length,e},t.SortedSet.prototype.union=function(t){var e,n,i;return this.length>=t.length?(e=this,n=t):(e=t,n=this),i=e.clone(),i.add.apply(i,n.toArray()),i},t.SortedSet.prototype.toJSON=function(){return this.toArray()},t.Index=function(){this._fields=[],this._ref="id",this.pipeline=new t.Pipeline,this.documentStore=new t.Store,this.tokenStore=new t.TokenStore,this.corpusTokens=new t.SortedSet,this.eventEmitter=new t.EventEmitter,this._idfCache={},this.on("add","remove","update",function(){this._idfCache={}}.bind(this))},t.Index.prototype.on=function(){var t=Array.prototype.slice.call(arguments);return this.eventEmitter.addListener.apply(this.eventEmitter,t)},t.Index.prototype.off=function(t,e){return this.eventEmitter.removeListener(t,e)},t.Index.load=function(e){e.version!==t.version&&t.utils.warn("version mismatch: current "+t.version+" importing "+e.version);var n=new this;return n._fields=e.fields,n._ref=e.ref,n.documentStore=t.Store.load(e.documentStore),n.tokenStore=t.TokenStore.load(e.tokenStore),n.corpusTokens=t.SortedSet.load(e.corpusTokens),n.pipeline=t.Pipeline.load(e.pipeline),n},t.Index.prototype.field=function(t,e){var e=e||{},n={name:t,boost:e.boost||1};return this._fields.push(n),this},t.Index.prototype.ref=function(t){return this._ref=t,this},t.Index.prototype.add=function(e,n){var i={},o=new t.SortedSet,r=e[this._ref],n=void 0===n?!0:n;this._fields.forEach(function(n){var r=this.pipeline.run(t.tokenizer(e[n.name]));i[n.name]=r,t.SortedSet.prototype.add.apply(o,r)},this),this.documentStore.set(r,o),t.SortedSet.prototype.add.apply(this.corpusTokens,o.toArray());for(var s=0;s<o.length;s++){var a=o.elements[s],h=this._fields.reduce(function(t,e){var n=i[e.name].length;if(!n)return t;var o=i[e.name].filter(function(t){return t===a}).length;return t+o/n*e.boost},0);this.tokenStore.add(a,{ref:r,tf:h})}n&&this.eventEmitter.emit("add",e,this)},t.Index.prototype.remove=function(t,e){var n=t[this._ref],e=void 0===e?!0:e;if(this.documentStore.has(n)){var i=this.documentStore.get(n);this.documentStore.remove(n),i.forEach(function(t){this.tokenStore.remove(t,n)},this),e&&this.eventEmitter.emit("remove",t,this)}},t.Index.prototype.update=function(t,e){var e=void 0===e?!0:e;this.remove(t,!1),this.add(t,!1),e&&this.eventEmitter.emit("update",t,this)},t.Index.prototype.idf=function(t){var e="@"+t;if(Object.prototype.hasOwnProperty.call(this._idfCache,e))return this._idfCache[e];var n=this.tokenStore.count(t),i=1;return n>0&&(i=1+Math.log(this.documentStore.length/n)),this._idfCache[e]=i},t.Index.prototype.search=function(e){var n=this.pipeline.run(t.tokenizer(e)),i=new t.Vector,o=[],r=this._fields.reduce(function(t,e){return t+e.boost},0),s=n.some(function(t){return this.tokenStore.has(t)},this);if(!s)return[];n.forEach(function(e,n,s){var a=1/s.length*this._fields.length*r,h=this,l=this.tokenStore.expand(e).reduce(function(n,o){var r=h.corpusTokens.indexOf(o),s=h.idf(o),l=1,u=new t.SortedSet;if(o!==e){var c=Math.max(3,o.length-e.length);l=1/Math.log(c)}return r>-1&&i.insert(r,a*s*l),Object.keys(h.tokenStore.get(o)).forEach(function(t){u.add(t)}),n.union(u)},new t.SortedSet);o.push(l)},this);var a=o.reduce(function(t,e){return t.intersect(e)});return a.map(function(t){return{ref:t,score:i.similarity(this.documentVector(t))}},this).sort(function(t,e){return e.score-t.score})},t.Index.prototype.documentVector=function(e){for(var n=this.documentStore.get(e),i=n.length,o=new t.Vector,r=0;i>r;r++){var s=n.elements[r],a=this.tokenStore.get(s)[e].tf,h=this.idf(s);o.insert(this.corpusTokens.indexOf(s),a*h)}return o},t.Index.prototype.toJSON=function(){return{version:t.version,fields:this._fields,ref:this._ref,documentStore:this.documentStore.toJSON(),tokenStore:this.tokenStore.toJSON(),corpusTokens:this.corpusTokens.toJSON(),pipeline:this.pipeline.toJSON()}},t.Index.prototype.use=function(t){var e=Array.prototype.slice.call(arguments,1);e.unshift(this),t.apply(this,e)},t.Store=function(){this.store={},this.length=0},t.Store.load=function(e){var n=new this;return n.length=e.length,n.store=Object.keys(e.store).reduce(function(n,i){return n[i]=t.SortedSet.load(e.store[i]),n},{}),n},t.Store.prototype.set=function(t,e){this.has(t)||this.length++,this.store[t]=e},t.Store.prototype.get=function(t){return this.store[t]},t.Store.prototype.has=function(t){return t in this.store},t.Store.prototype.remove=function(t){this.has(t)&&(delete this.store[t],this.length--)},t.Store.prototype.toJSON=function(){return{store:this.store,length:this.length}},t.stemmer=function(){var t={ational:"ate",tional:"tion",enci:"ence",anci:"ance",izer:"ize",bli:"ble",alli:"al",entli:"ent",eli:"e",ousli:"ous",ization:"ize",ation:"ate",ator:"ate",alism:"al",iveness:"ive",fulness:"ful",ousness:"ous",aliti:"al",iviti:"ive",biliti:"ble",logi:"log"},e={icate:"ic",ative:"",alize:"al",iciti:"ic",ical:"ic",ful:"",ness:""},n="[^aeiou]",i="[aeiouy]",o=n+"[^aeiouy]*",r=i+"[aeiou]*",s="^("+o+")?"+r+o,a="^("+o+")?"+r+o+"("+r+")?$",h="^("+o+")?"+r+o+r+o,l="^("+o+")?"+i,u=new RegExp(s),c=new RegExp(h),f=new RegExp(a),d=new RegExp(l),p=/^(.+?)(ss|i)es$/,m=/^(.+?)([^s])s$/,v=/^(.+?)eed$/,y=/^(.+?)(ed|ing)$/,g=/.$/,S=/(at|bl|iz)$/,w=new RegExp("([^aeiouylsz])\\1$"),x=new RegExp("^"+o+i+"[^aeiouwxy]$"),k=/^(.+?[^aeiou])y$/,b=/^(.+?)(ational|tional|enci|anci|izer|bli|alli|entli|eli|ousli|ization|ation|ator|alism|iveness|fulness|ousness|aliti|iviti|biliti|logi)$/,E=/^(.+?)(icate|ative|alize|iciti|ical|ful|ness)$/,_=/^(.+?)(al|ance|ence|er|ic|able|ible|ant|ement|ment|ent|ou|ism|ate|iti|ous|ive|ize)$/,F=/^(.+?)(s|t)(ion)$/,O=/^(.+?)e$/,P=/ll$/,N=new RegExp("^"+o+i+"[^aeiouwxy]$"),T=function(n){var i,o,r,s,a,h,l;if(n.length<3)return n;if(r=n.substr(0,1),"y"==r&&(n=r.toUpperCase()+n.substr(1)),s=p,a=m,s.test(n)?n=n.replace(s,"$1$2"):a.test(n)&&(n=n.replace(a,"$1$2")),s=v,a=y,s.test(n)){var T=s.exec(n);s=u,s.test(T[1])&&(s=g,n=n.replace(s,""))}else if(a.test(n)){var T=a.exec(n);i=T[1],a=d,a.test(i)&&(n=i,a=S,h=w,l=x,a.test(n)?n+="e":h.test(n)?(s=g,n=n.replace(s,"")):l.test(n)&&(n+="e"))}if(s=k,s.test(n)){var T=s.exec(n);i=T[1],n=i+"i"}if(s=b,s.test(n)){var T=s.exec(n);i=T[1],o=T[2],s=u,s.test(i)&&(n=i+t[o])}if(s=E,s.test(n)){var T=s.exec(n);i=T[1],o=T[2],s=u,s.test(i)&&(n=i+e[o])}if(s=_,a=F,s.test(n)){var T=s.exec(n);i=T[1],s=c,s.test(i)&&(n=i)}else if(a.test(n)){var T=a.exec(n);i=T[1]+T[2],a=c,a.test(i)&&(n=i)}if(s=O,s.test(n)){var T=s.exec(n);i=T[1],s=c,a=f,h=N,(s.test(i)||a.test(i)&&!h.test(i))&&(n=i)}return s=P,a=c,s.test(n)&&a.test(n)&&(s=g,n=n.replace(s,"")),"y"==r&&(n=r.toLowerCase()+n.substr(1)),n};return T}(),t.Pipeline.registerFunction(t.stemmer,"stemmer"),t.stopWordFilter=function(e){return e&&t.stopWordFilter.stopWords[e]!==e?e:void 0},t.stopWordFilter.stopWords={a:"a",able:"able",about:"about",across:"across",after:"after",all:"all",almost:"almost",also:"also",am:"am",among:"among",an:"an",and:"and",any:"any",are:"are",as:"as",at:"at",be:"be",because:"because",been:"been",but:"but",by:"by",can:"can",cannot:"cannot",could:"could",dear:"dear",did:"did","do":"do",does:"does",either:"either","else":"else",ever:"ever",every:"every","for":"for",from:"from",get:"get",got:"got",had:"had",has:"has",have:"have",he:"he",her:"her",hers:"hers",him:"him",his:"his",how:"how",however:"however",i:"i","if":"if","in":"in",into:"into",is:"is",it:"it",its:"its",just:"just",least:"least",let:"let",like:"like",likely:"likely",may:"may",me:"me",might:"might",most:"most",must:"must",my:"my",neither:"neither",no:"no",nor:"nor",not:"not",of:"of",off:"off",often:"often",on:"on",only:"only",or:"or",other:"other",our:"our",own:"own",rather:"rather",said:"said",say:"say",says:"says",she:"she",should:"should",since:"since",so:"so",some:"some",than:"than",that:"that",the:"the",their:"their",them:"them",then:"then",there:"there",these:"these",they:"they","this":"this",tis:"tis",to:"to",too:"too",twas:"twas",us:"us",wants:"wants",was:"was",we:"we",were:"were",what:"what",when:"when",where:"where",which:"which","while":"while",who:"who",whom:"whom",why:"why",will:"will","with":"with",would:"would",yet:"yet",you:"you",your:"your"},t.Pipeline.registerFunction(t.stopWordFilter,"stopWordFilter"),t.trimmer=function(t){var e=t.replace(/^\W+/,"").replace(/\W+$/,"");return""===e?void 0:e},t.Pipeline.registerFunction(t.trimmer,"trimmer"),t.TokenStore=function(){this.root={docs:{}},this.length=0},t.TokenStore.load=function(t){var e=new this;return e.root=t.root,e.length=t.length,e},t.TokenStore.prototype.add=function(t,e,n){var n=n||this.root,i=t[0],o=t.slice(1);return i in n||(n[i]={docs:{}}),0===o.length?(n[i].docs[e.ref]=e,void(this.length+=1)):this.add(o,e,n[i])},t.TokenStore.prototype.has=function(t){if(!t)return!1;for(var e=this.root,n=0;n<t.length;n++){if(!e[t[n]])return!1;e=e[t[n]]}return!0},t.TokenStore.prototype.getNode=function(t){if(!t)return{};for(var e=this.root,n=0;n<t.length;n++){if(!e[t[n]])return{};e=e[t[n]]}return e},t.TokenStore.prototype.get=function(t,e){return this.getNode(t,e).docs||{}},t.TokenStore.prototype.count=function(t,e){return Object.keys(this.get(t,e)).length},t.TokenStore.prototype.remove=function(t,e){if(t){for(var n=this.root,i=0;i<t.length;i++){if(!(t[i]in n))return;n=n[t[i]]}delete n.docs[e]}},t.TokenStore.prototype.expand=function(t,e){var n=this.getNode(t),i=n.docs||{},e=e||[];return Object.keys(i).length&&e.push(t),Object.keys(n).forEach(function(n){"docs"!==n&&e.concat(this.expand(t+n,e))},this),e},t.TokenStore.prototype.toJSON=function(){return{root:this.root,length:this.length}},function(t,e){"function"==typeof define&&define.amd?define(e):"object"==typeof exports?module.exports=e():t.lunr=e()}(this,function(){return t})}();
```
