## Supplementary material for "dsSurvival: Privacy preserving survival models for federated individual patient meta-analysis in DataSHIELD": Bookdown and documentation for using dsSurvival: plugin-bookdown.js

```
gitbook.require(["gitbook", "lodash", "jQuery"], function(gitbook, _, $) {

  var gs = gitbook.storage;

  gitbook.events.bind("start", function(e, config) {

    // add the Edit button (edit on Github)
    var edit = config.edit;
    if (edit && edit.link) gitbook.toolbar.createButton({
      icon: 'fa fa-edit',
      label: edit.text || 'Edit',
      position: 'left',
      onClick: function(e) {
        e.preventDefault();
        window.open(edit.link);
      }
    });

    // add the History button (file history on Github)
    var history = config.history;
    if (history && history.link) gitbook.toolbar.createButton({
      icon: 'fa fa-history',
      label: history.text || 'History',
      position: 'left',
      onClick: function(e) {
        e.preventDefault();
        window.open(history.link);
      }
    });

    // add the View button (file view on Github)
    var view = config.view;
    if (view && view.link) gitbook.toolbar.createButton({
      icon: 'fa fa-eye',
      label: view.text || 'View Source',
      position: 'left',
      onClick: function(e) {
        e.preventDefault();
        window.open(view.link);
      }
    });

    // add the Download button
    var down = config.download;
    var normalizeDownload = function() {
      if (!down || !(down instanceof Array) || down.length === 0) return;
      if (down[0] instanceof Array) return down;
      return $.map(down, function(file, i) {
        return [[file, file.replace(/.*[.]/g, '').toUpperCase()]];
      });
    };
    down = normalizeDownload(down);
    if (down) if (down.length === 1 && /[.]pdf$/.test(down[0][0])) {
      gitbook.toolbar.createButton({
        icon: 'fa fa-file-pdf-o',
        label: down[0][1],
        position: 'left',
        onClick: function(e) {
          e.preventDefault();
          window.open(down[0][0]);
        }
      });
    } else {
      gitbook.toolbar.createButton({
        icon: 'fa fa-download',
        label: 'Download',
        position: 'left',
        dropdown: $.map(down, function(item, i) {
          return {
            text: item[1],
            onClick: function(e) {
              e.preventDefault();
              window.open(item[0]);
            }
          };
        })
      });
    }

    // add the Information button
    var info = ['Keyboard shortcuts (<> indicates arrow keys):',
      '<left>/<right>: navigate to previous/next page',
      's: Toggle sidebar'];
    if (config.search !== false) info.push('f: Toggle search input ' +
      '(use <up>/<down>/Enter in the search input to navigate through search matches; ' +
      'press Esc to cancel search)');
    if (config.info !== false) gitbook.toolbar.createButton({
      icon: 'fa fa-info',
      label: 'Information about the toolbar',
      position: 'left',
      onClick: function(e) {
        e.preventDefault();
        window.alert(info.join('\n\n'));
      }
    });

    // highlight the current section in TOC
    var href = window.location.pathname;
    href = href.substr(href.lastIndexOf('/') + 1);
    // accentuated characters need to be decoded (#819)
    href = decodeURIComponent(href);
    if (href === '') href = 'index.html';
    var li = $('a[href^="' + href + location.hash + '"]').parent('li.chapter').first();
    var summary = $('ul.summary'), chaps = summary.find('li.chapter');
    if (li.length === 0) li = chaps.first();
    li.addClass('active');
    chaps.on('click', function(e) {
      chaps.removeClass('active');
      $(this).addClass('active');
      gs.set('tocScrollTop', summary.scrollTop());
    });

    var toc = config.toc;
    // collapse TOC items that are not for the current chapter
    if (toc && toc.collapse) (function() {
      var type = toc.collapse;
      if (type === 'none') return;
      if (type !== 'section' && type !== 'subsection') return;
      // sections under chapters
      var toc_sub = summary.children('li[data-level]').children('ul');
      if (type === 'section') {
        toc_sub.hide()
          .parent().has(li).children('ul').show();
      } else {
        toc_sub.children('li').children('ul').hide()
          .parent().has(li).children('ul').show();
      }
      li.children('ul').show();
      var toc_sub2 = toc_sub.children('li');
      if (type === 'section') toc_sub2.children('ul').hide();
      summary.children('li[data-level]').find('a')
        .on('click.bookdown', function(e) {
          if (href === $(this).attr('href').replace(/#.*/, ''))
            $(this).parent('li').children('ul').toggle();
        });
    })();

    // add tooltips to the <a>'s that are truncated
    $('a').each(function(i, el) {
      if (el.offsetWidth >= el.scrollWidth) return;
      if (typeof el.title === 'undefined') return;
      el.title = el.text;
    });

    // restore TOC scroll position
    var pos = gs.get('tocScrollTop');
    if (typeof pos !== 'undefined') summary.scrollTop(pos);

    // highlight the TOC item that has same text as the heading in view as scrolling
    if (toc && toc.scroll_highlight !== false && li.length > 0) (function() {
      // scroll the current TOC item into viewport
      var ht = $(window).height(), rect = li[0].getBoundingClientRect();
      if (rect.top >= ht || rect.top <= 0 || rect.bottom <= 0) {
        summary.scrollTop(li[0].offsetTop);
      }
      // current chapter TOC items
      var items = $('a[href^="' + href + '"]').parent('li.chapter'),
          m = items.length;
      if (m === 0) {
        items = summary.find('li.chapter');
        m = items.length;
      }
      if (m === 0) return;
      // all section titles on current page
      var hs = bookInner.find('.page-inner').find('h1,h2,h3'), n = hs.length,
          ts = hs.map(function(i, el) { return $(el).text(); });
      if (n === 0) return;
      var scrollHandler = function(e) {
        var ht = $(window).height();
        clearTimeout($.data(this, 'scrollTimer'));
        $.data(this, 'scrollTimer', setTimeout(function() {
          // find the first visible title in the viewport
          for (var i = 0; i < n; i++) {
            var rect = hs[i].getBoundingClientRect();
            if (rect.top >= 0 && rect.bottom <= ht) break;
          }
          if (i === n) return;
          items.removeClass('active');
          for (var j = 0; j < m; j++) {
            if (items.eq(j).children('a').first().text() === ts[i]) break;
          }
          if (j === m) j = 0;  // highlight the chapter title
          // search bottom-up for a visible TOC item to highlight; if an item is
          // hidden, we check if its parent is visible, and so on
          while (j > 0 && items.eq(j).is(':hidden')) j--;
          items.eq(j).addClass('active');
        }, 250));
      };
      bookInner.on('scroll.bookdown', scrollHandler);
      bookBody.on('scroll.bookdown', scrollHandler);
    })();

    // do not refresh the page if the TOC item points to the current page
    $('a[href="' + href + '"]').parent('li.chapter').children('a')
      .on('click', function(e) {
        bookInner.scrollTop(0);
        bookBody.scrollTop(0);
        return false;
      });

    var toolbar = config.toolbar;
    if (!toolbar || toolbar.position !== 'static') {
      var bookHeader = $('.book-header');
      bookBody.addClass('fixed');
      bookHeader.addClass('fixed')
      .css('background-color', bookBody.css('background-color'))
      .on('click.bookdown', function(e) {
        // the theme may have changed after user clicks the theme button
        bookHeader.css('background-color', bookBody.css('background-color'));
      });
    }

  });

  gitbook.events.bind("page.change", function(e) {
    // store TOC scroll position
    var summary = $('ul.summary');
    gs.set('tocScrollTop', summary.scrollTop());
  });

  var bookBody = $('.book-body'), bookInner = bookBody.find('.body-inner');
  var chapterTitle = function() {
    return bookInner.find('.page-inner').find('h1,h2').first().text();
  };
  var saveScrollPos = function(e) {
    // save scroll position before page is reloaded
    gs.set('bodyScrollTop', {
      body: bookBody.scrollTop(),
      inner: bookInner.scrollTop(),
      focused: document.hasFocus(),
      title: chapterTitle()
    });
  };
  $(document).on('servr:reload', saveScrollPos);

  // check if the page is loaded in an iframe (e.g. the RStudio preview window)
  var inIFrame = function() {
    var inIframe = true;
    try { inIframe = window.self !== window.top; } catch (e) {}
    return inIframe;
  };
  if (inIFrame()) {
    $(window).on('blur unload', saveScrollPos);
  }

  $(function(e) {
    var pos = gs.get('bodyScrollTop');
    if (pos) {
      if (pos.title === chapterTitle()) {
        if (pos.body !== 0) bookBody.scrollTop(pos.body);
        if (pos.inner !== 0) bookInner.scrollTop(pos.inner);
      }
    }
    if ((pos && pos.focused) || !inIFrame()) bookInner.find('.page-wrapper').focus();
    // clear book body scroll position
    gs.remove('bodyScrollTop');
  });

});
```
