## Supplementary material for "dsSurvival: Privacy preserving survival models for federated individual patient meta-analysis in DataSHIELD": Bookdown and documentation for using dsSurvival: plugin-clipboard.js

```
gitbook.require(["gitbook", "jQuery"], function(gitbook, $) {

  var copyButton = '<button type="button" class="copy-to-clipboard-button" title="Copy to clipboard" aria-label="Copy to clipboard"><i class="fa fa-copy"></i></button>';
  var clipboard;

  gitbook.events.bind("page.change", function() {

    if (!ClipboardJS.isSupported()) return;

    // the page.change event is thrown twice: before and after the page changes
    if (clipboard) {
      // clipboard is already defined
      // we can deduct that we are before page changes
      clipboard.destroy(); // destroy the previous events listeners
      clipboard = undefined; // reset the clipboard object
      return;
    }

    $(copyButton).prependTo("div.sourceCode");

    clipboard = new ClipboardJS(".copy-to-clipboard-button", {
      text: function(trigger) {
        return trigger.parentNode.textContent;
      }
    });

  });

});
```
