## Supplementary material for "dsSurvival: Privacy preserving survival models for federated individual patient meta-analysis in DataSHIELD": Bookdown and documentation for using dsSurvival: plugin-fontsettings.js

```
gitbook.require(["gitbook", "lodash", "jQuery"], function(gitbook, _, $) {
    var fontState;

    var THEMES = {
        "white": 0,
        "sepia": 1,
        "night": 2
    };

    var FAMILY = {
        "serif": 0,
        "sans": 1
    };

    // Save current font settings
    function saveFontSettings() {
        gitbook.storage.set("fontState", fontState);
        update();
    }

    // Increase font size
    function enlargeFontSize(e) {
        e.preventDefault();
        if (fontState.size >= 4) return;

        fontState.size++;
        saveFontSettings();
    };

    // Decrease font size
    function reduceFontSize(e) {
        e.preventDefault();
        if (fontState.size <= 0) return;

        fontState.size--;
        saveFontSettings();
    };

    // Change font family
    function changeFontFamily(index, e) {
        e.preventDefault();

        fontState.family = index;
        saveFontSettings();
    };

    // Change type of color
    function changeColorTheme(index, e) {
        e.preventDefault();

        var $book = $(".book");

        if (fontState.theme !== 0)
            $book.removeClass("color-theme-"+fontState.theme);

        fontState.theme = index;
        if (fontState.theme !== 0)
            $book.addClass("color-theme-"+fontState.theme);

        saveFontSettings();
    };

    function update() {
        var $book = gitbook.state.$book;

        $(".font-settings .font-family-list li").removeClass("active");
        $(".font-settings .font-family-list li:nth-child("+(fontState.family+1)+")").addClass("active");

        $book[0].className = $book[0].className.replace(/\bfont-\S+/g, '');
        $book.addClass("font-size-"+fontState.size);
        $book.addClass("font-family-"+fontState.family);

        if(fontState.theme !== 0) {
            $book[0].className = $book[0].className.replace(/\bcolor-theme-\S+/g, '');
            $book.addClass("color-theme-"+fontState.theme);
        }
    };

    function init(config) {
        var $bookBody, $book;

        //Find DOM elements.
        $book = gitbook.state.$book;
        $bookBody = $book.find(".book-body");

        // Instantiate font state object
        fontState = gitbook.storage.get("fontState", {
            size: config.size || 2,
            family: FAMILY[config.family || "sans"],
            theme: THEMES[config.theme || "white"]
        });

        update();
    };


    gitbook.events.bind("start", function(e, config) {
        var opts = config.fontsettings;
        if (!opts) return;
        
        // Create buttons in toolbar
        gitbook.toolbar.createButton({
            icon: 'fa fa-font',
            label: 'Font Settings',
            className: 'font-settings',
            dropdown: [
                [
                    {
                        text: 'A',
                        className: 'font-reduce',
                        onClick: reduceFontSize
                    },
                    {
                        text: 'A',
                        className: 'font-enlarge',
                        onClick: enlargeFontSize
                    }
                ],
                [
                    {
                        text: 'Serif',
                        onClick: _.partial(changeFontFamily, 0)
                    },
                    {
                        text: 'Sans',
                        onClick: _.partial(changeFontFamily, 1)
                    }
                ],
                [
                    {
                        text: 'White',
                        onClick: _.partial(changeColorTheme, 0)
                    },
                    {
                        text: 'Sepia',
                        onClick: _.partial(changeColorTheme, 1)
                    },
                    {
                        text: 'Night',
                        onClick: _.partial(changeColorTheme, 2)
                    }
                ]
            ]
        });


        // Init current settings
        init(opts);
    });
});
```
