## Supplementary material for "dsSurvival: Privacy preserving survival models for federated individual patient meta-analysis in DataSHIELD": Bookdown and documentation for using dsSurvival: plugin-search.js

```
gitbook.require(["gitbook", "lodash", "jQuery"], function(gitbook, _, $) {
    var index = null;
    var fuse = null;
    var _search = {engine: 'lunr', opts: {}};
    var $searchInput, $searchLabel, $searchForm;
    var $highlighted = [], hi, hiOpts = { className: 'search-highlight' };
    var collapse = false, toc_visible = [];

    function init(config) {
        // Instantiate search settings
        _search = gitbook.storage.get("search", {
            engine: config.search.engine || 'lunr',
            opts: config.search.options || {},
        });
    };

    // Save current search settings
    function saveSearchSettings() {
        gitbook.storage.set("search", _search);
    }

    // Use a specific index
    function loadIndex(data) {
        // [Yihui] In bookdown, I use a character matrix to store the chapter
        // content, and the index is dynamically built on the client side.
        // Gitbook prebuilds the index data instead: https://github.com/GitbookIO/plugin-search
        // We can certainly do that via R packages V8 and jsonlite, but let's
        // see how slow it really is before improving it. On the other hand,
        // lunr cannot handle non-English text very well, e.g. the default
        // tokenizer cannot deal with Chinese text, so we may want to replace
        // lunr with a dumb simple text matching approach.
        if (_search.engine === 'lunr') {
          index = lunr(function () {
            this.ref('url');
            this.field('title', { boost: 10 });
            this.field('body');
          });
          data.map(function(item) {
            index.add({
              url: item[0],
              title: item[1],
              body: item[2]
            });
          });
          return;
        }
        fuse = new Fuse(data.map((_data => {
            return {
                url: _data[0],
                title: _data[1],
                body: _data[2]
            };
        })), Object.assign(
            {
                includeScore: true,
                threshold: 0.1,
                ignoreLocation: true,
                keys: ["title", "body"]
            },
            _search.opts
        ));
    }

    // Fetch the search index
    function fetchIndex() {
        return $.getJSON(gitbook.state.basePath+"/search_index.json")
                .then(loadIndex);  // [Yihui] we need to use this object later
    }

    // Search for a term and return results
    function search(q) {
        let results = [];
        switch (_search.engine) {
            case 'fuse':
                if (!fuse) return;
                results = fuse.search(q).map(function(result) {
                    var parts = result.item.url.split('#');
                    return {
                        path: parts[0],
                        hash: parts[1]
                    };
                });
                break;
            case 'lunr':
            default:
                if (!index) return;
                results = _.chain(index.search(q)).map(function(result) {
                    var parts = result.ref.split("#");
                    return {
                        path: parts[0],
                        hash: parts[1]
                    };
                })
                .value();
        }

        // [Yihui] Highlight the search keyword on current page
        $highlighted = $('.page-inner')
          .unhighlight(hiOpts).highlight(q, hiOpts).find('span.search-highlight');
        scrollToHighlighted(0);

        return results;
    }

    // [Yihui] Scroll the chapter body to the i-th highlighted string
    function scrollToHighlighted(d) {
      var n = $highlighted.length;
      hi = hi === undefined ? 0 : hi + d;
      // navignate to the previous/next page in the search results if reached the top/bottom
      var b = hi < 0;
      if (d !== 0 && (b || hi >= n)) {
        var path = currentPath(), n2 = toc_visible.length;
        if (n2 === 0) return;
        for (var i = b ? 0 : n2; (b && i < n2) || (!b && i >= 0); i += b ? 1 : -1) {
          if (toc_visible.eq(i).data('path') === path) break;
        }
        i += b ? -1 : 1;
        if (i < 0) i = n2 - 1;
        if (i >= n2) i = 0;
        var lnk = toc_visible.eq(i).find('a[href$=".html"]');
        if (lnk.length) lnk[0].click();
        return;
      }
      if (n === 0) return;
      var $p = $highlighted.eq(hi);
      $p[0].scrollIntoView();
      $highlighted.css('background-color', '');
      // an orange background color on the current item and removed later
      $p.css('background-color', 'orange');
      setTimeout(function() {
        $p.css('background-color', '');
      }, 2000);
    }

    function currentPath() {
      var href = window.location.pathname;
      href = href.substr(href.lastIndexOf('/') + 1);
      return href === '' ? 'index.html' : href;
    }

    // Create search form
    function createForm(value) {
        if ($searchForm) $searchForm.remove();
        if ($searchLabel) $searchLabel.remove();
        if ($searchInput) $searchInput.remove();

        $searchForm = $('<div>', {
            'class': 'book-search',
            'role': 'search'
        });

        $searchLabel = $('<label>', {
            'for': 'search-box',
            'aria-hidden': 'false',
            'hidden': ''
        });

        $searchInput = $('<input>', {
            'id': 'search-box',
            'type': 'search',
            'class': 'form-control',
            'val': value,
            'placeholder': 'Type to search (Enter for navigation)',
            'title': 'Use Enter or the <Down> key to navigate to the next match, or the <Up> key to the previous match'
        });

        $searchLabel.append("Type to search");
        $searchLabel.appendTo($searchForm);
        $searchInput.appendTo($searchForm);
        $searchForm.prependTo(gitbook.state.$book.find('.book-summary'));
    }

    // Return true if search is open
    function isSearchOpen() {
        return gitbook.state.$book.hasClass("with-search");
    }

    // Toggle the search
    function toggleSearch(_state) {
        if (isSearchOpen() === _state) return;
        if (!$searchInput) return;

        gitbook.state.$book.toggleClass("with-search", _state);

        // If search bar is open: focus input
        if (isSearchOpen()) {
            gitbook.sidebar.toggle(true);
            $searchInput.focus();
        } else {
            $searchInput.blur();
            $searchInput.val("");
            gitbook.storage.remove("keyword");
            gitbook.sidebar.filter(null);
            $('.page-inner').unhighlight(hiOpts);
        }
    }

    function sidebarFilter(results) {
        gitbook.sidebar.filter(_.pluck(results, "path"));
        toc_visible = $('ul.summary').find('li:visible');
    }

    // Recover current search when page changed
    function recoverSearch() {
        var keyword = gitbook.storage.get("keyword", "");

        createForm(keyword);

        if (keyword.length > 0) {
            if(!isSearchOpen()) {
                toggleSearch(true); // [Yihui] open the search box
            }
            sidebarFilter(search(keyword));
        }
    }


    gitbook.events.bind("start", function(e, config) {
        // [Yihui] disable search
        if (config.search === false) return;
        init(config);
        collapse = !config.toc || config.toc.collapse === 'section' ||
          config.toc.collapse === 'subsection';

        // Pre-fetch search index and create the form
        fetchIndex()
        // [Yihui] recover search after the page is loaded
        .then(recoverSearch);


        // Type in search bar
        $(document).on("keyup", ".book-search input", function(e) {
            var key = (e.keyCode ? e.keyCode : e.which);
            // [Yihui] Escape -> close search box; Up/Down/Enter: previous/next highlighted
            if (key == 27) {
                e.preventDefault();
                toggleSearch(false);
            } else if (key == 38) {
              scrollToHighlighted(-1);
            } else if (key == 40 || key == 13) {
              scrollToHighlighted(1);
            }
        }).on("input", ".book-search input", function(e) {
            var q = $(this).val().trim();
            if (q.length === 0) {
                gitbook.sidebar.filter(null);
                gitbook.storage.remove("keyword");
                $('.page-inner').unhighlight(hiOpts);
            } else {
                var results = search(q);
                sidebarFilter(results);
                gitbook.storage.set("keyword", q);
            }
        });

        // Create the toggle search button
        gitbook.toolbar.createButton({
            icon: 'fa fa-search',
            label: 'Search',
            position: 'left',
            onClick: toggleSearch
        });

        // Bind keyboard to toggle search
        gitbook.keyboard.bind(['f'], toggleSearch);
    });

    // [Yihui] do not try to recover search; always start fresh
    // gitbook.events.bind("page.change", recoverSearch);
});
```
