## Supplementary material for "dsSurvival: Privacy preserving survival models for federated individual patient meta-analysis in DataSHIELD": Bookdown and documentation for using dsSurvival: plugin-sharing.js

```
gitbook.require(["gitbook", "lodash", "jQuery"], function(gitbook, _, $) {
    var SITES = {
        'github': {
            'label': 'Github',
            'icon': 'fa fa-github',
            'onClick': function(e) {
                e.preventDefault();
                var repo = $('meta[name="github-repo"]').attr('content');
                if (typeof repo === 'undefined') throw("Github repo not defined");
                window.open("https://github.com/"+repo);
            }
        },
        'facebook': {
            'label': 'Facebook',
            'icon': 'fa fa-facebook',
            'onClick': function(e) {
                e.preventDefault();
                window.open("http://www.facebook.com/sharer/sharer.php?u="+encodeURIComponent(location.href));
            }
        },
        'twitter': {
            'label': 'Twitter',
            'icon': 'fa fa-twitter',
            'onClick': function(e) {
                e.preventDefault();
                window.open("http://twitter.com/intent/tweet?text="+encodeURIComponent(document.title)+"&url="+encodeURIComponent(location.href)+"&hashtags=rmarkdown,bookdown");
            }
        },
        'linkedin': {
            'label': 'LinkedIn',
            'icon': 'fa fa-linkedin',
            'onClick': function(e) {
                e.preventDefault();
                window.open("https://www.linkedin.com/shareArticle?mini=true&url="+encodeURIComponent(location.href)+"&title="+encodeURIComponent(document.title));
            }
        },
        'weibo': {
            'label': 'Weibo',
            'icon': 'fa fa-weibo',
            'onClick': function(e) {
                e.preventDefault();
                window.open("http://service.weibo.com/share/share.php?content=utf-8&url="+encodeURIComponent(location.href)+"&title="+encodeURIComponent(document.title));
            }
        },
        'instapaper': {
            'label': 'Instapaper',
            'icon': 'fa fa-italic',
            'onClick': function(e) {
                e.preventDefault();
                window.open("http://www.instapaper.com/text?u="+encodeURIComponent(location.href));
            }
        },
        'vk': {
            'label': 'VK',
            'icon': 'fa fa-vk',
            'onClick': function(e) {
                e.preventDefault();
                window.open("http://vkontakte.ru/share.php?url="+encodeURIComponent(location.href));
            }
        },
        'whatsapp': {
            'label': 'Whatsapp',
            'icon': 'fa fa-whatsapp',
            'onClick': function(e) {
                e.preventDefault();
                var url = encodeURIComponent(location.href);
                window.open((isMobile() ? "whatsapp://send" : "https://web.whatsapp.com/send") + "?text=" + url);
            }
        },
    };

    function isMobile() {
      return !!navigator.maxTouchPoints;
    }

    gitbook.events.bind("start", function(e, config) {
        var opts = config.sharing;
        if (!opts) return;

        // Create dropdown menu
        var menu = _.chain(opts.all)
            .map(function(id) {
                var site = SITES[id];
                if (!site) return;
                return {
                    text: site.label,
                    onClick: site.onClick
                };
            })
            .compact()
            .value();

        // Create main button with dropdown
        if (menu.length > 0) {
            gitbook.toolbar.createButton({
                icon: 'fa fa-share-alt',
                label: 'Share',
                position: 'right',
                dropdown: [menu]
            });
        }

        // Direct actions to share
        _.each(SITES, function(site, sideId) {
            if (!opts[sideId]) return;

            gitbook.toolbar.createButton({
                icon: site.icon,
                label: site.label,
                title: site.label,
                position: 'right',
                onClick: site.onClick
            });
        });
    });
});
```
