## Supplementary material for "dsSurvival: Privacy preserving survival models for federated individual patient meta-analysis in DataSHIELD": Bookdown and documentation for using dsSurvival: header-attrs.js

```
// Pandoc 2.9 adds attributes on both header and div. We remove the former (to
// be compatible with the behavior of Pandoc < 2.8).
document.addEventListener('DOMContentLoaded', function(e) {
  var hs = document.querySelectorAll("div.section[class*='level'] > :first-child");
  var i, h, a;
  for (i = 0; i < hs.length; i++) {
    h = hs[i];
    if (!/^h[1-6]$/i.test(h.tagName)) continue;  // it should be a header h1-h6
    a = h.attributes;
    while (a.length > 0) h.removeAttribute(a[0].name);
  }
});
```
