## Supplementary material for "dsSurvival: Privacy preserving survival models for federated individual patient meta-analysis in DataSHIELD": Bookdown and documentation for using dsSurvival: forestplot_FINAL.R

```
########################################
# Better forest plot by Thodoris
########################################

########################
# load libraries
########################
library(knitr)
library(rmarkdown)
library(tinytex)
library(survival)
library(metafor)
library(meta)
library(ggplot2)
library(dsSurvivalClient)
require('DSI')
require('DSOpal')
require('dsBaseClient')


#######################
# connect to servers
#######################
builder <- DSI::newDSLoginBuilder()

builder$append(server="server1", url="http://192.168.56.100:8080/",
               user="administrator", password="datashield_test&", 
               table = "SURVIVAL.EXPAND_NO_MISSING1")

builder$append(server="server2", url="http://192.168.56.100:8080/",
               user="administrator", password="datashield_test&", 
               table = "SURVIVAL.EXPAND_NO_MISSING2")

builder$append(server="server3", url="http://192.168.56.100:8080/",
               user="administrator", password="datashield_test&", 
               table = "SURVIVAL.EXPAND_NO_MISSING3")

logindata <- builder$build()

connections <- DSI::datashield.login(logins = logindata, assign = TRUE, symbol = "D") 

ds.asNumeric(x.name = "D$cens",
             newobj = "EVENT",
             datasources = connections)

ds.asNumeric(x.name = "D$survtime",
             newobj = "SURVTIME",
             datasources = connections)

ds.asFactor(input.var.name = "D$time.id",
            newobj = "TID",
            datasources = connections)

ds.log(x = "D$survtime",
       newobj = "log.surv",
       datasources = connections)

ds.asNumeric(x.name = "D$starttime",
             newobj = "STARTTIME",
             datasources = connections)
ds.asNumeric(x.name = "D$endtime",
             newobj = "ENDTIME",
             datasources = connections)

dsSurvivalClient::ds.Surv(time='STARTTIME', time2='ENDTIME', 
                          event = 'EVENT', objectname='surv_object',
                          type='counting')

coxph_model_full <- dsSurvivalClient::ds.coxph.SLMA(formula = 'surv_object~D$age+D$female')

dsSurvivalClient::ds.coxph.SLMA(formula = 'survival::Surv(time=SURVTIME,event=EVENT)~D$age+D$female', 
                                dataName = 'D', 
                                datasources = connections)

coxph_model_strata <- dsSurvivalClient::ds.coxph.SLMA(formula = 'surv_object~D$age + 
                          survival::strata(D$female)')
summary(coxph_model_strata)

dsSurvivalClient::ds.coxphSummary(x = 'coxph_serverside')

input_logHR = c(coxph_model_full$server1$coefficients[1,2], 
                coxph_model_full$server2$coefficients[1,2], 
                coxph_model_full$server3$coefficients[1,2])

input_se    = c(coxph_model_full$server1$coefficients[1,3], 
                coxph_model_full$server2$coefficients[1,3], 
                coxph_model_full$server3$coefficients[1,3])

meta_model <- metafor::rma(input_logHR, sei = input_se, method = 'REML')

metafor::forest.rma(x = meta_model, digits = 4) 

studylabels <- c("Study 1","Study 2","Study 3")

#Variables should be formatted as a matrix
effects <- matrix(c(input_logHR
                    ,input_se)
                  ,dimnames = list(studylabels,c("logHR","SE"))
                  ,nrow=length(studylabels)
                  ,ncol=2)

mtotal <- metagen(TE=logHR, seTE=SE, data=as.data.frame(effects)
                  , method.tau = 'REML', prediction=T, hakn=F)

########################
# generate forest plot
########################
meta::forest(mtotal
             ,digits=4
             ,prediction=F
             ,leftlabs=c("Study","logHR","SE")
             ,xlim=c(1.03,1.05)
             )
```
