## Supplementary material for "dsSurvival: Privacy preserving survival models for federated individual patient meta-analysis in DataSHIELD": Bookdown and documentation for using dsSurvival: index.Rmd

````
--- 
title: "dsSurvival: Privacy preserving survival models in DataSHIELD"
author: "Soumya Banerjee and Tom R P Bishop"
date: "`r Sys.Date()`"
site: bookdown::bookdown_site
documentclass: book
bibliography: [book.bib, packages.bib]
biblio-style: apalike
link-citations: yes
description: "This is a bookdown demonstrating how to build privacy preserving survival models using dsSurvival in DataSHIELD. The output format for this example is bookdown::gitbook."
---

This is a bookdown with executable code demonstrating how to use the dsSurvival package to create privacy preserving survival models in DataSHIELD. dsSurvival builds privacy preserving survival models.


# Prerequisites

This is a bookdown with executable code demonstrating how to use the dsSurvival package to create privacy preserving survival models in DataSHIELD. dsSurvival builds privacy preserving survival models.

DataSHIELD is a platform for federated analysis of private data. DataSHIELD has a client-server architecture and this package has a client side and server side component.

The client side package is called dsSurvivalClient:

   https://github.com/neelsoumya/dsSurvivalClient

The server side package is called dsSurvival:

   https://github.com/neelsoumya/dsSurvival


Please install R. Then install R Studio 

   https://www.rstudio.com/products/rstudio/download/preview/


We assume that the reader is familiar with R and DataSHIELD.

We outline some of the other prerequisites that need to be installed below:


```{r eval=FALSE}


install.packages('devtools')

library(devtools)

devtools::install_github('neelsoumya/dsSurvivalClient')

devtools::install_github('datashield/dsBaseClient@6.1.1')

install.packages('rmarkdown')

install.packages('knitr')

install.packages('tinytex')

install.packages('metafor')

install.packages('DSOpal')

install.packages('DSI')

install.packages('opalr')


```

<!-- Remember each Rmd file contains one and only one chapter, and a chapter is defined by the first-level heading `#`. -->

<!-- To compile this example to PDF, you need XeLaTeX. You are recommended to install TinyTeX (which includes XeLaTeX <https://yihui.org/tinytex/>. -->

```{r include=FALSE}
# automatically create a bib database for R packages
knitr::write_bib(c(
  .packages(), 'bookdown', 'knitr', 'rmarkdown'
), 'packages.bib')
```
````
