## Supplementary material for "dsSurvival: Privacy preserving survival models for federated individual patient meta-analysis in DataSHIELD": Bookdown and documentation for using dsSurvival: installer_R.R

```
###############################################
# code to install all DataSHIELD R packages
# Usage:
#    R --no-save < installer_R.R
#    or
#    source('installer_R.R')
#
###############################################

install.packages('devtools')
library(devtools)
#devtools::install_github('https://github.com/datashield/dsBaseClient')
devtools::install_github('datashield/dsBaseClient@6.1.1')
library(dsBaseClient)
 
install.packages('DSOpal')
install.packages('DSI')
install.packages('opalr')


devtools::install_github('https://github.com/datashield/opal')
library(opal)

devtools::install_github('https://github.com/datashield/dsStatsClient')#, force = TRUE)
library(dsStatsClient)

devtools::install_github('https://github.com/datashield/dsGraphicsClient')#, force = TRUE)
library(dsGraphicsClient)

devtools::install_github('https://github.com/datashield/dsModellingClient')#, force = TRUE)

#library(dsModellingClient)
#devtools::install_github('https://github.com/datashield/DSOpal')#, force = TRUE)

install.packages('bookdown')
install.packages('rmarkdown')
install.packages('knitr')
install.packages('tinytex')
install.packages('metafor')

devtools::install_github('https://github.com/neelsoumya/dsSurvivalClient', 'privacy_survival_curves')#, force = TRUE)
library(dsSurvivalClient)
library(bookdown)
```
