## Supplementary material for "dsSurvival: Privacy preserving survival models for federated individual patient meta-analysis in DataSHIELD": Bookdown and documentation for using dsSurvival: packages.bib

@Manual{R-base,
  title = {R: A Language and Environment for Statistical Computing},
  author = {{R Core Team}},
  organization = {R Foundation for Statistical Computing},
  address = {Vienna, Austria},
  year = {2020},
  url = {https://www.R-project.org/},
}

@Manual{R-bookdown,
  title = {bookdown: Authoring Books and Technical Documents with R Markdown},
  author = {Yihui Xie},
  year = {2021},
  note = {R package version 0.24},
  url = {https://CRAN.R-project.org/package=bookdown},
}

@Manual{R-knitr,
  title = {knitr: A General-Purpose Package for Dynamic Report Generation in R},
  author = {Yihui Xie},
  year = {2021},
  note = {R package version 1.36},
  url = {https://yihui.org/knitr/},
}

@Manual{R-rmarkdown,
  title = {rmarkdown: Dynamic Documents for R},
  author = {JJ Allaire and Yihui Xie and Jonathan McPherson and Javier Luraschi and Kevin Ushey and Aron Atkins and Hadley Wickham and Joe Cheng and Winston Chang and Richard Iannone},
  year = {2021},
  note = {R package version 2.11},
  url = {https://CRAN.R-project.org/package=rmarkdown},
}

@Book{bookdown2016,
  title = {bookdown: Authoring Books and Technical Documents with {R} Markdown},
  author = {Yihui Xie},
  publisher = {Chapman and Hall/CRC},
  address = {Boca Raton, Florida},
  year = {2016},
  note = {ISBN 978-1138700109},
  url = {https://bookdown.org/yihui/bookdown},
}

@Book{knitr2015,
  title = {Dynamic Documents with {R} and knitr},
  author = {Yihui Xie},
  publisher = {Chapman and Hall/CRC},
  address = {Boca Raton, Florida},
  year = {2015},
  edition = {2nd},
  note = {ISBN 978-1498716963},
  url = {https://yihui.org/knitr/},
}

@InCollection{knitr2014,
  booktitle = {Implementing Reproducible Computational Research},
  editor = {Victoria Stodden and Friedrich Leisch and Roger D. Peng},
  title = {knitr: A Comprehensive Tool for Reproducible Research in {R}},
  author = {Yihui Xie},
  publisher = {Chapman and Hall/CRC},
  year = {2014},
  note = {ISBN 978-1466561595},
  url = {http://www.crcpress.com/product/isbn/9781466561595},
}

@Book{rmarkdown2018,
  title = {R Markdown: The Definitive Guide},
  author = {Yihui Xie and J.J. Allaire and Garrett Grolemund},
  publisher = {Chapman and Hall/CRC},
  address = {Boca Raton, Florida},
  year = {2018},
  note = {ISBN 9781138359338},
  url = {https://bookdown.org/yihui/rmarkdown},
}

@Book{rmarkdown2020,
  title = {R Markdown Cookbook},
  author = {Yihui Xie and Christophe Dervieux and Emily Riederer},
  publisher = {Chapman and Hall/CRC},
  address = {Boca Raton, Florida},
  year = {2020},
  note = {ISBN 9780367563837},
  url = {https://bookdown.org/yihui/rmarkdown-cookbook},
}


@article{Banerjee2022,
abstract = {Achieving sufficient statistical power in a survival analysis usually requires large amounts of data from different sites. Sensitivity of individual-level data, ethical and practical considerations regarding data sharing across institutions could be a potential challenge for achieving this added power. Hence we implemented a federated meta-analysis approach of survival models in DataSHIELD, where only anonymous aggregated data are shared across institutions, while simultaneously allowing for exploratory, interactive modelling. In this case, meta-analysis techniques to combine analysis results from each site are a solution, but a manual analysis workflow hinders exploration. Thus, the aim is to provide a framework for performing meta-analysis of Cox regression models across institutions without manual analysis steps for the data providers. We introduce a package (dsSurvival) which allows privacy preserving meta-analysis of survival models, including the calculation of hazard ratios. Our tool can be of great use in biomedical research where there is a need for building survival models and there are privacy concerns about sharing data. A tutorial in bookdown format with code, diagnostics, plots and synthetic data is available here: https://neelsoumya.github.io/dsSurvivalbookdown/ All code is available from the following repositories: https://github.com/neelsoumya/dsSurvivalClient/ https://github.com/neelsoumya/dsSurvival/ {\#}{\#}{\#} Competing Interest Statement The authors have declared no competing interest.},
author = {Banerjee, Soumya and Sofack, Ghislain and Papakonstantinou, Thodoris and Avraam, Demetris and Burton, Paul and Z{\"{o}}ller, Daniela and Bishop, Tom RP},
doi = {10.1101/2022.01.04.471418},
journal = {bioRxiv},
mendeley-groups = {cam{\_}project,My{\_}PAPERS},
month = {jan},
pages = {2022.01.04.471418},
publisher = {Cold Spring Harbor Laboratory},
title = {{dsSurvival: Privacy preserving survival models for federated individual patient meta-analysis in DataSHIELD}},
year = {2022}
}
