## Supplementary material for "dsSurvival: Privacy preserving survival models for federated individual patient meta-analysis in DataSHIELD": Bookdown and documentation for using dsSurvival: README.md

## Introduction

[![License](https://img.shields.io/badge/license-GPLv3-blue.svg)](https://www.gnu.org/licenses/gpl-3.0.html)

This is a bookdown with executable code demonstrating how to use the dsSurvival package to create privacy preserving survival models in DataSHIELD. dsSurvival builds privacy preserving survival models.

DataSHIELD is a platform for federated analysis of private data. This package can be used to build survival models, Cox proportional hazards models or Cox regression models.

The complete bookdown is available here:

https://neelsoumya.github.io/dsSurvival_bookdown


DataSHIELD has a client-server architecture and this package has a client side and server side component.

  * The server side package is called dsSurvival:
      
      https://github.com/neelsoumya/dsSurvival

  * The client side package is called dsSurvivalClient:
      
      https://github.com/neelsoumya/dsSurvivalClient


## Bookdown

The complete bookdown, tutorial, vignette with executable code and synthetic data is available here:

https://neelsoumya.github.io/dsSurvival_bookdown


## Quick start

Please install R and R Studio 

   https://www.rstudio.com/products/rstudio/download/preview/


Install the following packages:


```r 
install.packages('devtools')
library(devtools)
devtools::install_github('neelsoumya/dsSurvivalClient')
devtools::install_github('datashield/dsBaseClient@6.1.1')
install.packages('rmarkdown')
install.packages('knitr')
install.packages('tinytex')
install.packages('metafor')
install.packages('DSOpal')
install.packages('DSI')
install.packages('opalr')
```


Follow the tutorial in bookdown format with executable code:

https://neelsoumya.github.io/dsSurvival_bookdown/


## Full Installation


* Install R and R Studio 

* In R, install the following packages

```r 
  
  install.packages('devtools')
  library(devtools)
  devtools::install_github('neelsoumya/dsSurvivalClient')
  install.packages('bookdown')
  devtools::install_github('datashield/dsBaseClient@6.1.1')
  install.packages('rmarkdown')
  install.packages('knitr')
  install.packages('tinytex')
  install.packages('metafor')
  install.packages('DSOpal')
  install.packages('DSI')
  install.packages('opalr')
  
  ```

  or

  ```r 

  R --no-save < installer_R.R
  
  ```
  
  or
  
  run the following script in R `installer_R.R`
  
 
 
 Install R Studio and the development environment as described below:

    https://data2knowledge.atlassian.net/wiki/spaces/DSDEV/pages/12943461/Getting+started

 Install the virtual machines as described below:

    https://data2knowledge.atlassian.net/wiki/spaces/DSDEV/pages/931069953/Installation+Training+Hub-+DataSHIELD+v6

    https://data2knowledge.atlassian.net/wiki/spaces/DSDEV/pages/1657634881/Testing+100+VM

    https://data2knowledge.atlassian.net/wiki/spaces/DSDEV/pages/1657634898/Tutorial+6.1.0+100+VM

 Install dsBase and dsSurvival on Opal server in the Virtual Machine (type neelsoumya/dsSurvival and main in the textboxes)
  
 
  
  
## Usage

See the bookdown below for a complete tutorial:

https://neelsoumya.github.io/dsSurvival_bookdown


A minimal example of a book based on R Markdown and **bookdown** (https://github.com/rstudio/bookdown). Please see the page "Get Started" at https://bookdown.org/home/about/ for how to compile this example.

The bookdown can be compiled by typing the following commands:

  ```r 
  
  library(bookdown)

  bookdown::serve_book()
  
  ```


## Contact

Soumya Banerjee and Tom R.P. Bishop


## Citation

If you use this code, please cite the following DOI


[![DOI](https://zenodo.org/badge/DOI/10.5281/zenodo.4806588.svg)](https://doi.org/10.5281/zenodo.4806588)


and the following

dsSurvival: Privacy preserving survival models for federated individual patient meta-analysis in DataSHIELD

https://www.biorxiv.org/content/10.1101/2022.01.04.471418v1

https://doi.org/10.1101/2022.01.04.471418
