## Supplementary material for "dsSurvival: Privacy preserving survival models for federated individual patient meta-analysis in DataSHIELD": Bookdown and documentation for using dsSurvival: survival_bookdown.Rmd

````
--- 
title: "dsSurvival: Survival models in DataSHIELD"
author: "Soumya Banerjee and Tom Bishop"
date: "`r Sys.Date()`"
site: bookdown::bookdown_site
documentclass: book
bibliography: [book.bib, packages.bib]
biblio-style: apalike
link-citations: yes
description: "This is a minimal example of using the bookdown package to write a book. The output format for this example is bookdown::gitbook."
---

# Prerequisites

This is a _sample_ book written in **Markdown**. You can use anything that Pandoc's Markdown supports, e.g., a math equation $a^2 + b^2 = c^2$.

The **bookdown** package can be installed from CRAN or Github:

```{r eval=FALSE}
install.packages("bookdown")
# or the development version
# devtools::install_github("rstudio/bookdown")
```

Remember each Rmd file contains one and only one chapter, and a chapter is defined by the first-level heading `#`.

To compile this example to PDF, you need XeLaTeX. You are recommended to install TinyTeX (which includes XeLaTeX): <https://yihui.org/tinytex/>.

```{r include=FALSE}
# automatically create a bib database for R packages
knitr::write_bib(c(
  .packages(), 'bookdown', 'knitr', 'rmarkdown'
), 'packages.bib')
```

<!--chapter:end:index.Rmd-->

---
output:
  pdf_document: default
  html_document: default
---
# Introduction {#intro}

This is a document that outlines a vignette for implementing survival models
and meta-analyzing hazard ratios in the DataSHIELD platform.

You can label chapter and section titles using `{#label}` after them, e.g., we can reference Chapter \@ref(intro). If you do not manually label them, there will be automatic labels anyway, e.g., Chapter \@ref(methods).

Figures and tables with captions will be placed in `figure` and `table` environments, respectively.

```{r nice-fig, fig.cap='Here is a nice figure!', out.width='80%', fig.asp=.75, fig.align='center'}
par(mar = c(4, 4, .1, .1))
plot(pressure, type = 'b', pch = 19)
```

Reference a figure by its code chunk label with the `fig:` prefix, e.g., see Figure \@ref(fig:nice-fig). Similarly, you can reference tables generated from `knitr::kable()`, e.g., see Table \@ref(tab:nice-tab).

```{r nice-tab, tidy=FALSE}
knitr::kable(
  head(iris, 20), caption = 'Here is a nice table!',
  booktabs = TRUE
)
```

You can write citations, too. For example, we are using the **bookdown** package [@R-bookdown] in this sample book, which was built on top of R Markdown and **knitr** [@xie2015].


# Computational workflow

The computational steps are outlined below. 
The first step is connecting to the server and loading the survival data.
We assume that the reader is familiar with these details.

```
devtools::install_github('neelsoumya/dsSurvivalClient', 'privacy_survival_curves')

library(knitr)
library(rmarkdown)
library(tinytex)
library(survival)
library(metafor)
library(ggplot2)
library(survminer)
library(dsSurvivalClient)
require('DSI')
require('DSOpal')
require('dsBaseClient')
builder <- DSI::newDSLoginBuilder()

 builder$append(server="server1", url="https://opal-sandbox.mrc-epid.cam.ac.uk",
                user="dsuser", password="password", 
               table = "SURVIVAL.EXPAND_NO_MISSING1")
 builder$append(server="server2", url="https://opal-sandbox.mrc-epid.cam.ac.uk",
               user="dsuser", password="password", 
               table = "SURVIVAL.EXPAND_NO_MISSING2")
 builder$append(server="server3", url="https://opal-sandbox.mrc-epid.cam.ac.uk",
               user="dsuser", password="password", 
               table = "SURVIVAL.EXPAND_NO_MISSING3")	       

logindata <- builder$build()
connections <- DSI::datashield.login(logins = logindata, assign = TRUE, symbol = "D") 

```


```{r, include = FALSE}
####################
# Load library
####################
library(knitr)
library(rmarkdown)
library(tinytex)
library(survival)
library(metafor)
library(ggplot2)
# library(survminer)
#library(dsSurvival)
library(dsSurvivalClient)
require('DSI')
require('DSOpal')
require('dsBaseClient')
#######################
# Get data
#######################
builder <- DSI::newDSLoginBuilder()

 # builder$append(server="server1", url="https://opal-sandbox.mrc-epid.cam.ac.uk",
 #                user="dsuser", password="password", 
 #               table = "SURVIVAL.EXPAND_NO_MISSING1")
 # builder$append(server="server2", url="https://opal-sandbox.mrc-epid.cam.ac.uk",
 #               user="dsuser", password="password", 
 #               table = "SURVIVAL.EXPAND_NO_MISSING2")
 # builder$append(server="server3", url="https://opal-sandbox.mrc-epid.cam.ac.uk",
 #               user="dsuser", password="password", 
 #               table = "SURVIVAL.EXPAND_NO_MISSING3")	 
 

builder$append(server = "study1",
              url = "http://192.168.56.100:8080/",
              user = "administrator", password = "datashield_test&",
              table = "SURVIVAL.EXPAND_NO_MISSING1", driver = "OpalDriver")
builder$append(server = "study2",
              url = "http://192.168.56.100:8080/",
              user = "administrator", password = "datashield_test&",
              table = "SURVIVAL.EXPAND_NO_MISSING2", driver = "OpalDriver")
builder$append(server = "study3",
              url = "http://192.168.56.100:8080/",
              user = "administrator", password = "datashield_test&",
              table = "SURVIVAL.EXPAND_NO_MISSING3", driver = "OpalDriver")

logindata <- builder$build()

##############
# login
##############
# Log onto the remote Opal training servers
connections <- DSI::datashield.login(logins = logindata, assign = TRUE, symbol = "D") 

```


# Creating server-side variables for survival analysis

We now outline some steps for analysing survival data.

* make sure that the outcome variable is numeric 

```
ds.asNumeric(x.name = "D$cens",
             newobj = "EVENT",
             datasources = connections)
ds.asNumeric(x.name = "D$survtime",
             newobj = "SURVTIME",
             datasources = connections)
```

*  convert time id variable to a factor 

```
ds.asFactor(input.var.name = "D$time.id",
            newobj = "TID",
            datasources = connections)
```

*  create in the server-side the log(survtime) variable

```
ds.log(x = "D$survtime",
       newobj = "log.surv",
       datasources = connections)
```

*  create start time variable

```
ds.asNumeric(x.name = "D$starttime",
             newobj = "STARTTIME",
             datasources = connections)
ds.asNumeric(x.name = "D$endtime",
             newobj = "ENDTIME",
             datasources = connections)
```


```{r, include=FALSE}
# make sure that the outcome is numeric 
ds.asNumeric(x.name = "D$cens",
             newobj = "EVENT",
             datasources = connections)
ds.asNumeric(x.name = "D$survtime",
             newobj = "SURVTIME",
             datasources = connections)
# convert time id variable to a factor 
ds.asFactor(input.var.name = "D$time.id",
            newobj = "TID",
            datasources = connections)
# create in the server-side the log(survtime) variable
ds.log(x = "D$survtime",
       newobj = "log.surv",
       datasources = connections)
# create start time variable
ds.asNumeric(x.name = "D$starttime",
             newobj = "STARTTIME",
             datasources = connections)
# create end time variable
ds.asNumeric(x.name = "D$endtime",
             newobj = "ENDTIME",
             datasources = connections)
# check which variables exist
dsBaseClient::ds.ls()
# call new function modify NAMESPACE and DESCRIPTION
# call coxph server side
# client side function is here:
# https://github.com/neelsoumya/dsBaseClient/blob/absolute_newbie_client/R/ds.coxph.SLMA.R
# server side function is here:
# https://github.com/neelsoumya/dsBase/blob/absolute_newbie/R/coxphSLMADS.R
```


# Create survival object and call ds.coxph.SLMA()


* use constructed Surv object in *ds.coxph.SLMA()*

```
dsSurvivalClient::ds.Surv(time='STARTTIME', time2='ENDTIME', 
                      event = 'EVENT', objectname='surv_object',
                      type='counting')
coxph_model_full <- dsSurvivalClient::ds.coxph.SLMA(formula = 'surv_object~D$age+D$female')
```

* use direct inline call to *survival::Surv()*

```
dsSurvivalClient::ds.coxph.SLMA(formula = 'survival::Surv(time=SURVTIME,event=EVENT)~D$age+D$female', 
                                dataName = 'D', 
                                datasources = connections)
```

* call with *survival::strata()*

```
coxph_model_strata <- dsSurvivalClient::ds.coxph.SLMA(formula = 'surv_object~D$age + 
                          survival::strata(D$female)')
summary(coxph_model_strata)
```


```{r, include=FALSE}
# 1. use constructed surv object in coxph
dsSurvivalClient::ds.Surv(time='STARTTIME', time2='ENDTIME', event = 'EVENT', objectname='surv_object', type='counting')
coxph_model_full <- dsSurvivalClient::ds.coxph.SLMA(formula = 'surv_object~D$age+D$female')
# 2. use direct inline call
dsSurvivalClient::ds.coxph.SLMA(formula = 'survival::Surv(time=SURVTIME,event=EVENT)~D$age+D$female', dataName = 'D', datasources = connections)
# 3. Call with strata()
coxph_model_strata <- dsSurvivalClient::ds.coxph.SLMA(formula = 'surv_object~D$age+survival::strata(D$female)')
summary(coxph_model_strata)
```

# Summary of survival objects

We can also summarize a server-side object of type *survival::Surv()*
using a call to *ds.coxphSummary()*. This will provide a non-disclosive
summary of the server-side object. An example call is shown below:


```
dsSurvivalClient::ds.coxphSummary(x = 'coxph_serverside')
```


```{r, include=FALSE}
#################################
# summary of coxphSLMA
#################################
# TODO:
# dsBaseClient::ds.summary(x = 'surv_object')
# dsBaseClient::ds.class(x = 'surv_object')
# dsBaseClient::ds.mean(x='surv_object')
#################################
# TODO: Plot survival curves
#################################
# fit <- survival::survfit(formula = 'surv_object~D$age+D$female', data = 'D')
# need ds.survfit() and survfitDS()
# fit_model <- ds.survfit(coxph_model[1])
# plot(fit_model)
# TODO:
# plot(survfit_km, fun="cloglog")
# TODO: 
# ggplot like functionality see other functions
# In dsBaseClient::
# ds.survfit()
#         datashield.aggregate("survfitDS", ....)
#          return (the fit model)
# In dsBase::
# survfitDS(coxph_model)  
#               fit_model <- survival::survfit(coxph_model, newdata = 'D')
#               return (fit_model)
# TODO: dsSurvivalClient::ds.survfit(formula='surv_object~1', objectname='survfit_object')
# verify that object has been created
dsBaseClient::ds.ls()
# ERROR summary of survfit not allowed
# dsBaseClient::ds.summary(x='survfit_object')
# dsBaseClient::ds.mean(x='survfit_object')
```


# Diagnostics for Cox proportional hazards models

We have also created functions to test for the assumptions of 
Cox proportional hazards models.

```
dsSurvivalClient::ds.coxphSLMAassign(formula = 'surv_object~D$age+D$female',
                            objectname = 'coxph_serverside')
dsSurvivalClient::ds.cox.zphSLMA(fit = 'coxph_serverside')
dsSurvivalClient::ds.coxphSummary(x = 'coxph_serverside')
```

A diagnostic summary is shown below. 

```{r, echo=FALSE}
dsSurvivalClient::ds.coxphSLMAassign(formula = 'surv_object~D$age+D$female', objectname = 'coxph_serverside')
dsSurvivalClient::ds.cox.zphSLMA(fit = 'coxph_serverside')
dsSurvivalClient::ds.coxphSummary(x = 'coxph_serverside')
```


# Meta-analyze hazard ratios

We now outline how the hazard ratios from the survival models are meta-analyzed. 
We use the *metafor* package for meta-analysis. We show the summary of an 
example meta-analysis and a forest plot below. The forest plot shows a basic
example of meta-analyzed hazard ratios from a survival model (analyzed in
dsSurvivalClient).

The log-hazard ratios and their standard errors from each study can be
found after running *ds.coxphSLMA()*

The hazard ratios can then be meta-analyzed:

```
input_logHR = c(coxph_model_full$study1$coefficients[1,2], 
		coxph_model_full$study2$coefficients[1,2], 
		coxph_model_full$study3$coefficients[1,2])
input_se    = c(coxph_model_full$study1$coefficients[1,3], 
		coxph_model_full$study2$coefficients[1,3], 
		coxph_model_full$study3$coefficients[1,3])
metafor::rma(log_hazard_ratio, sei = se_hazard_ratio, method = 'REML')
```

A summary of this meta-analyzed model is shown below.


```{r, echo=FALSE}

# list of hazard ratios for first parameter (age) over 3 studies 
input_logHR = c(coxph_model_full$study1$coefficients[1,2], coxph_model_full$study2$coefficients[1,2], coxph_model_full$study3$coefficients[1,2])
input_se    = c(coxph_model_full$study1$coefficients[1,3], coxph_model_full$study2$coefficients[1,3], coxph_model_full$study3$coefficients[1,3])
meta_model <- metafor::rma(input_logHR, sei = input_se, method = 'REML')
summary(meta_model)
#######################################################
# forest plots of final meta-analyzed hazard ratios
#######################################################
```

We now show a forest plot with the meta-analyzed hazard ratios. The hazard ratios come from the dsSurvivalClient function *ds.coxphSLMA()*. The hazard ratios are meta-analyzed using the *metafor* package.

```{r fig.cap="Example forest plot of meta-analyzed hazard ratios.", echo=FALSE}

metafor::forest.rma(x = meta_model)

```    


\newpage
   
```{r, include=FALSE}

#############################################
# disconnect
#############################################
DSI::datashield.logout(conns = connections)

```


* https://github.com/datashield
  
  * http://www.metafor-project.org
  
  * https://github.com/neelsoumya/dsBase

  * https://github.com/neelsoumya/dsBaseClient

  * https://github.com/neelsoumya/dsSurvival

  * https://github.com/neelsoumya/dsSurvivalClient
  
  * https://github.com/neelsoumya/datashield_testing_basic
  

<!--chapter:end:01-intro.Rmd-->


<!--chapter:end:03-method.Rmd-->


<!--chapter:end:04-application.Rmd-->

# Summary

This bookdown demonstrates how to build privacy poreserving survival models using dsSurvival in DataSHIELD.

<!--chapter:end:05-summary.Rmd-->

`r if (knitr::is_html_output()) '
# References {-}
'`

<!--chapter:end:06-references.Rmd-->
````
