## Supplementary material for "dsSurvival: Privacy preserving survival models for federated individual patient meta-analysis in DataSHIELD": Vignette and bookdown with executable code and synthetic data that requires only client side package installation

*2022-01-07*

### Chapter 1 Prerequisites

This is a bookdown with executable code demonstrating how to use the dsSurvival package to create privacy preserving survival models in DataSHIELD. dsSurvival builds privacy preserving survival models.

DataSHIELD is a platform for federated analysis of private data. DataSHIELD has a client-server architecture and this package has a client side and server side component.

The client side package is called dsSurvivalClient:

https://github.com/neelsoumya/dsSurvivalClient

The server side package is called dsSurvival:

https://github.com/neelsoumya/dsSurvival

Please install R. Then install R Studio

https://www.rstudio.com/products/rstudio/download/preview/

We assume that the reader is familiar with R and DataSHIELD.

We outline some of the other prerequisites that need to be installed below:

```r
install.packages('devtools')

library(devtools)

devtools::install_github('neelsoumya/dsSurvivalClient')

devtools::install_github('datashield/dsBaseClient@6.1.1')

install.packages('rmarkdown')

install.packages('knitr')

install.packages('tinytex')

install.packages('metafor')

install.packages('DSOpal')

install.packages('DSI')

install.packages('opalr')
```

```
## surv_object~D$age+D$female
```

```
##
   [------------------------------------------------------------------------------
  Checking study1 (coxph_serverside <- coxphSLMAassignDS(formula = surv_object ~ D$age -
  Checking study2 (coxph_serverside <- coxphSLMAassignDS(formula = surv_object ~ D$age -
  Checking study3 (coxph_serverside <- coxphSLMAassignDS(formula = surv_object ~ D$age -
  Waiting...  (coxph_serverside <- coxphSLMAassignDS(formula = surv_object ~ D$age + D$f
  Checking study1 (coxph_serverside <- coxphSLMAassignDS(formula = surv_object ~ D$age -
  Checking study2 (coxph_serverside <- coxphSLMAassignDS(formula = surv_object ~ D$age -
  Checking study3 (coxph_serverside <- coxphSLMAassignDS(formula = surv_object ~ D$age -
  Waiting...  (coxph_serverside <- coxphSLMAassignDS(formula = surv_object ~ D$age + D$f
  Checking study1 (coxph_serverside <- coxphSLMAassignDS(formula = surv_object ~ D$age -
  Finalizing assignment study1 (coxph_serverside <- coxphSLMAassignDS(formula = surv_obj
  Checking study2 (coxph_serverside <- coxphSLMAassignDS(formula = surv_object ~ D$age -
  Finalizing assignment study2 (coxph_serverside <- coxphSLMAassignDS(formula = surv_obj
  Checking study3 (coxph_serverside <- coxphSLMAassignDS(formula = surv_object ~ D$age -
  Finalizing assignment study3 (coxph_serverside <- coxphSLMAassignDS(formula = surv_obj
  Assigned expr. (coxph_serverside <- coxphSLMAassignDS(formula = surv_object ~ D$age +
```

```
## NULL
```

```
##
   [-------------------------------------------------------------------------------
  Checking study1 (cox.zphSLMADS("coxph_serverside", "km", TRUE, FALSE, TRUE)) [-------
  Checking study2 (cox.zphSLMADS("coxph_serverside", "km", TRUE, FALSE, TRUE)) [-------
  Checking study3 (cox.zphSLMADS("coxph_serverside", "km", TRUE, FALSE, TRUE)) [-------
  Waiting...  (cox.zphSLMADS("coxph_serverside", "km", TRUE, FALSE, TRUE)) [-----------
  Checking study1 (cox.zphSLMADS("coxph_serverside", "km", TRUE, FALSE, TRUE)) [-------
  Checking study2 (cox.zphSLMADS("coxph_serverside", "km", TRUE, FALSE, TRUE)) [-------
  Checking study3 (cox.zphSLMADS("coxph_serverside", "km", TRUE, FALSE, TRUE)) [-------
  Waiting...  (cox.zphSLMADS("coxph_serverside", "km", TRUE, FALSE, TRUE)) [-----------
  Checking study1 (cox.zphSLMADS("coxph_serverside", "km", TRUE, FALSE, TRUE)) [-------
  Getting aggregate study1 (cox.zphSLMADS("coxph_serverside", "km", TRUE, FALSE, TRUE))
  Checking study2 (cox.zphSLMADS("coxph_serverside", "km", TRUE, FALSE, TRUE)) [=>-----
  Getting aggregate study2 (cox.zphSLMADS("coxph_serverside", "km", TRUE, FALSE, TRUE))
  Checking study3 (cox.zphSLMADS("coxph_serverside", "km", TRUE, FALSE, TRUE)) [===>---
  Getting aggregate study3 (cox.zphSLMADS("coxph_serverside", "km", TRUE, FALSE, TRUE))
  Aggregated (cox.zphSLMADS("coxph_serverside", "km", TRUE, FALSE, TRUE)) [============
```

```
## $study1
##          chisq df    p
## D$age    1.022  1 0.31
## D$female 0.364  1 0.55
## GLOBAL   1.239  2 0.54
##
## $study2
##             chisq df    p
## D$age    -254.681  1 1.00
## D$female    0.991  1 0.32
## GLOBAL   -222.531  2 1.00
##
## $study3
##          chisq df       p
## D$age    15.27  1 9.3e-05
## D$female  8.04  1  0.0046
## GLOBAL   23.31  2 8.7e-06
```

```
##
   [-----------------------------------------------------------------------------------
  Checking study1 (coxphSummaryDS("coxph_serverside")) [-------------------------------
  Checking study2 (coxphSummaryDS("coxph_serverside")) [-------------------------------
  Checking study3 (coxphSummaryDS("coxph_serverside")) [-------------------------------
  Waiting...  (coxphSummaryDS("coxph_serverside")) [-----------------------------------
  Checking study1 (coxphSummaryDS("coxph_serverside")) [-------------------------------
  Checking study2 (coxphSummaryDS("coxph_serverside")) [-------------------------------
  Checking study3 (coxphSummaryDS("coxph_serverside")) [-------------------------------
  Waiting...  (coxphSummaryDS("coxph_serverside")) [-----------------------------------
  Checking study1 (coxphSummaryDS("coxph_serverside")) [-------------------------------
  Getting aggregate study1 (coxphSummaryDS("coxph_serverside")) [=====>----------------
  Checking study2 (coxphSummaryDS("coxph_serverside")) [=======>-----------------------
  Getting aggregate study2 (coxphSummaryDS("coxph_serverside")) [===========>----------
  Checking study3 (coxphSummaryDS("coxph_serverside")) [===============>---------------
  Getting aggregate study3 (coxphSummaryDS("coxph_serverside")) [================>----
  Aggregated (coxphSummaryDS("coxph_serverside")) [=====================================
```

```
metafor::forest.rma(x = meta_model, digits = 4)
```

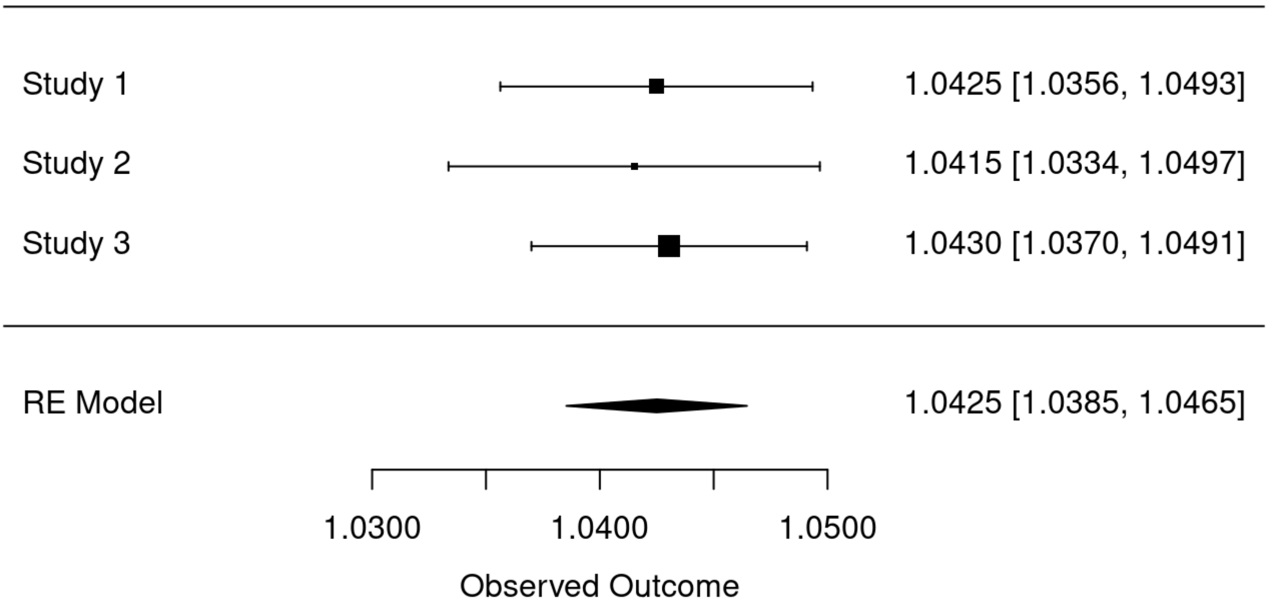


Figure 3.1: Example forest plot of meta-analyzed hazard ratios.

Finally, once you have finished your analysis, you can disconnect from the server(s) using the following command:

```
DSI::datashield.logout(conns = connections)
```

- https://github.com/datashield
- http://www.metafor-project.org
- https://github.com/neelsoumya/dsSurvival
- https://github.com/neelsoumya/dsSurvivalClient

### Chapter 4 Summary

This bookdown shows how to build privacy preserving survival models using dsSurvival in DataSHIELD. You can read more at:

- https://github.com/neelsoumya/dsSurvivalClient
- https://github.com/neelsoumya/dsSurvival
- https://neelsoumya.github.io/dsSurvival_bookdown/
- https://github.com/datashield
